# Genetic regulation of anthocyanin biosynthesis in *Cornus* species: The roles of R2R3-MYB transcription factors

**DOI:** 10.1101/2025.06.24.661385

**Authors:** Žaklina Pavlović, Miriam Payá-Milans, Marzena Nowakowska, Matthew L. Huff, Kimberly D. Gwinn, Robert N. Trigiano, Marcin Nowicki

## Abstract

Flowering dogwood (*Cornus florida* L.) and Asian dogwood (*C. kousa* F. Buerger ex Hance) are popular deciduous ornamental trees native to a wide range of the eastern and southeastern United States and East Asia, respectively. Anthocyanin pigments enhance desirable pink or dark red colored bracts in dogwoods. Although anthocyanin biosynthesis is one of the best-studied biological processes in nature, genomic and genetic resources to understand the molecular regulation of its synthesis in dogwoods are still lacking. Two classes of genes control anthocyanin production; both structural genes and MYB transcription factors may function as positive or negative regulators of anthocyanin biosynthesis. To reveal the molecular mechanisms that govern color production in ornamental dogwoods, mature bracts of three cultivars of *C. florida* (white bracts: ‘Cloud Nine’; red bracts: ‘Cherokee Brave’ and ‘Cherokee Chief’) and two cultivars of *C. kousa* (light green bracts: ‘Greensleeves’ and mid-tone pink bracts ‘Rosy Teacups’) were sampled when color was maximally visible. Differential gene expression analysis of the RNAseq data identified 1,156 differentially expressed genes in *C. florida* and 1,396 in *C. kousa*. Phylogenetic analysis with functional orthologues in other plants grouped the candidate R2R3-MYB identified in this study into two distinct subgroups. *CfMYB2*, *CfMYB3*, and *CkMYB*2 belonged to subgroup 4, whereas *CfMYB1 80* and *CkMYB1* clustered in subgroup 5. The former repress anthocyanin and proanthocyanidin synthesis in flowering and Asian dogwoods, whereas the latter increase it. Our study contributes to understanding processes behind anthocyanin production and lays foundation to future development of molecular markers for faster development of desirable red-bracted dogwoods.

## INTRODUCTION

Flowering dogwood (*Cornus florida* L.) and Asian dogwood (*C. kousa* F. Buerger ex Hance) are popular understory and deciduous ornamental trees of the *Cornus* genus, which includes about 60 species of trees and shrubs mostly found throughout the temperate regions of North America, Europe, and Asia (Eyde 1987, Cappiello and Shadow 2005, Wadl, Windham et al. 2014). Native to a wide range of states in the eastern and southeastern United States (US), flowering dogwoods owe their exceptional appeal to showy snow-white, pink, or red bracts, red fall foliage, and drupes (Witte, Windham et al. 2000, Cappiello and Shadow 2005). Asian dogwoods also are valued for their spring display of white, pink, or red bracts, brilliant-red fall foliage, colorful fruits, and exfoliating bark, and were introduced from Eastern Asia to the US because of their generally high resistance to insects and pathogens and overall better stress tolerance compared to flowering dogwood (Ranney, Grand and Knighten 1995, Li, Mmbaga et al. 2009, Wadl, Windham et al. 2014).

The majority of plant pigments are flavonoids, which are ubiquitous across the plant kingdom, and typically show functions in insect pollination and seed dispersal, hormone signaling, and abiotic and biotic defense (Liu, Osbourn and Ma 2015, Cao, Li et al. 2020, Wu, Wen et al. 2022). The desirable pink or dark red colors in bracts of flowering and Asian dogwoods are enhanced by biosynthesis of anthocyanin pigments (Kabir, Bautista et al. 2022). Two classes of genes control biosynthesis of anthocyanins and other phenylpropanoid-derived compounds: (1) structural genes encoding enzymes participating in formation of anthocyanins and (2) regulatory genes networks consisting of three types of transcription factors (TFs): v-myb avian myeloblastosis viral oncogene homolog proteins (MYBs), basic helix-loop-helix proteins (bHLHs), and WD repeat proteins (WDRs) (Allan, Hellens and Laing 2008, Dubos, Stracke et al. 2010, Zhang, Butelli and Martin 2014).

Anthocyanin profiles in both dogwood species are similar, with major fruit anthocyanins as cyanidin 3-O-galactoside in *C. florida* and cyanidin 3-O-glucoside in *C. kousa*, respectively (Du, Wang and Francis 1974, Eyde 1987, Vareed, Reddy et al. 2006, Lev-Yadun and Gould 2009). Anthocyanins are specialized flavonoid metabolites found in plants and play an important role in conferring defense against plant pathogens and abiotic stresses such as UV radiation, drought, or cold (Jin, Cominelli et al. 2000, Lev-Yadun and Gould 2009, Zhang, Wang et al. 2019, Gong, Li et al. 2020, Wajahat 2023). Moreover, their impact on plant crop quality contributes to the horticultural, industrial, and nutritional value of various plant products, such as apple (*Malus domestica* Borkh.) (Lin-Wang, Bolitho et al. 2010, Meng, Zhang et al. 2016), chrysanthemum (*Chrysanthemum morifolium* (Ramat.) Hemst.) (Liu, Xiang et al. 2015), eggplant (*Solanum melongena* L.) (Zhang, Hu et al. 2014), grape (*Vitis vinifera* L.) (Czemmel, Heppel and Bogs 2012), morning glory (*Ipomea purporea* L.) (Park, Ishikawa et al. 2007), petunia (*Petunia hybrida* D. Don ex W. H. Baxter) (Park, Ishikawa et al. 2007, Albert, Lewis et al. 2009, Albert, Lewis et al. 2011), royal lily (*Lilium regale* E. H. Wilson) (Yamagishi 2011), snapdragon (*Antirrhinum majus* L.) (Naing, Ai et al. 2018), or strawberry (*Fragaria vesca* L.) (Aharoni, De Vos et al. 2001), among many others. Their antioxidant activity has recently evoked tractable attention in protection against cancer, strokes, and other chronic human disorders (Zhang, Butelli and Martin 2014, Kodama, Brinch-Pedersen et al. 2018). Anthocyanins produced in the leaves of *C. kousa* were found to have anti-obesity effects (Khan, Shin et al. 2018).

Although flavonoid pigment biosynthesis is one of the best-studied biological processes in nature and R2R3-MYBs are widely distributed in the plant kingdom (Pei, Huang et al. 2024), key players involved in phenylpropanoid pathway and anthocyanin regulation in flowering and Asian dogwoods have not yet been identified. This means that the regulatory mechanisms of anthocyanin biosynthesis in *Cornus* species remain poorly understood. That knowledge gap delays the development of functional markers or the application of marker-assisted selection in dogwood breeding. We hypothesize that *Cornus* species have evolved divergent MYB regulatory networks to achieve convergent red bract phenotypes, thus providing a model for studying evolutionary plasticity in flavonoid pathways. Therefore, this work aimed to investigate the global transcription profiles of MYB TFs and structural genes using the RNA sequencing (RNAseq) approach within various cultivars to identify candidate R2R3-MYB transcription factors that are either positive or negative regulators of the anthocyanin pathway and other flavonoid compounds.

## MATERIAL AND METHODS

### Plant Materials

Four- week -old bracts were collected and immediately frozen in liquid nitrogen. Bracts from three cultivars of *C. florida* (white bracts: ‘Cloud Nine’; red bracts: ‘Cherokee Brave’ and ‘Cherokee Chief’) and two cultivars of *C. kousa* (light green bracts: ‘Greensleeves,’ and light to medium pink ‘Rosy Teacups’) were harvested when color was maximally visible (April and May 2017) (Figure 1A). Trees were grown at the University of Tennessee Knoxville, USA greenhouse and were about 5 years old and disease-free.

**Figure 1.**
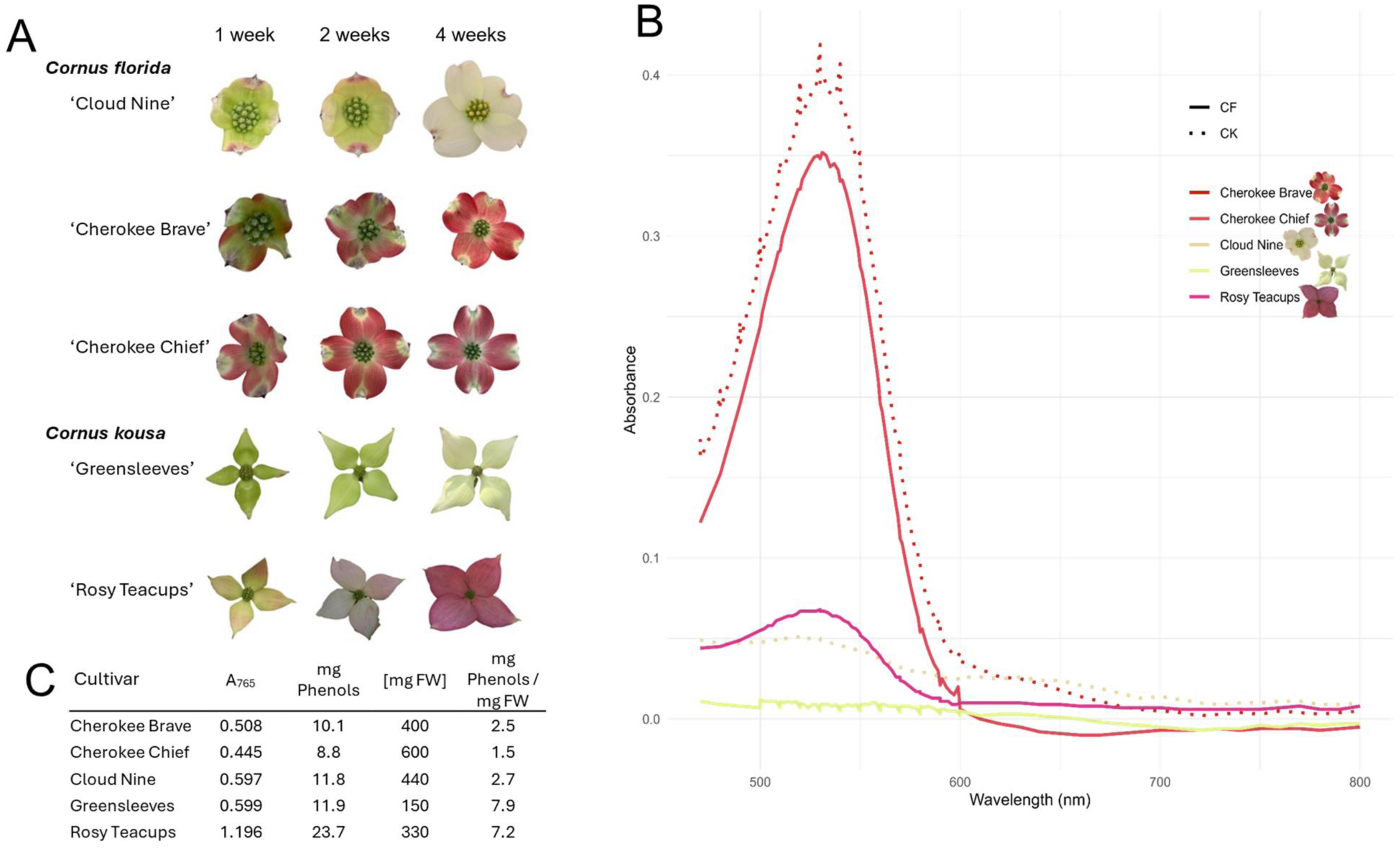
Phenotypes of *Cornus florida* and *C*. *kousa* bracts. (**A**) Representative bract phenotypes at 1, 2, and 4 weeks post-emergence for five cultivars: *C. florida* ‘Cloud Nine’, ‘Cherokee Brave’, ‘Cherokee Chief’; *C. kousa* ‘Greensleeves’, and ‘Rosy Teacups’. (**B**) Absorbance spectra (400–to–800 nm) of anthocyanin extracts from 4-week-old bracts. (**C**) Total phenolic content measured by Folin-Ciocalteu assay at 765 nm. Values are expressed as mg gallic acid equivalents (GAE) per mg fresh weight (FW).

### Anthocyanin isolation and spectral analyses

About 2 g of frozen samples per cultivar were homogenized in liquid nitrogen, and acetone extraction was performed as follows. Frozen tissue was ground to a fine powder. Two sequential extractions with ice-cold acetone (1:1 w:v) were performed, followed by centrifugation (21,000 × *g*, 4°C). Supernatants were pooled and re-extracted with 70% acetone containing 0.01% HCl (v/v). Combined acetone extracts were mixed with chloroform (1:1 v/v), vortexed, and centrifuged (21,000 × *g*, 4°C). The aqueous phase was collected and concentrated using a rotary evaporator (35 °C, 5 min under vacuum with 4 °C cooling). Final extracts were stored at −80 °C. Absorbance spectra (400 to 800 nm) were collected using Shimadzu UV-VIS Spectrophotometer UB-1900i (Shimadzu Scientific Instruments Inc., Coulmabia, MD). Total phenolic content was quantified using the Folin-Ciocalteu method (Folin and Ciocalteu 1927). Gallic acid (50–500 mg/L in H₂O) served as the reference. Reactions included 20 µL sample or standard and 100 µL Folin-Ciocalteu reagent with 300 µL 7.5% Na₂CO₃ (w/v). Absorbance was measured at 755 nm and 765 nm (validation step; primary quantification at 765 nm). Phenolic content was expressed as gallic acid equivalents (GAE) using the standard curve (R² = 0.9792 at 765 nm).

### RNA Extraction and Sequencing

RNA was extracted using 500 mg of frozen tissue per sample with the GeneAll® Ribospin™ Plant Total RNA Kit (GeneAll Biotechnology, Seoul, Korea), following the manufacturer’s instructions. Bract samples were homogenized using zirconia beads at 6 m/s for 30 sec in a Bead mill 24 (Fisher Scientific, Waltham, MA). Subsequently, total RNA integrity was assessed with Agilent 2100 Bioanalyzer system using the RNA 6000 Nano LabChip® kit (Agilent Technologies, Palo Alto, CA). Four of the 20 isolated RNA samples were below the required quality threshold (RIN >=6.0). The 16 RNA samples from three cultivars of flowering dogwood and two of Asian dogwood were submitted to GeneWiz (South Plainfield, NJ) for library preparation and sequencing using Illumina HiSeq 2500, which generated 150 bp paired end reads. All analyzed cultivars had at least two biological replicates.

### Quality Control

RNAseq yielded between 20 and 37 million reads per library, which were subjected to error correction using Rcorrector (RNAseq error CORRECTOR) (Song and Florea 2015). Read quality control was performed using FastQC v0.11.4 (Andrews, Krueger et al. 2010). Adaptors, short (<30 bases), and low-quality reads were trimmed using SKEWER v0.2.2 (Jiang, Lei et al. 2014). Subsequently, read quality control was reassessed using FastQC v0.11.4 (Andrews, Krueger et al. 2010) (Supplementary Table T1).

### Read Alignment and Gene Quantification

Reference genomes for flowering and Asian dogwood were not available at this study’s onset. Therefore, trimmed reads were mapped to the *C. florida* or *C. kousa* transcriptomes (Yu, Xiang et al. 2017) using Genomic short-read Nucleotide Alignment Program (GSNAP version 2018-01-31) (Wu, Reeder et al. 2016). Aligned reads were used to generate a gene-specific count matrix across samples using a standalone script provided by HTSeq (Anders, Pyl and Huber 2015), which counts the number of reads mapped to each gene.

### Differential Expression Analysis and Functional Enrichment

To identify R2R3-MYB TFs and structural genes involved in anthocyanin biosynthesis, gene counts were analyzed for gene-level differential expression using DESeq2 (Love, Huber and Anders 2014). This package provides an option to test differential expression by shrinking estimators of the negative binomial distribution. Count results from HTSeq were imported with an underlying experimental design that modeled the bract color as the effect of treatment while adjusting for cultivar or species as factors. Genes with less than one count across samples were removed. Regularized log transformation function was used to calculate sample distances using t-SNE plot functionality in the R package *tsne* (Donaldson and Donaldson 2010). The Wald test (Ghosh 1992) was employed to identify differentially expressed genes (DEGs) between two sample classes, red-bracted and non-pigmented for either species. Significant DEGs were identified with adjusted p-value < 0.05 after multiple testing using the Benjamini-Hochberg method (Benjamini and Hochberg 1995). Heatmaps with the expression profiles of DEGs after mapping to each of the two *Cornus* species transcriptomes were generated using the R package *pheatmap* with correlation distances (Pearson 1895). TransDecoder v3.0.0 was used to predict encoded peptides based on *C. florida* and *C. kousa* transcriptomes. TransDecoder identified all open reading frames (ORFs) that were at least 50 amino acids long in the transcripts. These ORFs were then scanned for homology to known proteins using blastp (Altschul, Gish et al. 1990) against the UniProtKB/TrEMBL database (Boutet, Lieberherr et al. 2007) for plants and searched against Pfam (Finn, Coggill et al. 2016) for conserved protein domains with HMMER v3.1b2 (Potter, Luciani et al. 2018). TransDecoder was rerun with the outputs generated from blastp and HMMER to improve the prediction of coding regions. All predicted peptides were used on InterProScan 5.21-60.0 software for ortholog assignment and annotation of GO terms. Additionally, DEG transcripts were uploaded to the KAAS server for KEGG (Kyoto Encyclopedia of Genes and Genomes) (Kanehisa and Goto 2000). Finally, a custom database for the GAPDH housekeeping gene in eudicots was used to devise putative sequences in *C. florida* and *C. kousa* transcriptomes (see below).

### Gene Ontology enrichment analysis

To gain insights into the functions of the DEGs, genes were annotated and analyzed through gene ontology (GO) by AgriGO v2.0 (Tian, Liu et al. 2017) and the plugin BiNGO v3.0.3 (Maere, Heymans and Kuiper 2005) for Cytoscape v3.3.0 using the *C. florida* or *C. kousa* transcriptomes (Yu, Xiang et al. 2017) GO annotations as background reference. BiNGO was used to calculate overrepresented GO terms in lists of up- and down-regulated DEGs using a hypergeometric test and the Benjamini and Hochberg method to adjust *p*-values, considering significantly enriched GO terms with FDR < 0.05. Additionally, AgriGO was used to perform Singular Enrichment Analysis of GO terms in the lists of DEGs using the hypergeometric statistical test method and the Yekutieli multi-test adjustment method for FDR correction under dependency (Benjamini and Yekutieli 2001) under the significance level of 0.05. Complete GO was selected as the Gene Ontology type. REVIGO (Supek, Bošnjak et al. 2011) was used to summarize and visualize the results of AgriGO.

### Phylogenetic analysis and the conserved motif discovery

Phylogenic relationships of selected R2R3-MYB protein sequences associated with phenylpropanoid pathways from various plant species downloaded from NCBI were inferred using the Neighbor-Joining method (Saitou and Nei 1987) implemented into MEGA11 (Stecher, Tamura and Kumar 2020, Tamura, Stecher and Kumar 2021). The evolutionary distances were computed using the p-distance method (Nei and Kumar 2000) and are expressed in the units of the number of amino acid substitutions per site. This analysis involved 105 amino acid sequences and had 768 informative positions in the final dataset. All ambiguous positions were removed for each sequence pair (pairwise deletion option). Bootstrap values are shown as percentages of 1,000 replicates whenever greater than 50%. The subgroups labels S1 through S28 are designated as per previous studies (Stracke, Werber and Weisshaar 2001, Dubos, Stracke et al. 2010, Shelton, Stranne et al. 2012, Liu, Osbourn and Ma 2015). The species and accession numbers of labeled MYB proteins are provided in Supplementary Table T2. Multiple expectation maximization for Motif Elicitation (MEME), an online bioinformatic tool (http://memesuite.org/), has been used to search for characterized distinctive motifs in MYB repressors, such as EAR with conserved sequence LxLxL and SID interacting motif containing GY/FDFLGL conserved sequence. The species and accession numbers of R2R3-MYB repressor proteins are given in Supplementary Table T3.

### Reverse Transcription Quantitative PCR (RT-qPCR)

Bracts were collected from three cultivars of *C. florida* and two cultivars of *C*. *kousa* one week after emergence, followed by bract collections at two and four weeks after emergence. Samples were collected during the flowering period for three consecutive years (April 2019 – May 2021). Three biological replicates were used for each time point collection. From the total RNA extracted from all samples as described above (Ribospin), cDNA was prepared using qScript ™ cDNA SuperMix (Quantabio, Beverly, MA, USA) according to the manufacturer’s instructions. Primers for the six and four consistently up-or down-regulated candidate regulatory and structural genes were designed based on *C. florida* and *C. kousa* sequences, respectively (Supplementary Table T4). Primers were designed using Primer-BLAST NCBI Genbank with amplification lengths between 70 and 200 bp (Supplementary Table T4). Gene expression was analyzed using PerfeCTa SYBR Green Fast Mix (Quantabio, Beverly, MA, USA). A reaction volume of 20 μL contained 1μL of previously diluted cDNA (1:10), 2 μL of gene-specific primers (1 μM each), and 12 μL of SYBR Green Fast Mix. RT-qPCR experiments were performed on QuantStudio™ 6 Flex (Applied Biosystems, Waltham, MA, USA), and the program was run for 40 cycles, each consisting of 15 s at 95 °C and 1 min at 60 °C. All samples were run in three technical replicates, and no-template controls were included in all plates. The expression of each gene, respectively, was normalized to GAPDH expression for each plant at 1 week post emergence using the 2^-ΔΔCT^ method (Livak and Schmittgen 2001). Data analysis was performed using R v4.2.0 with the following packages: ggplot2 v3.5.1, dplyr v1.1.4, tidyr v1.3.1, multcomp v1.4-26, emmeans v1.10.5, stringr v1.5.1, and broom v1.0.6. RT-qPCR data for each gene was analyzed using a two-way ANOVA (Kaufmann and Schering 2007) to examine the effects of time after bracts emergence and cultivar on that gene’s expression. Post-hoc analyses were performed using Tukey’s HSD test with Sidak adjustment for main effects and Tukey adjustment for interactions at α = 0.05. Compact letter displays were generated to visualize significant differences between groups. Mean expression values were calculated for each Cultivar and Stage.

### RNA-seq Data Processing and Pathway-Centric Analysis

RNA sequencing data from 13 SRA libraries representing the five *Cornus* cultivars were processed through a reproducible computational pipeline. Raw reads were quality-controlled using FastQC and trimmed with Trimmomatic to remove adapters and low-quality bases. *De novo* transcriptome assemblies for *C. florida* and *C. kousa* were generated using Trinity v2.15.1, followed by protein-coding sequence prediction via TransDecoder. Pathway-specific gene identification focused on 14 anthocyanin biosynthesis steps (e.g., PAL, CHS, ANS), as advised by KEGG (Kanehisa and Goto 2000). Putative paralogs were annotated by querying assembled transcripts against reference protein sequences (UniProt/SwissProt) using BLASTP (e-value ≤1e-20, query coverage ≥80%). Reciprocal BLAST validation ensured orthology, and isoforms were resolved using Salmon-based transcript quantification.

Expression values from multiple SRA replicates per cultivar were aggregated using a two-tiered approach: paralogs within each pathway step were averaged, and biological replicates were combined via median expression values. Zero counts, representing undetected expression, were treated as missing data and excluded from calculations, consistent with recommendations for handling sparse expression values in RNAseq studies (Biesecker 2013). Metadata integration linked SRA accessions to cultivar phenotypes through structured column headers (e.g., “Cherokee Brave_SRR21411472”), which enabled cross-referencing of molecular data with bract coloration traits. Normalized expression values were analyzed using a hierarchical statistical approach: (1) Cultivar-level comparisons: One-way ANOVA identified significant expression differences (α = 0.05) across cultivars, with Tukey’s HSD post-hoc testing (critical Q = 3.74–4.58, gene-specific degrees of freedom) assigning letter groupings (A/B/C) to distinguish non-significant pairs. (2) Phenotype-species interactions: Two-way ANOVA evaluated the main effects of species (*C. florida* vs. *C. kousa*) and phenotype (red/pink vs. white/green), with Benjamini-Hochberg correction controlling false discovery rates. This integrative methodology enabled robust identification of regulatory nodes (e.g., *05_CHI* in *C. florida*, *12_ANS* in *C. kousa*) underlying species-specific anthocyanin accumulation patterns, while maintaining transparency across 148 paralogs and 14 enzymatic steps.

## RESULTS

### Color Variation in Dogwood Bracts

The variation in anthocyanins content and color intensity in flowering and Asian dogwood cultivars depended on the season, climate, and stage of plant development. Among *C. florida* red-bracted cultivars used in this study, ‘Cherokee Chief’ mature bracts exhibited the deepest rose-red blooms with beige to pale green edges and highlights in the center (Figure 1A). ‘Cherokee Brave’ displayed reddish-pink bracts with a pale green center, which was prevalent in the early stages of development (Figure 1A). ‘Cloud Nine’ was the only pure white-bracted cultivar, which occasionally had dark purple notches (Figure 1A). Asian dogwood cultivar ‘Rosy Teacups’ (Figure 1A) manifested light to medium pink bracts, whereas ‘Greensleeves’ (Figure 1A) produced white bracts with green to creamy highlights.

Absorbance maxima of anthocyanins isolated from 4-week-old bracts was between 520 and 540 nm (Figure 1B). This is in agreement with known spectra of two major anthocyanin species detected in the fruits *C. florida* and *C. kousa*, cyanidin-3-O-glucoside and cyanidin-3-O-galactoside, respectively. These spectra confirm production of similar anthocyanins in fruits and in bracts of both species. A minor peak in the same range detected for ‘Cloud Nine’ could originate from the colored bracts edges or anthocyanins that are not visual to the bare-eye (Figure 1A). Lack of such a peak in ‘Greensleeves’ was not due to diminished phenolics metabolism; indeed, this cultivar had the highest phenolics levels of all analyzed samples (7.9 mg phenols per mg fresh weight; Figure 1C). Similarly, the white-bracted ‘Cloud Nine’ had the highest levels of phenolics among the three analyzed cultivars of *C. florida*.

### RNA sequencing and alignment

RNAseq was performed with individual libraries for each set of bracts collected from each tree, yielding between ∼28 million to ∼ 39.2 million raw paired end reads per library. After the adaptor and low-quality sequences trimming, more than 71% and 74% of total clean reads were successfully mapped to flowering and Asian dogwood transcriptomes, respectively, because their reference genomes and annotations were not publicly available at this study’s completion (Supplementary Table T1).

### Phylogenetic analysis

Phylogenetic analysis of R2R3-MYB TFs involved in regulating phenylpropanoid-derived compounds, such as monolignols, anthocyanins, proanthocyanidins, flavonols, and isoflavonoids, was conducted to assess the candidate roles of identified *C. florida* and *C. kousa* R2R3-MYB genes (Supplementary Figure F1). CfMYB1 80 and CkMYB1 protein sequences were clustered to subgroup 5, to which activators of anthocyanin and proanthocyanidin synthesis are classified. All of these activators harbor R3 repeat [D/E]Lx2[R/K]x3Lx6Lx3R, which is an interacting bHLH motif (Grotewold, Sainz et al. 2000, Zimmermann, Heim et al. 2004). Correspondingly, Pfam database (Finn, Coggill et al. 2016) was used for protein and domain sequence classification. MYB-like DNA binding domains were detected in CfMYB2, CfMYB3, CfMYB5 78, CfMYB6 30, CkMYB1, and CkMYB2 protein sequences. All six sequences belonged to Pfam clan CLO123, which contains a diverse range of DNA-binding domains with a bHLH motif. Notably, MYB-like DNA binding domains were not detected in CfMYB1 80 protein sequence, which could be due to sequence fragmentation. CfMYB2, CfMYB3, and CkMYB2 protein sequences were clustered to subgroup 4. Majority of known MYB protein repressors negatively regulate phenylpropanoid metabolism and belong to this subgroup 4. In contrast, C-terminal sequence of these proteins contains either C1 GDIP or C2 EAR conserved motifs (Jin, Cominelli et al. 2000). CfMYB5 78 and CfMYB6 30 did not cluster with any designated subgroups; that could be due to lineage- or species-specific “orphan” genes or the phylogenetic analysis methodology used here (Supplementary Figure F1). MEME, an online tool, was used to search for GDIP and EAR motives (Figure 2A). CfMYB2, CfMYB3 in *C. florida*, and CkMYB2 in *C. kousa* contained EAR pdLNLD/ELxiG/S motif located at C-terminal behind MYB R repeat (Figure 2B).

**Figure 2.**
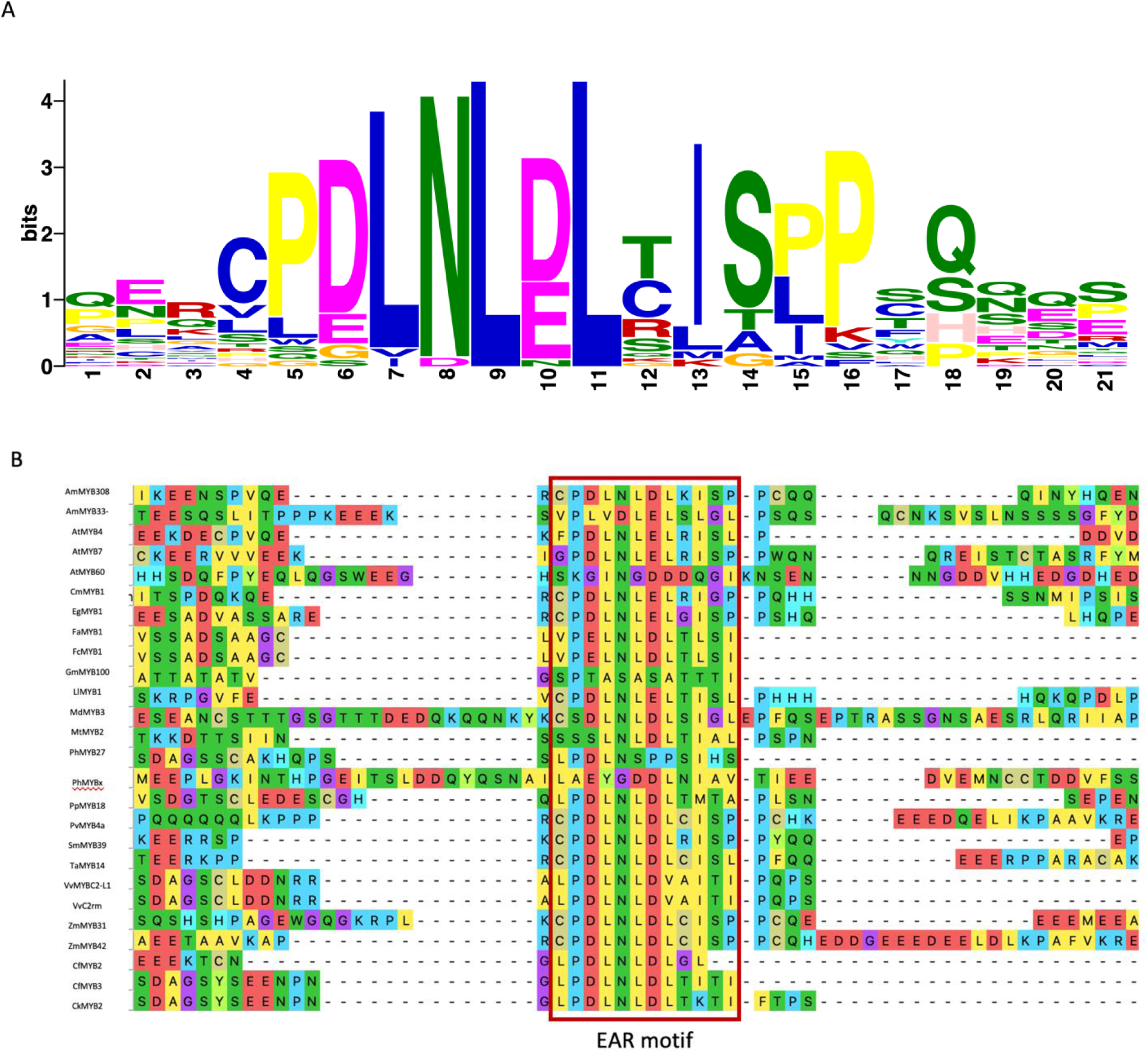
Positional enrichment of Ethylene-responsive element binding factor-associated Amphiphilic Repression (EAR) motif in R2R3-MYB transcription factors of *Cornus florida* and *C*. *kousa*. (**A**) Distribution of the Ethylene-responsive element binding factor-associated Amphiphilic Repression (EAR) motif in 26 R2R3-MYB protein sequences from *C. florida* and *C. kousa*. (**B**) Multiple sequence alignment showing EAR motif localization at the C-terminal region of CfMYB2, CfMYB3, and CkMYB2, downstream of the MYB R repeat.

### DEGs Involved in Anthocyanin Biosynthesis Pathway

Differential gene expression analysis identified 1,156 significant DEGs in *C. florida,* of which 615 were upregulated and 541 were downregulated, based on the thresholds for log2 fold change |log_2_FC| ≥ 1 and false discovery rate (FDR) ≤ 0.05 (Supplementary Table T5). In *C. kousa*, 1,396 were identified as DEGs, of which 753 were upregulated and 643 were downregulated (Supplementary Table T6). Sample distances were represented using t-SNE plots and heatmaps to evaluate the overall transcriptomes of three flowering dogwood cultivars (Figure 3A-B) and two Asian dogwood cultivars (Figure 3C-D). The 13 transcriptome profiles were separated into five clusters that correlated with bract color and cultivar when reads were mapped to either *C. florida* or *C. kousa* transcriptome. GO enrichment analysis was applied to identify the functions and biological pathways involved in the DEGs (Supplementary Table T7-8). Moreover, DEGs for both species were employed for the KEGG pathway enrichment using AgriGO v2.0 analysis (Tian, Liu et al. 2017) and yielded significant GO terms (FDR < 0.05) involved in pathways such as plant hormone signal transduction, metabolic pathways, and membrane transporters belonging to MATE (Multidrug and Toxic Compound Extrusion) which are involved in the transport of anthocyanins to the vacuole (Supplementary Figure F2A-B). This study’s goal was to better understand phenylpropanoid metabolism regulation, mainly anthocyanin regulation; hence, MYB TFs as key regulators were studied further. The seven candidates *R2R3-MYB* genes were identified as differentially expressed using the RNAseq data at an adjusted *p*-value or FDR < 0.05 when comparing *C. florida* red and white-bracted cultivars and *C. kousa* pink and light green-bracted cultivars, respectively. Candidate genes designated as *CfMYB1 80*, *CfMYB2*, *CfMYB3*, *CfMYB5* 78, *CfMYB6 30, CkMYB1*, and *CkMYB*2 were identified as downregulated in non-red bracted *C. florida* and *C. kousa* cultivars. Additionally, three structural genes involved in anthocyanin biosynthesis – *CHI*, *CHS*, and *F3H* – were downregulated in non-red bracted *C. florida* and *C. kousa* cultivars.

**Figure 3.**
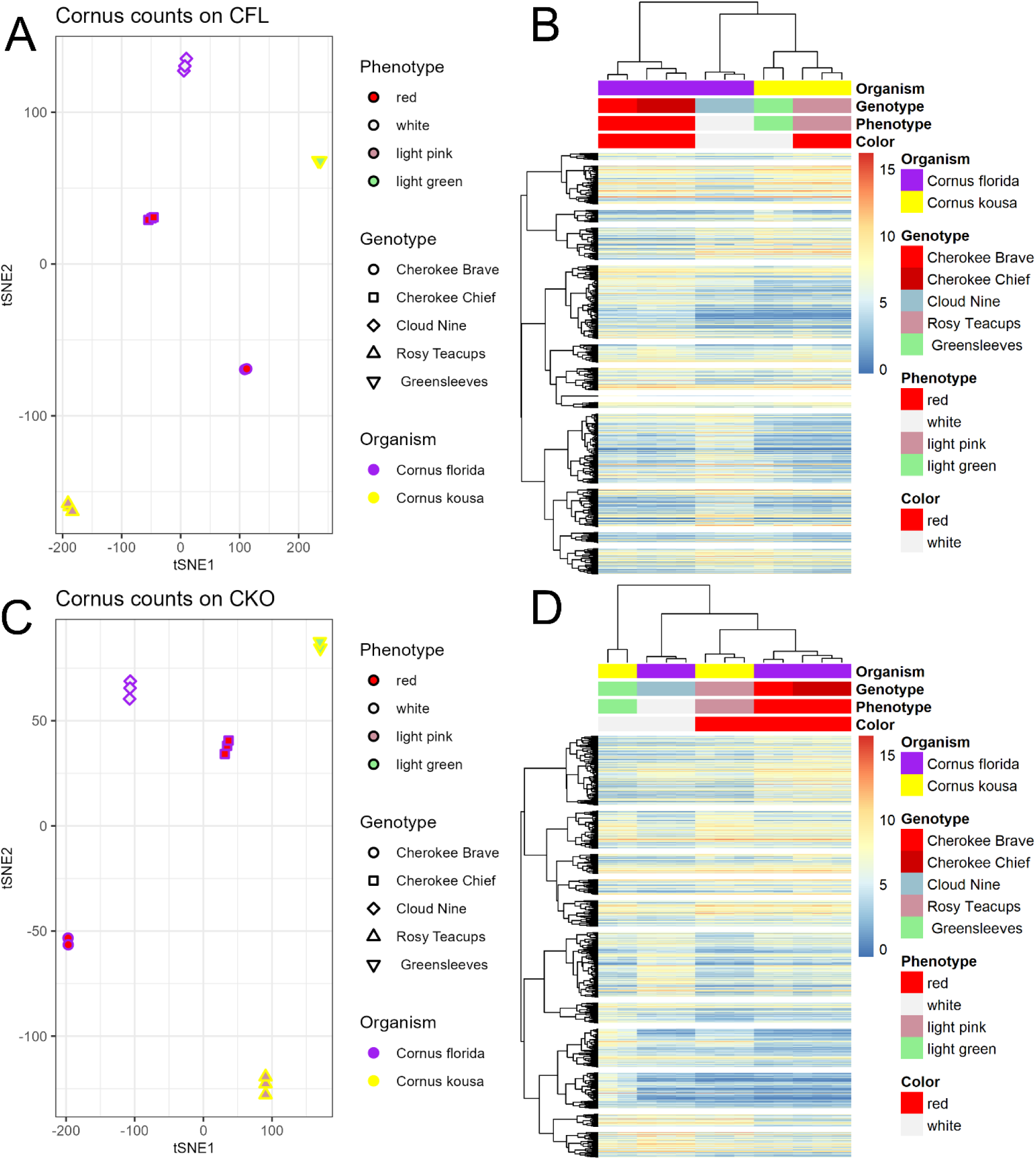
Transcriptomic clustering and expression heatmaps of *Cornus* bract samples. **(A, C)** Hierarchical clustering of 13 RNA-seq samples mapped to the *C. florida* (**A**) and *C. kousa* (**C**) transcriptomes, grouped by cultivar and bract color. (**B**, **D**) Heatmaps of normalized expression values for bract color-associated DEGs, showing clustering by species (**B**) and by color (**D**).

### Expression Dynamics of R2R3-MYB Transcription Factors Across Dogwood Cultivars and Developmental Stages

The qRT-PCR analysis revealed distinct, statistically robust patterns of gene expression for both regulatory (R2R3-MYBs; Figure 4) and structural genes (Supplementary Figure 3) involved in the anthocyanin biosynthesis pathway in *Cornus florida* and *C. kousa*. Expression data were evaluated across multiple cultivars, species, and three developmental stages, with biological replicates that provided high statistical power. Two-way and three-way ANOVA, followed by Tukey HSD post-hoc tests, demonstrated that cultivar, developmental stage, and their interaction frequently had significant effects on the transcriptional regulation of key genes.

**Figure 4.**
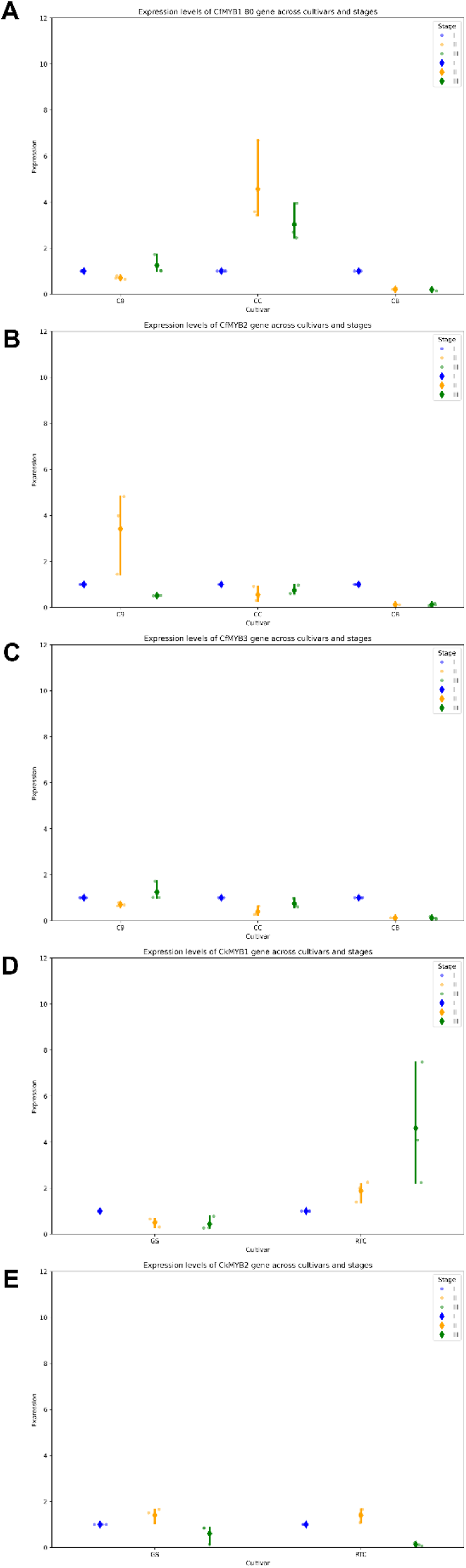
qRT-PCR validation of candidate R2R3-MYB gene expression in *Cornus* bracts. Expression levels of (**A**) CfMYB1_80, (**B**) CfMYB2, (**C**) CfMYB3, (**D**) CkMYB1, and (**E**) CkMYB2 across three developmental stages (1, 2, and 4 weeks post-emergence) in five cultivars. Expression normalized to GAPDH and analyzed using the 2^^−ΔΔCT^ method. Error bars represent standard deviation of three biological replicates.

***CfMYB1_80*** (Figure 4A): For the gene *CfMYB1_80*, cultivar had a significant effect (F=30.81, p=0.000002), and the cultivar × stage interaction was also significant (F=9.56, p=0.00025). Post-hoc groupings indicated that only the ‘Cherokee Chief’ at stages II and III were significantly different from all other combinations, consistent with the role of subgroup 5 MYBs as positive regulators of anthocyanin and proanthocyanidin biosynthesis, particularly in red-bracted cultivars during late development.

***CfMYB2*** (Figure 4B): A significant main effect of both cultivar (F=10.04, p=0.0012) and stage (F=5.21, p=0.0164), as well as a highly significant cultivar × stage interaction (F=8.79, p=0.0004), was observed for *CfMYB2* expression. Post-hoc analysis revealed that the ‘Cloud Nine’ sampled at stage II was uniquely elevated compared to all other cultivar-stage combinations (*p*<0.05), which suggested a distinct role for this gene in suppressing anthocyanin accumulation at this specific developmental window. This fine-tuned, temporally restricted expression is consistent with the function of subgroup 4 MYBs as negative regulators of flavonoid biosynthesis.

***CfMYB3*** (Figure 4C): This gene displayed significant main effects for both cultivar (F=5.79, p=0.0089) and stage (F=6.41, p=0.0059), with significant interaction between both factors (F=8.54, p=0.0005). Post-hoc HSD revealed that ‘Cherokee Brave’ at stages II and II, as well as ‘Cherokee Chief’ at stages II and III were significantly different from ‘Cloud Nine’ sampled at various stages, which indicated that *CfMYB3* is differentially regulated across genetic backgrounds and developmental phases. This pattern supports the hypothesis that *CfMYB3* contributes to the suppression of anthocyanin synthesis in specific cultivars and stages, in line with its phylogenetic placement among known repressors.

***CkMYB1*** (Figure 4D): In *C. kousa*, *CkMYB1* expression was significantly affected by cultivar (F=6.75, p=0.019) and the cultivar × stage interaction (F=5.49, p=0.020). ‘Rosy Teacups’ sampled at stage III showed significantly higher expression of *CkMYB1* than all other combinations, suggesting a cultivar-specific, developmentally regulated activation of anthocyanin synthesis in this pink-bracted variety.

***CkMYB2*** (Figure 4E): *CkMYB2* was significantly regulated by stage (F=23.60, p=0.00002), with no significant cultivar or interaction effects. Tukey HSD revealed that stage II had the highest expression, followed by stage i, and the lowest levels were seen at the latest stage III. This developmental regulation is consistent with the role of subgroup 4 MYBs in temporally restricting anthocyanin accumulation.

### Expression Dynamics of Biosynthesis Genes Across Dogwood Cultivars and Developmental Stages

***CHI*** (Supplementary Figure F3A): The chalcone isomerase (CHI) expression was strongly affected by cultivar (F=11.28, p=0.0002), stage (F=12.50, p=0.0001), and species × cultivar (F=8.74, p=0.0010), with a significant three-way interaction (F=2.98, p=0.035). The highest expression was observed in the *C. florida* ‘Cloud Nine’ stage II, which was significantly different from all other combinations. This suggests that CHI is a major regulatory checkpoint for anthocyanin accumulation in dogwood bracts, with strong genetic and developmental control. The consistent upregulation in ‘Cloud Nine’ as compared with RNA-seq data solidifies CHI’s role as a key flux control point. Phenotype-specific discrepancies in white cultivars suggest post-transcriptional modulation, necessitating protein-level validation.

***CHS*** (Supplementary Figure F3B): The chalcone synthase (CHS) was significantly regulated by stage (F=10.85, p=0.0003), species × cultivar (F=14.75, p=0.00004), and species × cultivar × stage (F=6.18, p=0.00094). The *C. florida* ‘Cloud Nine’ at stage II again showed the highest expression, consistent with this gene’s role as the first committed step in flavonoid biosynthesis. The disconnect between transcript abundance and statistical significance in RNA-seq, alongside qRT-PCR’s stage-specific spikes, implies post-transcriptional regulation (e.g., translational control or protein stability). This decoupling underscores the limitation of relying solely on mRNA data for pathway flux interpretation.

***F3H*** (Supplementary Figure F3C): The flavanone 3-hydroxylase (F3H) expression was significantly influenced by developmental stage (F=5.93, p=0.0081), but not by cultivar or interaction of the factors. Tukey HSD post-hoc tests indicated that the latest-sampled stage III was significantly lower than stages I and II (p<0.05), which suggested a developmentally regulated downregulation of this early biosynthetic gene as bracts mature. Whereas RNA-seq identified genetic background-driven regulation of this gene (see above), qRT-PCR highlighted temporal specificity, particularly in C. kousa. This suggests stage-specific sampling bias in RNA-seq’s whole-bract approach, which may average temporal fluctuations.

Collectively, these results demonstrate that the regulation of anthocyanin biosynthesis in dogwood bracts is highly complex and involves multiple layers of control by R2R3-MYB transcription factors and structural genes, with strong effects of genetic background, developmental stage, and interactions. The fine-scale, temporally restricted expression of MYB repressors and activators suggests that dogwoods have evolved distinct, cultivar-specific regulatory circuits to modulate brightly colored bracts, a trait of high horticultural and ecological significance (ALBORNOZ, ROSAS and LÓPEZ 2023).

### Expression Patterns in the Anthocyanin Biosynthesis Pathway

RNA-seq analysis across the 14 enzymatic steps of the anthocyanin biosynthesis pathway revealed distinct, multi-tiered regulation associated with species identity, cultivar, and bract coloration (Figure 5; Supplementary Table T9). Our three-tiered statistical approach (cultivar vs. species vs. species × phenotype) uncovered regulatory patterns obscured in single-level analyses, which resolved contradictions between transcript abundance and phenotypic outcomes. Three complementary analyses-per-cultivar (Figure 5A), species × phenotype Figure 5B), (and per-species (Figure 5C) demonstrated that 13/14 genes exhibited significant expression differences (ANOVA Type III, p < 0.05), with only chalcone synthase (04_CHS) showing conserved expression across all tiers (p > 0.05). This comprehensive approach uncovered evolutionarily divergent regulatory strategies underlying similar floral phenotypes.

**Figure 5:**
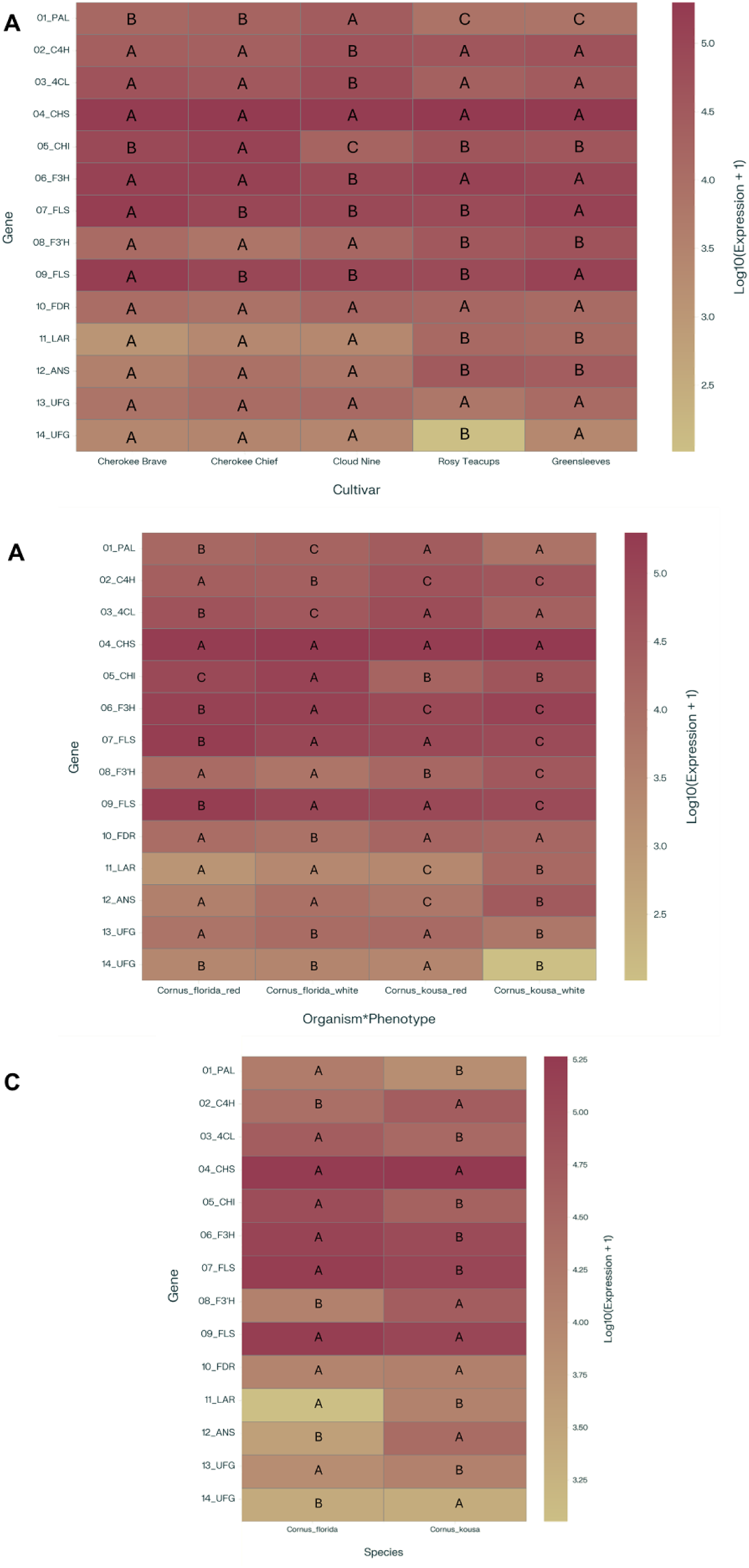
Heatmap of phenylpropanoid pathway gene expression across *Cornus* cultivars. Log10-transformed weighted mean expression values for 14 pathway genes across five cultivars. (**A**) Cultivar-level expression; (**B**) phenotype and species-level; (**C**) species-level. Color gradient: low (whitish #CCBF85) to high (crimson #943A51). Letters indicate statistically homogeneous groups (Tukey HSD, α = 0.05).

The white-bracted *C. florida* cultivar ‘Cloud Nine’ emerged as a metabolic outlier and exhibited statistically distinct expression patterns (Tukey HSD, p < 0.001) for key upstream enzymes. 01_PAL was 2.2-fold higher than ‘Cherokee Brave’ (28,648 vs. 14,502 TPM); 02_C4H was 1.9-fold elevated vs. ‘Rosy Teacups’ (47,740 vs. 40,262 TPM); and 03_4CL showed 3.8-fold increase vs. *C. kousa* ‘Greensleeves’ (67,521 vs. 17,718 TPM). These findings position ‘Cloud Nine’ as a model for upstream flux control, with prioritized phenylpropanoid precursor synthesis. In contrast, *C. kousa* cultivars specialized in late-pathway modification. The 11_LAR (leucoanthocyanidin reductase) expression was 7.0-fold higher in ‘Rosy Teacups’, red-bracted *C. kousa* cultivar, than ‘Cherokee Brave’ (17,987 vs. 2,574 TPMp < 0.001), which suggested species-specific diversion into proanthocyanidins. Despite extreme 04_CHS expression in ‘Rosy Teacups’ (198,697 TPM), statistical analysis revealed no significant differences across cultivars, species, or phenotypes (p = 0.312–0.406). This unexpected conservation at the pathway’s first committed step implies regulatory constraint, with flux controlled downstream through species-specific mechanisms.

Chalcone isomerase 05_CHI emerged as a nexus of evolutionary divergence and exemplified complex regulatory plasticity with tier-specific groupings. Under per-cultivar aggregation ‘Cloud Nine’ was distinct from either *C. kousa* cultivar or from *C. florida* red-bracted cultivars (p < 0.001). The same pattern emerged when aggregated at species × phenotype tier, with ‘Cloud Nine’ being distinct from *C. florida* red or *C. kousa* white/red (p < 0.001). The 4.8-fold difference between *C. florida* red (86,898 TPM) and *C. kousa* ‘Greensleeves’ (4,013 TPM) highlights the 05_CHI’s role as a key phenotypic checkpoint, with species employing distinct mechanisms to modulate flavanone production.

The interplay between species and phenotype uncovered divergent molecular strategies for red bract development. In *C. florida* red cultivars, 05_CHI expression reached exceptional levels (86,898 TPM; p < 0.001), which underscored this enzyme’s role in directing flux toward flavanone synthesis. By contrast, *C. kousa* ‘Rosy Teacups’ prioritized late-step modification, with 12_ANS (anthocyanidin synthase) expression peaking at 36,125 TPM (p = 0.004), a critical enzyme for generating colored anthocyanidins. A paradox emerged in white-bracted cultivars, where 12_ANS expression was 2.1-fold higher in *C. kousa* white (36,125 TPM) than in its red counterparts (17,987 TPM; p = 0.004). This counterintuitive pattern suggests non-canonical roles for 12_ANS in pigment stabilization or UV protection and disentangled the transcript abundance from visible pigmentation and hinted at post-transcriptional regulatory mechanisms (Ray, Mishra et al. 2023).

The statistical grouping patterns across analytical tiers provide a compelling insight into convergent evolution. Both dogwood species achieve vivid red bracts, but *C. florida* relies on upstream commitment through elevated 01_PAL and 05_CHI expression, whereas *C. kousa* employs downstream diversification via 08_F3’H and 12_ANS activation. White phenotypes, despite their visual similarity, achieve convergent downregulation through distinct pathways: *C. florida* white-bracted ‘Cloud Nine’ reduces flux at the pathway’s entry point (01_PAL, 18,375 TMP), whereas *C. kousa* ‘Greensleeves’ suppresses terminal steps (11_LAR, 11,737 TPM).

Floral color evolution in *Cornus* spp. thus emerges as a model for regulatory plasticity, where species traverse unique molecular trajectories to converge on similar phenotypes. The conserved expression of 04_CHS alongside species-specific late-step regulation illustrates how metabolic pathways balance evolutionary constraint with innovation-a paradigm for probing the molecular underpinnings of biodiversity (Treder, Klamkowski et al. 2023). This duality of conservation and divergence underscores the anthocyanin pathway’s remarkable adaptability, to offer new avenues for exploring the genetic basis of floral diversification.

## DISCUSSION

### Importance of Anthocyanin Regulation in Ornamental Dogwoods

Ornamental plants constitute an economically vital component of agriculture and horticulture. Aesthetically important traits such as leaf morphology, flower color, size, and scent, bark color, and plant habitat improve the living ambiance and enhance cultivating sentiment (Linde, Yan and Debener 2007, Wadl, Saxton et al. 2011, Zheng, Li et al. 2021). The whole-genome sequences and draft genome sequences of 69 ornamental plants were published between 2012 and the end of 2020 (Zheng, Li et al. 2021). Yet, less than 20 of these are ornamental trees, and among them only a few produced red, pink, or violet blooms, such as *Prunus mume* SIEB. et ZUCC. (Zhang, Chen et al. 2012), *P*. *serrulate* L. (Yi, Yu et al. 2020), and *P*. *yedoensis* Matsumura (Baek, Choi et al. 2018), *Cercis canadensis* L. (Stai, Yadav et al. 2019), *Osmanthus fragrans* Lour. (Yang, Yue et al. 2018), *Handroanthus impetiginosus* Mart. Ex DC (Silva-Junior, Grattapaglia et al. 2018) and *Bombax ceiba* L. (Gao, Wang et al. 2018). Moreover, candidate genes controlling anthocyanin synthesis, an essential pigment for flower and leaf color, have not been identified in many ornamental woody plants. Recently, the anthocyanin profile has been determined in *C. canadensis* but not the gene machinery orchestrating its production (Veazie, Ma and Werner 2017).

To carry that momentum further, here we aimed to uncover the molecular mechanisms behind bracts color in flowering and Asian dogwoods, important woody ornamentals. Integration of the biochemical spectral analyses of bract extracts with RNAseq and its confirmation using qRT-PCR, as well as the phylogenetic analyses, identified genes involved in anthocyanin production in ornamental dogwoods. To synthesize these multilayered findings, we constructed a regulatory network model (Figure 6) that illustrates the possible interactions between R2R3-MYB transcription factors and structural genes across the anthocyanin biosynthesis pathway. This schematic highlights the cultivar-specific expression patterns and regulatory divergence between *C. florida* and *C. kousa*, thereby providing a visual framework for understanding the molecular basis of bract pigmentation. Our findings generate a great springboard for future functional studies and molecular breeding efforts.

**Figure 6.**
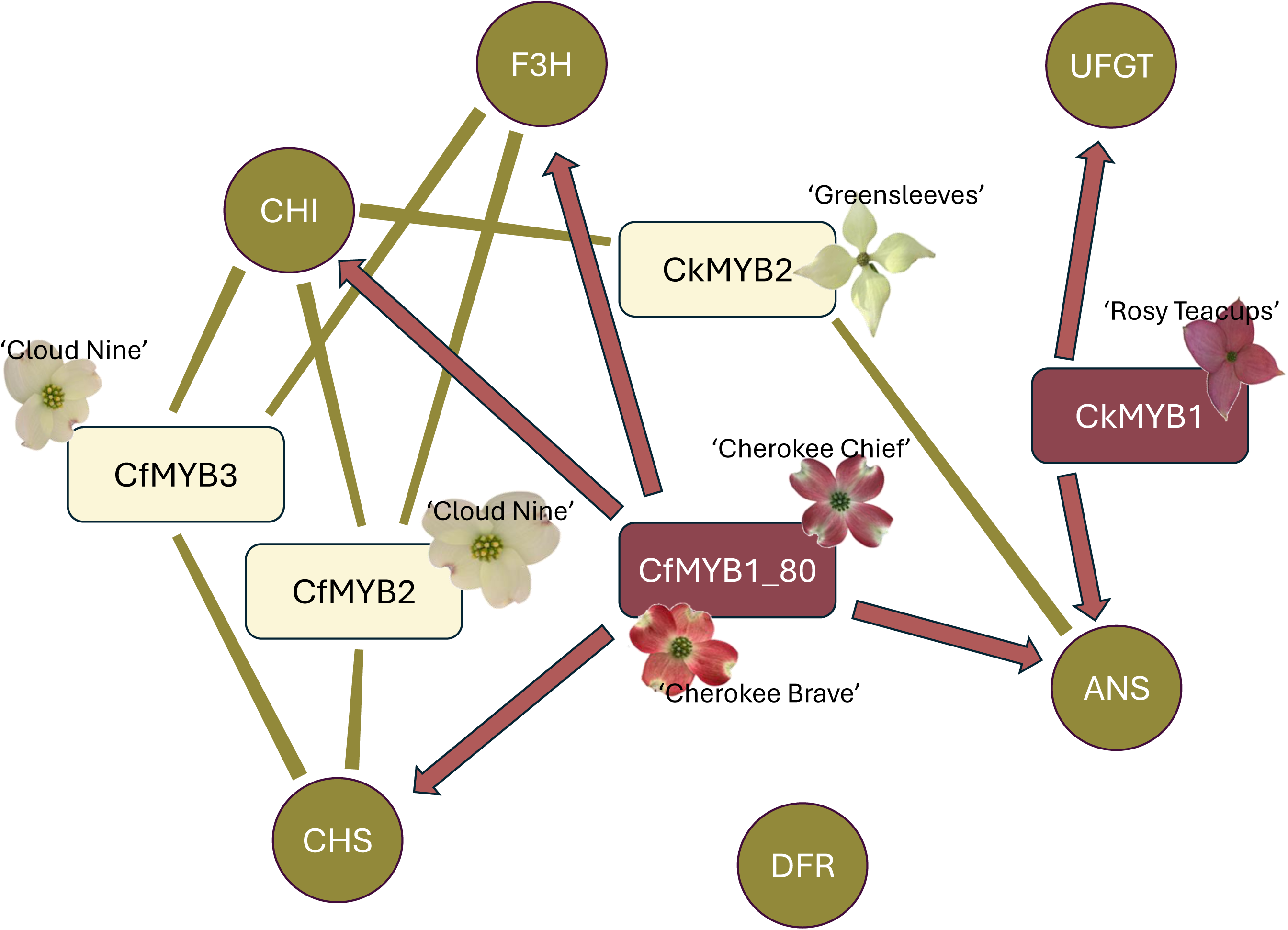
Regulatory network of R2R3-MYB transcription factors and anthocyanin biosynthetic genes. Nodes represent MYB TFs (red = activators, green = repressors) and structural genes (green). Edges indicate regulatory interactions: arrows = activation, rectangles = repression. Gene expression patterns are cultivar-specific, based on RNA-seq and RT-qPCR data. Cultivars are labeled.

### Challenges in Dogwood Breeding and the Role of Molecular Tools

In addition to providing nourishment for wildlife due to the high level of fat, protein, and available calcium in fruit (Thomas 1969), flowering and Asian dogwoods are important ornamental crops essential for the economy of Tennessee, Oregon, North Carolina, and Ohio, among other states, and in 2019, total dogwood sales exceeded USD31 million annually (USDA-NASS 2021). Although more than 150 cultivars are commercially available for both species, consumer demands for specific traits of interest, such as dark red bract or foliage display, resistance to pathogens, variegated foliage, and dwarf or weeping habit, have not been satisfied (Witte, Windham et al. 2000, Cappiello and Shadow 2005, Wadl, Windham et al. 2014, Nowicki, Houston et al. 2020). For instance, white bracted cultivars Kay’s Appalachian Mist, Jean’s Appalachian Snow, and Karen’s Appalachian Blush are resistant to dogwood powdery mildew pathogen (*Erysiphe pulchra* [Cooke & Peck]; (Windham, Witte and Trigiano 2003)). But they lack desirable dark red or pink bracts of cultivars belonging to the Cherokee series, such as ‘Cherokee Brave’, ‘Cherokee Chief’, ‘Cherokee Sunset’, ‘Cherokee Daybreak’, and ‘Appalachian Sunrise’ (personal comm. R. N. Trigiano) which are generally considered either moderately or susceptible to *E. pulchra* (Thurn, Lamb and Eshenaur 2019).

In contrast, Asian dogwoods typically lack variation in bracts colors and do not display a range of bract colors from white to dark red as flowering dogwoods do (Cappiello and Shadow 2005, Molnar, Muehlbauer et al. 2017). ‘Rosy Teacups’, ‘Miss Satomi’, and ‘Rutpink’ are among the few cultivars exhibiting dark to bright red or pink bracts (Trigiano, Ament et al. 2004, Molnar, Muehlbauer et al. 2017). Moreover, a hybrid breeding program between *C. florida* and *C. kousa* was initiated almost a half-century ago, aiming to enhance ornamental traits and develop attractive landscape trees that do not require extensive maintenance (Molnar, Muehlbauer et al. 2017). Notably, time-demanding development of dogwood cultivars typically depends on repeated selection for desirable traits. To achieve development of new cultivars with desirable traits, several challenges need to be overcome, such as a long juvenile period (3 to 5 years), self-incompatibility, inbreeding depression, and high susceptibility of flowering dogwood to two fungal foliar pathogens causing dogwood anthracnose (*Discula destructiva*) or powdery mildew (*E. pulchra*) (Gunatilleke and Gunatilleke 1984, Ament, Windham and Trigiano 2000, Reed 2004, Wang, Wadl et al. 2009, Hadziabdic, Fitzpatrick et al. 2010, Hagan, Akridge et al. 2011, Wadl, Saxton et al. 2011, Mantooth, Hadziabdic et al. 2017, Nowicki, Redington et al. 2025). These challenges highlight the need for molecular approaches to accelerate breeding programs and develop cultivars with desired traits. Red and pink bracted blossoms with their variations foster the attractive ornamental appeal, and in flowering and Asian dogwoods, anthocyanin pigments provide the basis for bract and foliage coloration (Vareed, Reddy et al. 2006, Wadl, Wang et al. 2010, Wadl, Saxton et al. 2011). However, to our knowledge, structural and regulatory genes to control the biosynthesis of anthocyanins and other phenylpropanoid-derived compounds in flowering and Asian dogwoods have not been reported yet. Moreover, genetic resources, quality reference genome, and annotation for *Cornus* sp. are still under active development, though certain genomic information is available (Bewick and Leebens-Mack 2020).

### R2R3-MYB Transcription Factors as Key Regulators of Anthocyanin Biosynthesis

Among the genes controlling anthocyanin production, R2R3-MYB TFs fine-tune flavonoid production and their biological functions, such as involvement in developmental and cell differentiation processes, response to environmental stress, and metabolite biosynthesis, and they have been functionally characterized in cereals, vegetables, fruit, and ornamental crops (Du, Zhang et al. 2009, Dubos, Stracke et al. 2010, Liu, Osbourn and Ma 2015, Wang, Ma et al. 2019, Zhang, Wang et al. 2019, Ding, Patterson et al. 2020, Gong, Li et al. 2020, Yan, Chen et al. 2020, Yan, Li et al. 2020). Concomitantly, to date, a whooping 61 R2R3-MYB TFs have been identified in herbaceous ornamental plants such as *Petunia* sp. (Quattrocchio, Wing et al. 1999, Albert, Lewis et al. 2009, Albert, Lewis et al. 2011), *Rosa* sp. (Lin-Wang, Bolitho et al. 2010), *Lilium* sp. (Nakatsuka, Haruta et al. 2008, Yamagishi, Shimoyamada et al. 2010, Yamagishi 2011, Yamagishi, Yoshida and Nakayama 2012, Yamagishi 2016), *Antirrhinum* sp. (Martin, Prescott et al. 1991, Moyano, Martínez-Garcia and Martin 1996, Schwinn, Venail et al. 2006, Shang, Venail et al. 2011), *Gerbera* sp. (Elomaa, Uimari et al. 2003) but only one in a woody plant, *P*. *meme* (Zhang, Hao et al. 2017).

The MYB TF family is one of the largest in plants and regulates a broad range of metabolic pathways, including responses to abiotic stimuli, regulation of benzenoid, glucosinolate, phenylpropanoid, and terpenoid biosynthesis, and control of developmental and cell differentiation process, e.g., organ formation, cuticle development, and trichome initiation and branching (Shin, Choi et al. 2002, Liu, Osbourn and Ma 2015, Zhao, Cheng et al. 2018, Tan, Man et al. 2019, Fichman, Zandalinas et al. 2020, Gong, Li et al. 2020). MYB TFs are characterized by a highly conserved N-terminal DNA-binding domain repeat (R) and a variable C-terminal regulatory region (C). Each MYB repeat contains approximately 50 amino acids, including unique DNA motifs and MYB binding sites. Based on the sequence similarities, MYB domain repeats are named R1, R2, and R3 depending on the number of repeats, ranging from 1 to 4 repeats. Accordingly, the MYB gene superfamily is divided into the following four families: 1R-MYB, 2R-MYB (R2R3), 3R-MYB, and 4R-MYB. (Lipsick 1996, Rosinski and Atchley 1998, Kranz, Scholz and Weisshaar 2000, Dubos, Stracke et al. 2010, Jiang and Rao 2020).

### Comparative Expression and Phylogenetic Insights into MYB Regulators

The first plant R2R3-MYB gene, COLOREDI, was identified in *Zea mays* and is crucial for enhancing anthocyanin biosynthesis in the aleurone tissue (Klempnauer, Gonda and Bishop 1982). Since the identification of COLOREDI, the scientific interest in R2R3-MYB family has yielded insights and breeding progress in many economically important crops. The constant development of fast and cost-effective next-generation sequencing technologies has enabled the identification of more than 600 R2R3-MYB genes across 130 plant species (Morozova and Marra 2008, Wu, Wen et al. 2022, Yin, Guo et al. 2022). Although the consensus regarding the classification schemes and nomenclature uniformity of R2R3-MYB TFs is still lacking, mainly because of lineage-specific gene loss or expansion, differences in sample coverage, and the type of phylogenetic analysis applied, R2R3-MYB TFs have been classified into 23 to 90 subgroups (Du, Liang et al. 2015, Jiang and Rao 2020, Li, Wen et al. 2020). Accordingly, each R2R3-MYB subgroup regulates a specific metabolic pathway (Wu, Wen et al. 2022).

Anthocyanin and proanthocyanidins activators of subgroup 5 required specific bHLH TFs to encode transcription of structural genes (Chen, Hu et al. 2019, Ma and Constabel 2019, Wu, Wen et al. 2022). In contrast, MYB repressors, which act to reduce anthocyanin biosynthesis, belong exclusively to subgroup 4 and contain the N-terminal regions with a DNA-binding domain as well. The N-terminal region of MYBs is highly conserved, and its function usually depends on the C-terminal region’s distinct motifs. The C1 LlsrGIDPxT/sHRxI/L region and C2 region pdLNLD/ELxiG/S are referred to as GIDP and EAR motif (Ethylene-responsive element-binding factor associated Amphiphilic Repression motif), respectively. EAR is the most common transcriptional repression motif found in plants. The C3 and C4 motifs are also present in a small segment of the R2R3-MYB structural domain (Ohta, Matsui et al. 2001, Kagale and Rozwadowski 2011, Chen, Hu et al. 2019, Ma and Constabel 2019). Majority of the identified and functionally characterized MYBs involved in anthocyanin biosynthesis, such as MdMYB1, MdMYBA, and MdMYB3 in apples, ROSEA1 and ROSEA2 in snapdragon, AN2, DPL, and PHZ in petunia positively regulate the transcriptional expression of structural genes involved in this pathway (Moyano, Martínez-Garcia and Martin 1996, Schwinn, Venail et al. 2006, Albert, Lewis et al. 2011, Albert, Davies et al. 2014). Upstream genes (early biosynthetic genes) such as CHI (chalcone isomerase), CHS (chalcone synthase), and F3H (flavanone 3-hydroxylase), are the key enzymes to mediate and regulate anthocyanin biosynthesis, which is branched from the phenylpropanoid pathway. In comparison, DFR (dihydroflavonol 4-reductase), ANS (anthocyanidin synthase), and UFGT (flavonoid 3-O-glucosyltransferase) are downstream genes (late biosynthetic genes) (Pelletier, Burbulis and Winkel-Shirley 1999, Takos, Jaffé et al. 2006, Cominelli, Gusmaroli et al. 2008, Vimolmangkang, Han et al. 2013).

### Structural Gene Expression and Developmental Timing

Our transcriptome study aimed to identify important genes involved in phenylpropanoid metabolite biosynthesis, particularly anthocyanin and proanthocyanidins production, using RNAseq and RT-qPCR for validation. Five R2R3-MYB TFs were detected as downregulated in *C. florida* and only two in *C. kousa*, which are lower numbers compared to other plant species such as 126 genes in *A. thaliana* (Stracke, Werber and Weisshaar 2001), 192 in *Populus* (Wilkins, Nahal et al. 2009), 244 in soybean (Du, Yang et al. 2012), or 406 in *Gossypium hirsutum* L. (Wang, Ma et al. 2019). Genome unavailability and mapping of the RNAseq reads to *C. florida* and *C. kousa* leaf transcriptomes likely affected our ability to mine more R2R3-MYB TFs. In contrast, downregulation of the R2R3-MYB TFs and of the upstream structural genes *CHS*, *CHI,* and *F3H* might result from RNAseq of physiologically fully developed bracts at four weeks after emergence – a possible artifact of sampling at the stage where the transcripts are past their peak levels. Moreover, expression levels of *CHS* and *F3H,* key enzymes involved in flavonoid and anthocyanin synthesis, were increased in emerging two weeks old bracts of ‘Cloud Nine’, ‘Cherokee Chief’, and ‘Rosy Teacups’. As we focused on detailed analyses of DEGs, late biosynthetic genes were not detected as significantly up- or downregulated, which could possibly limit our detection of other MYB TFs or upstream structural genes in dogwoods. In apples, MdMYB10 orchestrates the synthesis of anthocyanins in the peel, flesh, and color of foliage at the early stage (Espley, Hellens et al. 2007); whereas MdMYB10b regulates the color of the fruit cortex at the later phases (Chagné, Lin-Wang et al. 2013). Collectively, this confirms previous findings that the anthocyanins content varies greatly depending on the species’ genetic background, environmental conditions, and stage of plant development (Wadl, Saxton et al. 2011, Jaakola 2013, Mattioli, Francioso et al. 2020). Contrastingly, others (Salvatierra, Pimentel et al. 2013) showed that expression levels of early biosynthetic genes are not crucial in the flavonoid pathway for pigment manifestation. Finally, phenylpropanoid biosynthesis is a complex metabolic grid with many branches and alternative metabolic routes (Yan, Chen et al. 2020, Yan, Li et al. 2020, Chen, Nowicki et al. 2023).

Functions and characteristics of R2R3-MYB proteins have been studied extensively, and their comparative phylogenetic analysis brought to light both the diversity and the conservation in this gene family (Stracke, Werber and Weisshaar 2001, Wilkins, Nahal et al. 2009, Du, Yang et al. 2012, Wang, Tang et al. 2019, Wang, Ma et al. 2019). Also, studies in soybean and petunia indicated that anthocyanin activators and repressors could be identified by phylogenetic analysis (Elomaa, Uimari et al. 2003, Albert, Lewis et al. 2011). Accordingly, phylogenetic analysis grouped the candidate R2R3-MYB proteins identified in this study in two major subgroups, 4 and 5. R2R3-MYB TFs CfMYB2, CfMYB3, and CkMYB2 clustered in subgroup 4, which genes negatively regulate phenylpropanoid metabolism. Consistently, repressors of this group were identified in various plant species, including strawberry FaMYB1 (*Fragaria* x *ananassa* Duchesne) (Aharoni, De Vos et al. 2001) and FcMYB1 (*Fragaria chiloensis*) (Salvatierra, Pimentel et al. 2013), petunia PhMYB27 (Albert, Davies et al. 2014), grapevine VvMYBC2-L1/ and VvMYB4-like (Cavallini, Matus et al. 2015, Pérez-Díaz, Pérez-Díaz et al. 2016), poplar PtrMYB182 and PtrMYB57 (Yoshida, Ma and Constabel 2015, Wan, Li et al. 2017), crabapple MdMYB3 (Tian, Zhang et al. 2017), peach PpMYB17-20 (Zhou, Peng et al. 2016), and Chinese narcissus NtMYB2 (Anwar, Wang et al. 2018).

PhMYB27 is part of the R2R3-MYB, bHLH, and WDR complex and functions as an anthocyanin repressor in petunia petals (Albert, Davies et al. 2014). *NtMYB2* suppresses red pigmentation in the petals and corona of Chinese narcissus by repressing the transcript levels of structural genes involved in the anthocyanin biosynthetic pathway (Anwar, Wang et al. 2018). Likewise, expression levels of *CfMYB2* significantly increased in ‘Cloud Nine’ white bracts at two weeks after emergence, whereas *CfMYB3* levels slightly increased in the same cultivar when four weeks old. CfMYB2 showed protein sequence similarity with AtMYB4. *AtMYB4* is known to negatively regulate the sinapate ester biosynthesis induced by UV-B in *Arabidopsis*; therefore, it is a regulator of the phenylpropanoid pathway through the expression of the cinnamate-4-hydroxylase (*C4H*) gene (Jin, Cominelli et al. 2000). Moreover, phylogenetic analysis indicated strong sequence similarity of CfMYB2 with FaMYB1 from strawberry and PhMYB27 from petunia (Aharoni, De Vos et al. 2001, Albert, Lewis et al. 2011), which appear to be R2R3-MYB suppressors of structural genes involved in the anthocyanin pathway. CfMYB3 and CkMYB2 protein sequences were most similar to AtMYB3, which affects sinapoyl malate and negatively regulates anthocyanin biosynthesis in *Arabidopsis*. Its activity is regulated by corepressors NIGHT LIGHT-INDUCIBLE (LNK1) and CLOCK-REGULATED (LNK2), which mediate the binding of AtMYB3 to the *C4H* (Zhou, Zhang et al. 2017). Phylogenetic analyses of CfMYB3 and CkMYB2 placed them closest to grapes repressors VVMYBC2-L1 and VVC2rm, which are involved in suppressing anthocyanin and proanthocyanidins production (Cavallini, Matus et al. 2015). Surprisingly, *CkMYB2* transcript levels increased in both *C. kousa* cultivars used in this study, which may explain the color variation pink intensity of ‘Rosy Teacup’ bracts. Finally, in grapes and petunia, R2R3-MYB repressors affect both anthocyanins and proanthocyanidins, thereby suggesting that their role is less specific compared to R2R3-MYB activators (Cavallini, Matus et al. 2015, Ma and Constabel 2019). CfMYB1 80 and CkMYB1 were clustered to subgroup 5 with other anthocyanin and proanthocyanidin R2R3-MYB activators. In ‘Cherokee Chief’, which bracts displayed the darkest shade of red, expression levels of *CfMYB1 80* were significantly increased when bracts were two and four weeks old. Also, the expression levels of *CkMYB1* were comparably greater in the pink bracted cultivar ‘Rosy Teacup’ over ‘Greensleeves’. Anthocyanin-encoding genes belonging to subgroup 5 were reported in maize *aleurone 1* (C1) (Carey, Strahle et al. 2004) and *purple leaf* (PL) (Cone, Cocciolone et al. 1993) and in orchid *OgMYB1* (Chiou and Yeh 2008). *OgMYB1* undergoes active expression during floral development in red sepal and petal tissues, to regulate the color patterns in *Oncidium* spp. Gower Ramsey (Chiou and Yeh 2008).

Another R2R3 MYB TFs of subgroup 5 encompass the proanthocyanidins or condensed tannins activators. They are colorless flavonoid polymers in many plants’ leaves and seeds. They confer defense against pathogens and herbivore feeding (Dixon, Xie and Sharma 2005). Proanthocyanidins have not been reported in *C. florida* and *C. kousa*. In this study, CfMYB1 80 and CkMYB1 showed the highest sequence similarity with AtMYB123 protein. *AtMYB123* regulates the production of proanthocyanidins in Arabidopsis, regulating *DIHYDROFLAVONOL-4-REDUCTASE* (*DFR*) gene synthesis (Nesi, Jond et al. 2001). R2R3-MYB TFs regulating proanthocyanidins activation were identified in strawberry FaMYB11 (Schaart, Dubos et al. 2013), lotus LjTT2a, LjTT2b, and LjTT2c (Yoshida, Iwasaka et al. 2008), kiwi fruit DkMYB2 and DkMYB4 (Akagi, Ikegami et al. 2009) among other plant species.

## CONCLUSIONS

Our results provided an essential foundation for further analysis of R2R3-MYB TFs that may be responsible for anthocyanin and proanthocyanidins regulation in bracts of *C. florida* and *C. kousa*. Based on our data, future efforts could be directed toward elucidating how these and other candidate genes regulate complex phenylpropanoid metabolism in these species. Moreover, functional characterization of these genes through CRISPR-Cas9, ectopic plant leaf expressions, or genetic engineering will enable heterologous production of anthocyanins. The envisioned development of functional markers or the application of marker-assisted selection in complicated dogwood breeding will allow early selection of red-bracted dogwood cultivars for gainful commercial release.

## Acknowledgements

Authors gratefully recognize the Hidden Hollow Nursery at Belvidere, TN for providing us with trees for this research. Ms. Sarah L. Boggess MS and Grace Pietsch are gratefully recognized for technical assistance and plant maintenance. The Staton Lab members are gratefully recognized for their help with bioinformatic analyses.

## Ethics and Data Availability Statement

No protected species were used in this study.

All relevant data are within the paper and its Supporting Information files. Raw RNAseq data are available at NCBI under the short reads accession numbers SRR21411460 through SRR21411472. Sequences of MYBs identified in this research are available in GenBank: OP382880 through OP382884.

## Supplementary Figures

**Supplementary Figure F1.**
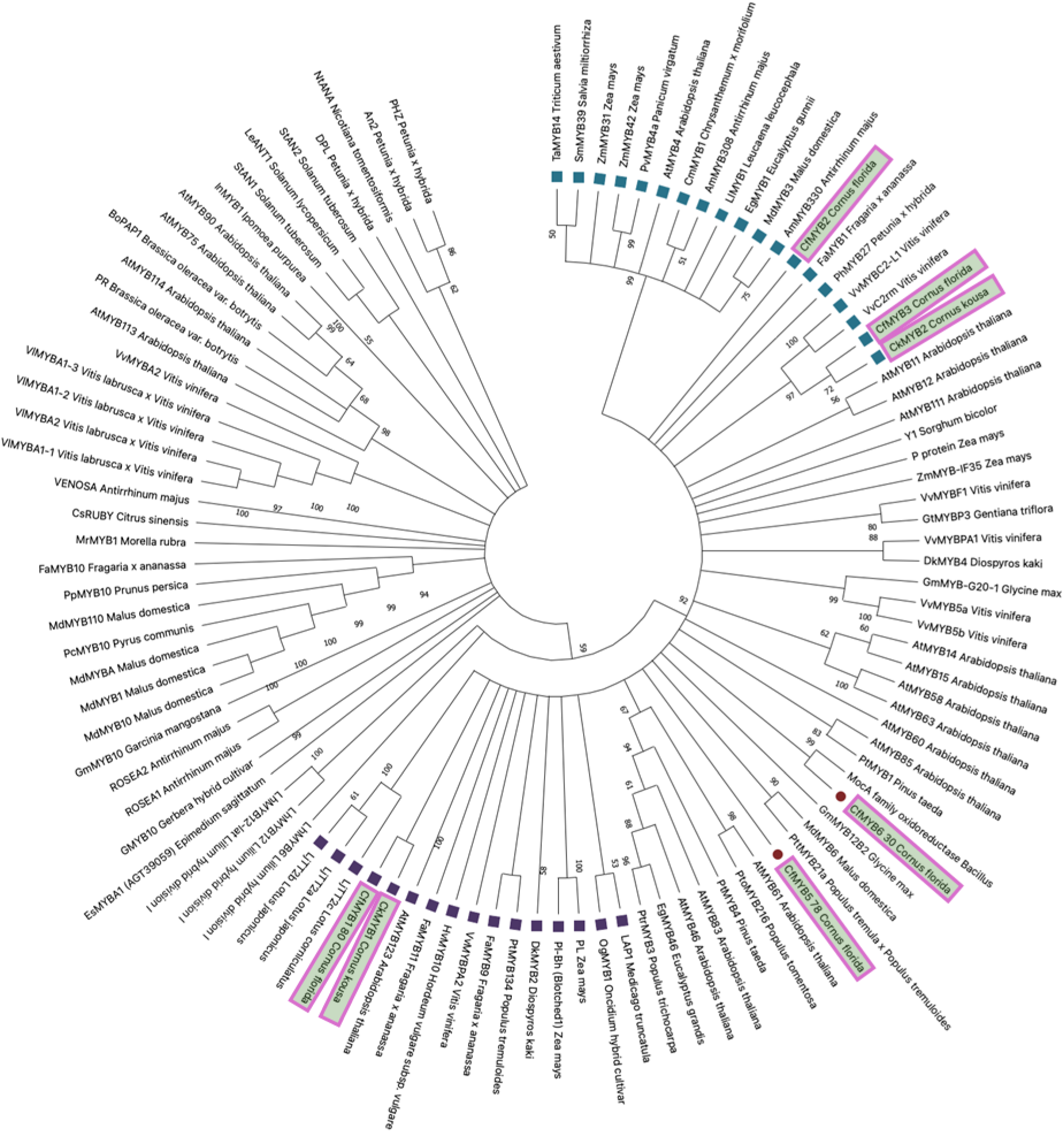
Phylogenetic tree of R2R3-MYB transcription factors. Constructed using neighbor-joining method with 105 full-length MYB protein sequences. Bootstrap values (>50%) shown. Subgroups S1–S28 labeled per Stracke et al. (2001). *Cornus* MYBs highlighted.

**Supplementary Figure 2.**
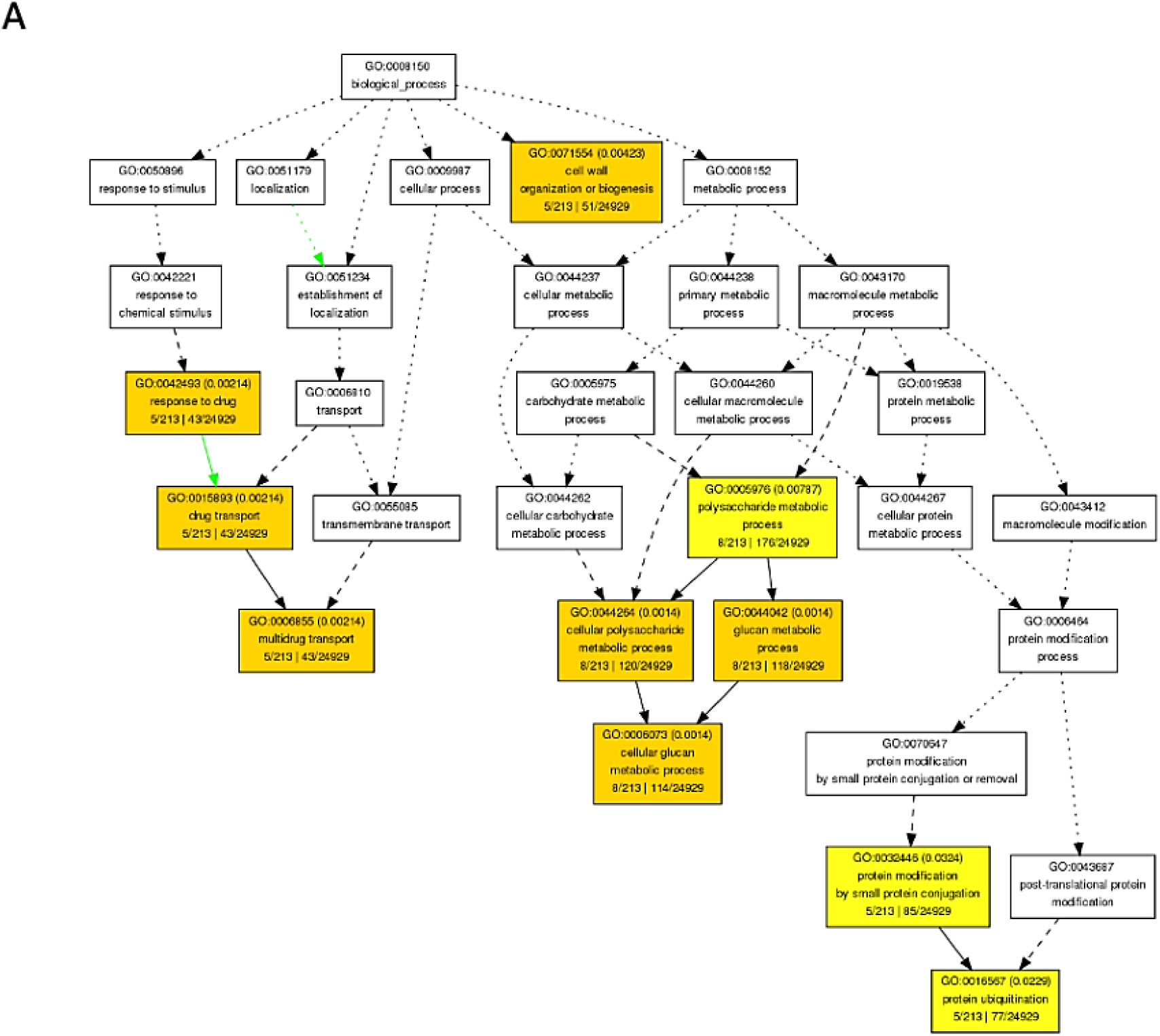
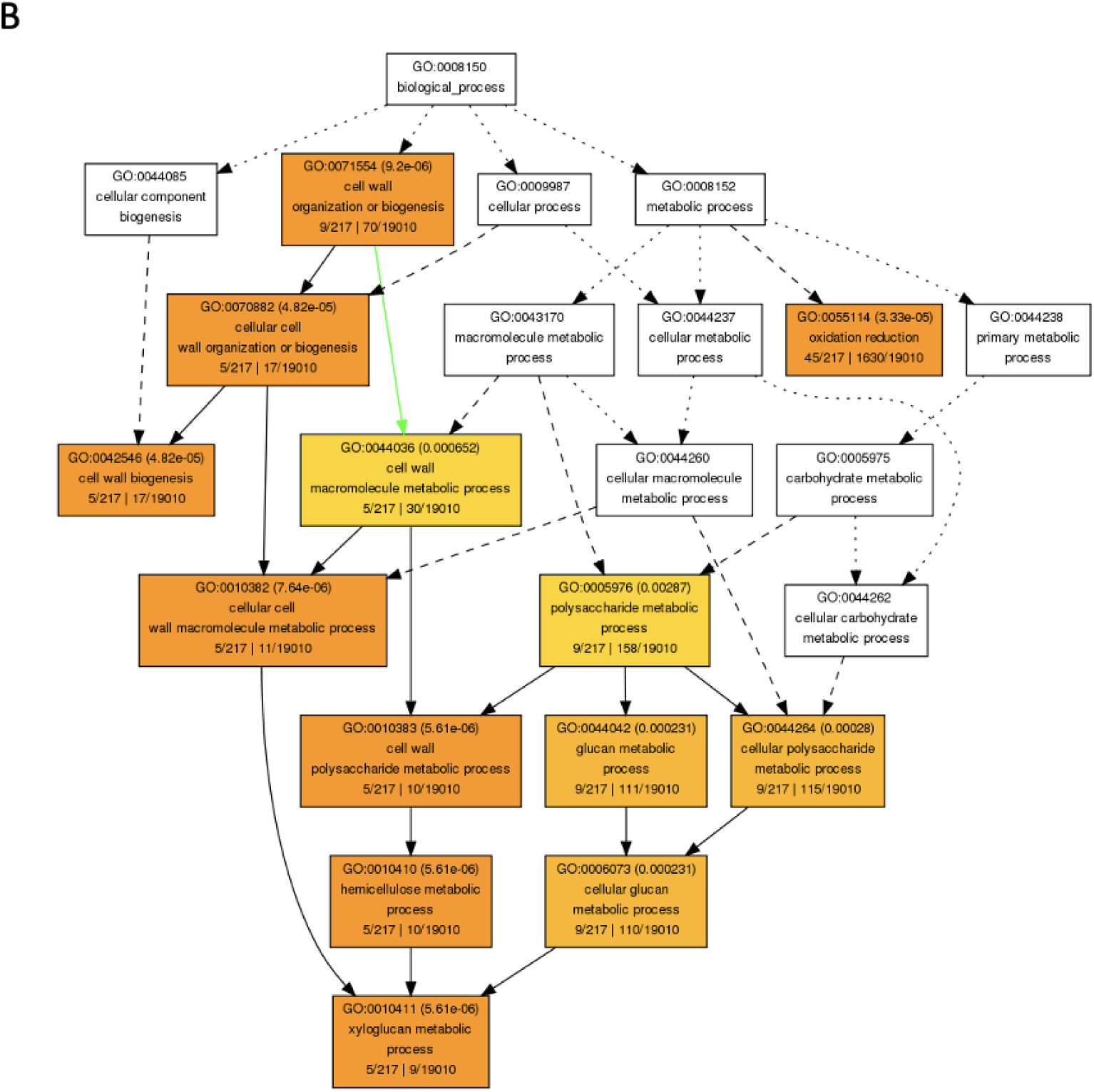
Supplementary Figure F2. GO enrichment networks of DEGs in *Cornus* transcriptomes. **(A)** GO terms enriched in DEGs upregulated in *C. florida*. (**B**) GO terms enriched in DEGs upregulated in *C. kousa*. See Supplementary Tables 7 and 8 for full term lists.

**Supplementary Figure 3.**
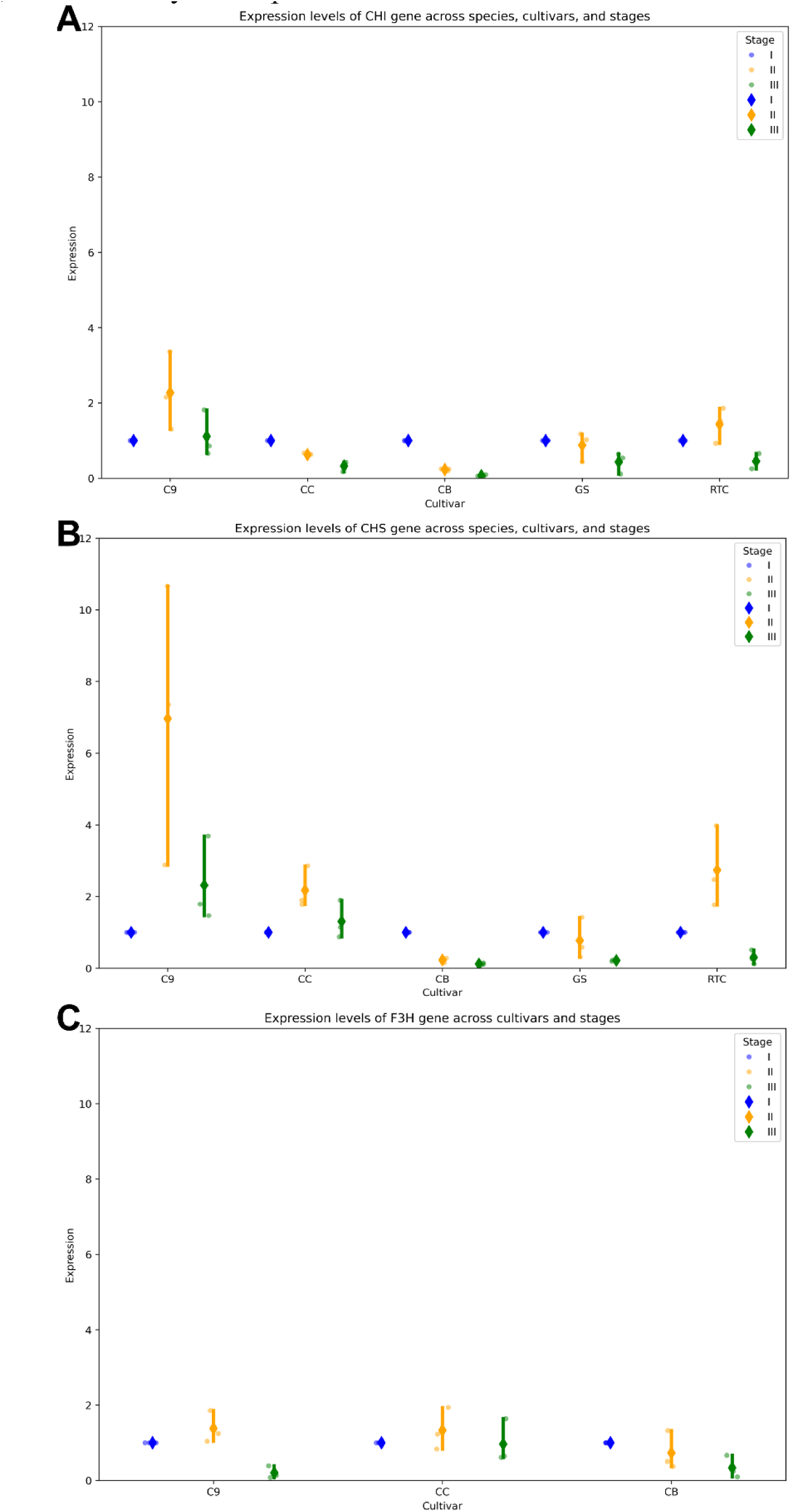
RT-qPCR validation of structural gene expression. Expression of (**A**) CHI, (**B**) CHS, and (**C**) F3H in five cultivars across three developmental stages. Cultivars abbreviated: C9 = ‘Cloud Nine’, CB = ‘Cherokee Brave’, CC = ‘Cherokee Chief’, GS = ‘Greensleeves’, RTC = ‘Rosy Teacups’.

## Supplementary Tables

**Supplementary Table T1.** Metadata for RNA-seq samples used in this study, including species, phenotype, cultivar, SRA accession numbers, total reads, and mapping statistics to C. florida and C. kousa transcriptomes.

| Sample name | Organism | Phenotype | Genotype | SRA accession | Numbers of reads | Numbers of uniquely mapped to *Cornus florida* transcriptome (CFT) | Percentage of uniquely mapped reads to CFT | Numbers of uniquely mapped to *Cornus kousa* transcriptome (CKT) | Percentage of uniquely mapped reads to CKT |
| --- | --- | --- | --- | --- | --- | --- | --- | --- | --- |
| 1*1 | *Cornus florida* | red | Cherokee Brave | SRR21411472 | 34140879 | 24818316.5 | 72.69% | 27686788.5 | 81.10% |
| 1*2 | *Cornus florida* | red | Cherokee Brave | SRR21411471 | 35366608 | 20950536 | 59.24% | 24000797.5 | 67.86% |
| 1*3 | *Cornus florida* | red | Cherokee Chief | SRR21411467 | 31649684 | 23509163 | 74.28% | 24587183.5 | 77.69% |
| 1*6 | *Cornus florida* | white | Cloud Nine | SRR21411466 | 28399202 | 21460142.5 | 75.57% | 21425521 | 75.44% |
| 1*10 | *Cornus kousa* | light pink | Rosy Teacups | SRR21411465 | 29668132 | 22965778.5 | 77.41% | 23004720.5 | 77.54% |
| 1*13 | *Cornus kousa* | light green | Greensleeves | SRR21411464 | 33082227 | 26184769 | 79.15% | 25745124.5 | 77.82% |
| 2*1 | *Cornus florida* | red | Cherokee Chief | SRR21411463 | 35782681 | 24181390 | 67.58% | 24398398.5 | 68.18% |
| 2*2 | *Cornus florida* | red | Cherokee Chief | SRR21411462 | 34453910 | 26047989.5 | 75.60% | 25499764.5 | 74.01% |
| 2*3 | *Cornus florida* | white | Cloud Nine | SRR21411461 | 28919895 | 23053520.5 | 79.72% | 22646851.5 | 78.31% |
| 2*4 | *Cornus florida* | white | Cloud Nine | SRR21411460 | 37510289 | 26976299 | 71.92% | 26287991 | 70.08% |
| 2*5 | *Cornus kousa* | light pink | Rosy Teacups | SRR21411470 | 38487414 | 25455388.5 | 66.14% | 28869402.5 | 75.01% |
| 2*5 | *Cornus kousa* | light pink | Rosy Teacups | SRR21411469 | 39321965 | 23799064.5 | 60.52% | 27216224.5 | 69.21% |
| 2*8 | *Cornus kousa* | light green | Greensleeves | SRR21411468 | 30532090 | 18797761 | 61.57% | 21492150.5 | 70.39% |

**Supplementary Table T2.**
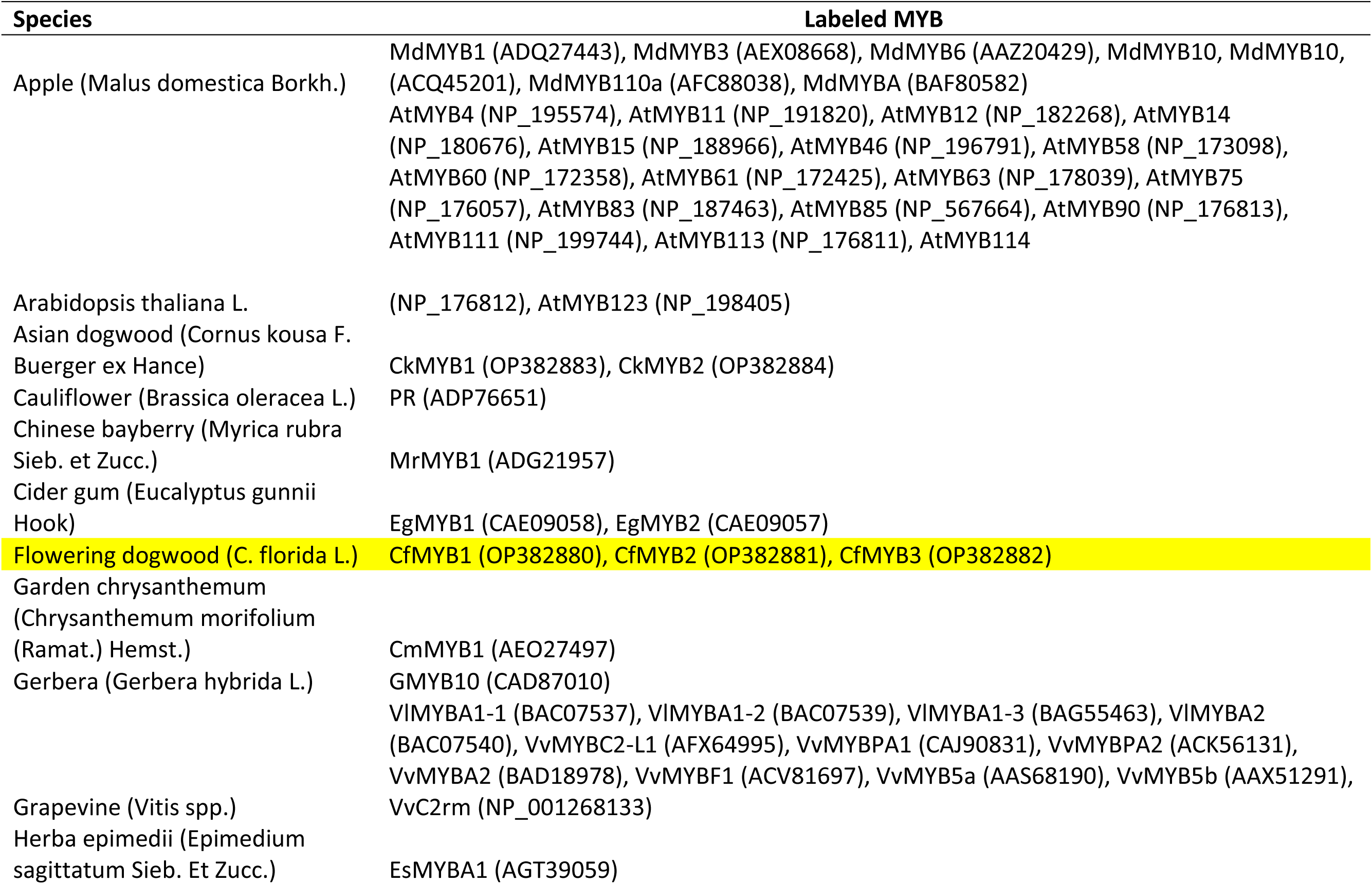
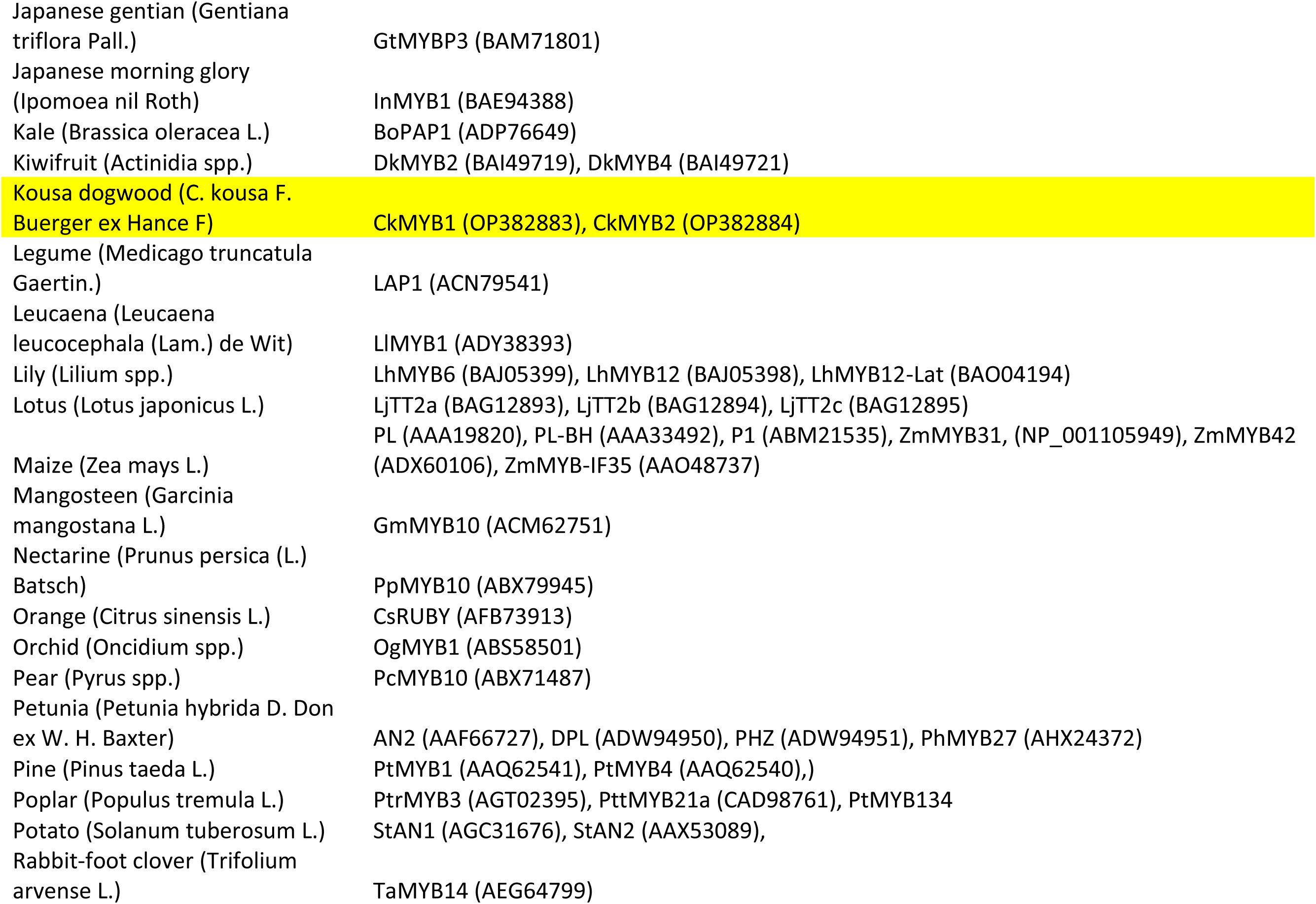
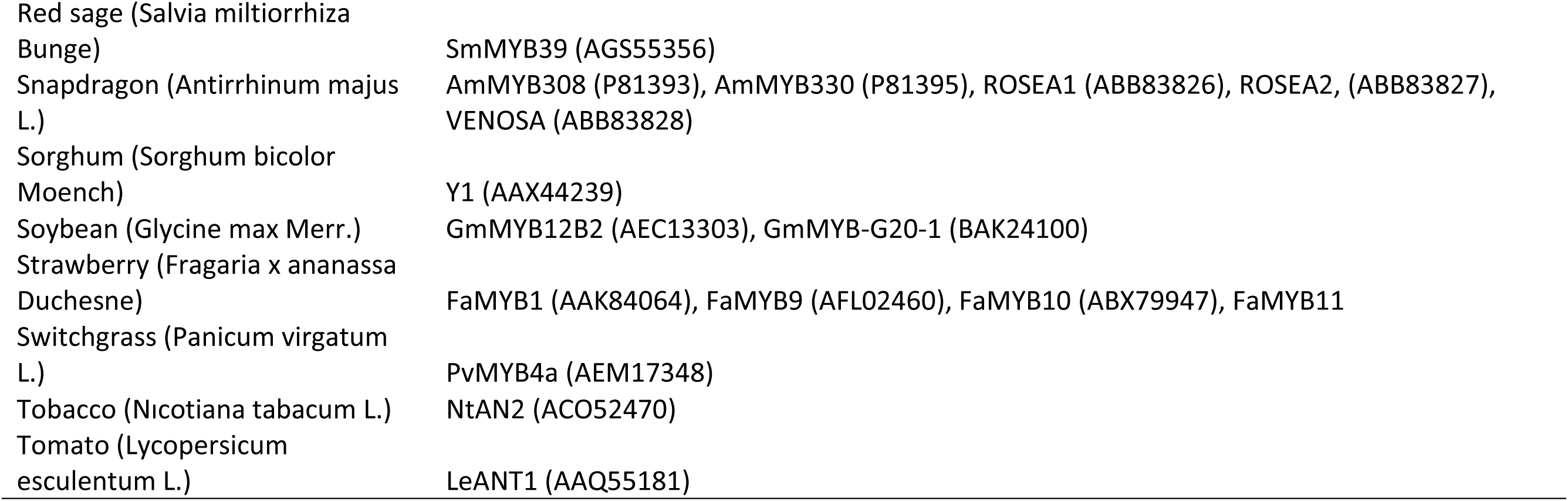
Species names and GenBank accession numbers of MYB transcription factors used for phylogenetic analysis of R2R3-MYB proteins.

**Supplementary Table T3.**
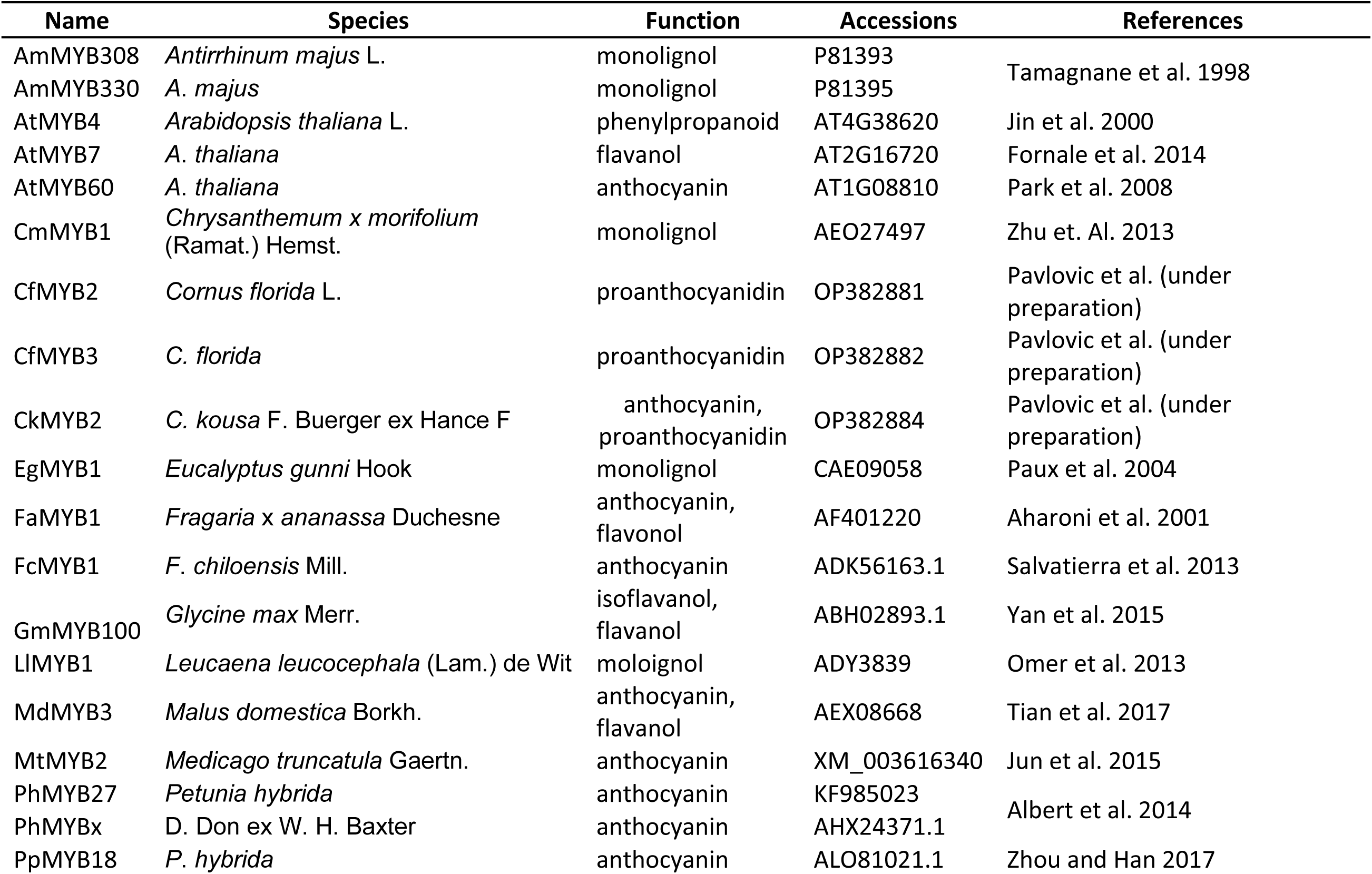
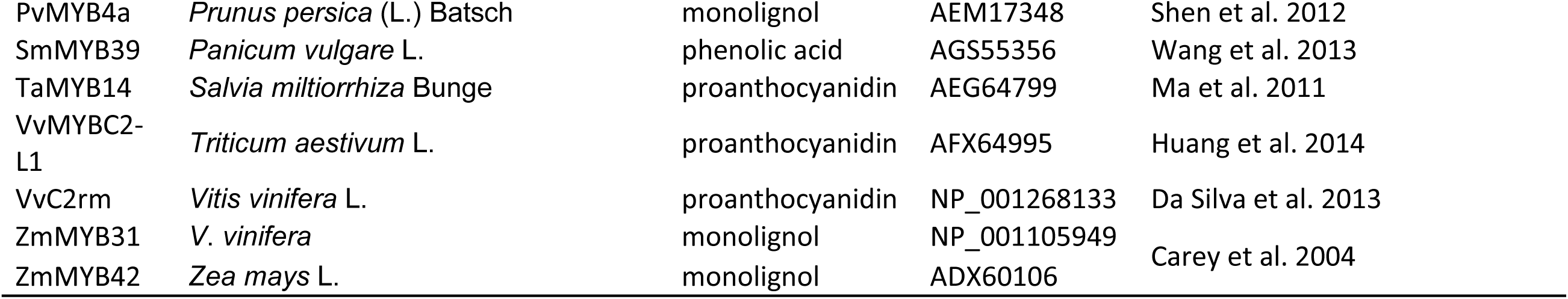
R2R3-MYB repressor proteins used for EAR motif analysis, including species, gene function, accession numbers, and references.

**Supplementary Table T4.**
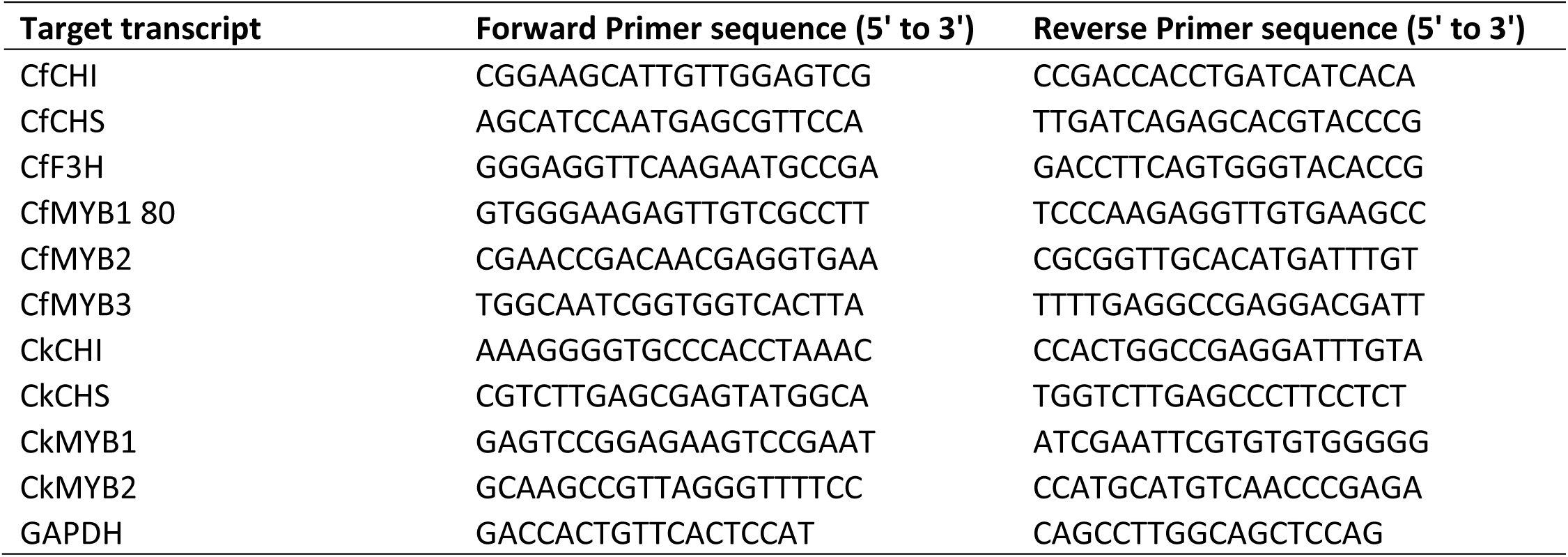
Primer sequences used for RT-qPCR validation of candidate MYB and structural genes. Includes target gene, forward and reverse primer sequences, expected amplicon size, and transcriptome contig IDs.

**Supplementary Table T5.**
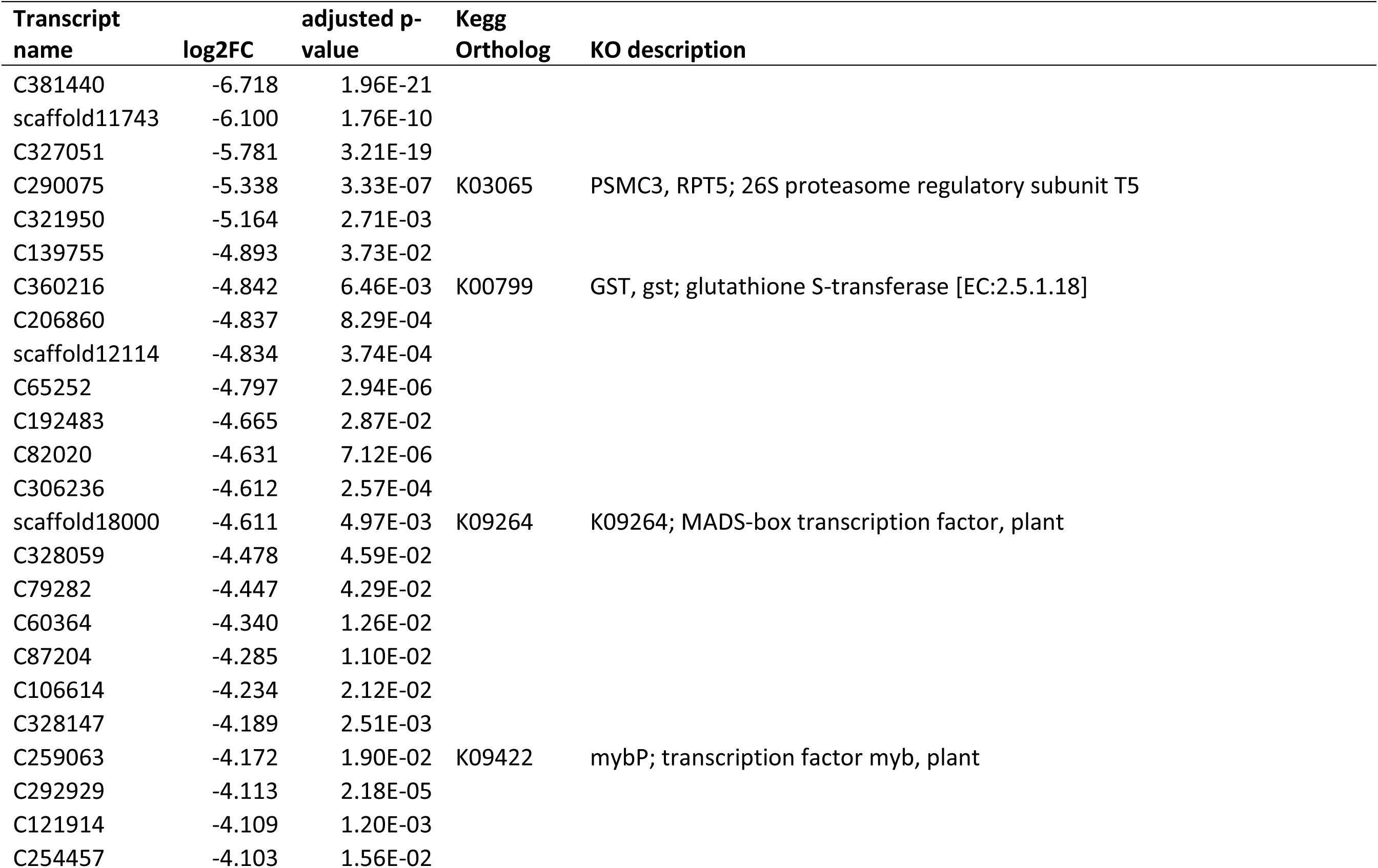
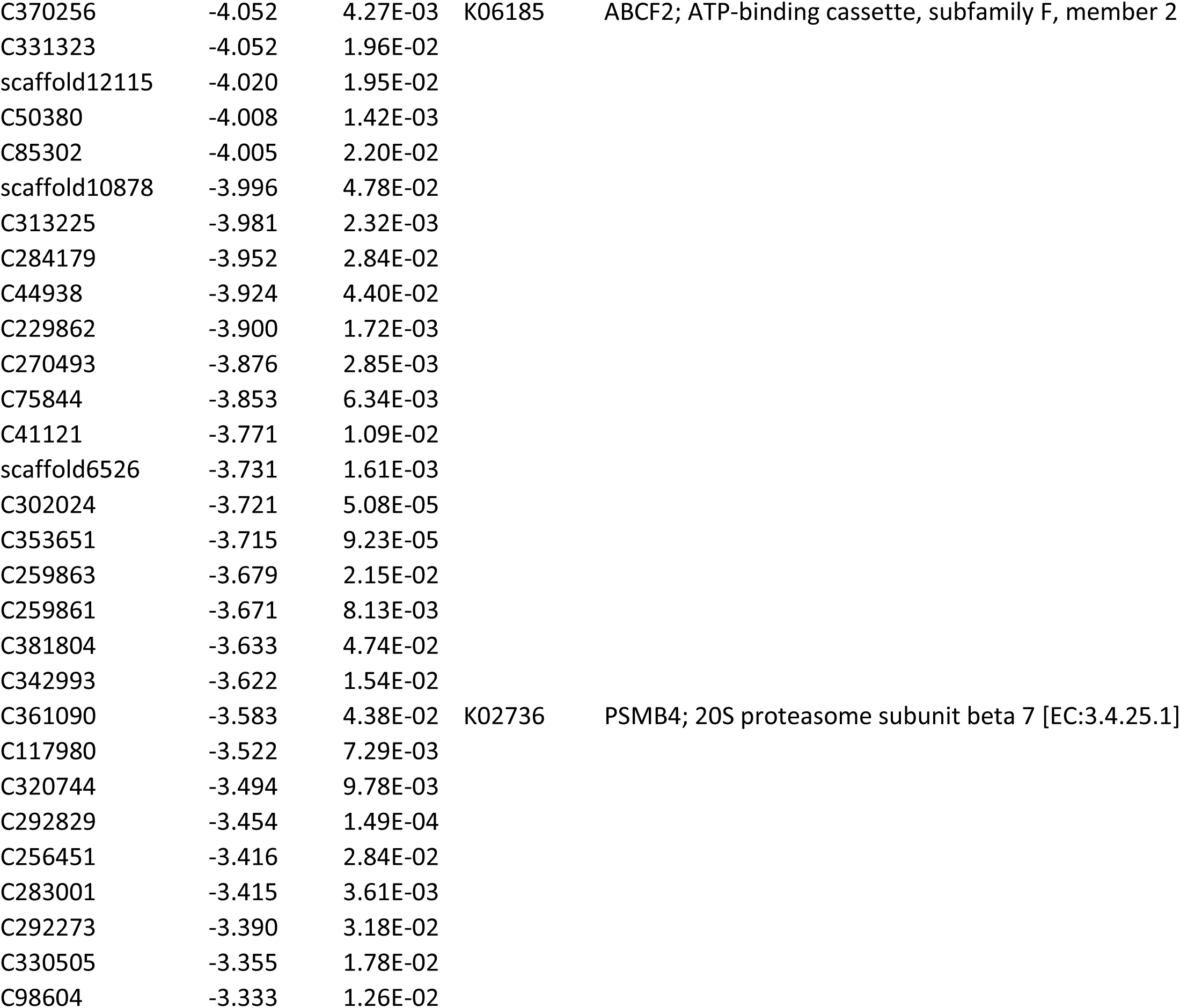
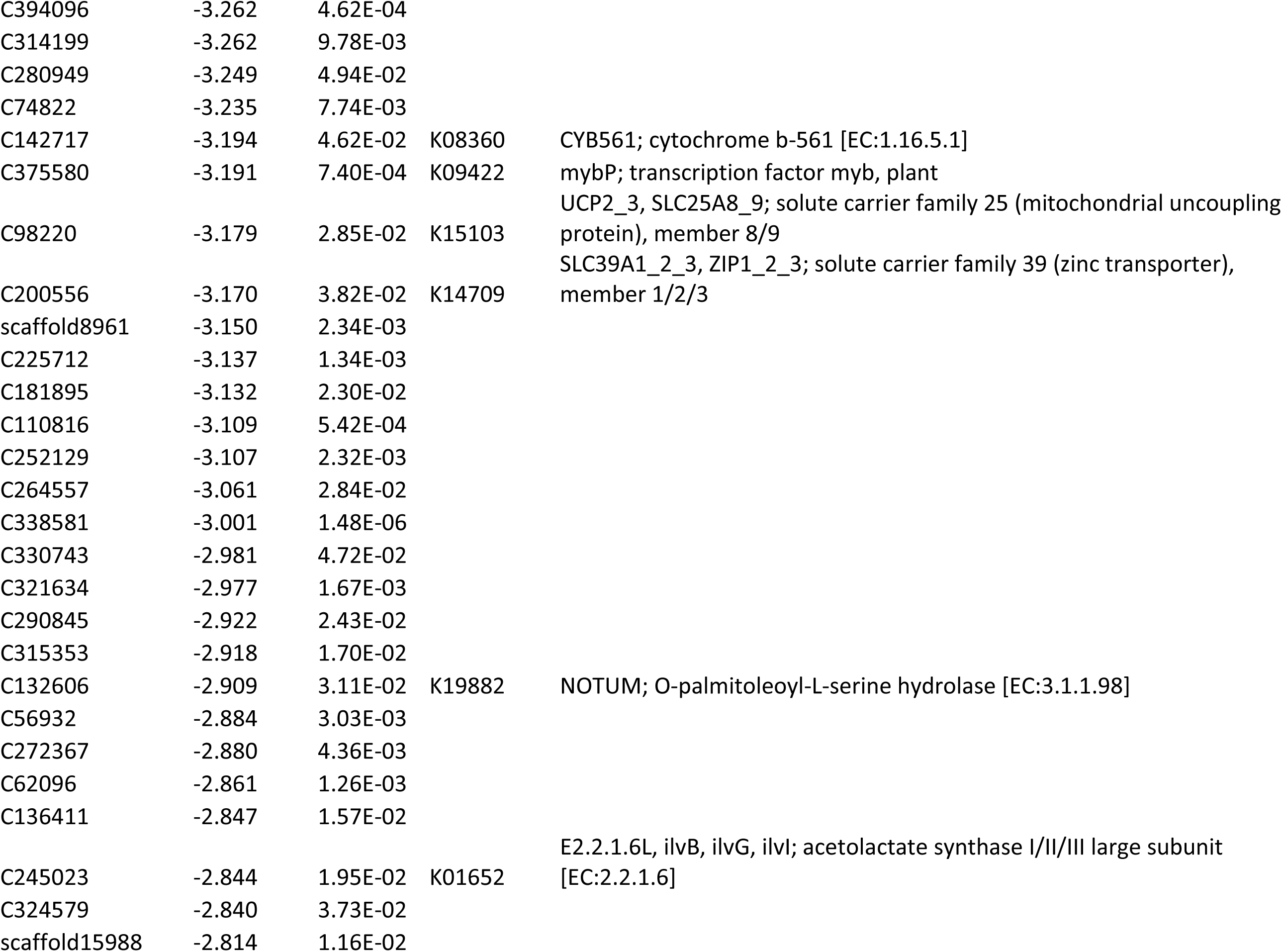
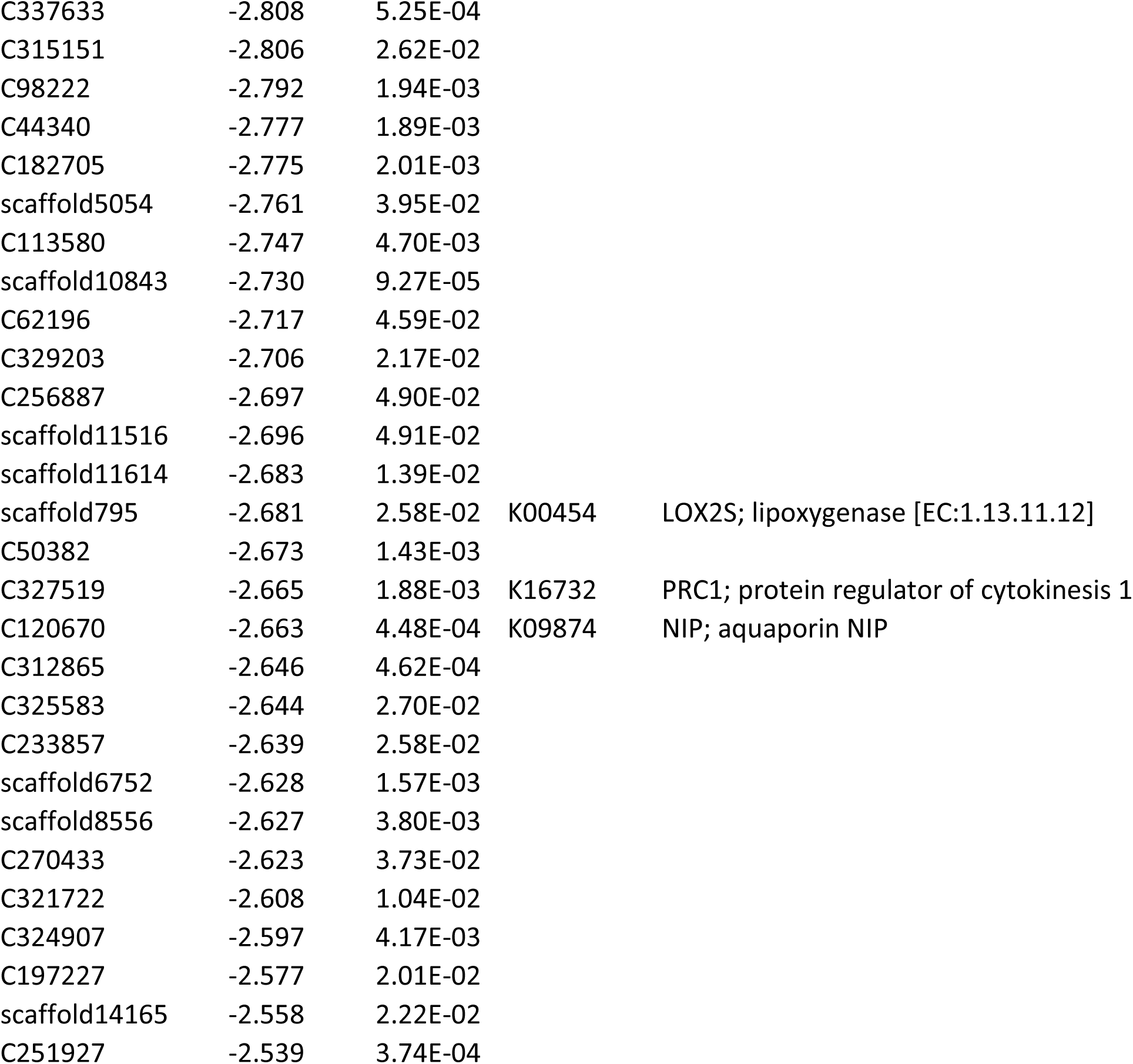
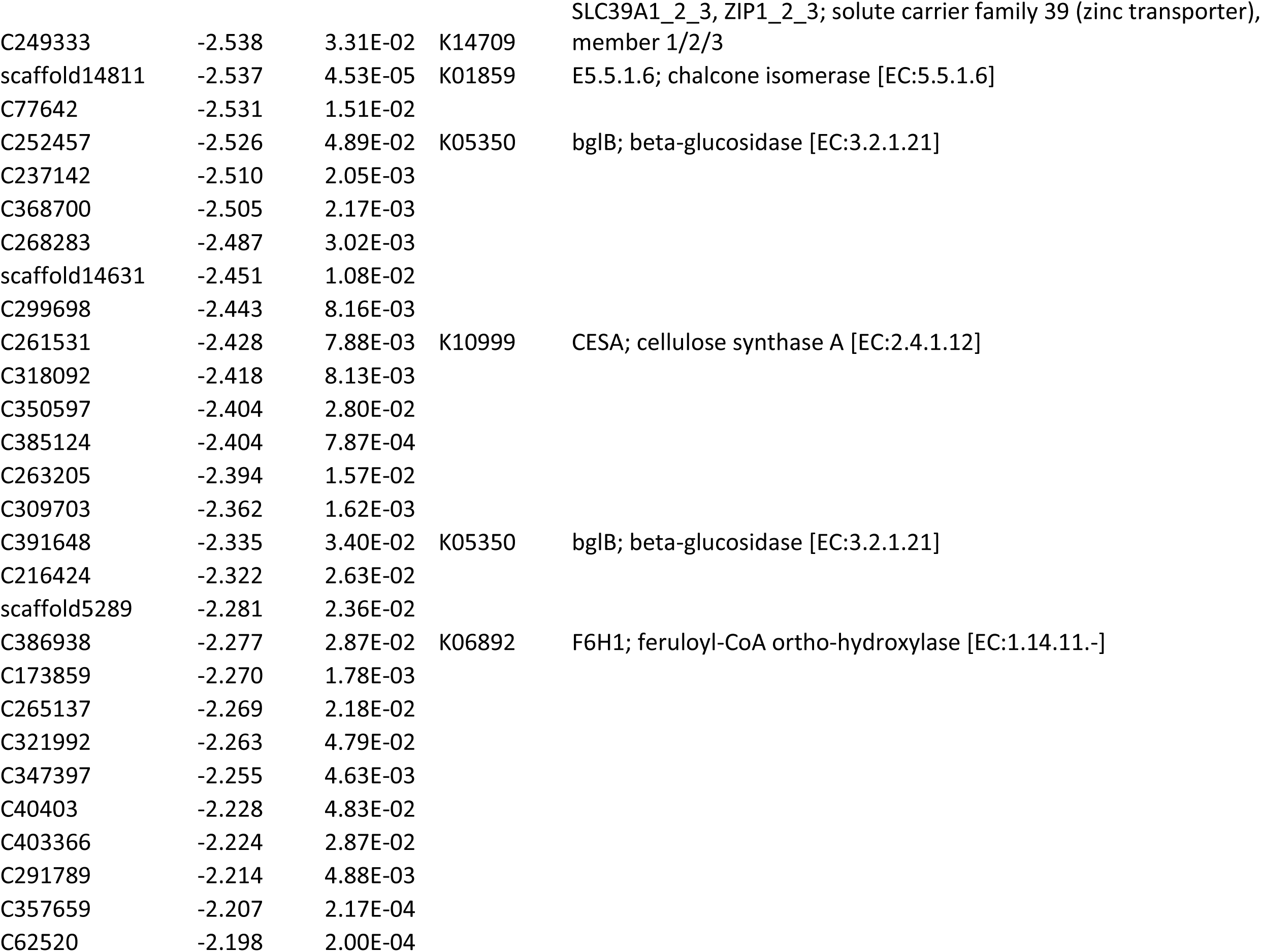
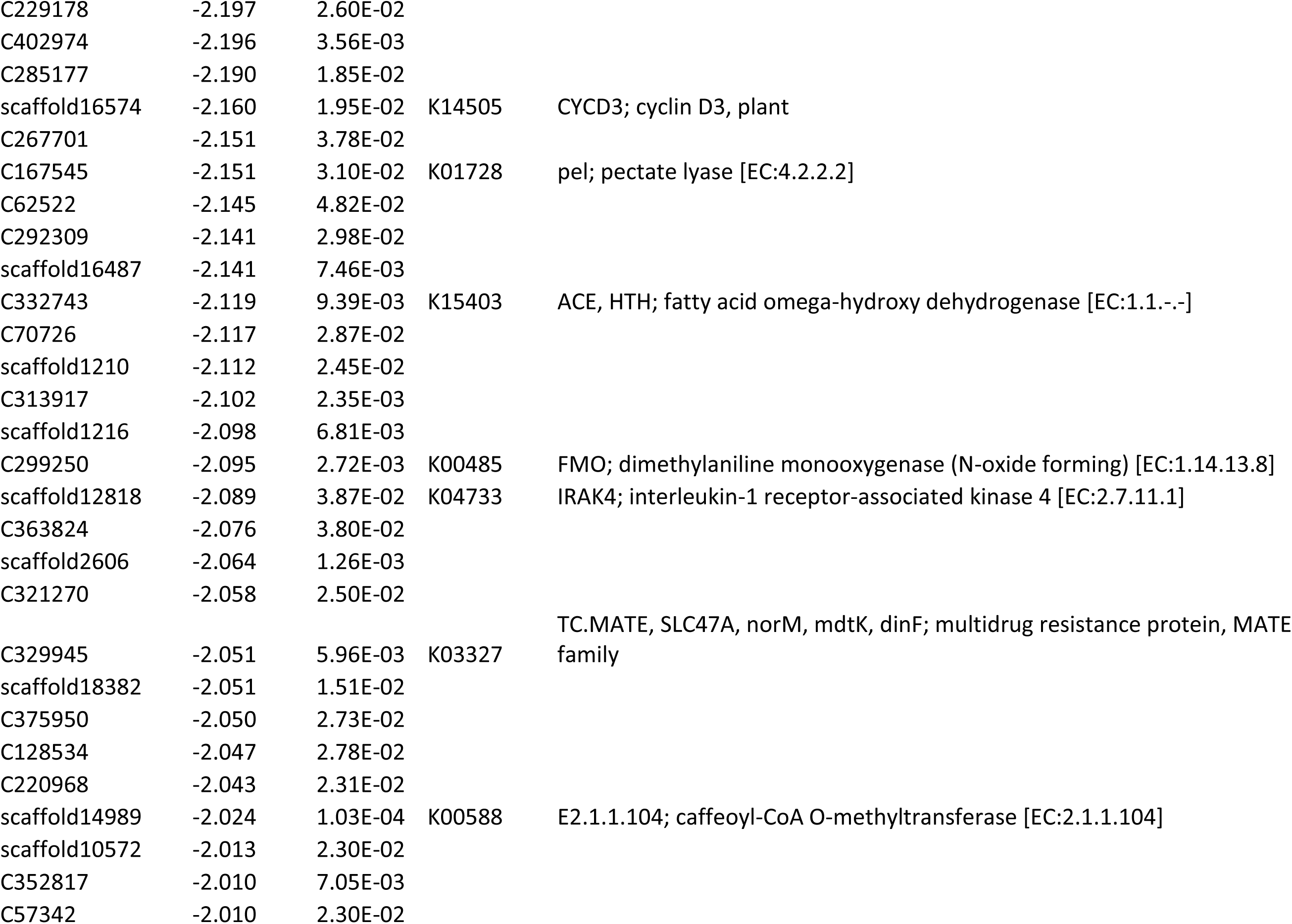
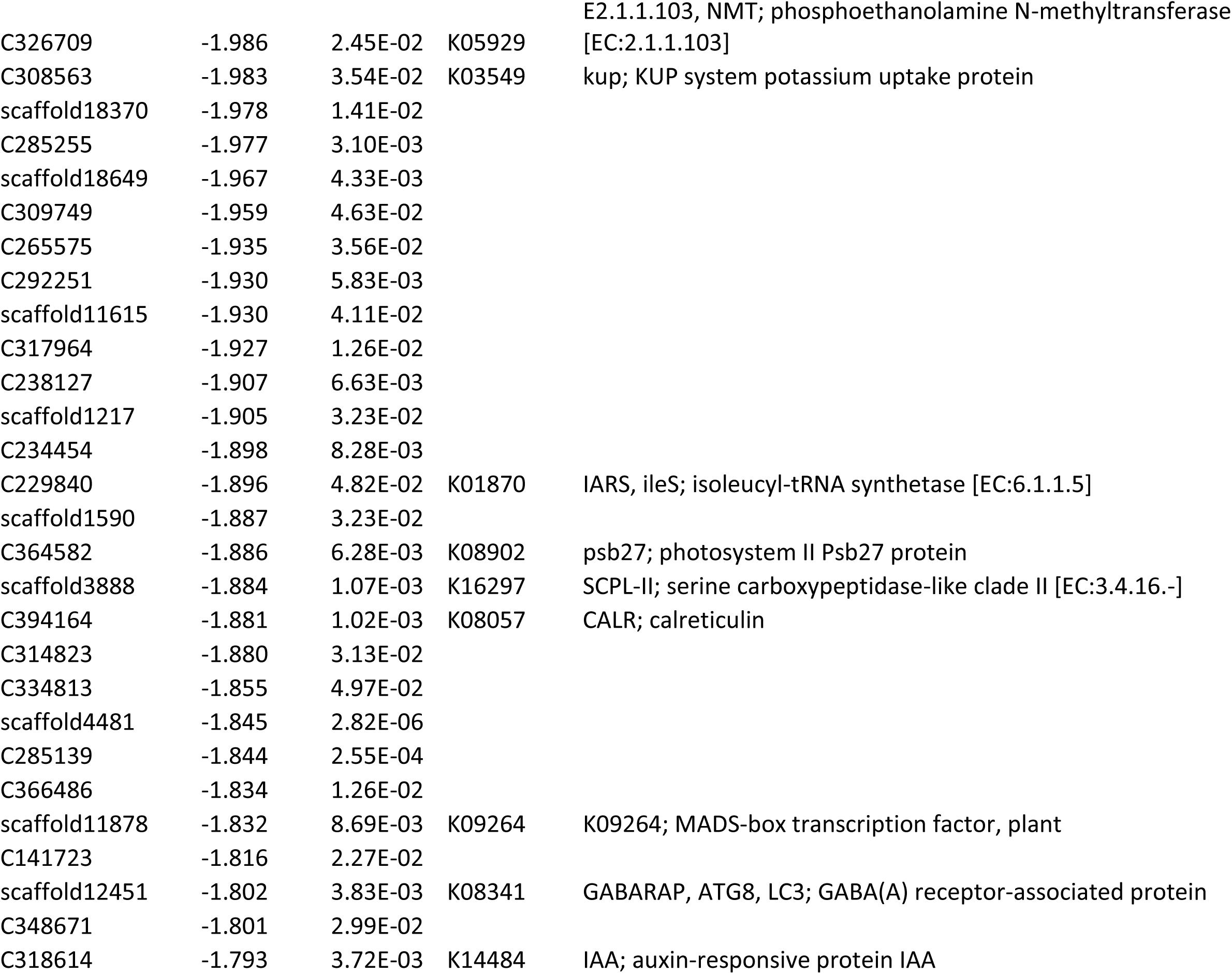
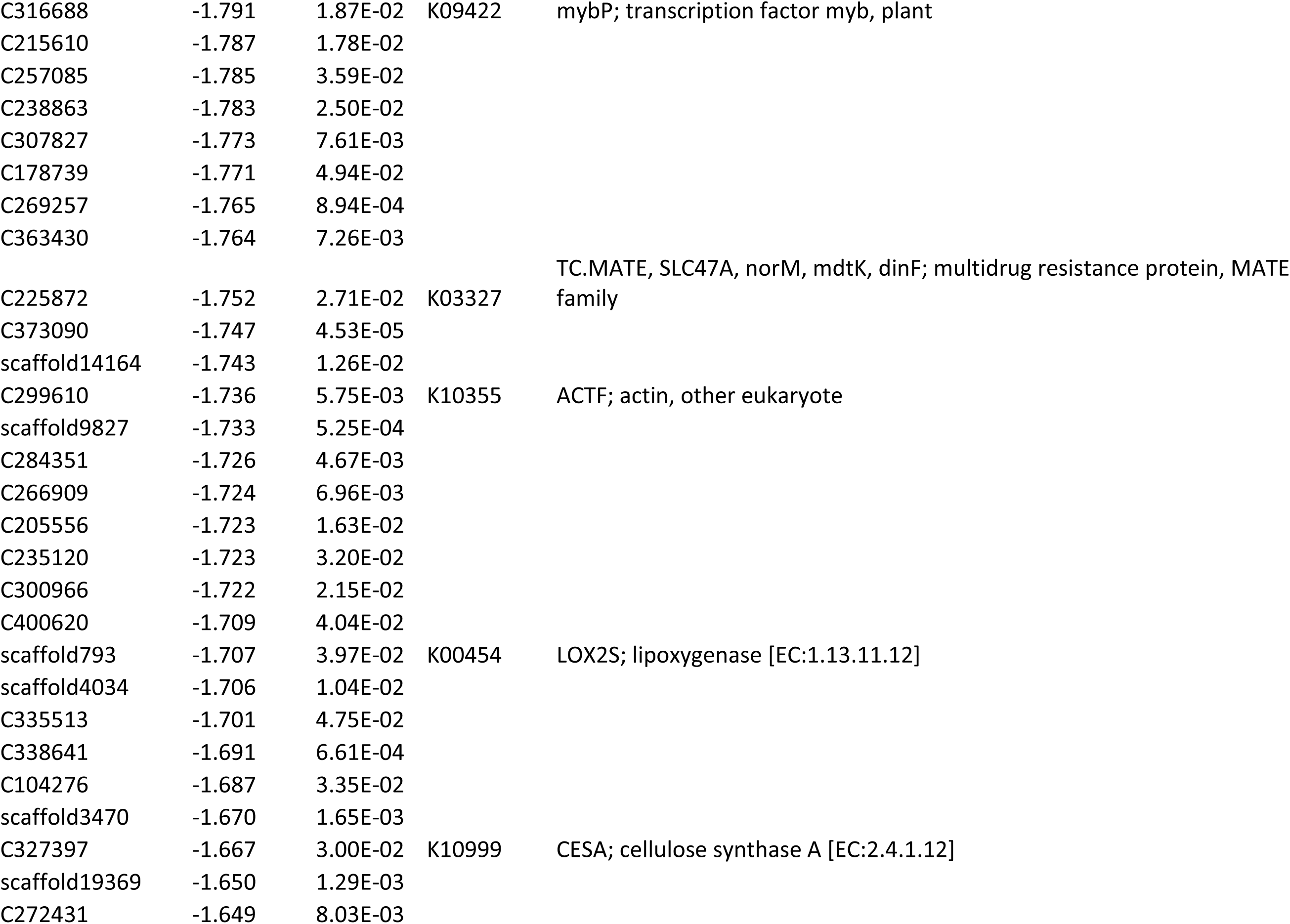
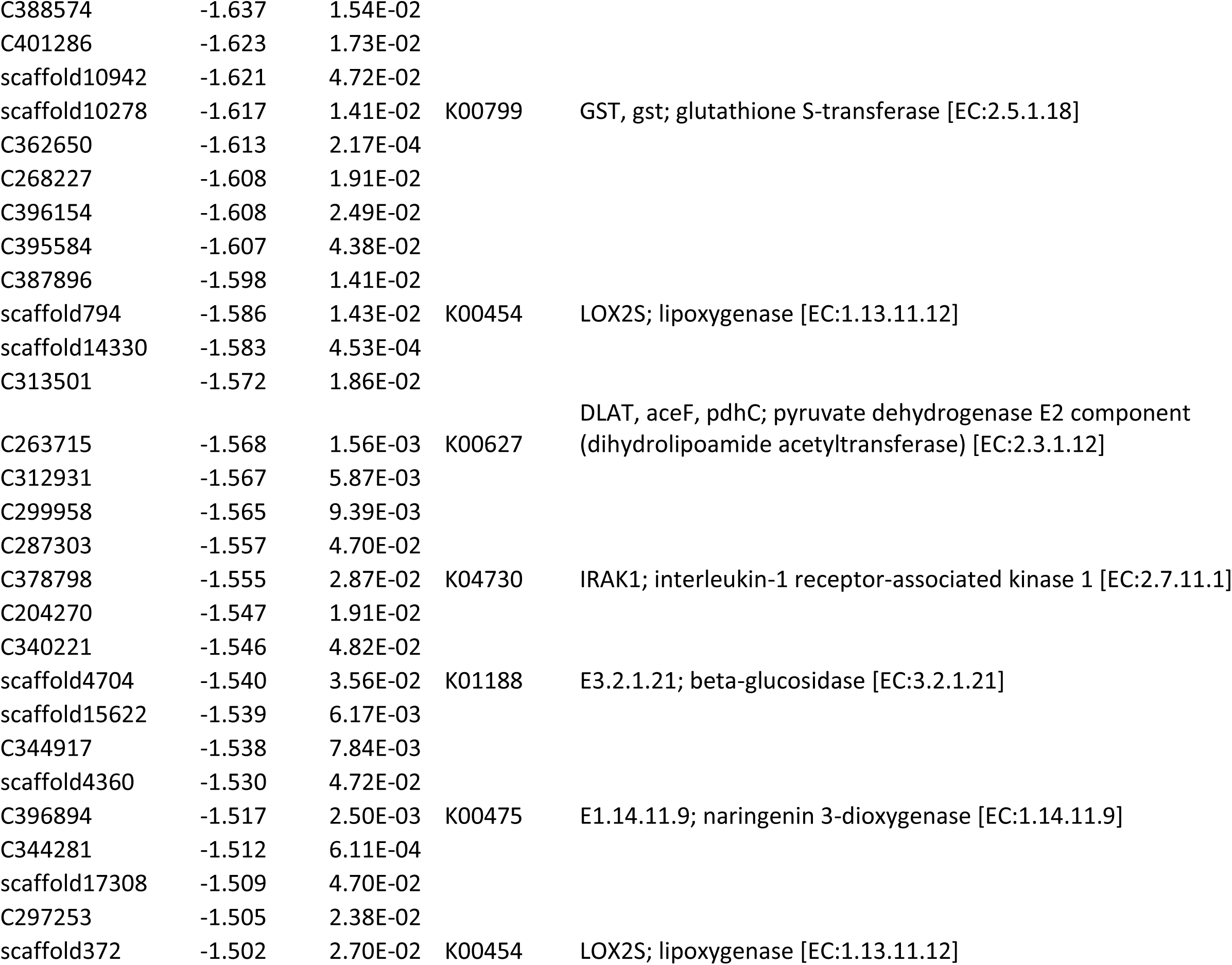
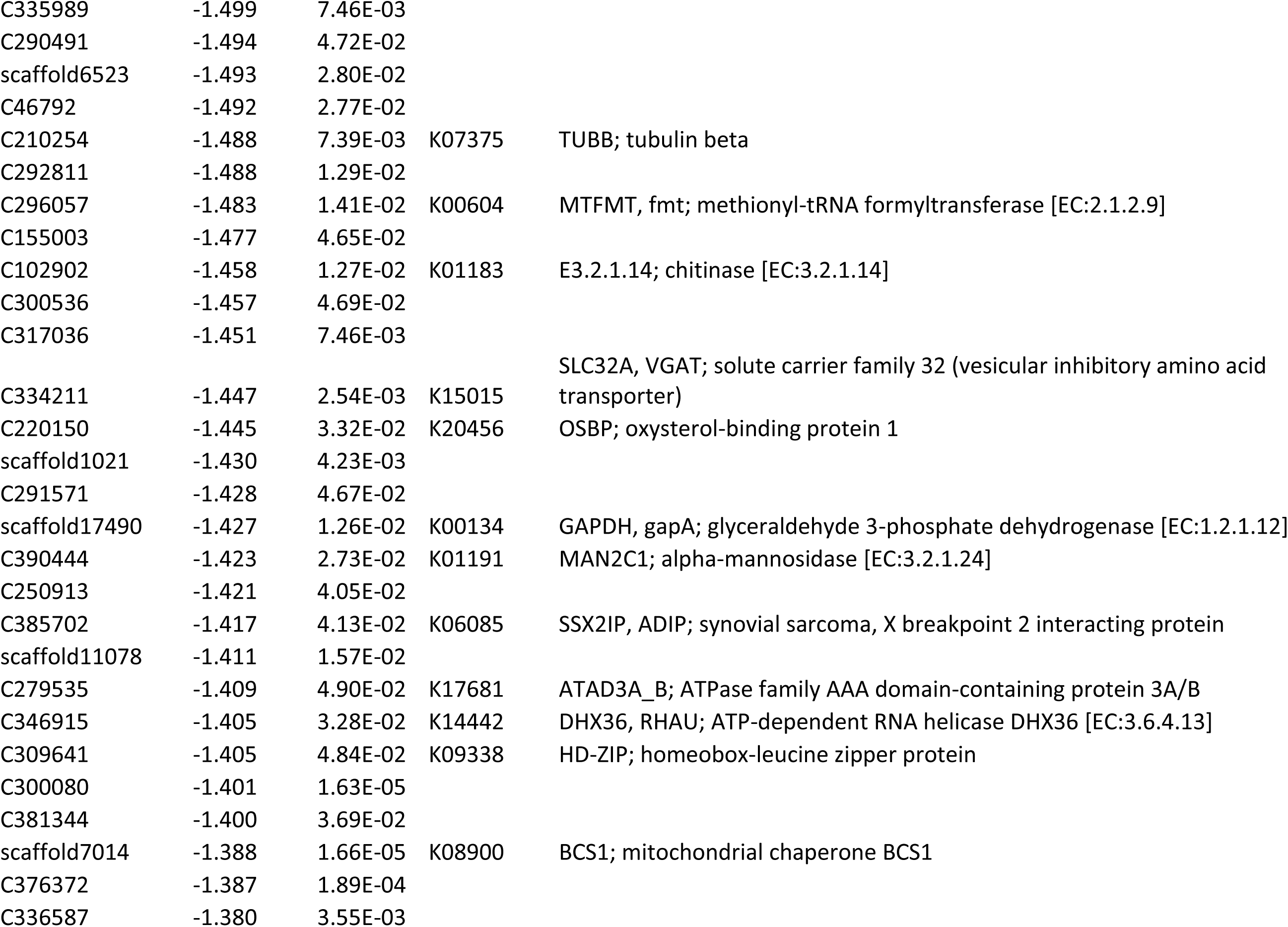
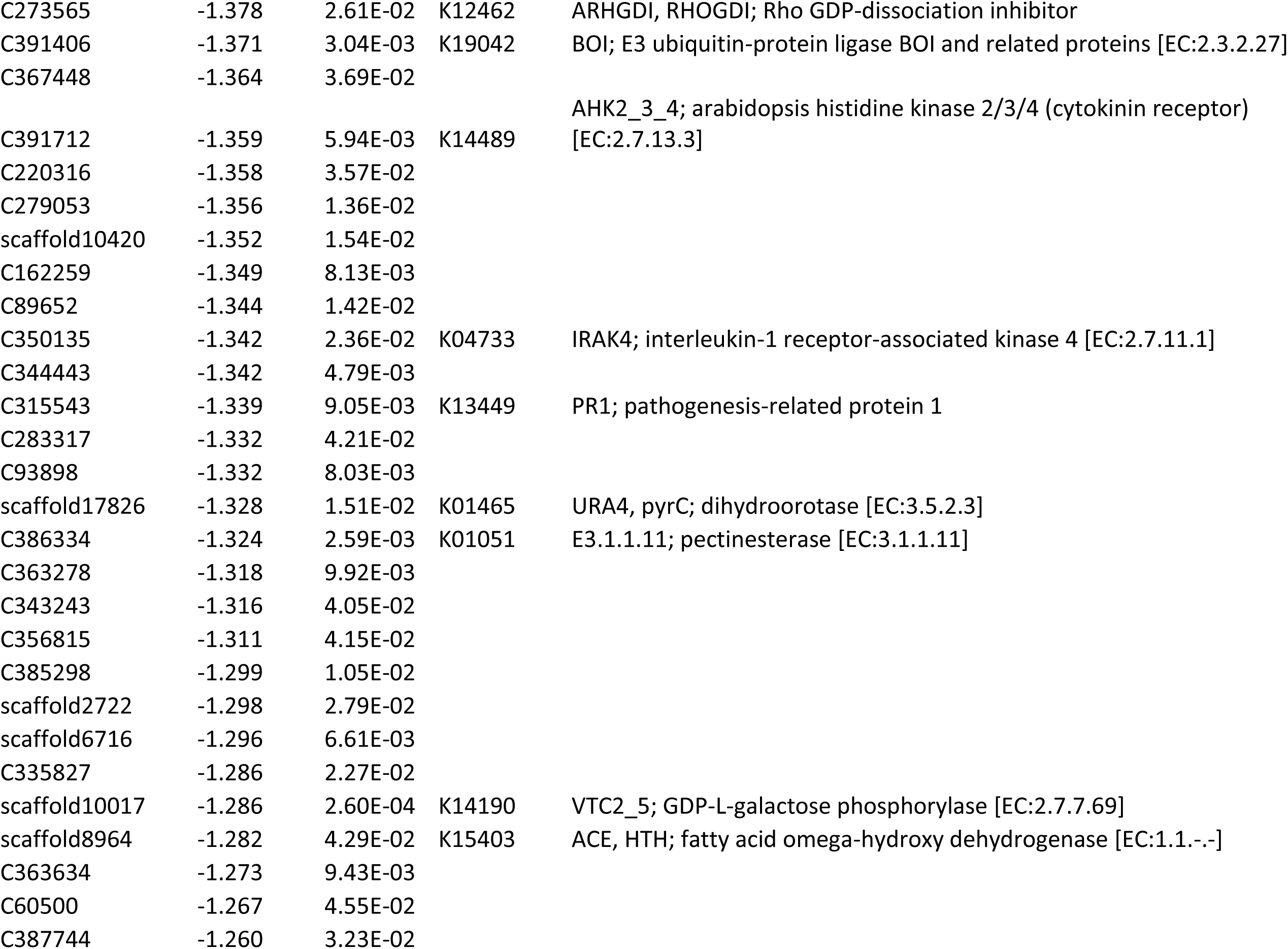
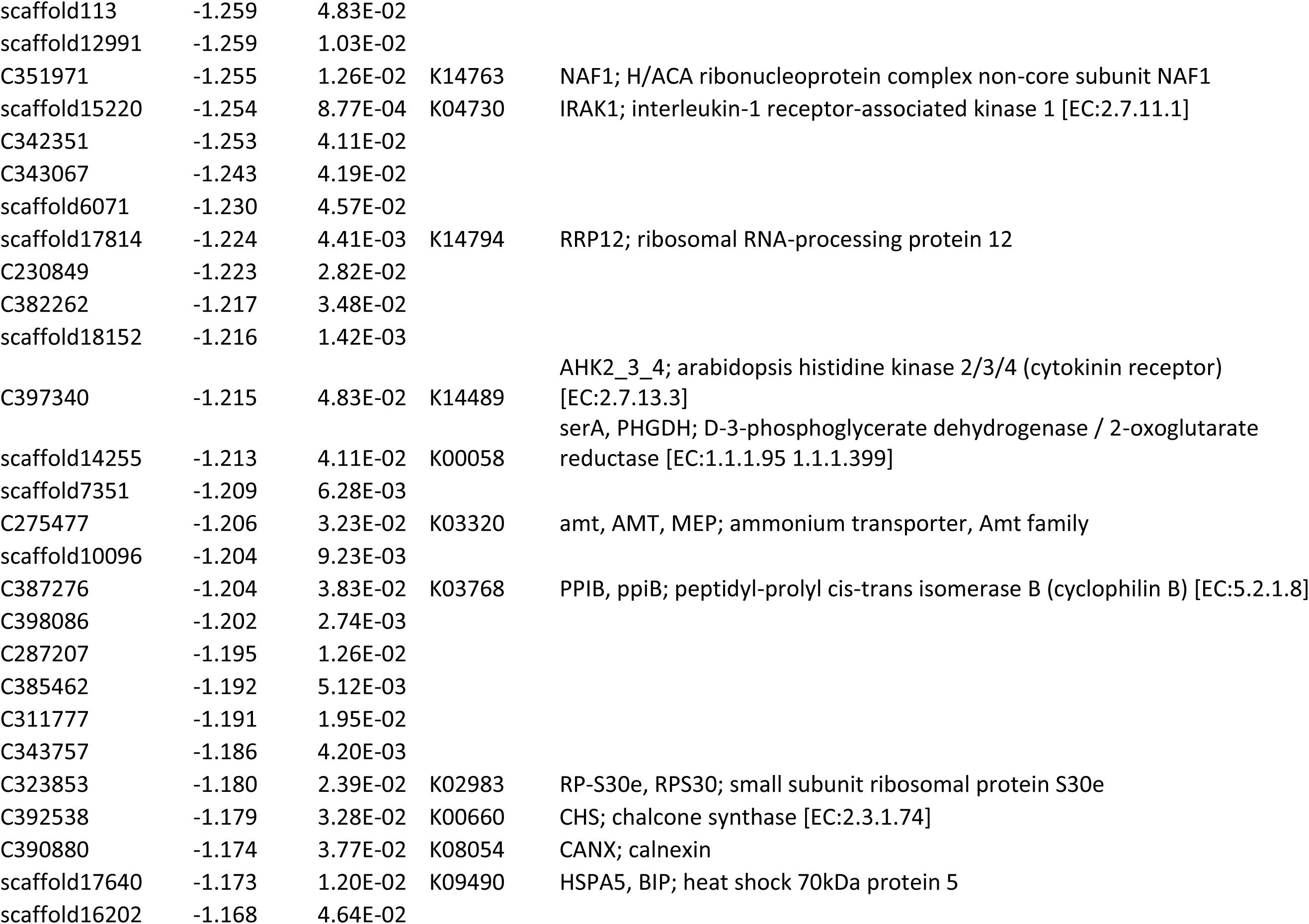
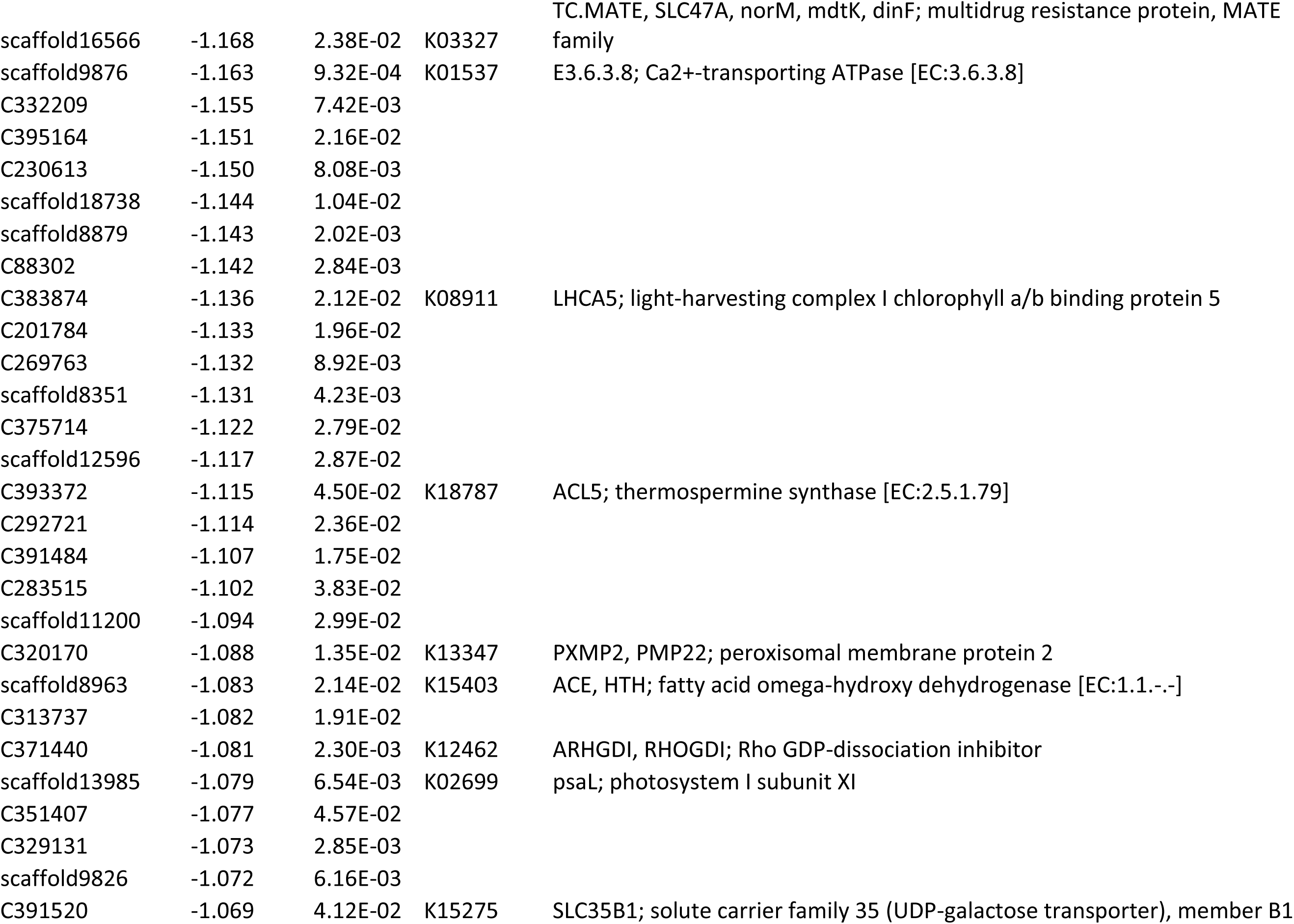
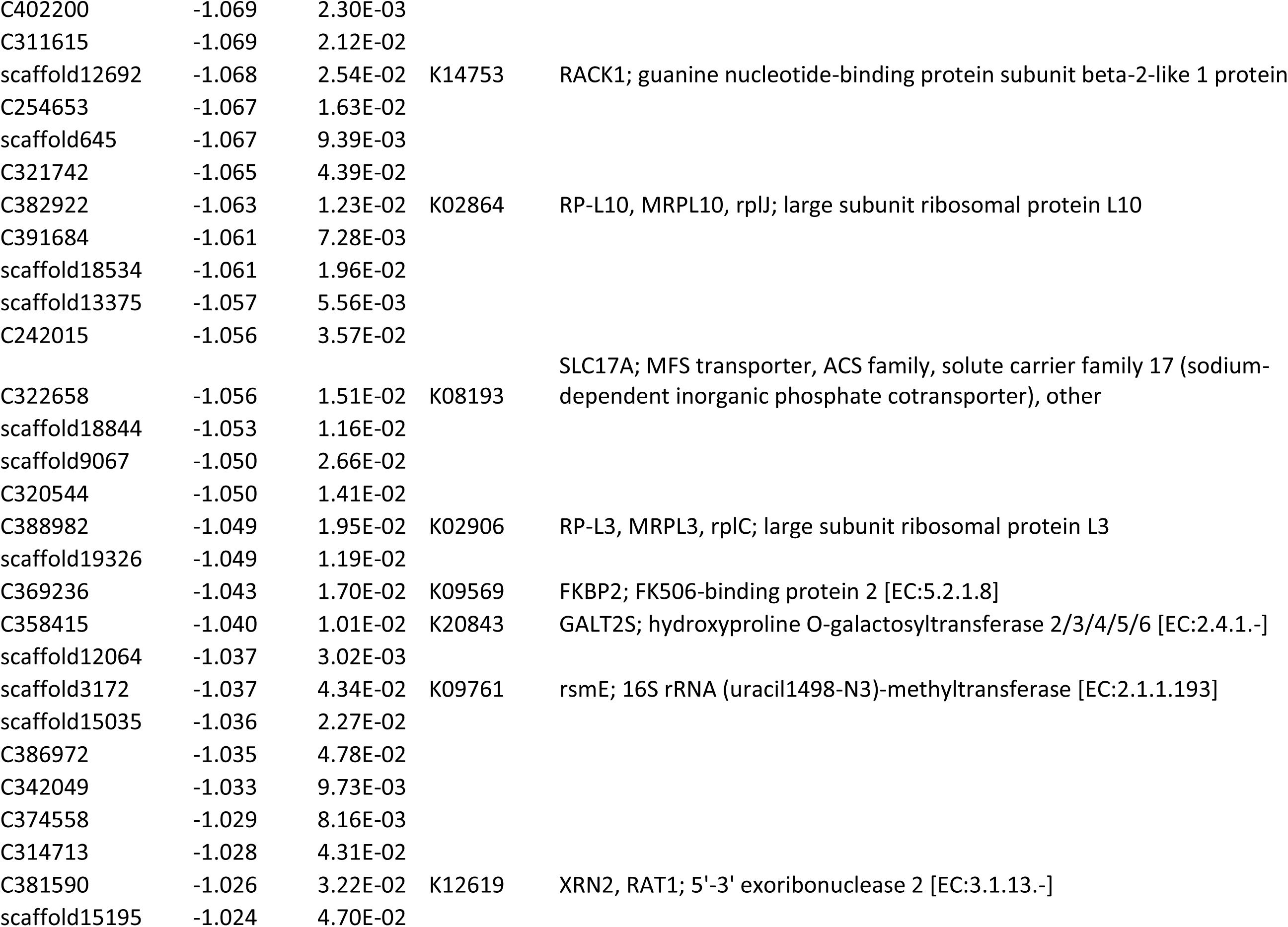
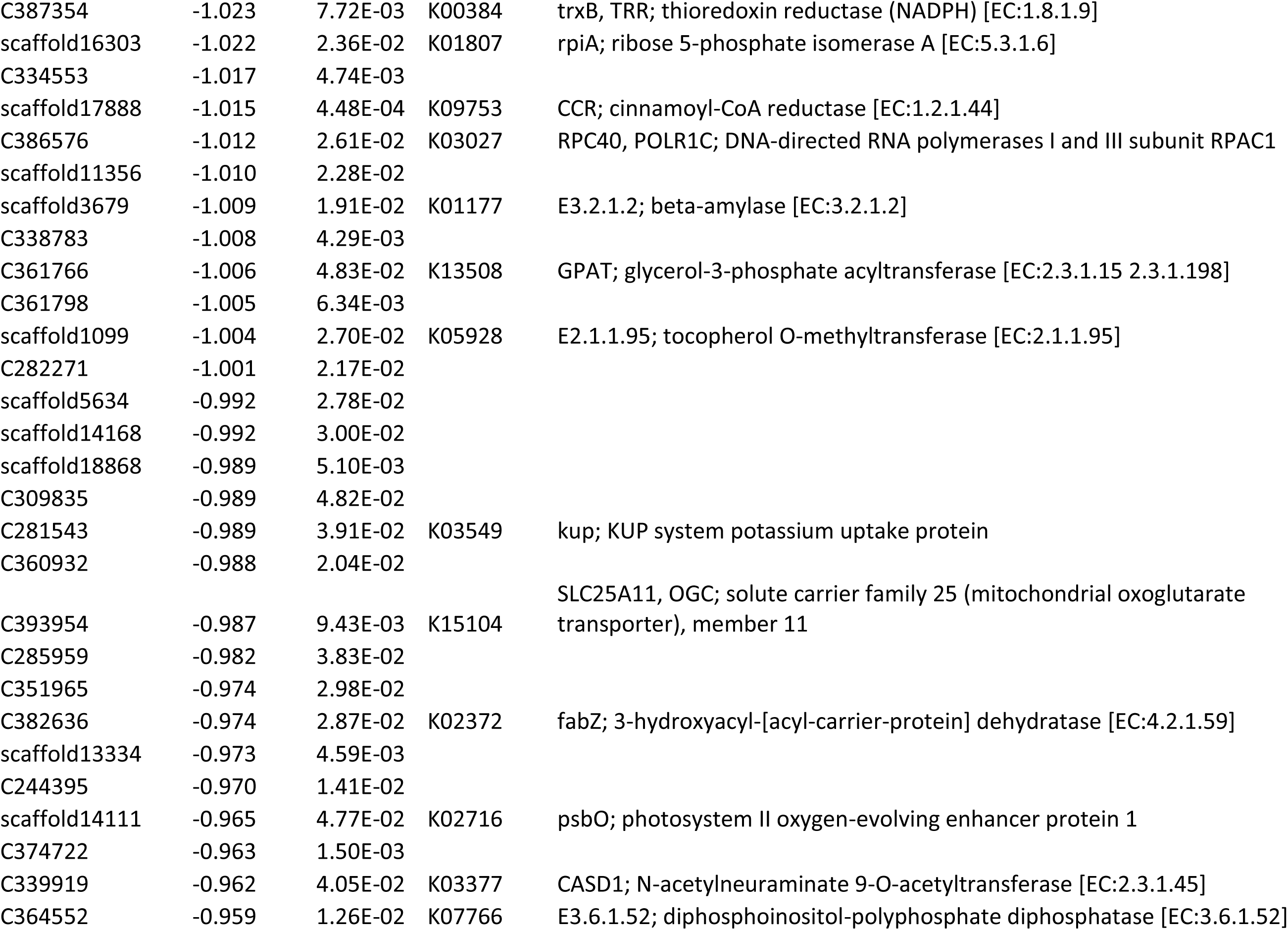
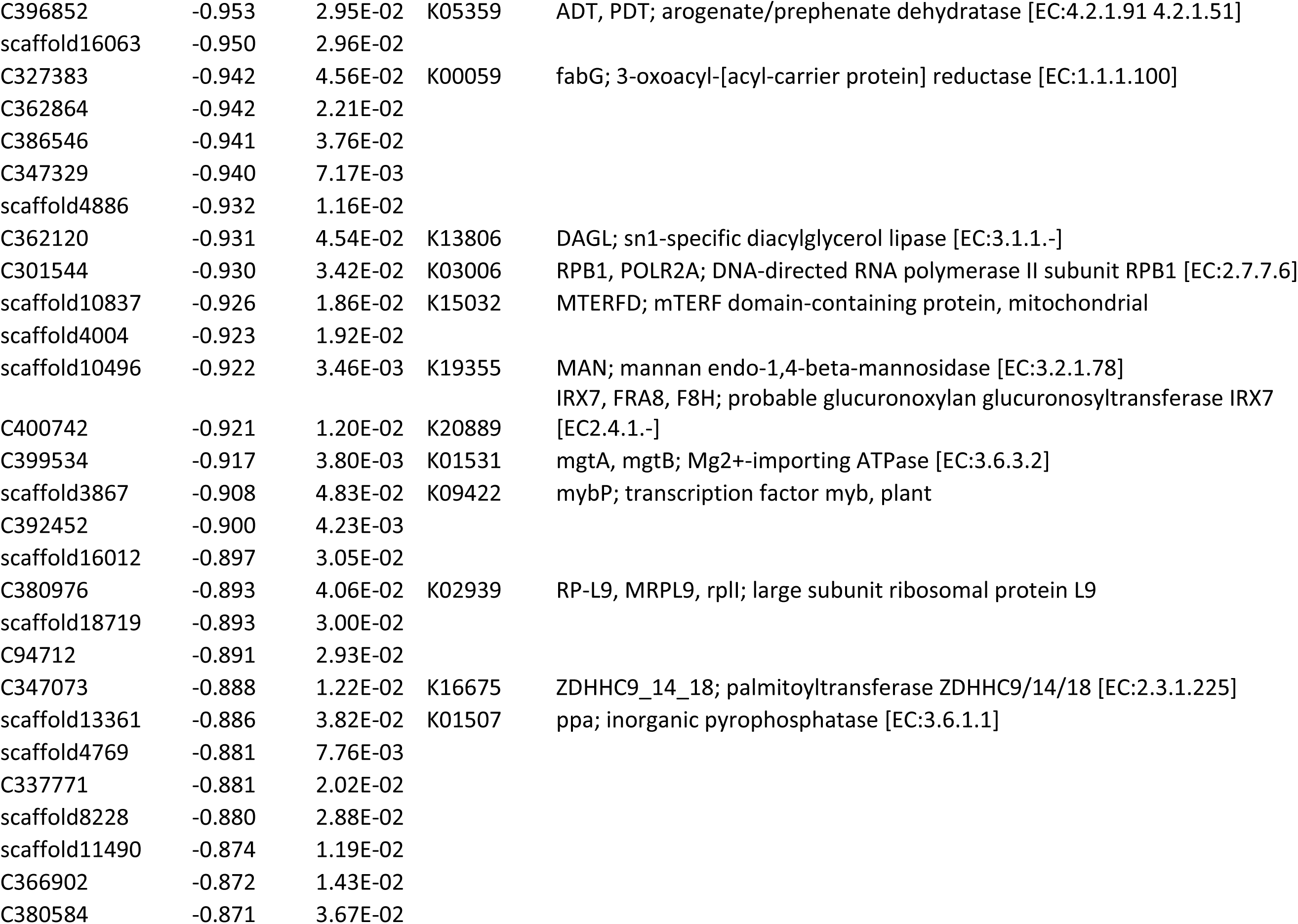
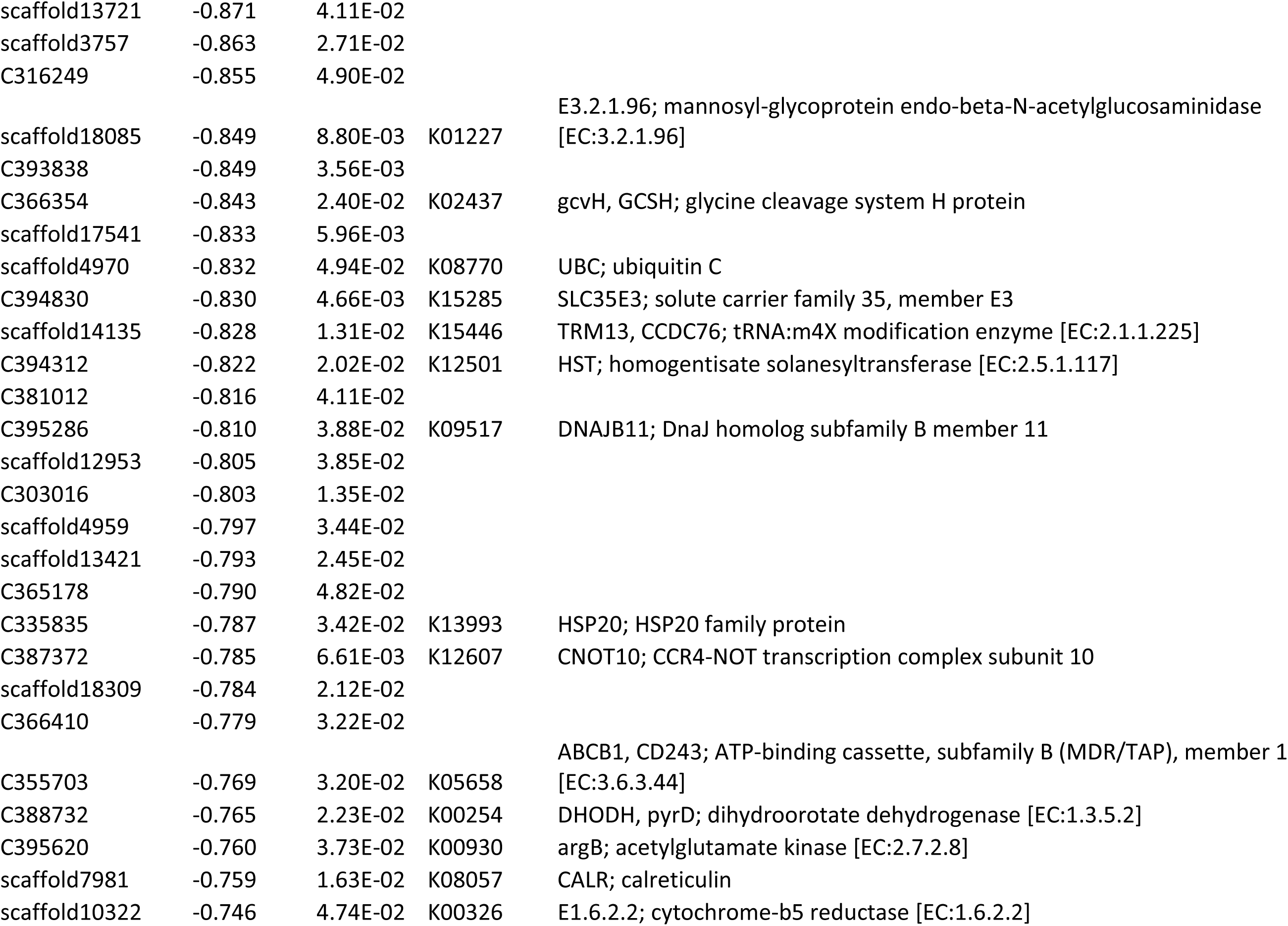
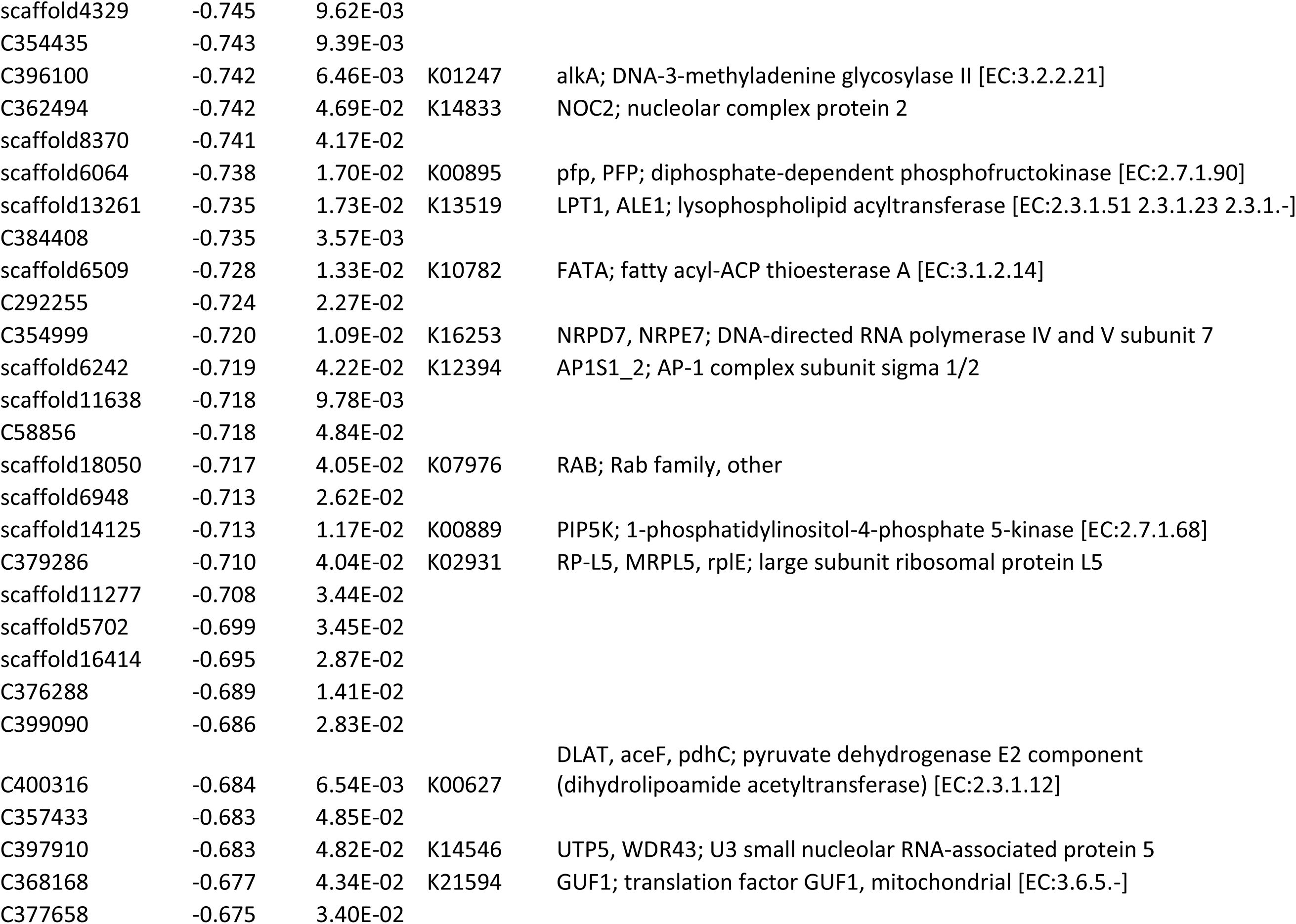
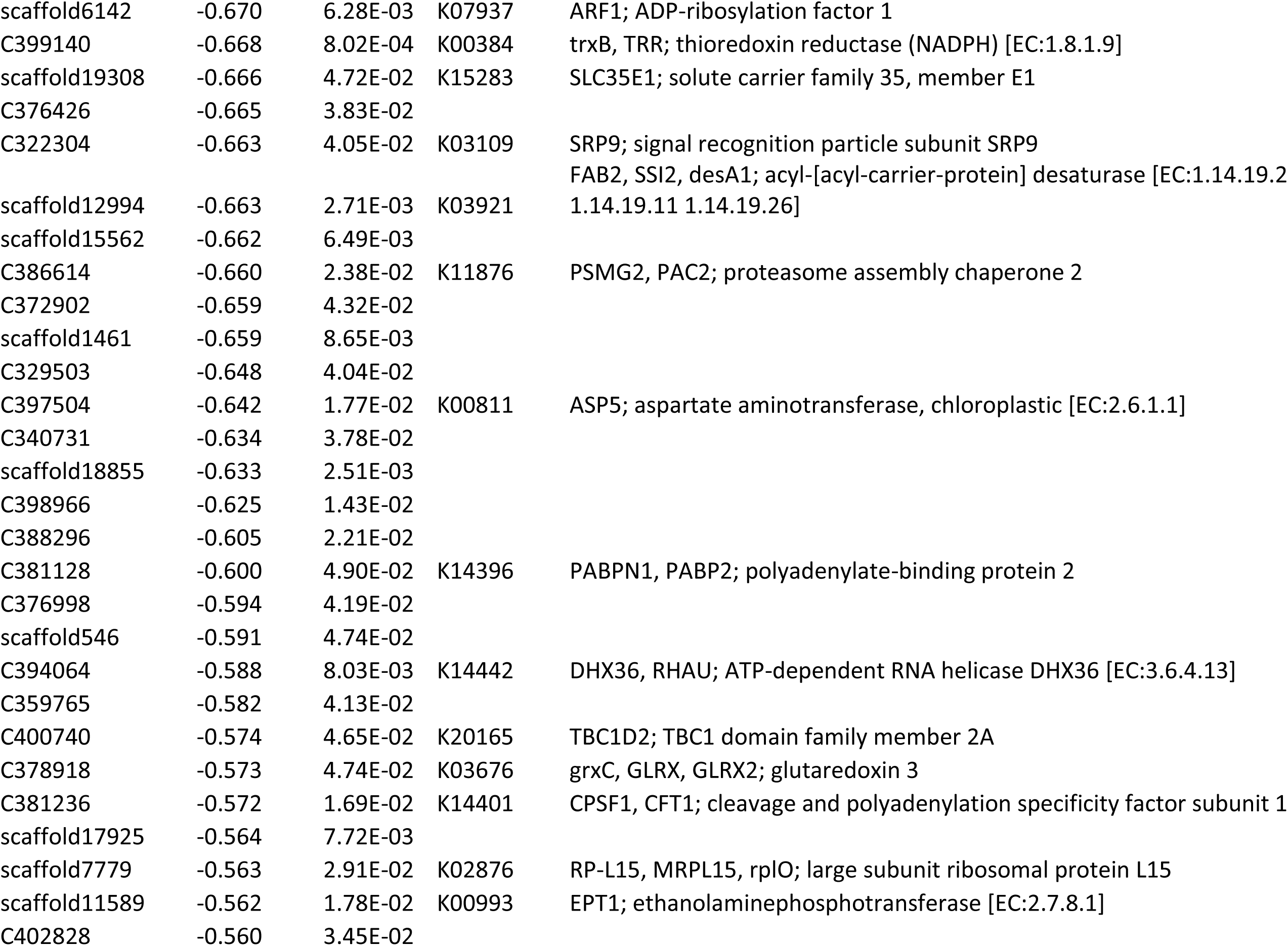
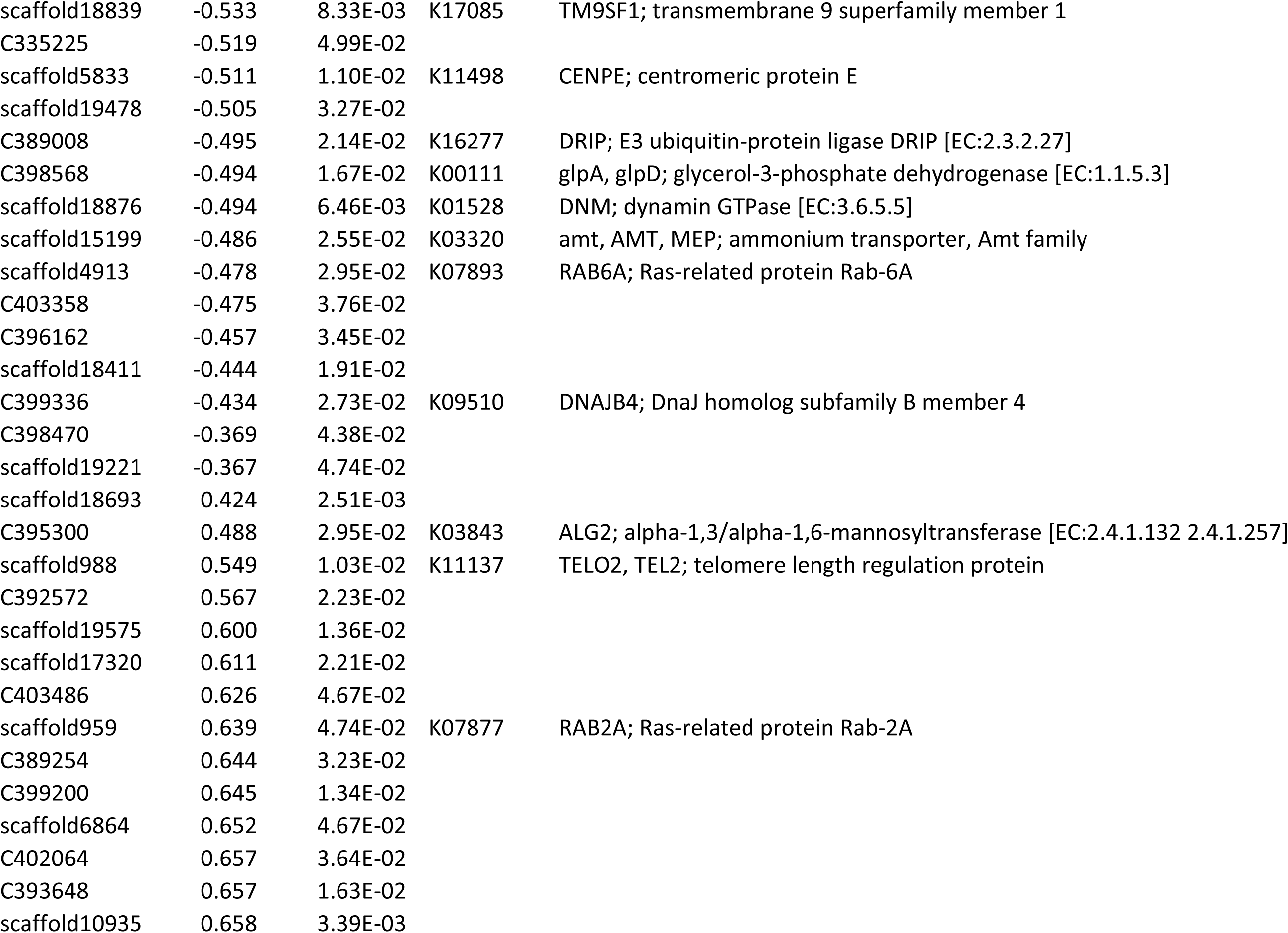
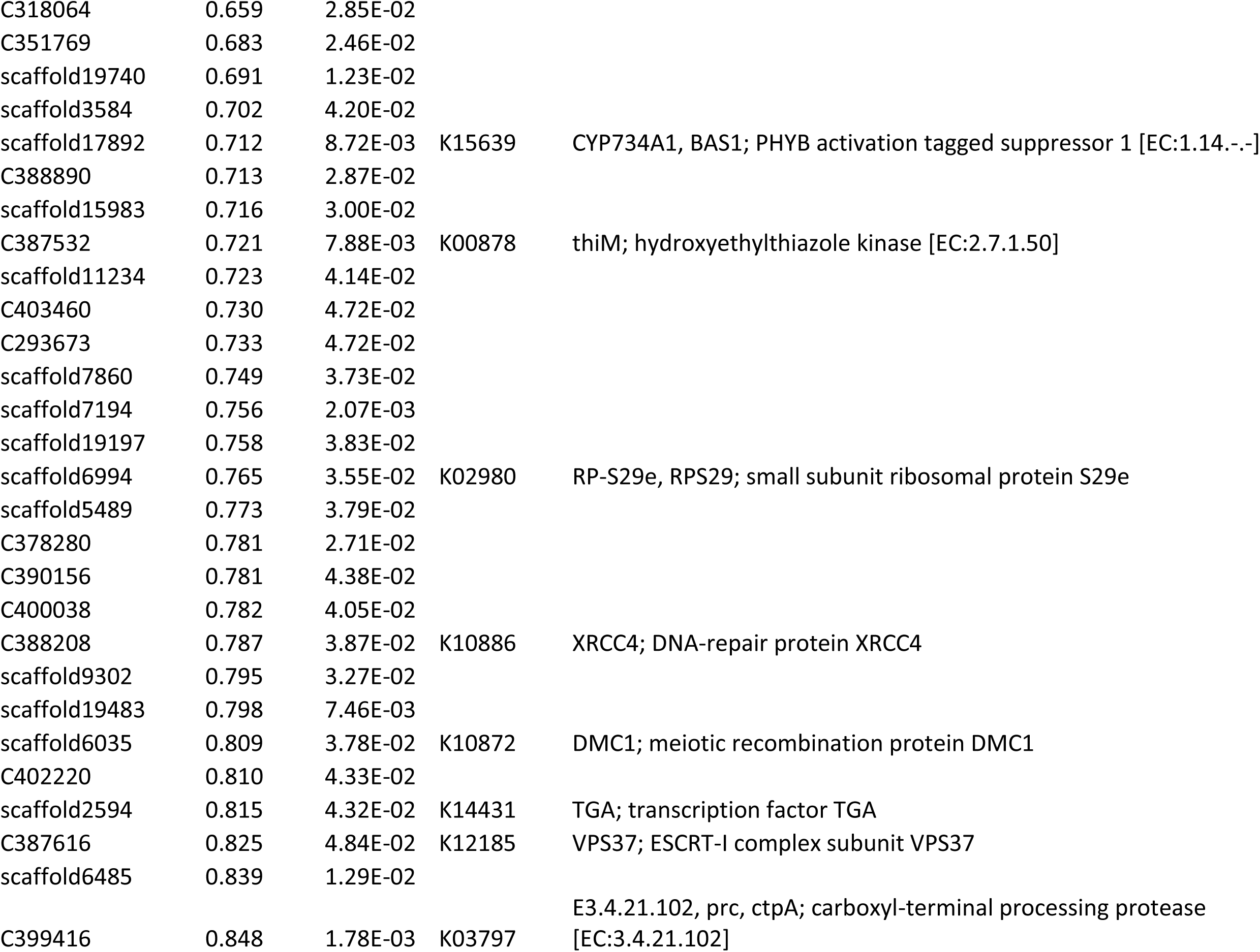
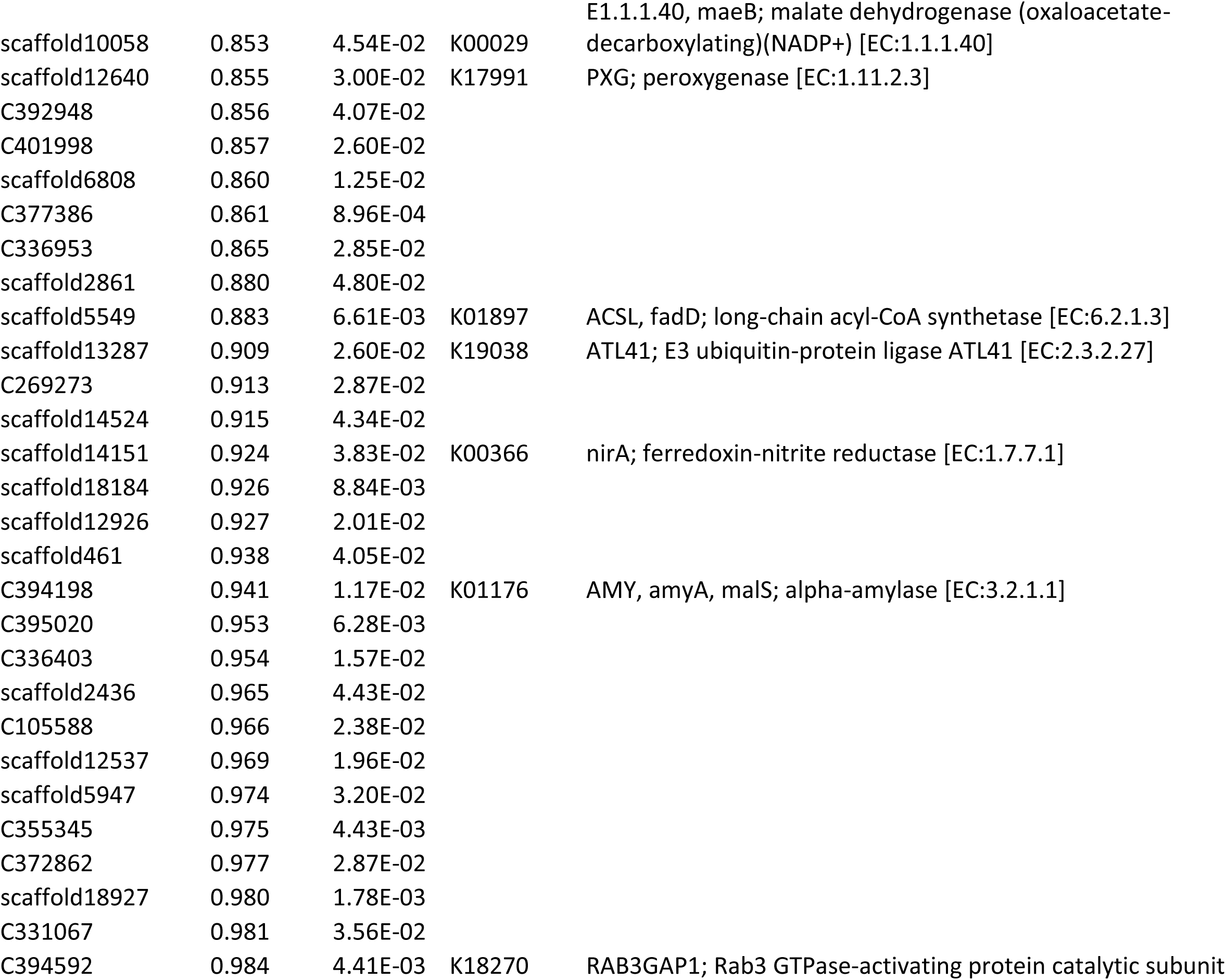
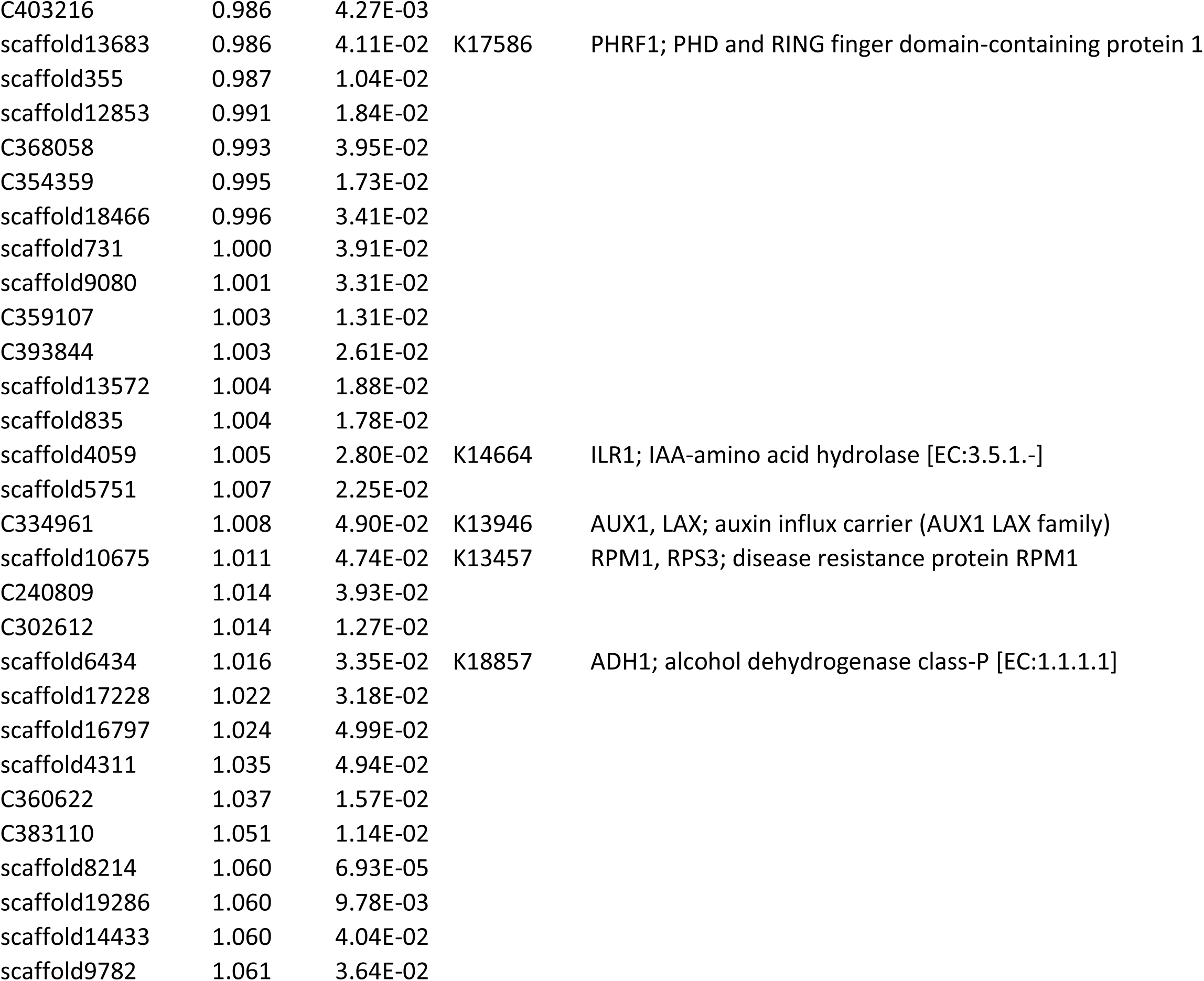
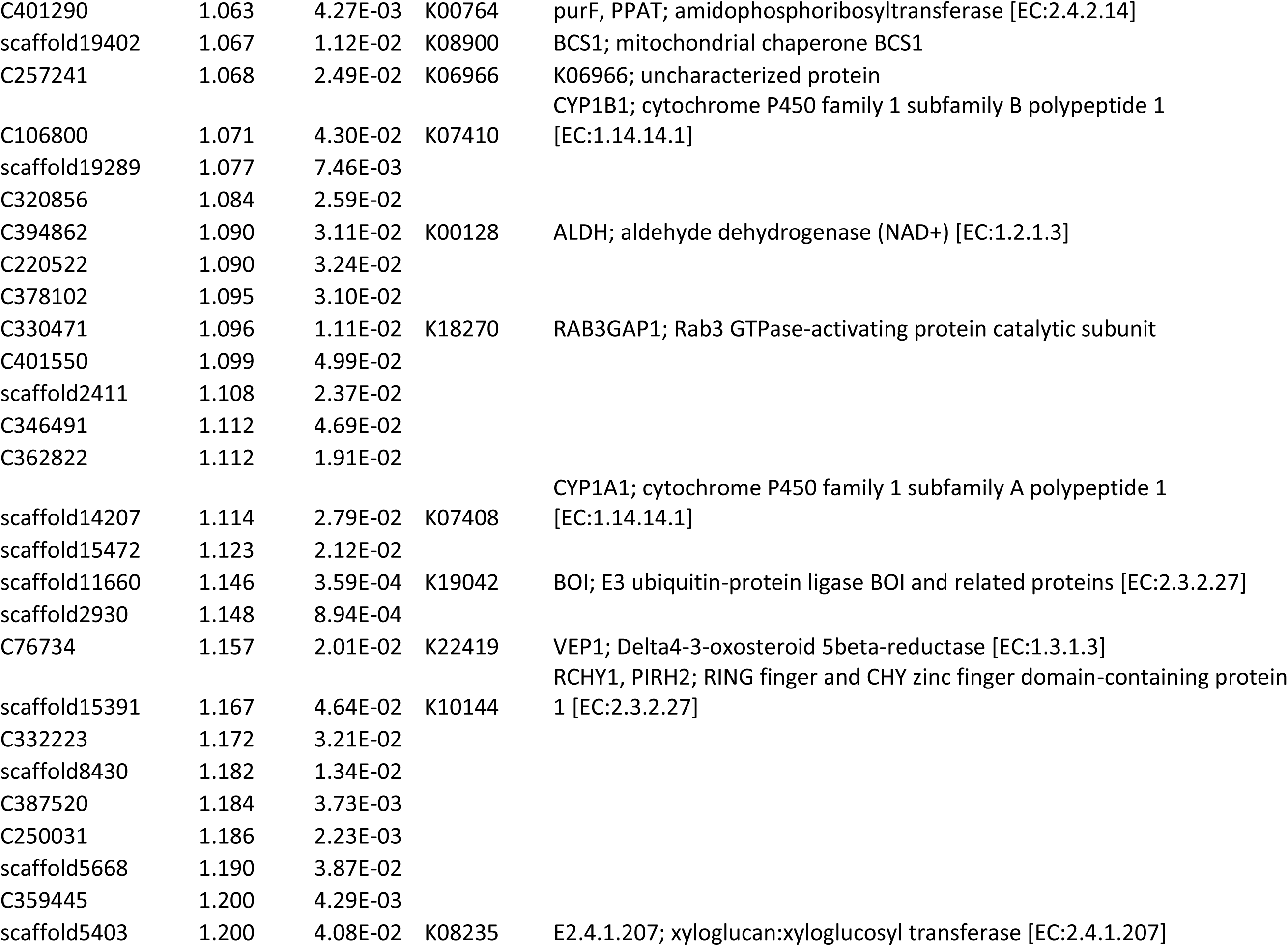
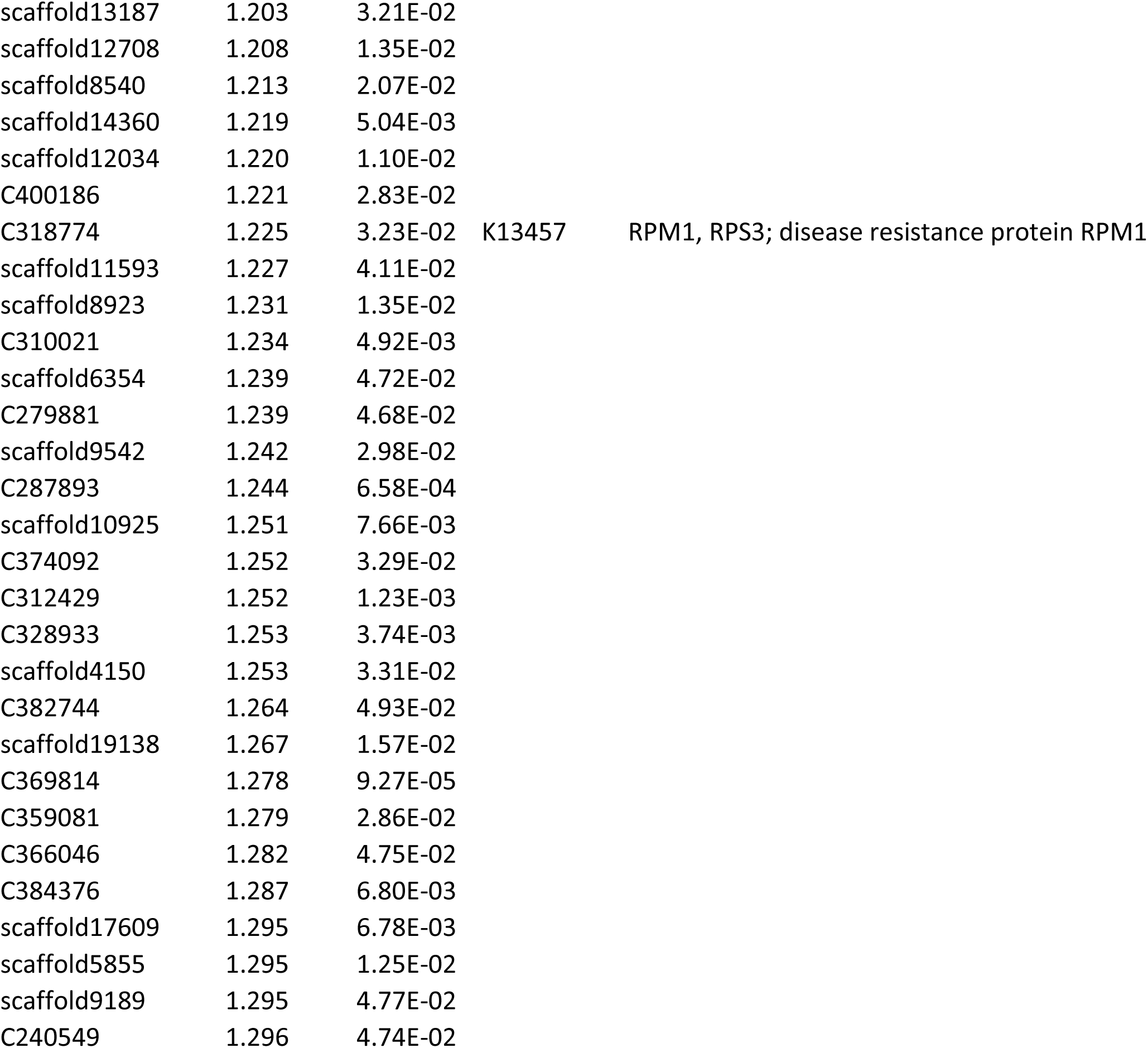
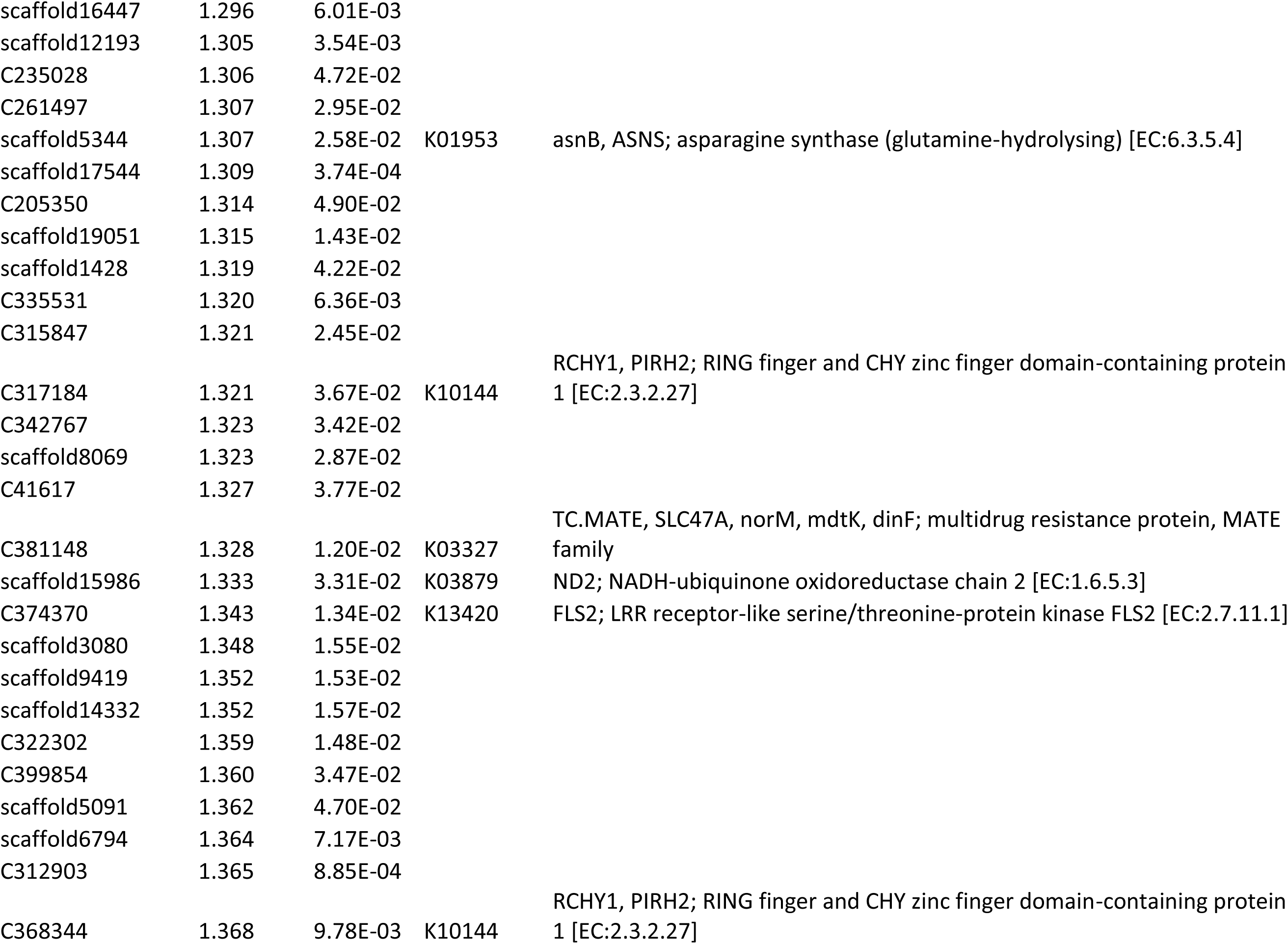
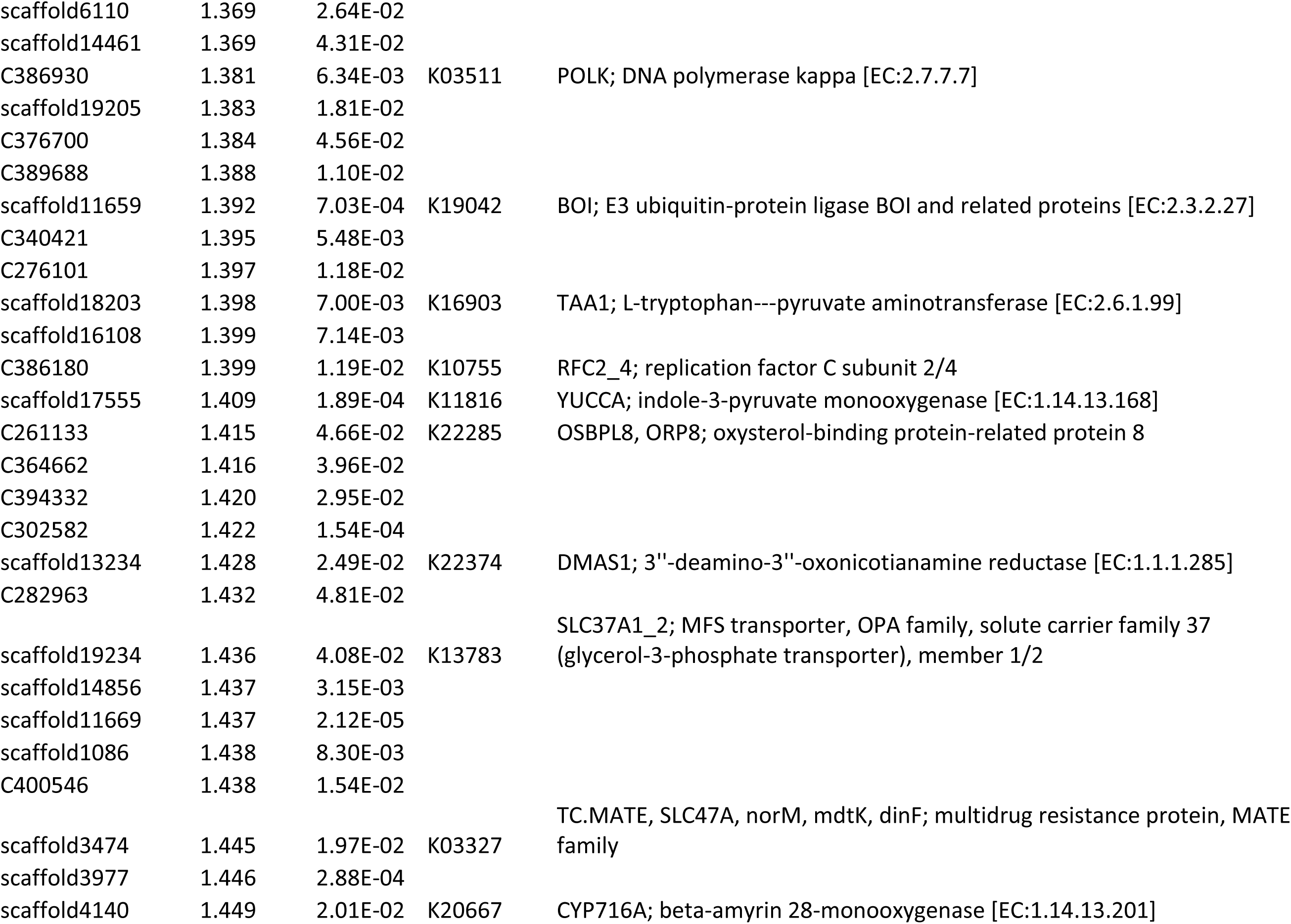
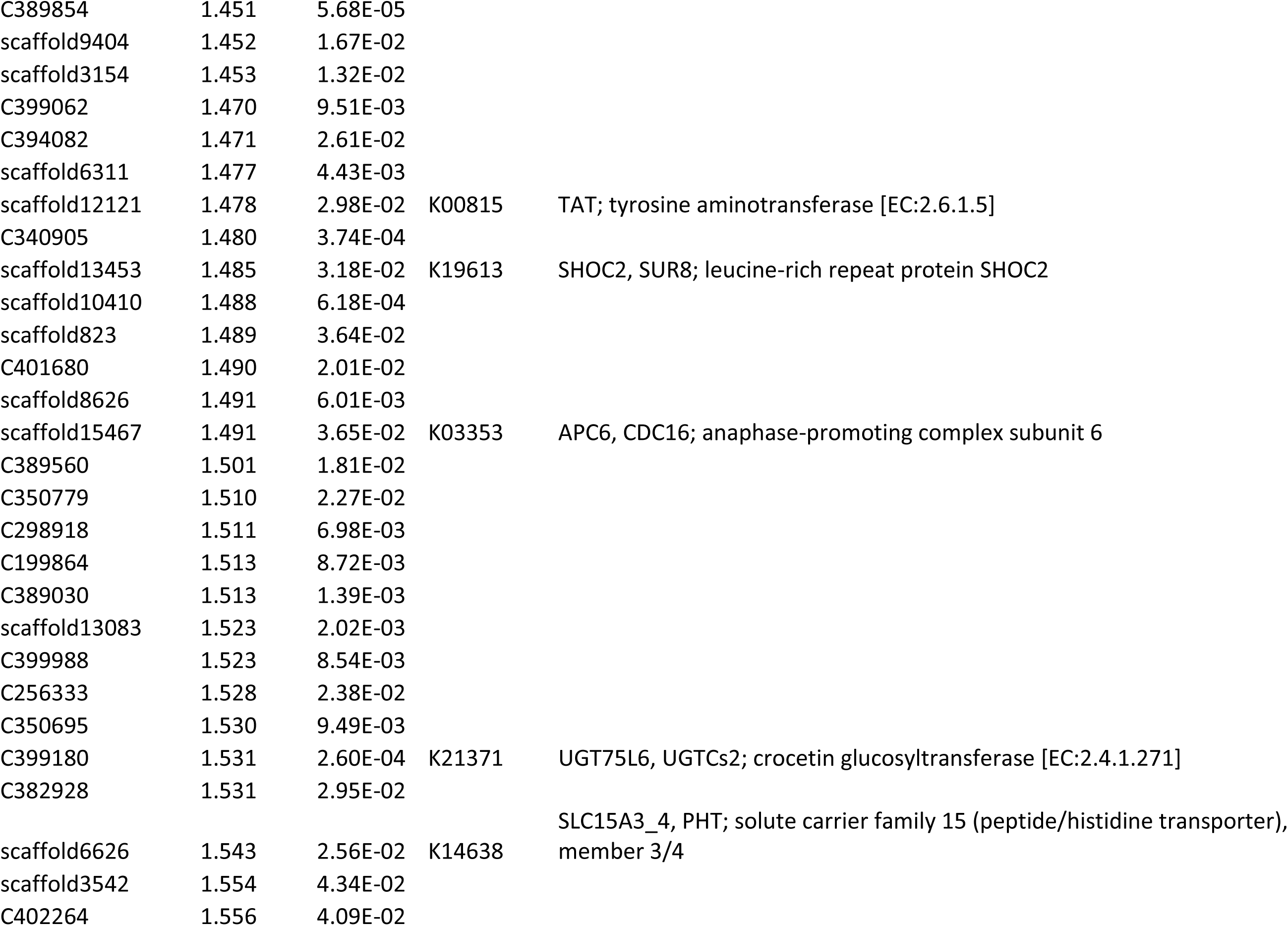
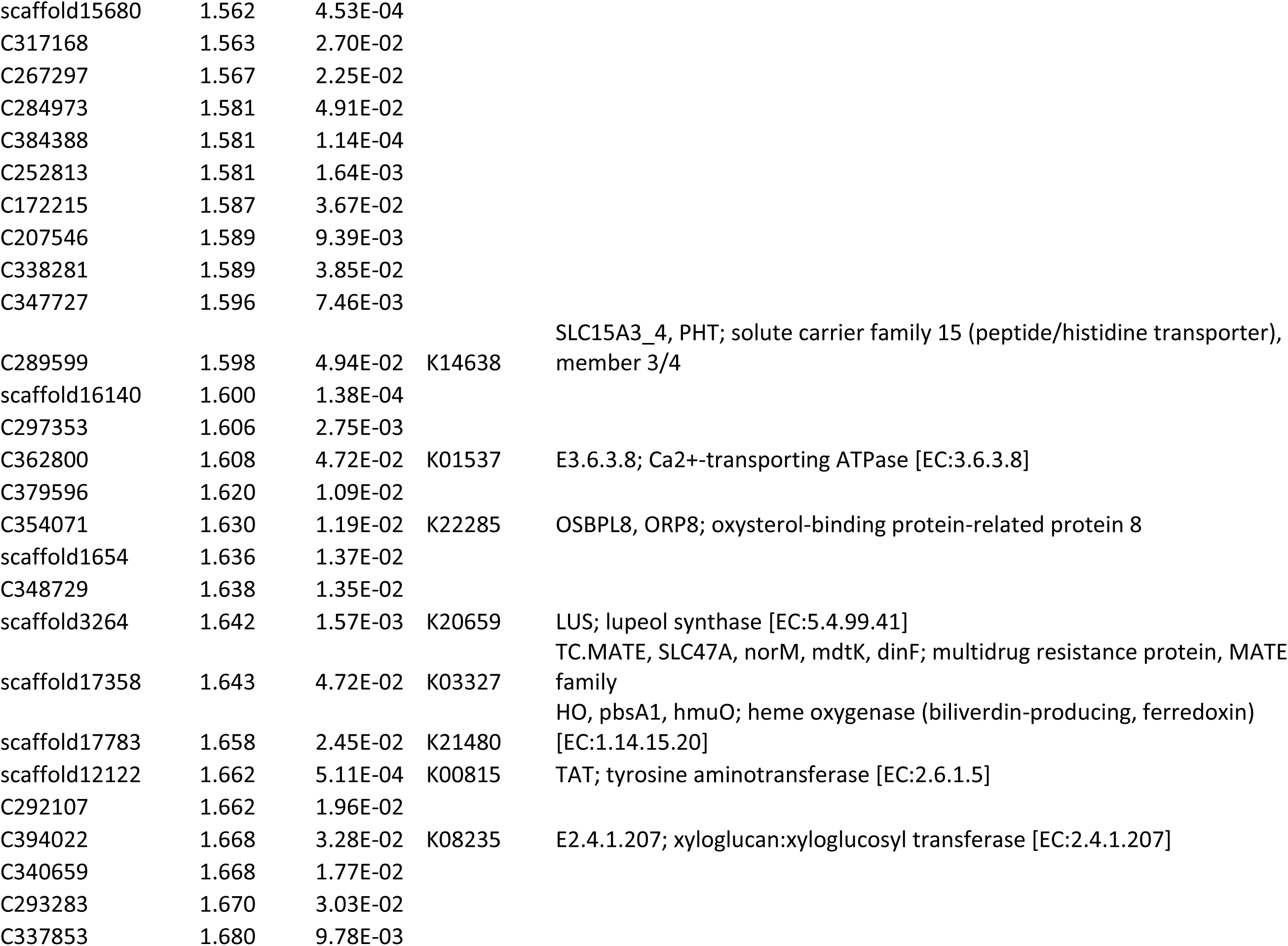
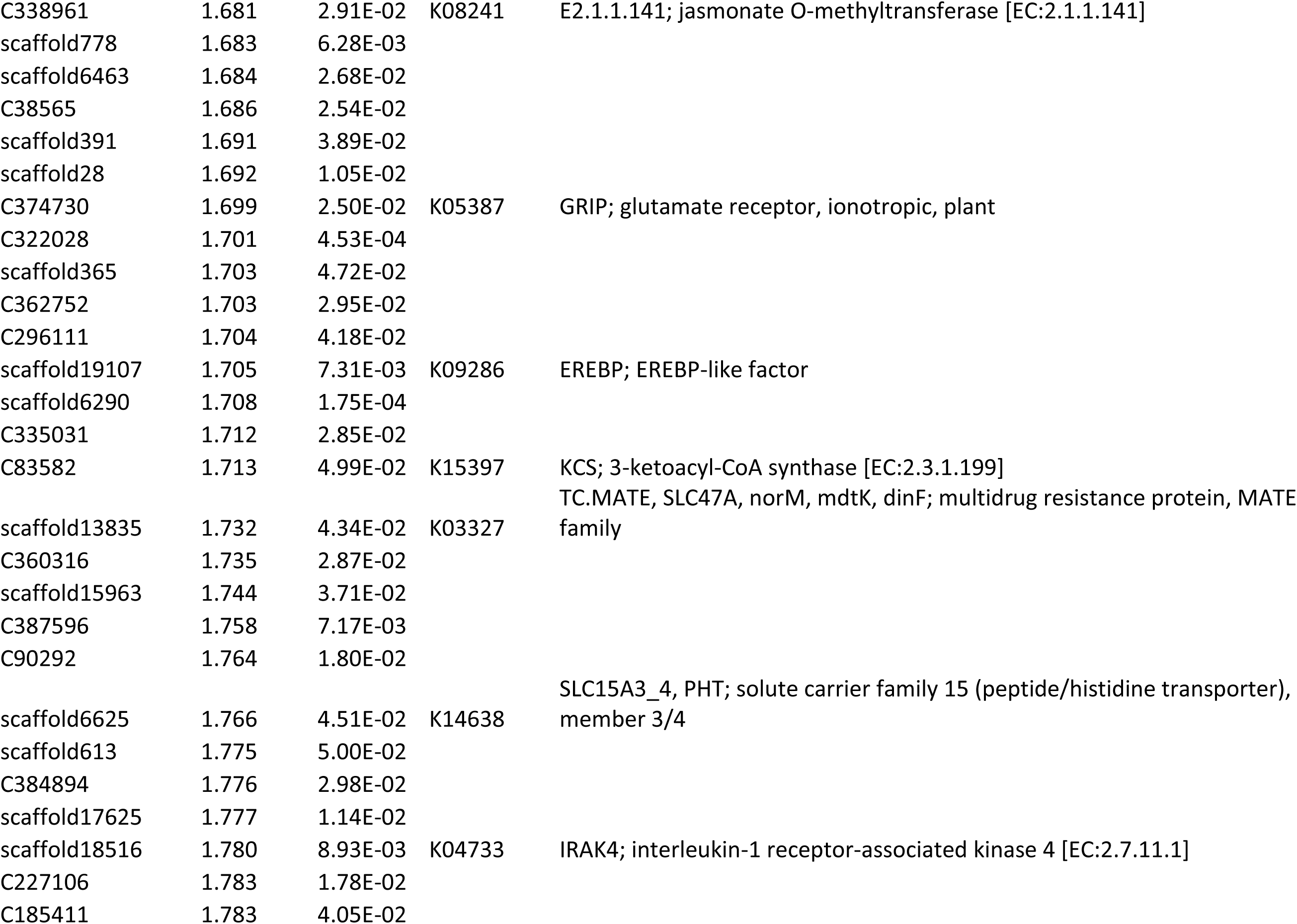
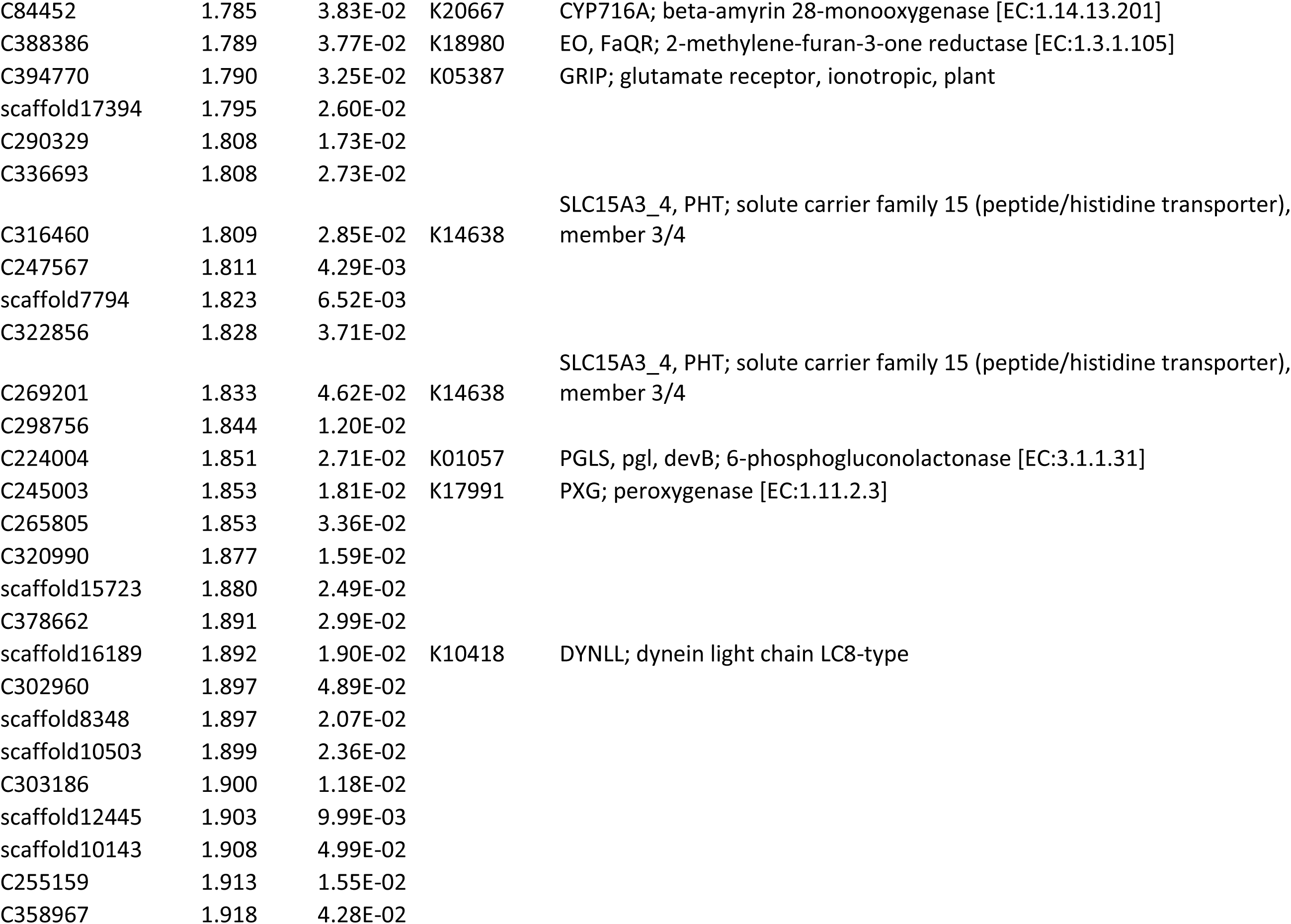
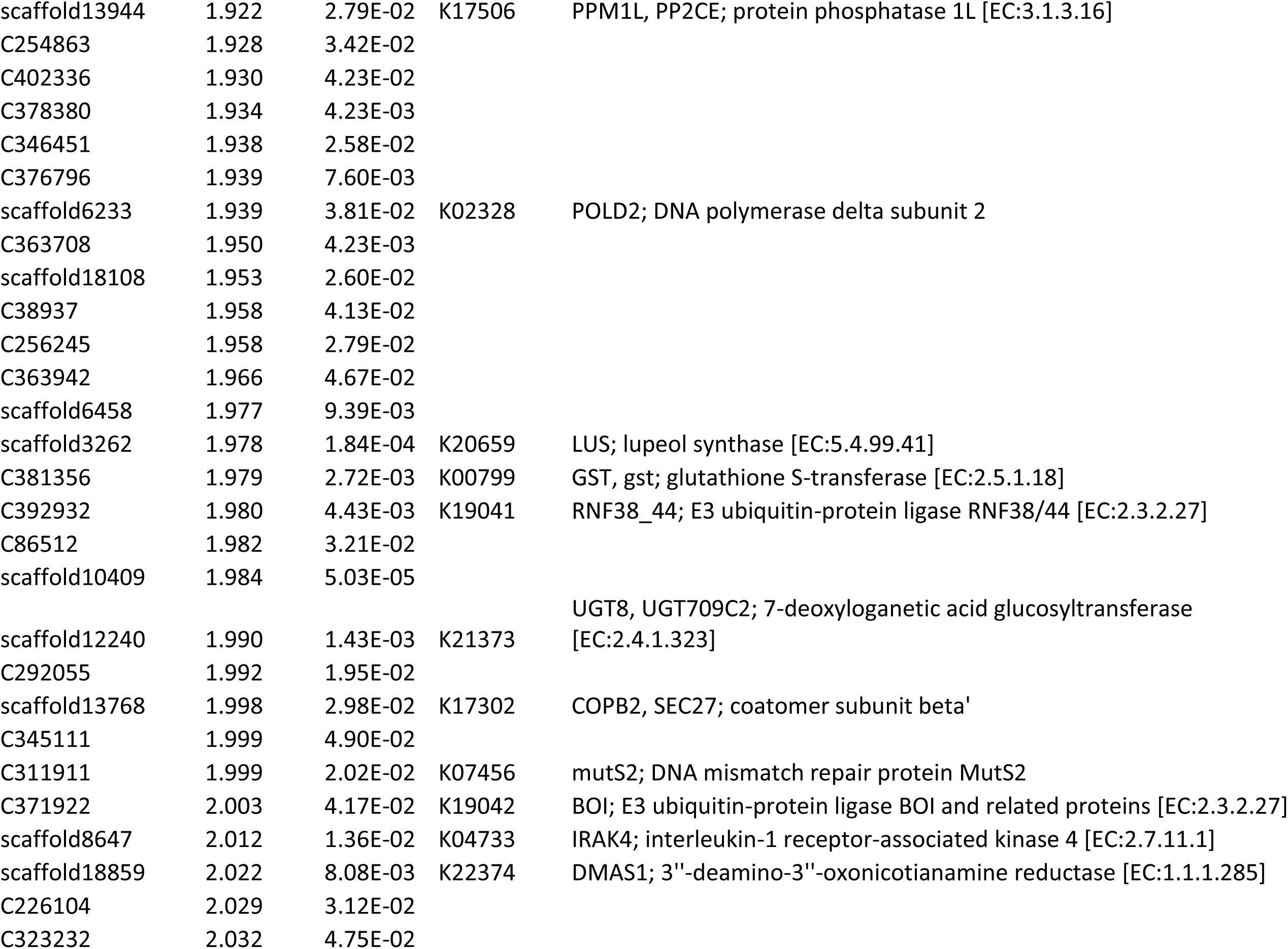
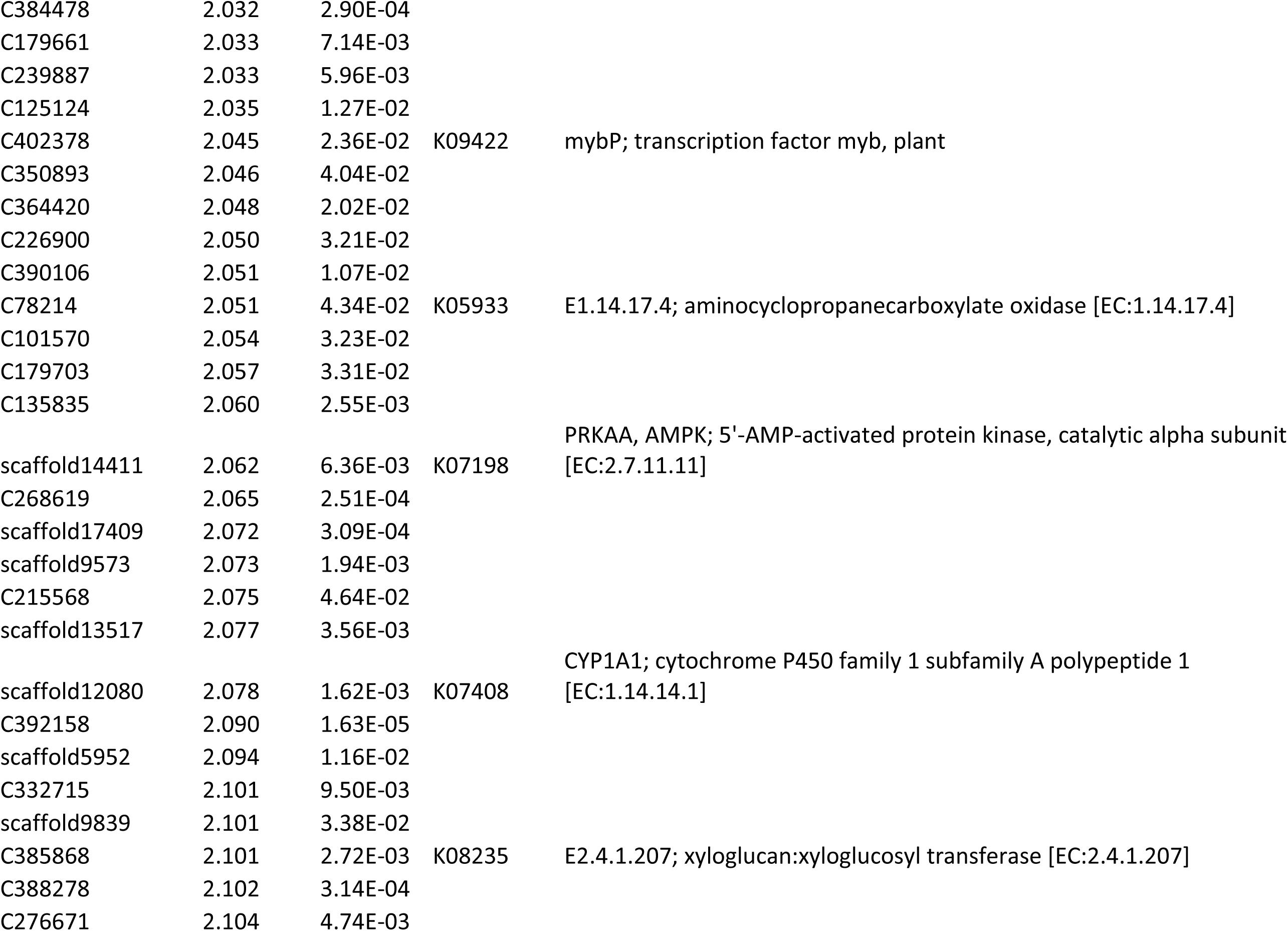
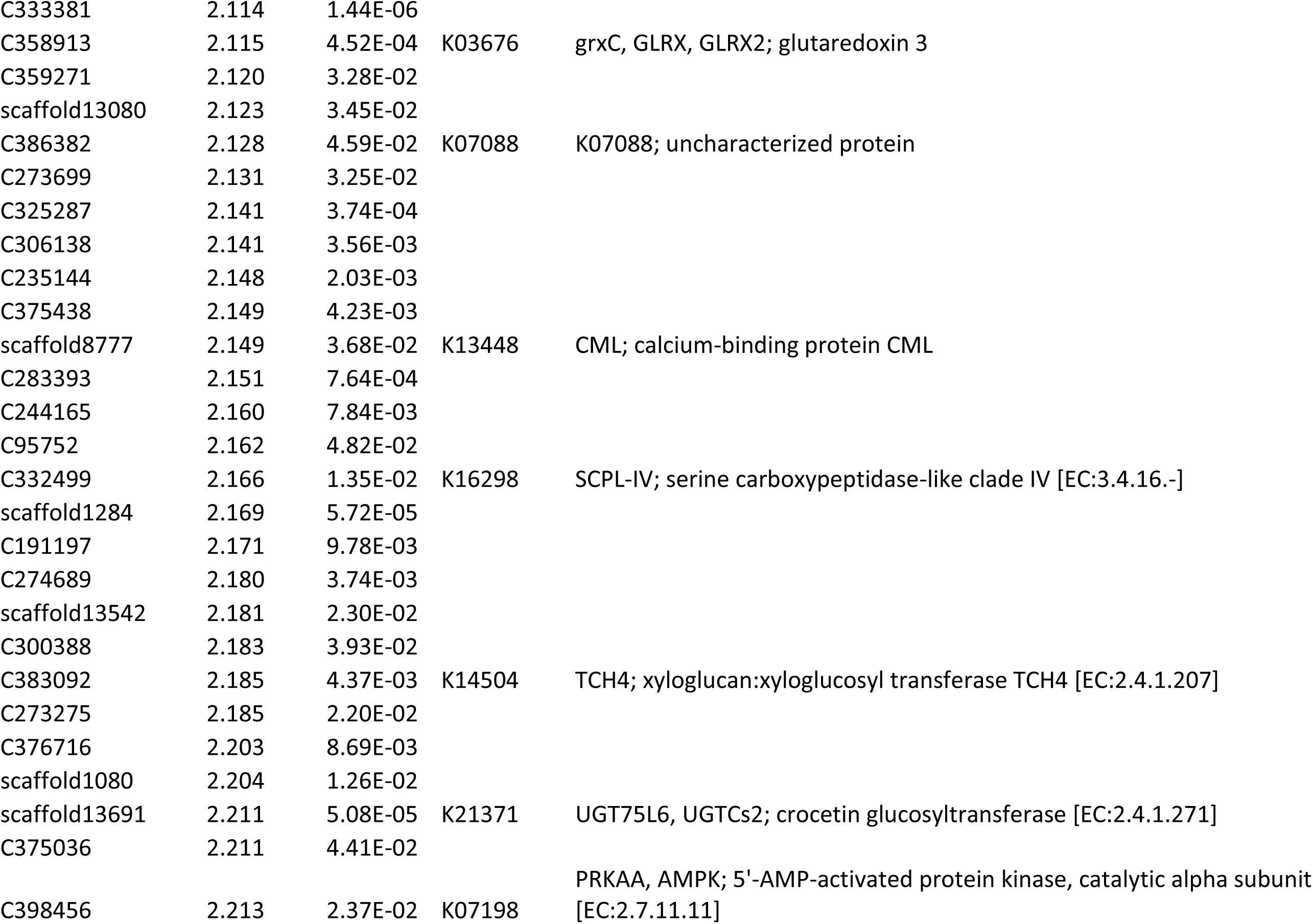
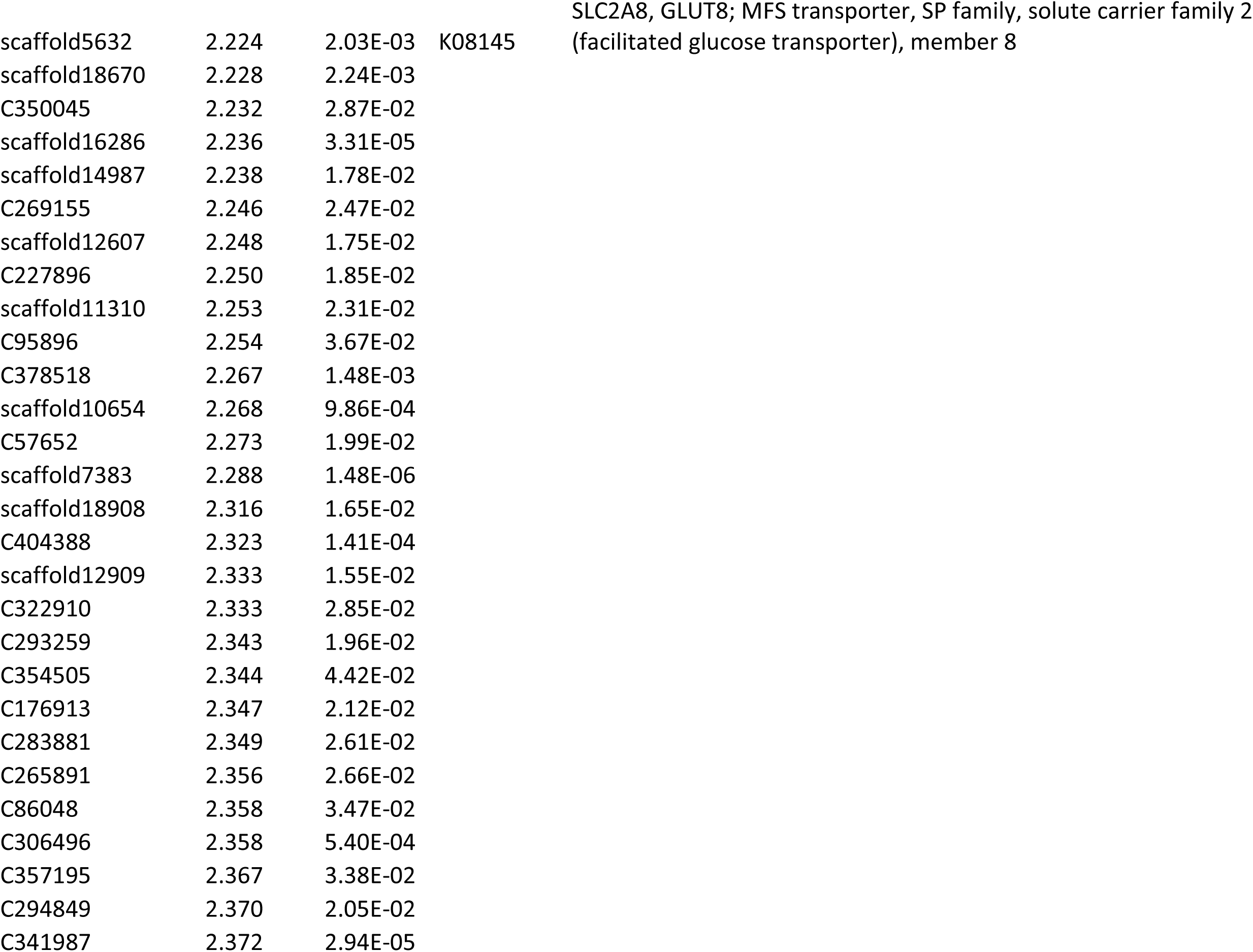
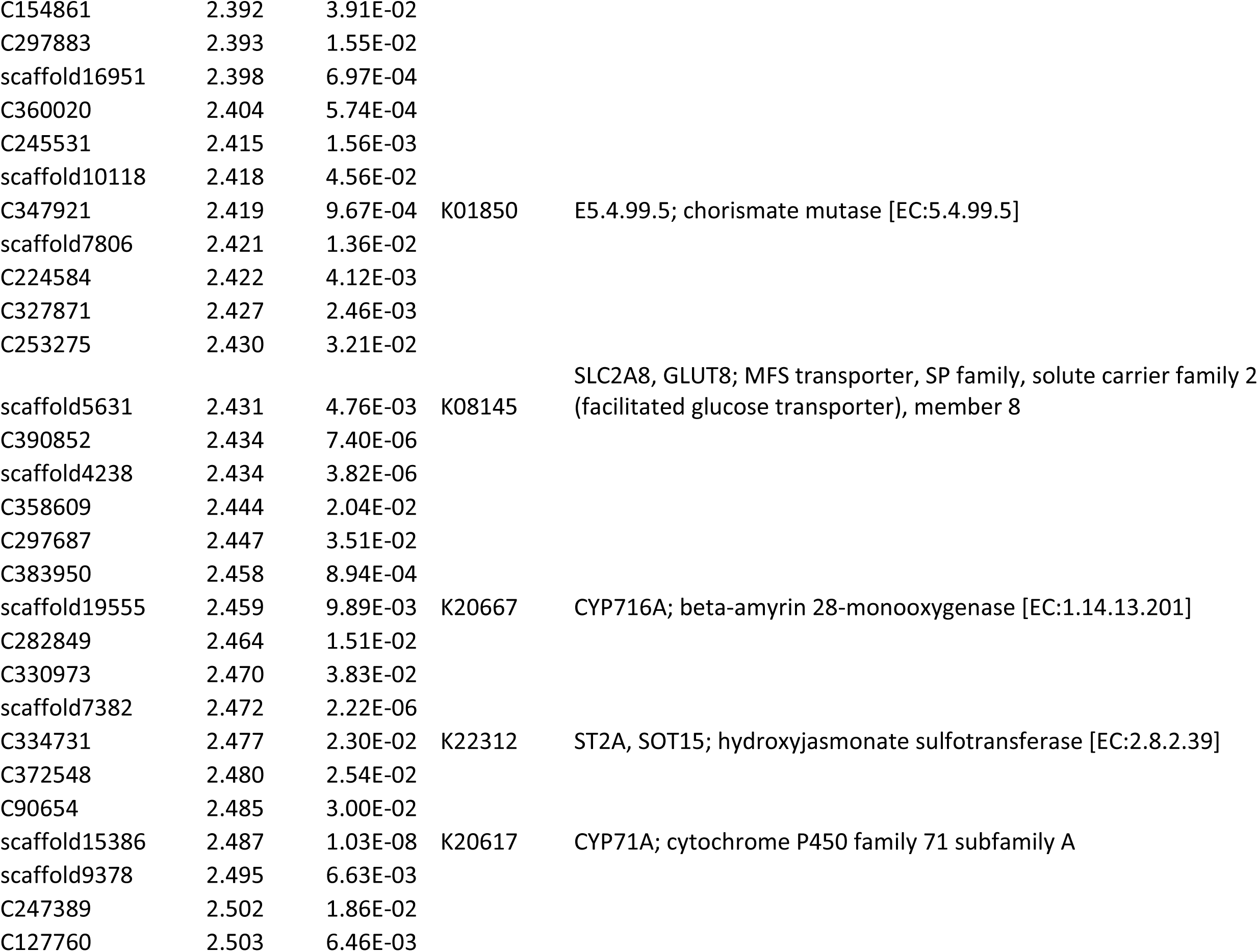
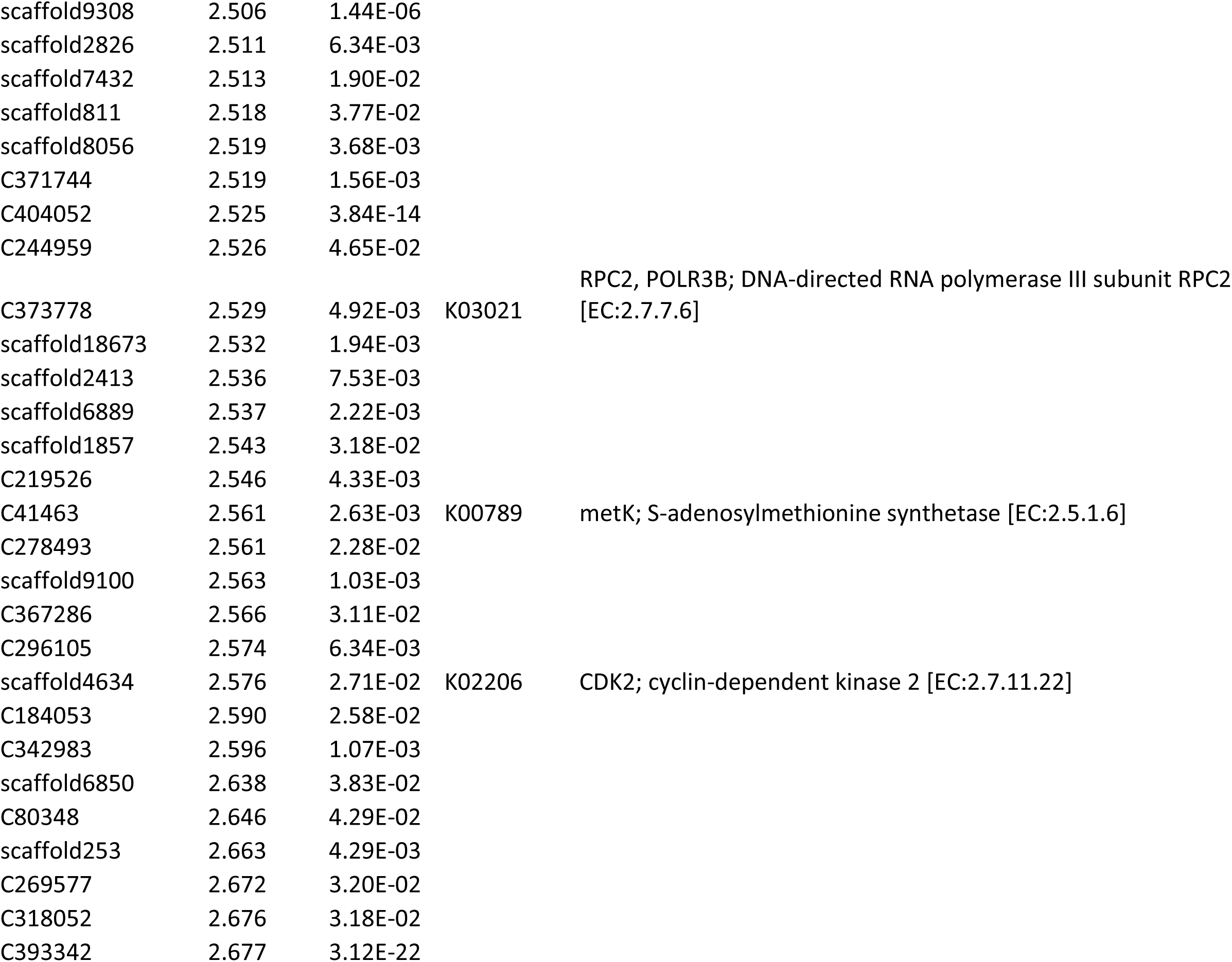
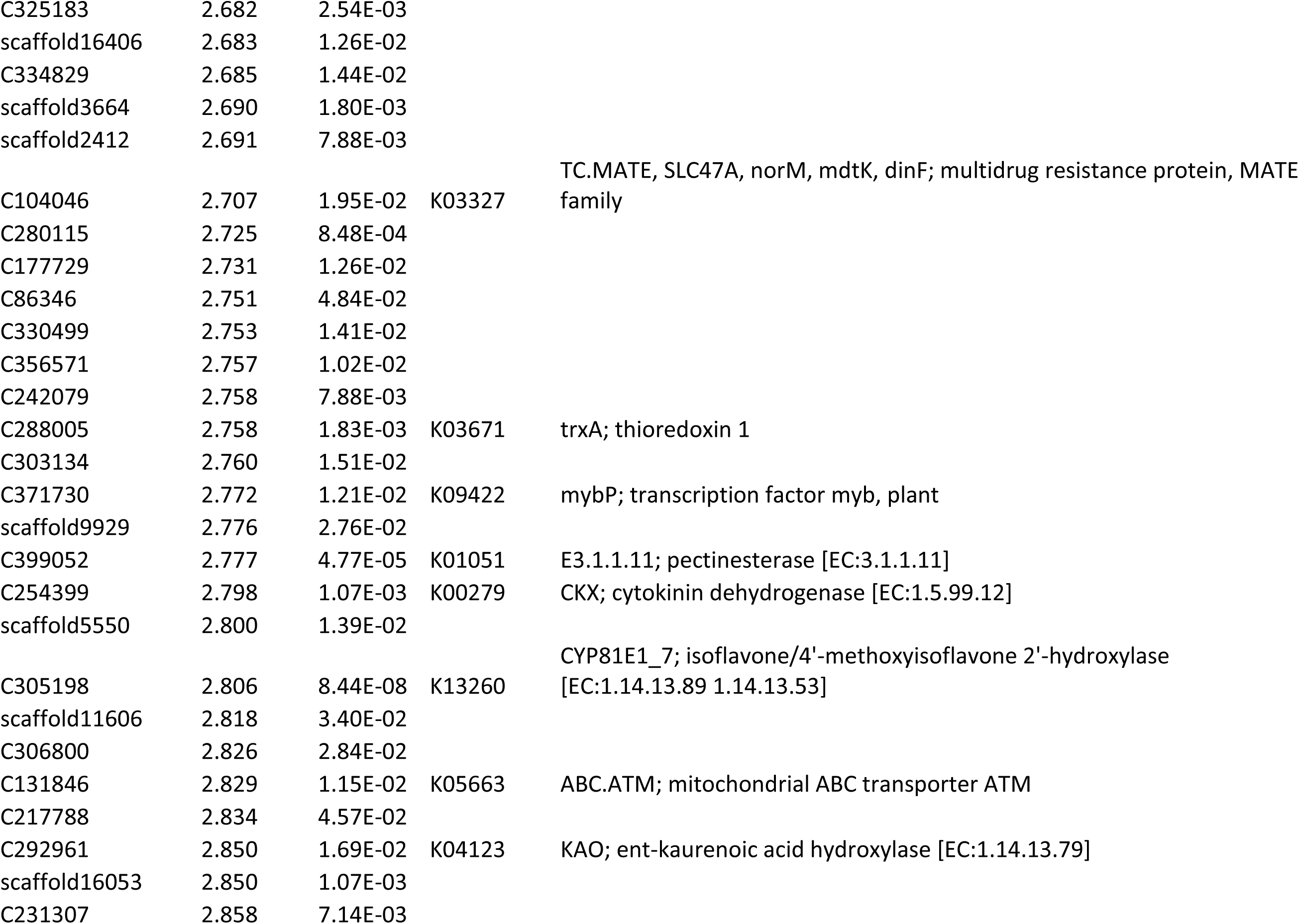
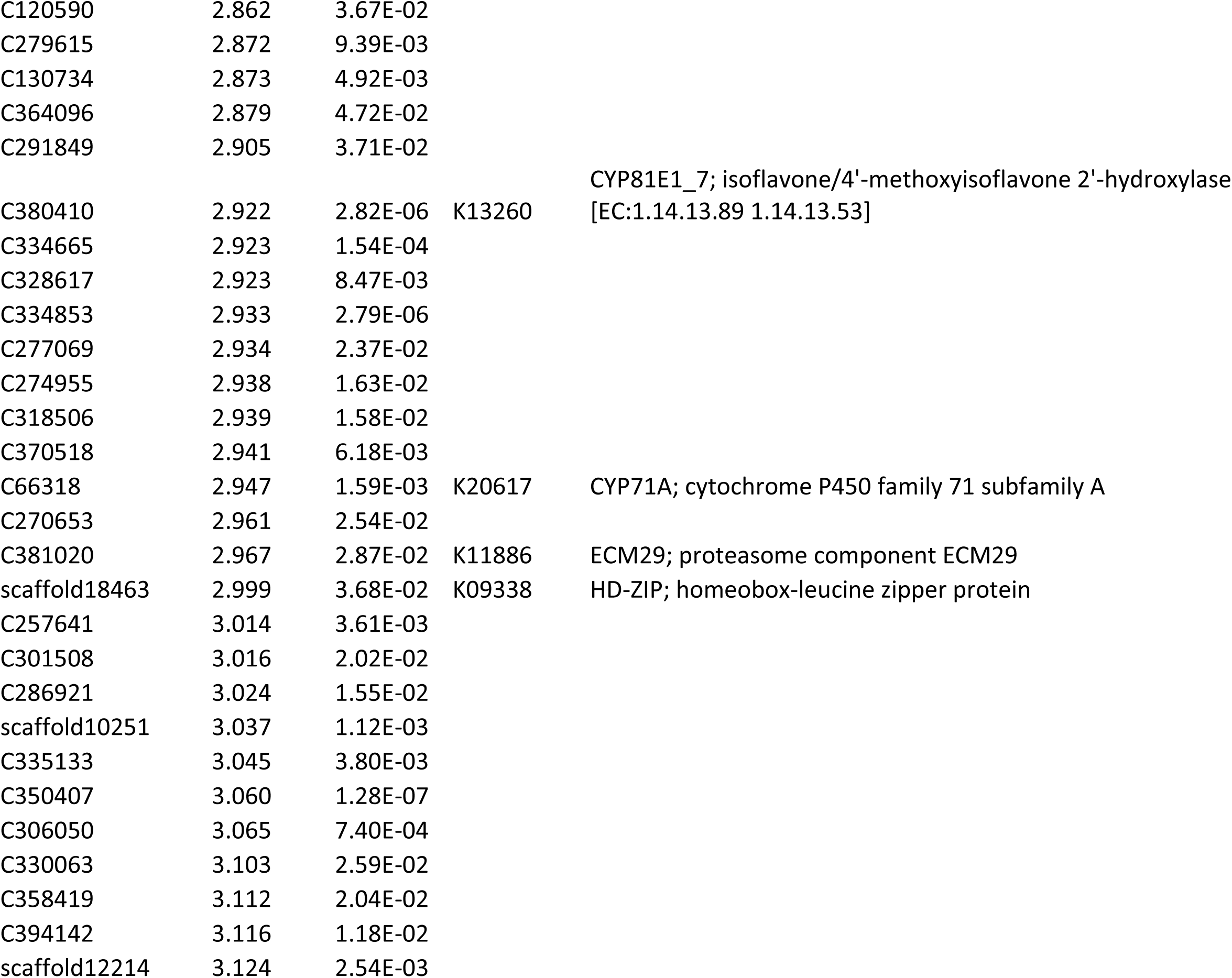
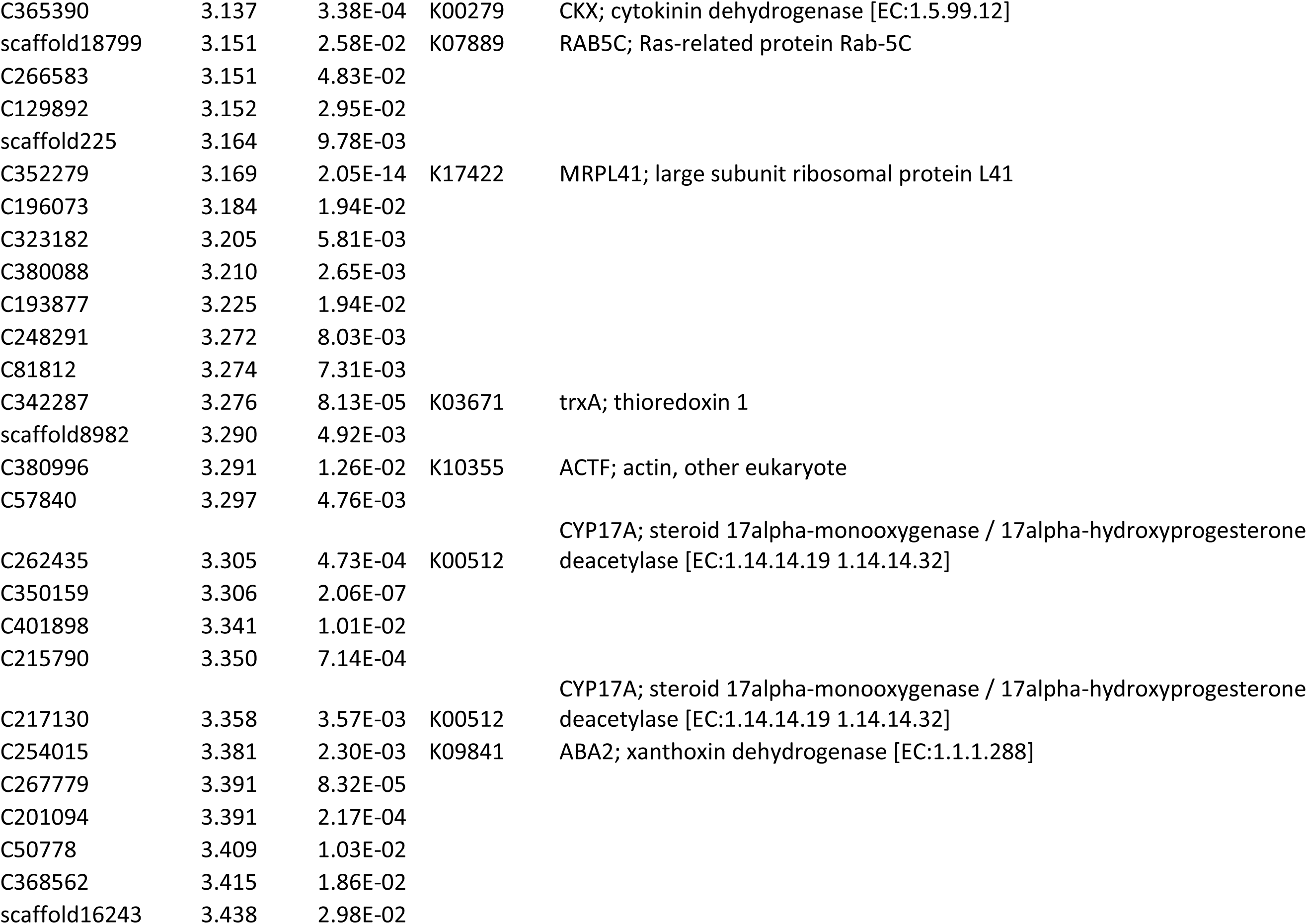
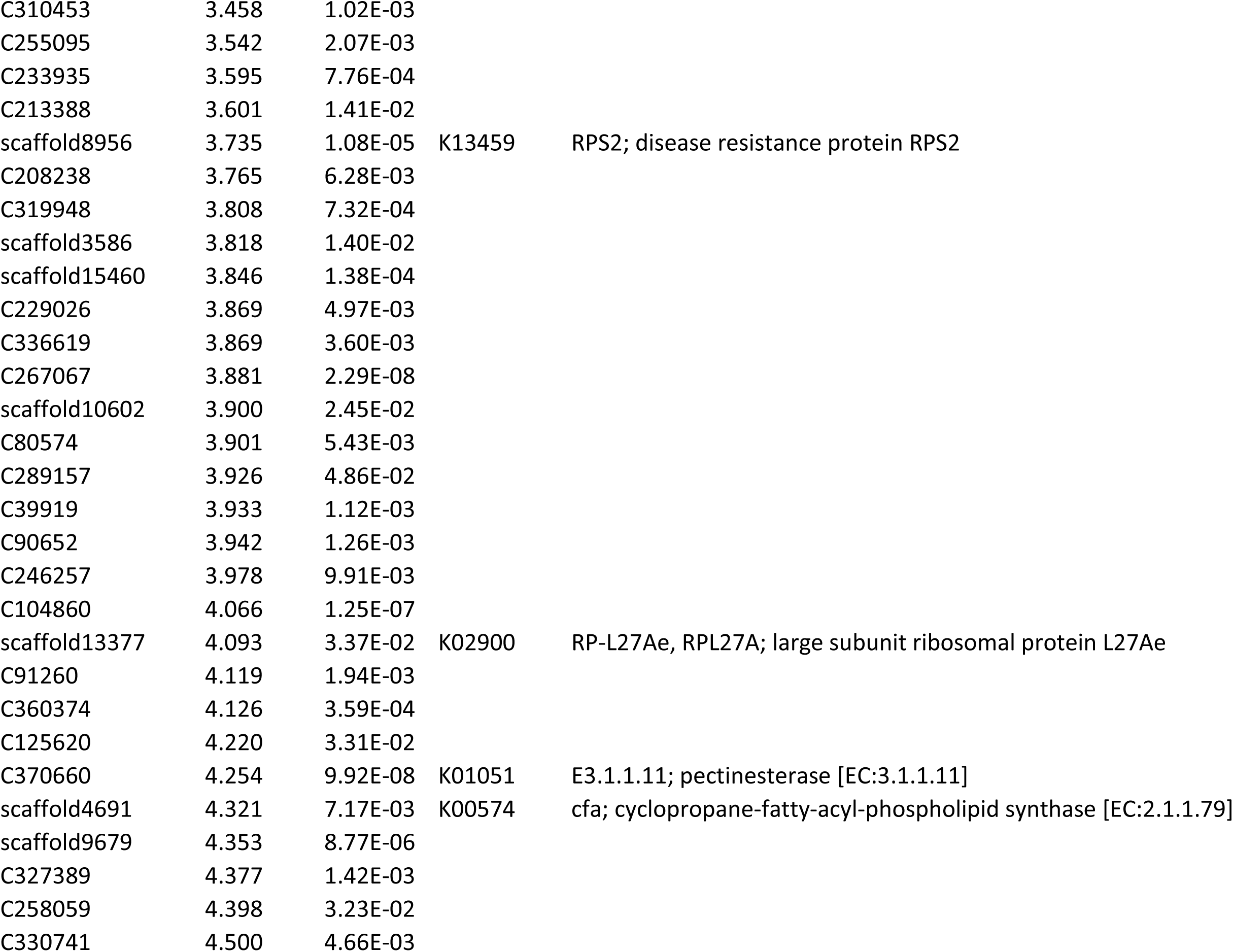
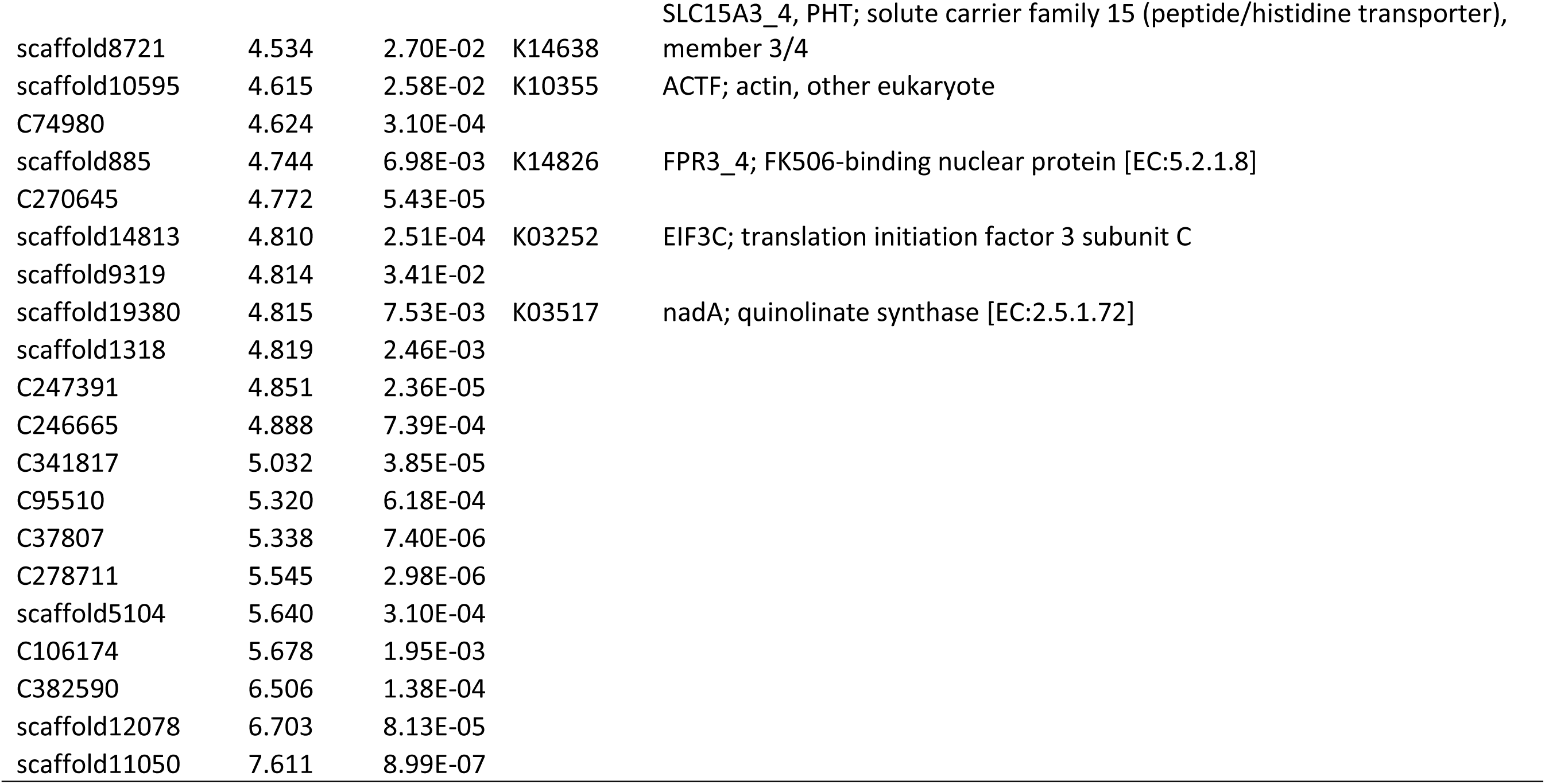
Differentially expressed genes (DEGs) identified between white- and red-bracted cultivars of C. florida. Includes transcript ID, log2 fold change, adjusted p-value, KEGG ortholog, and functional annotation.

**Supplementary Table T6.**
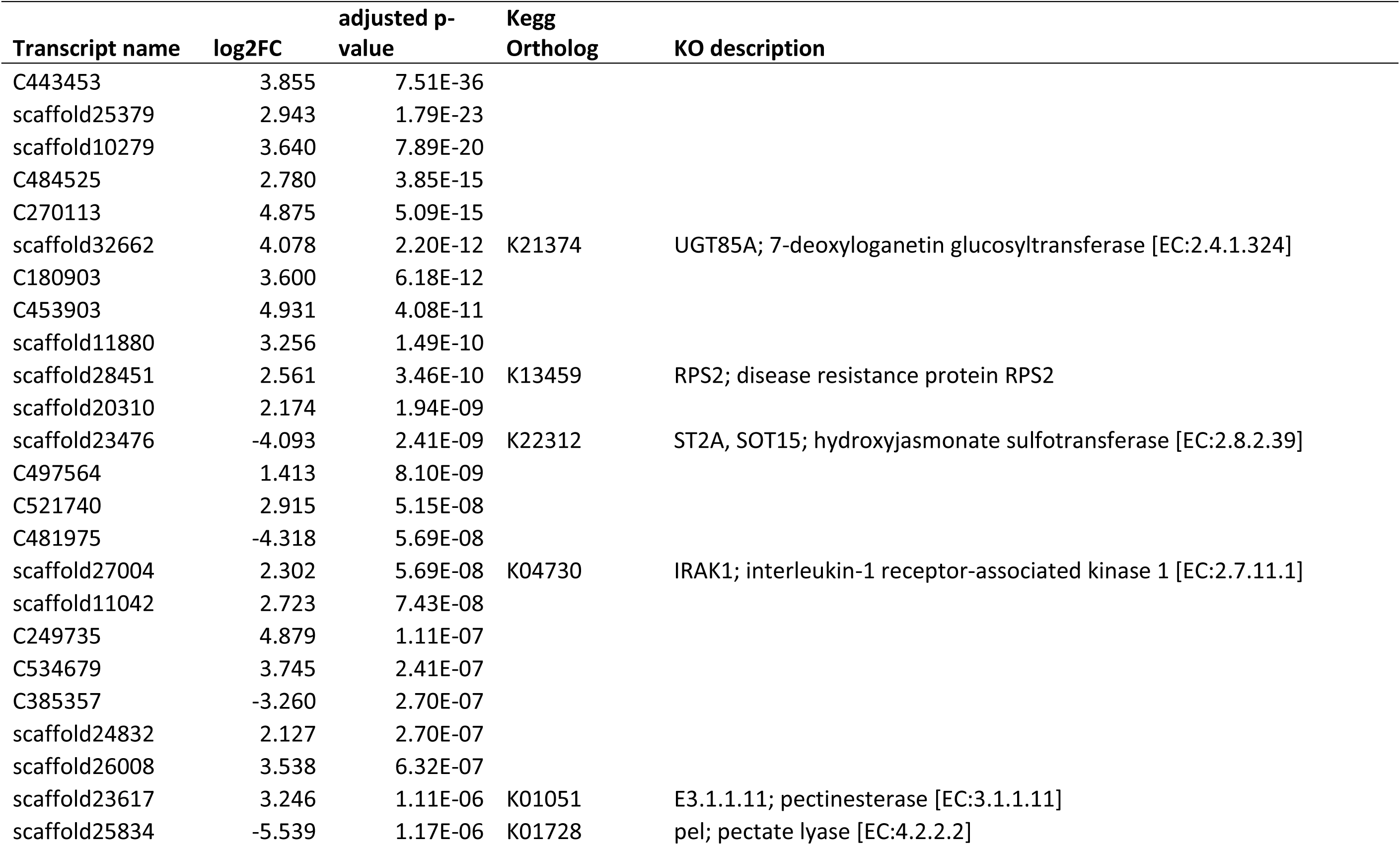
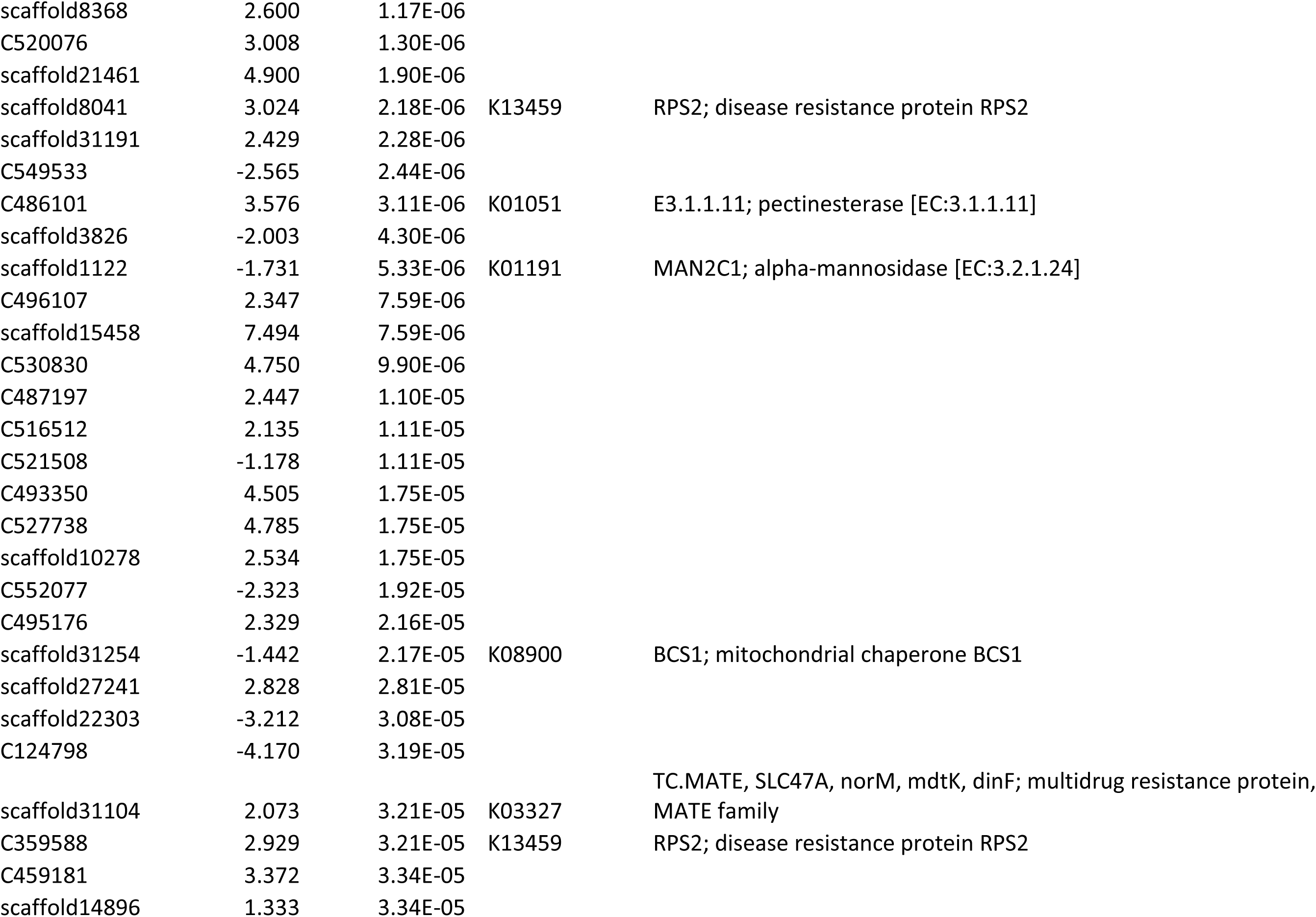
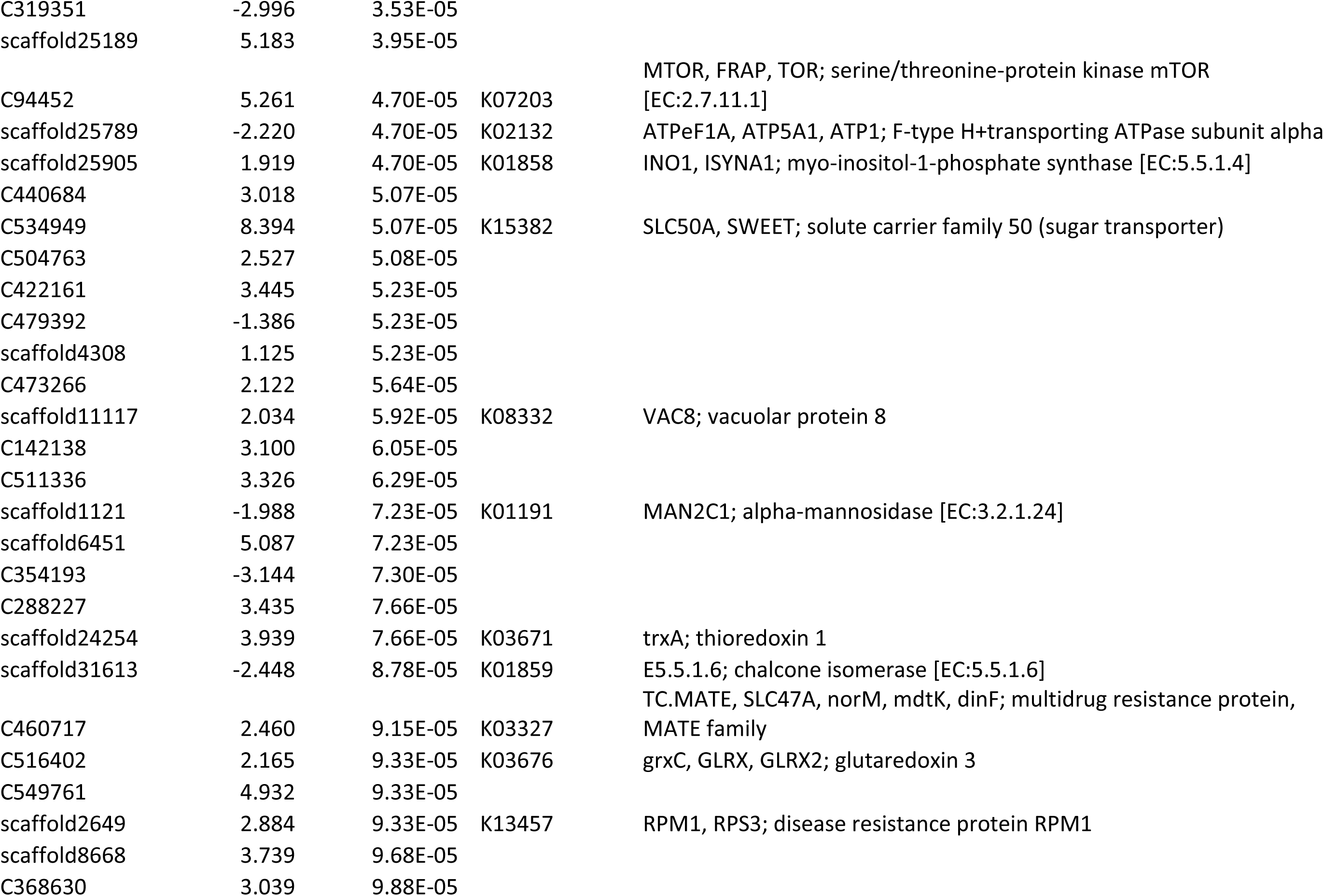
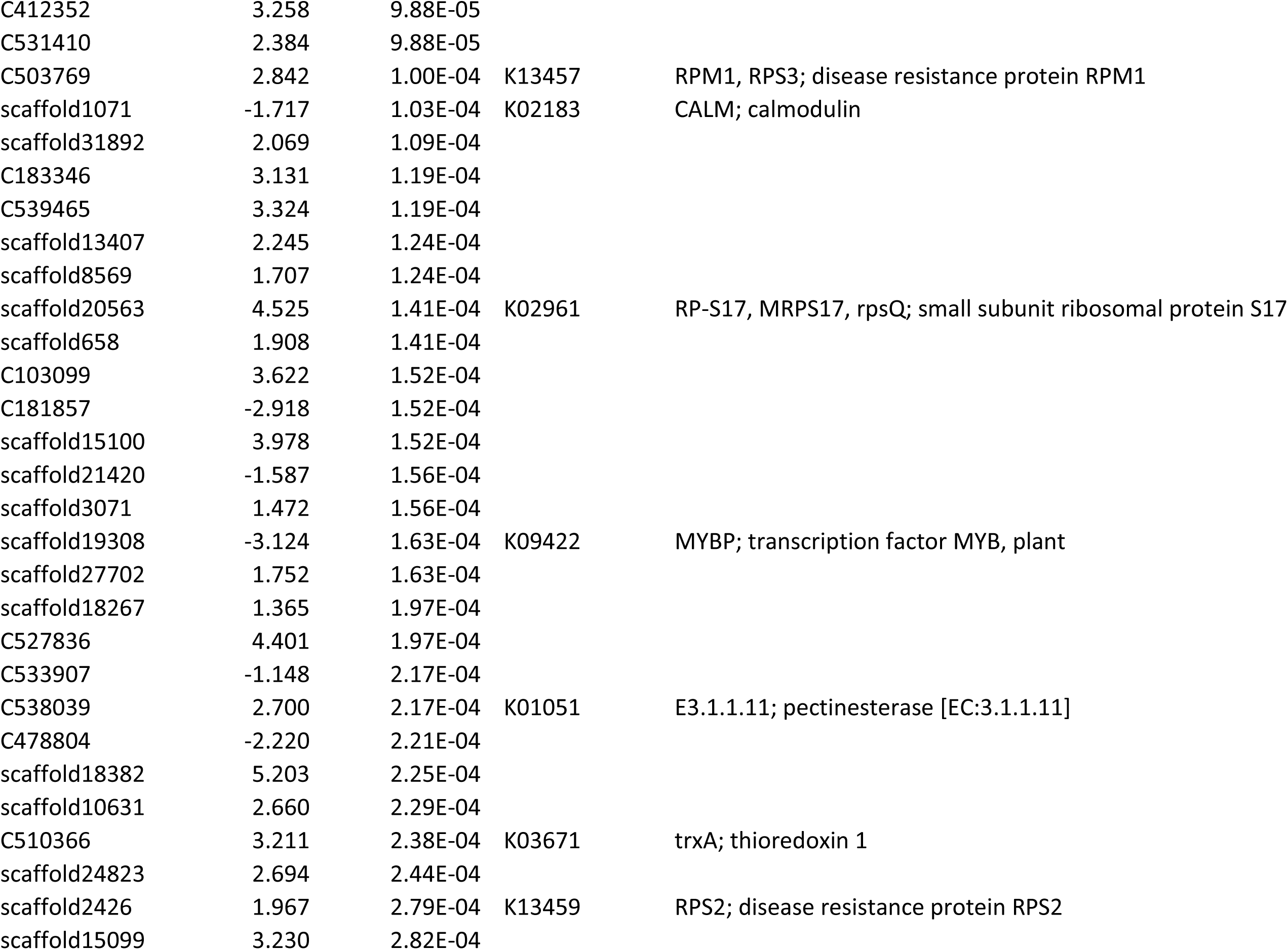
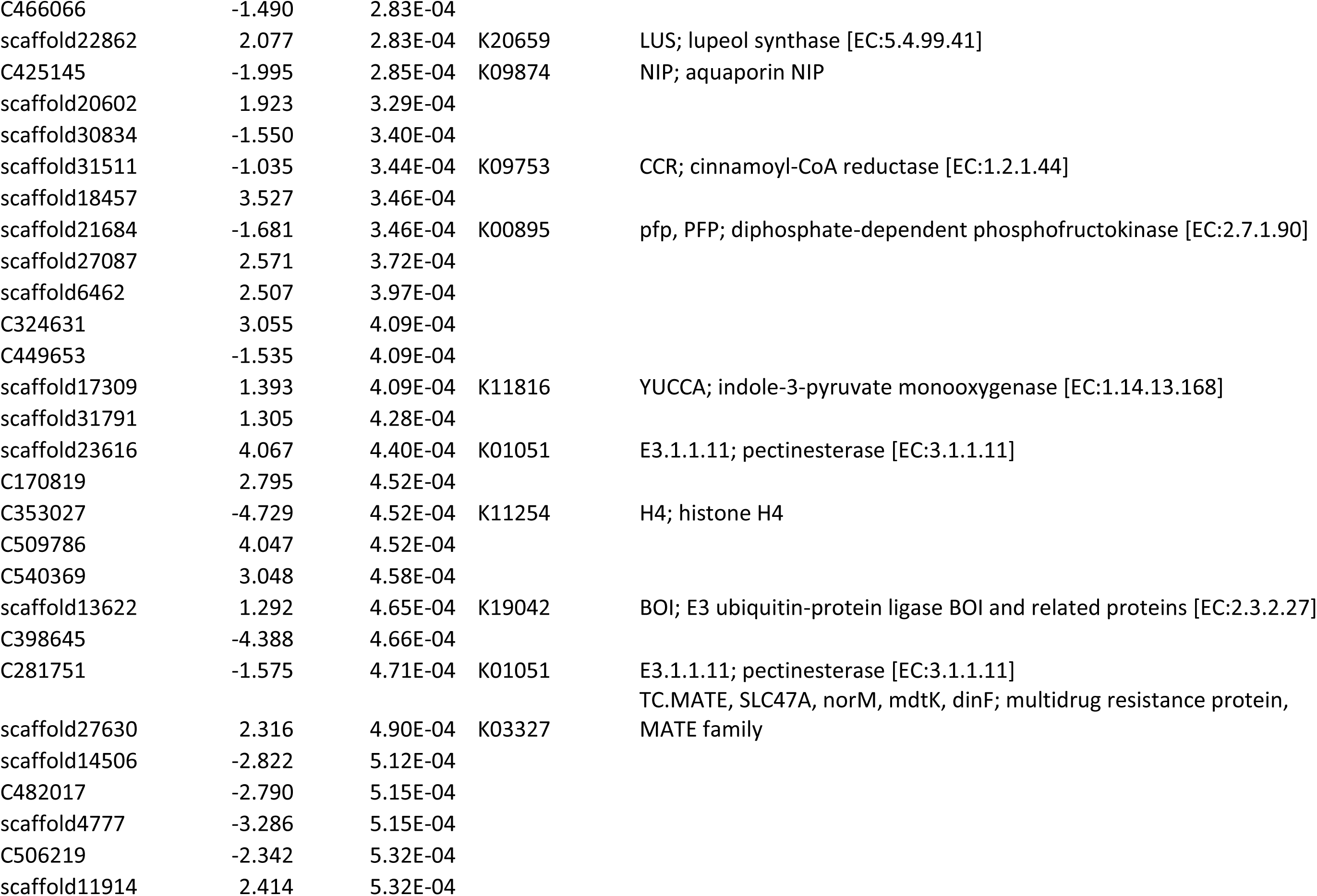
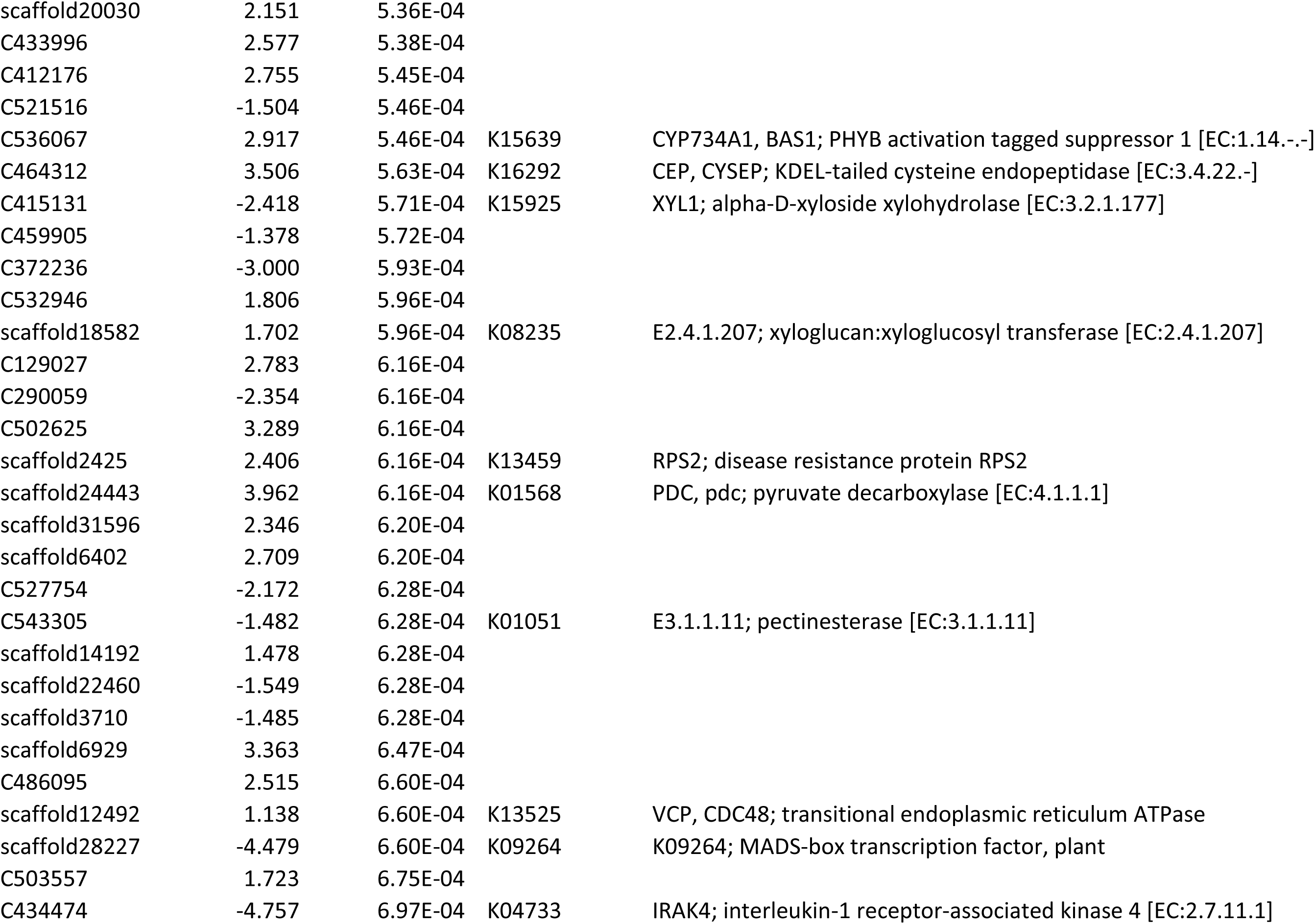
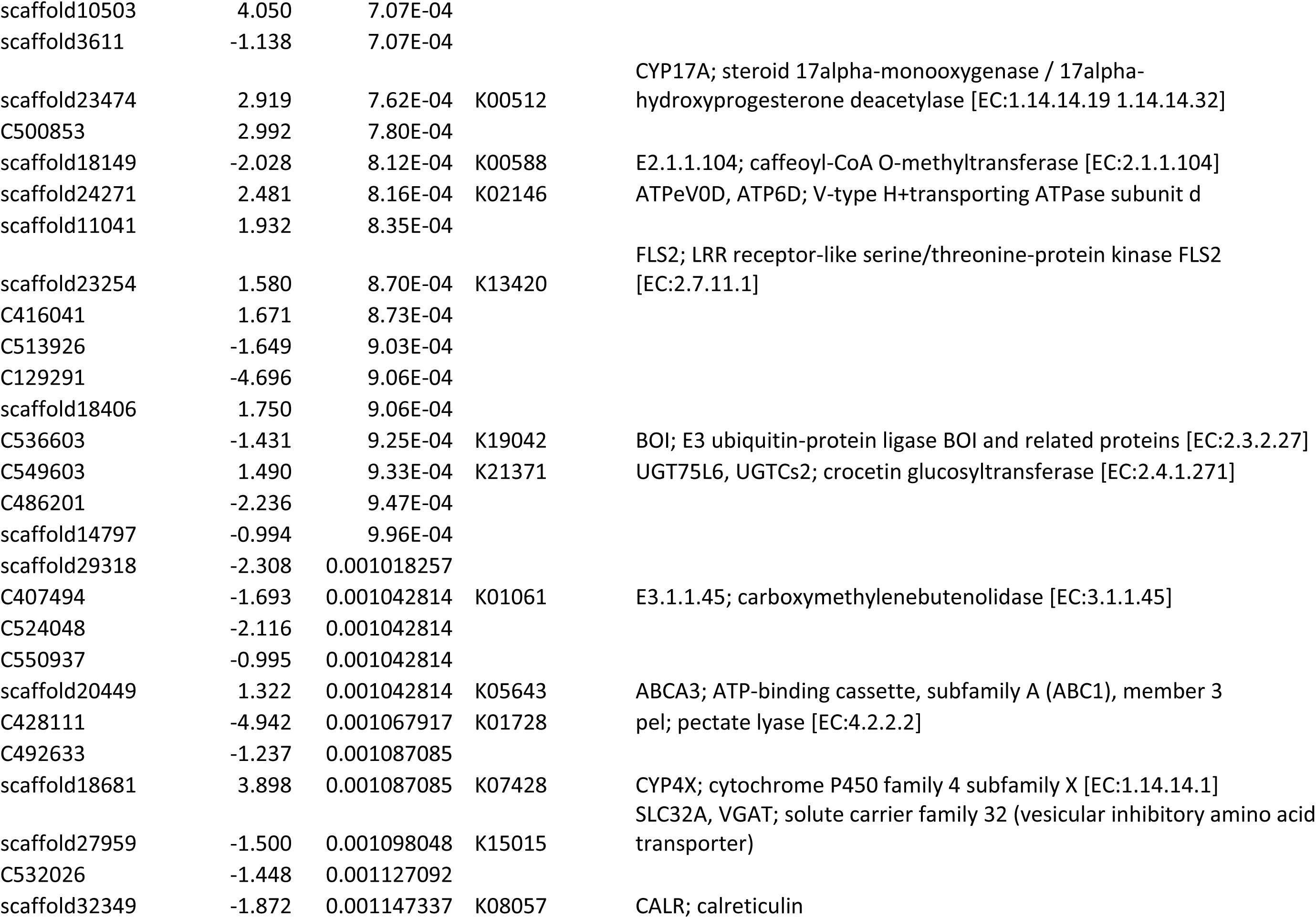
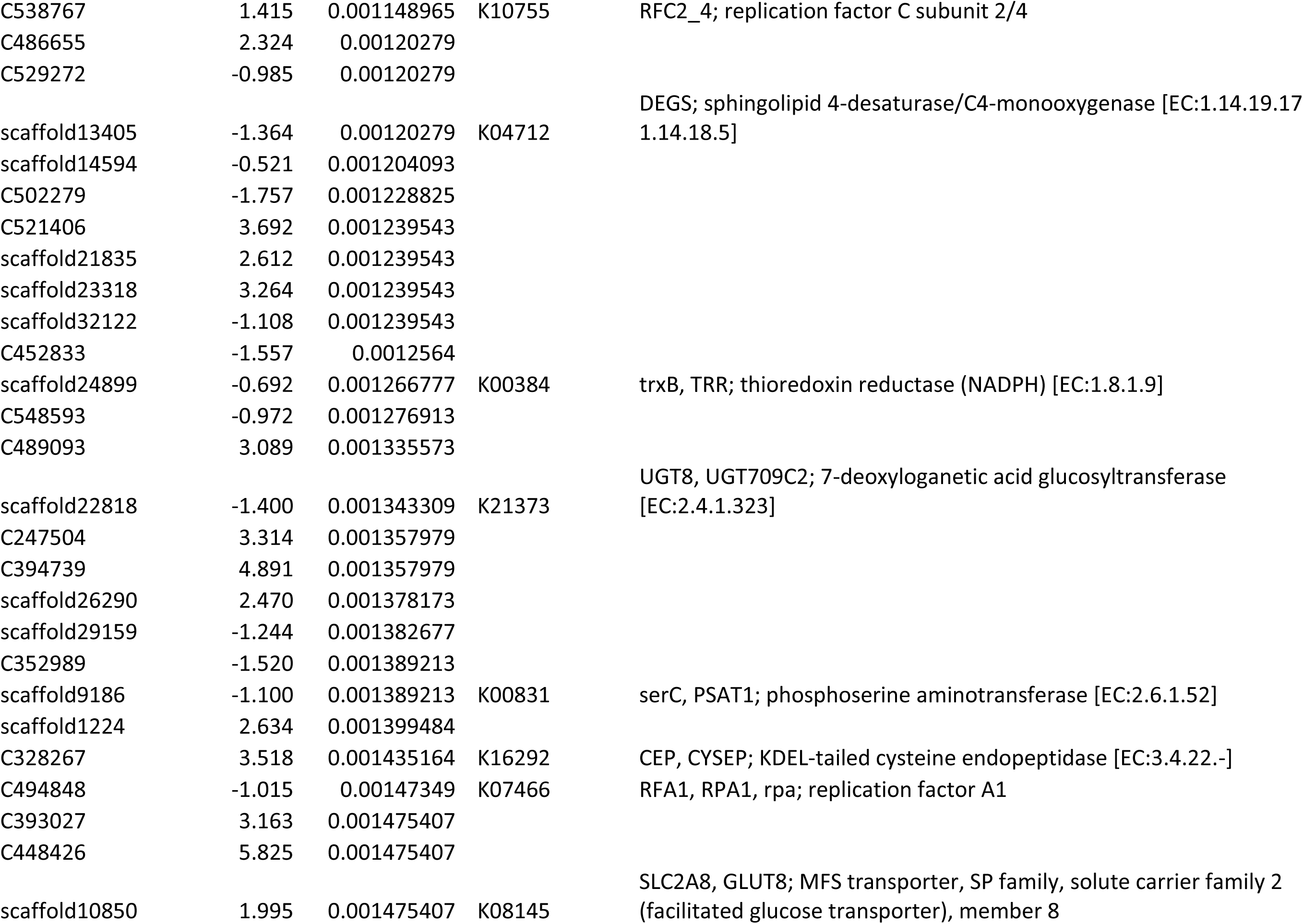
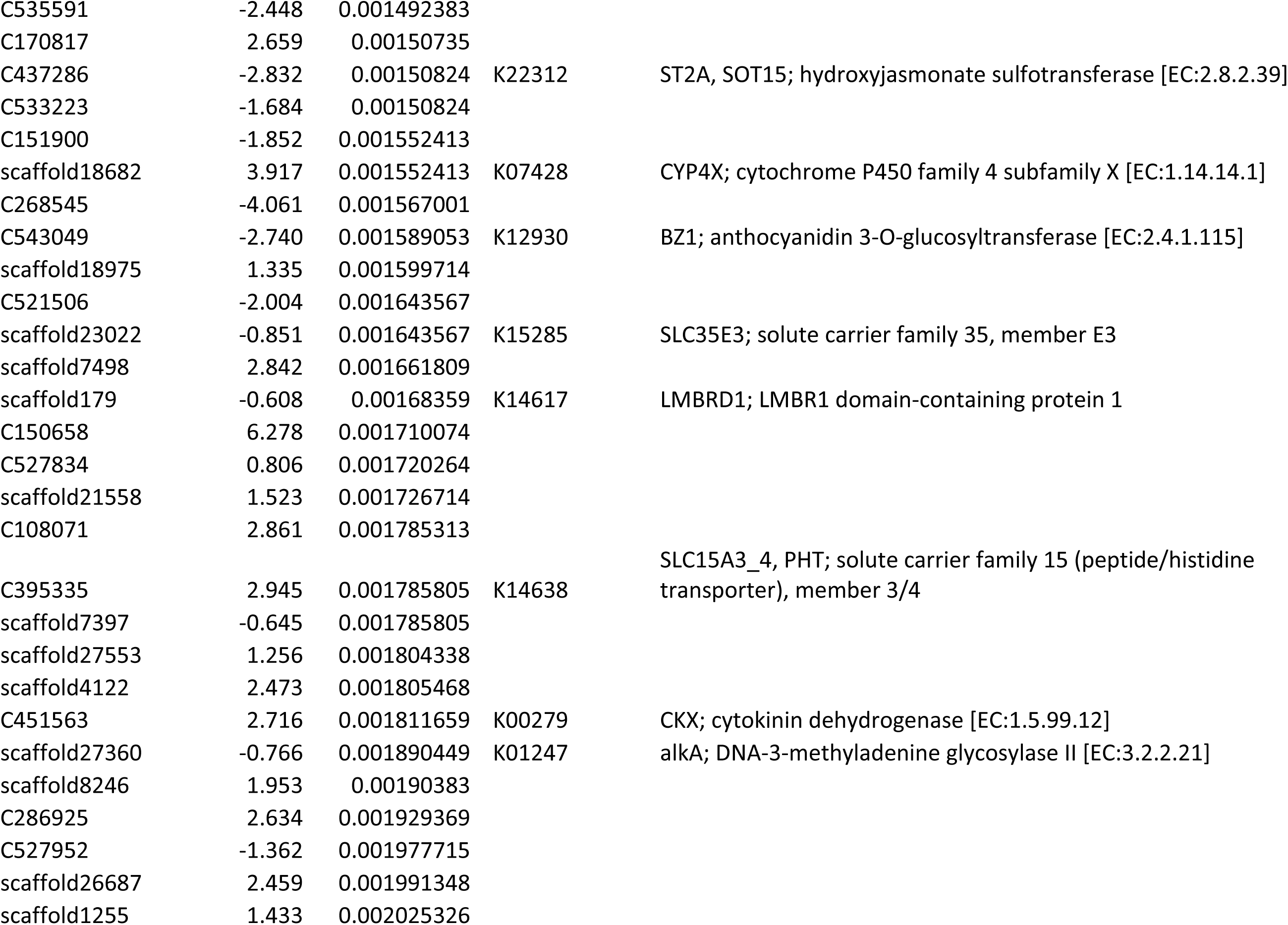
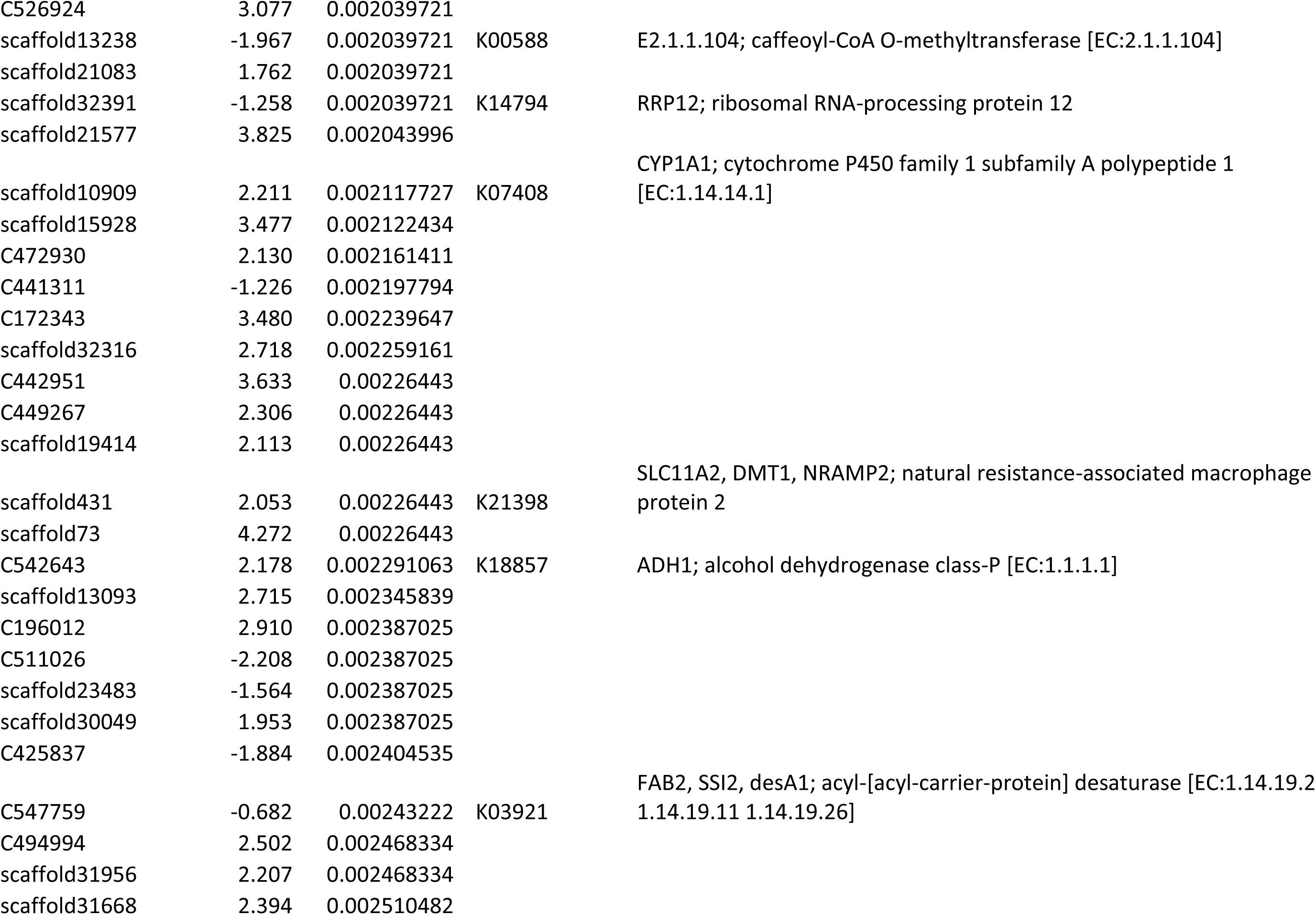
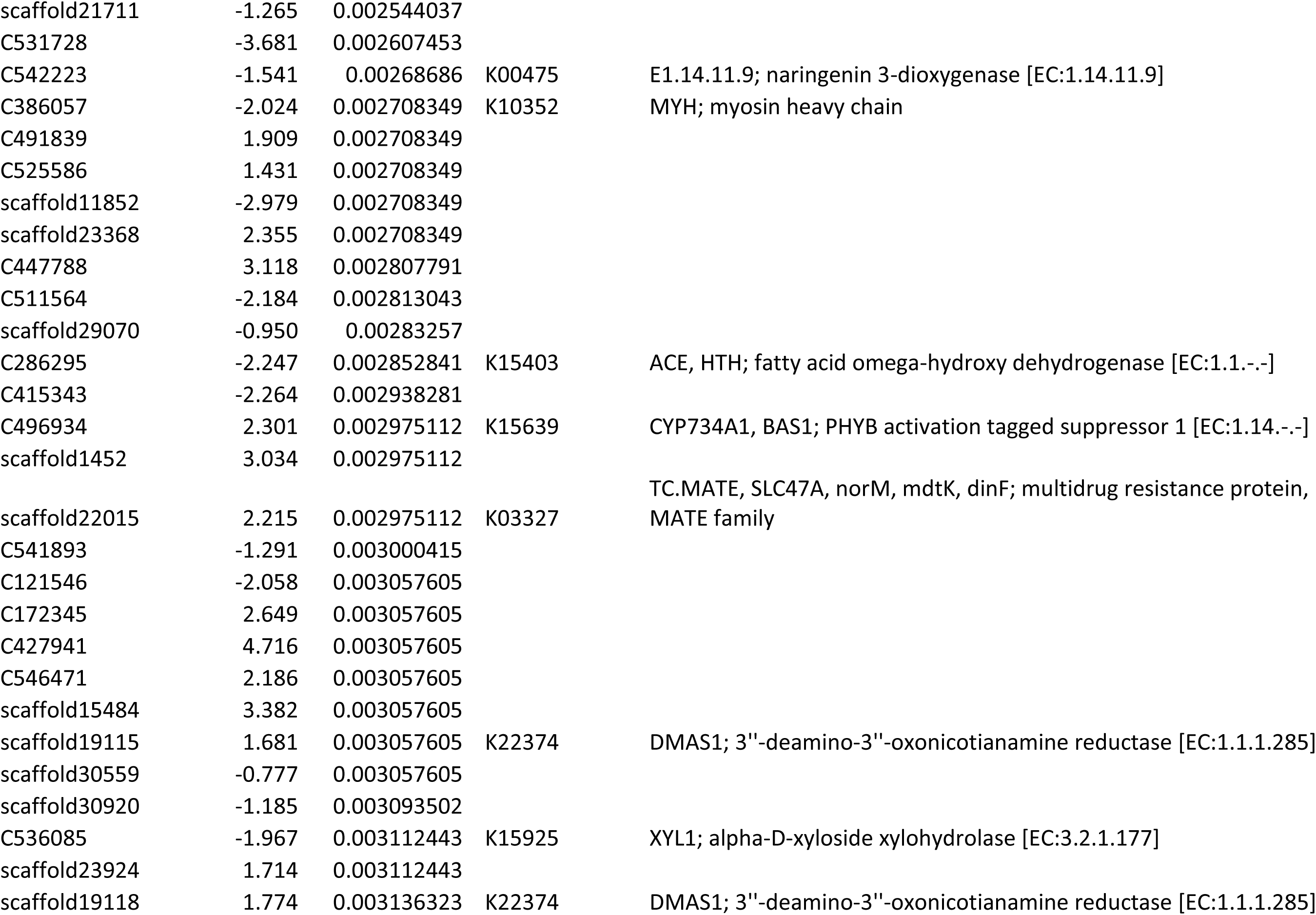
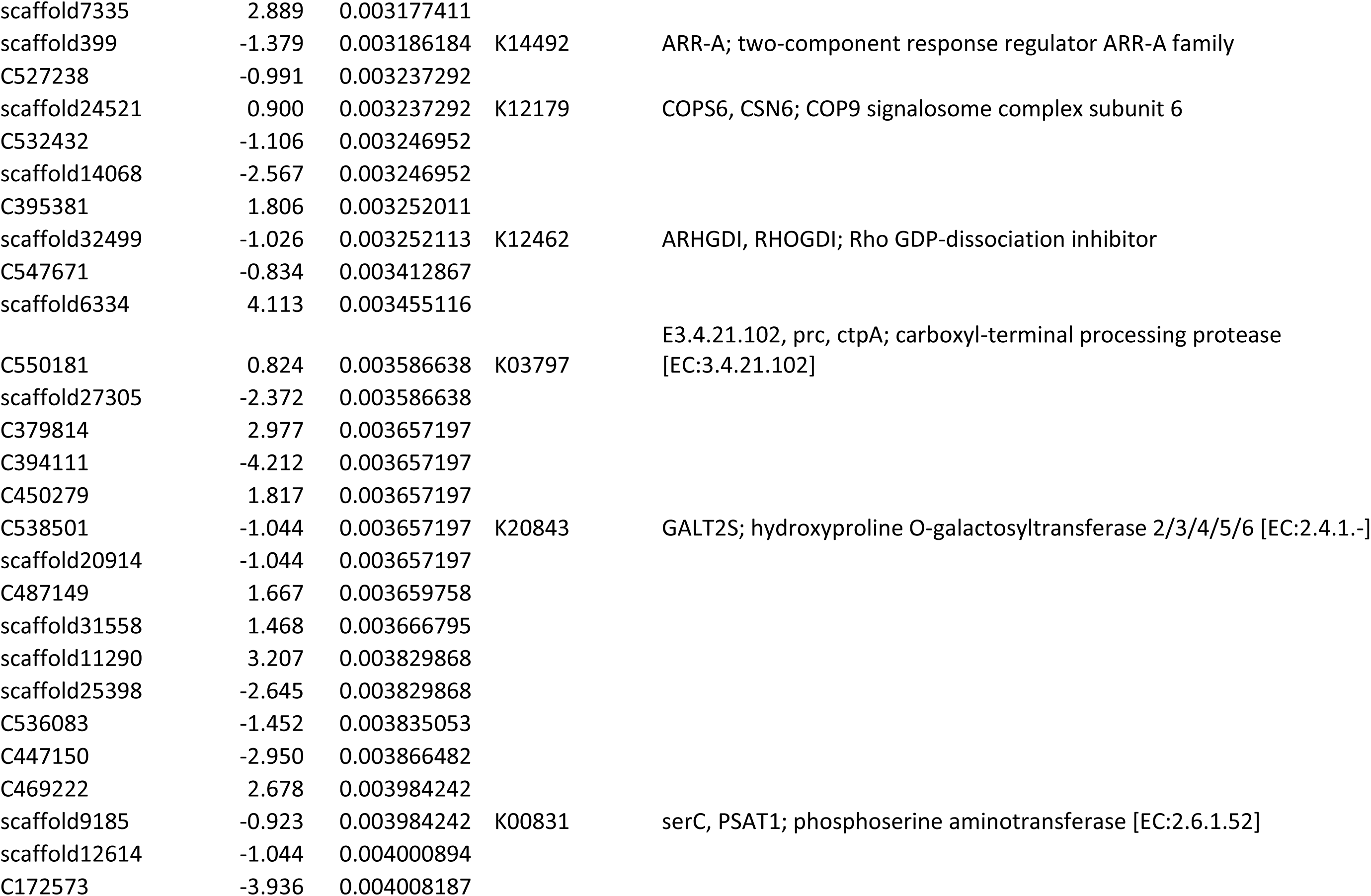
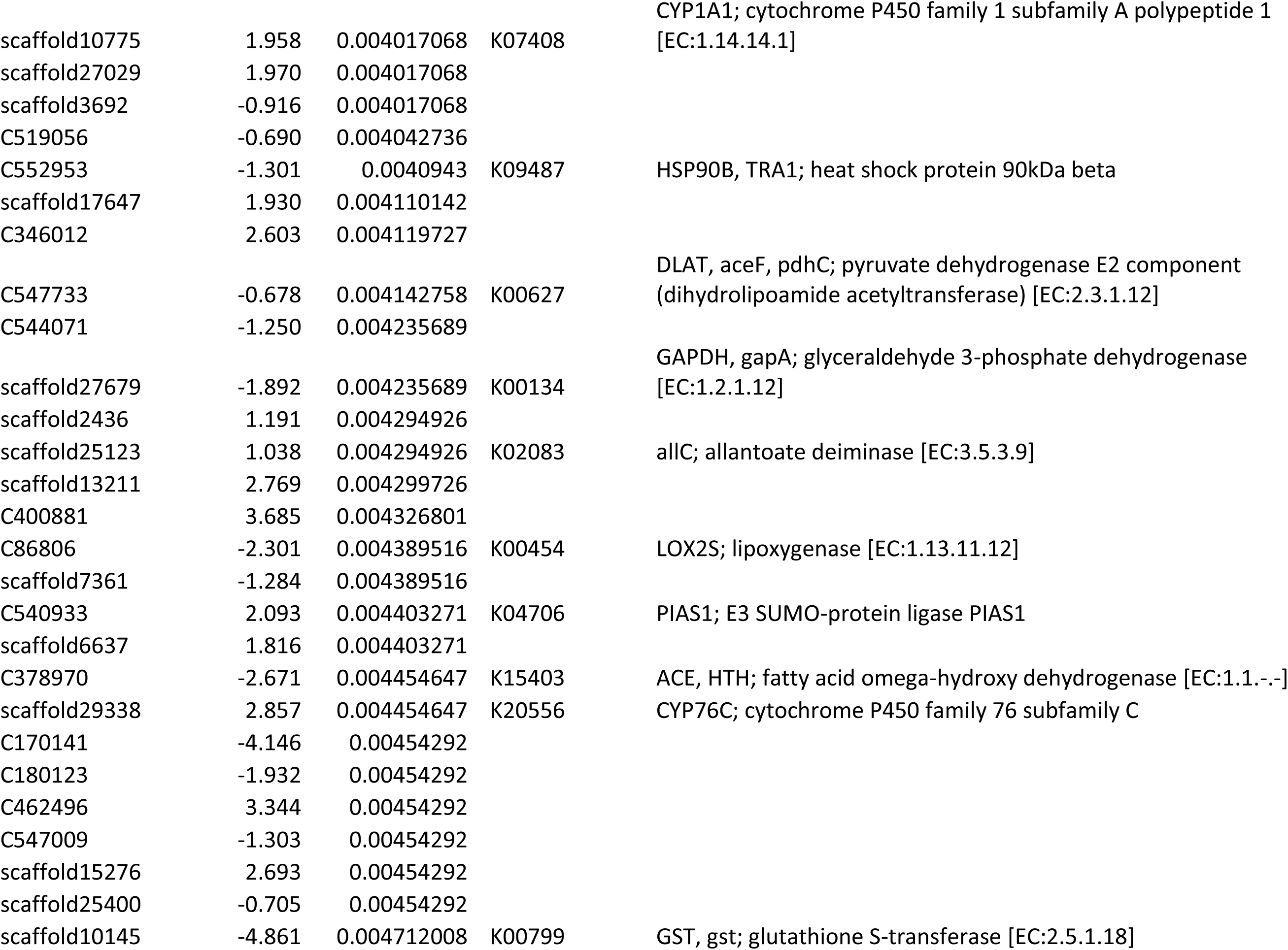
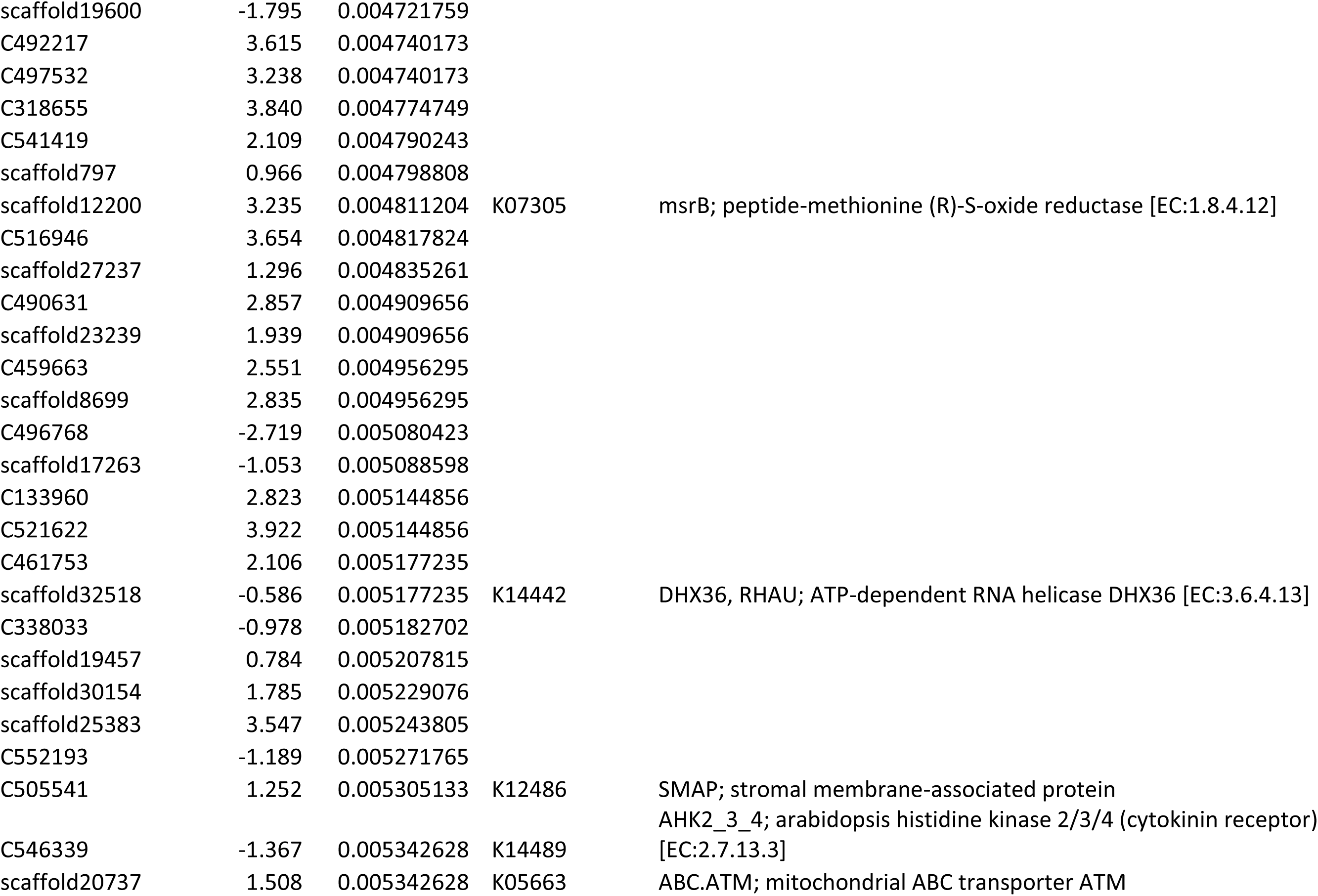
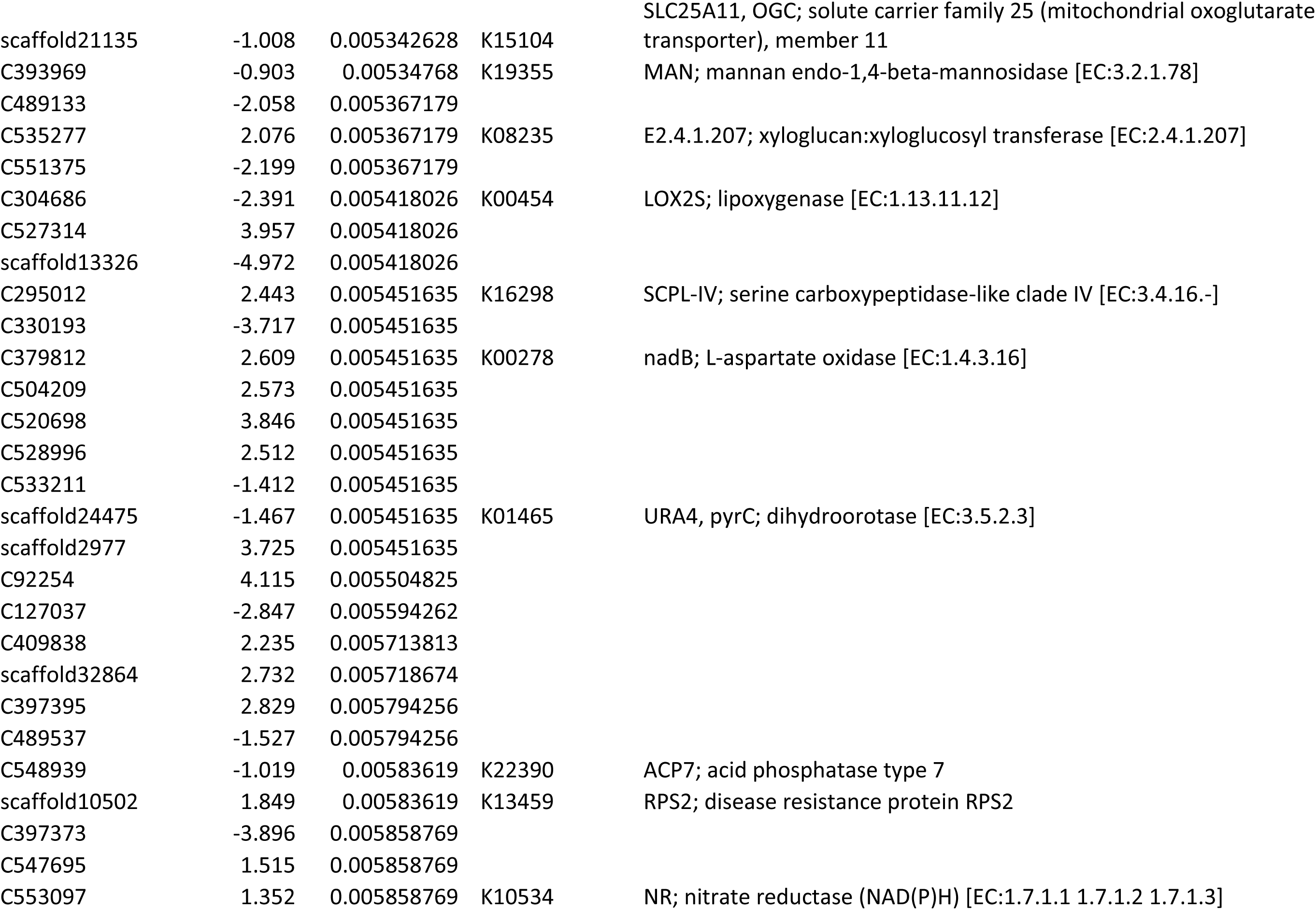
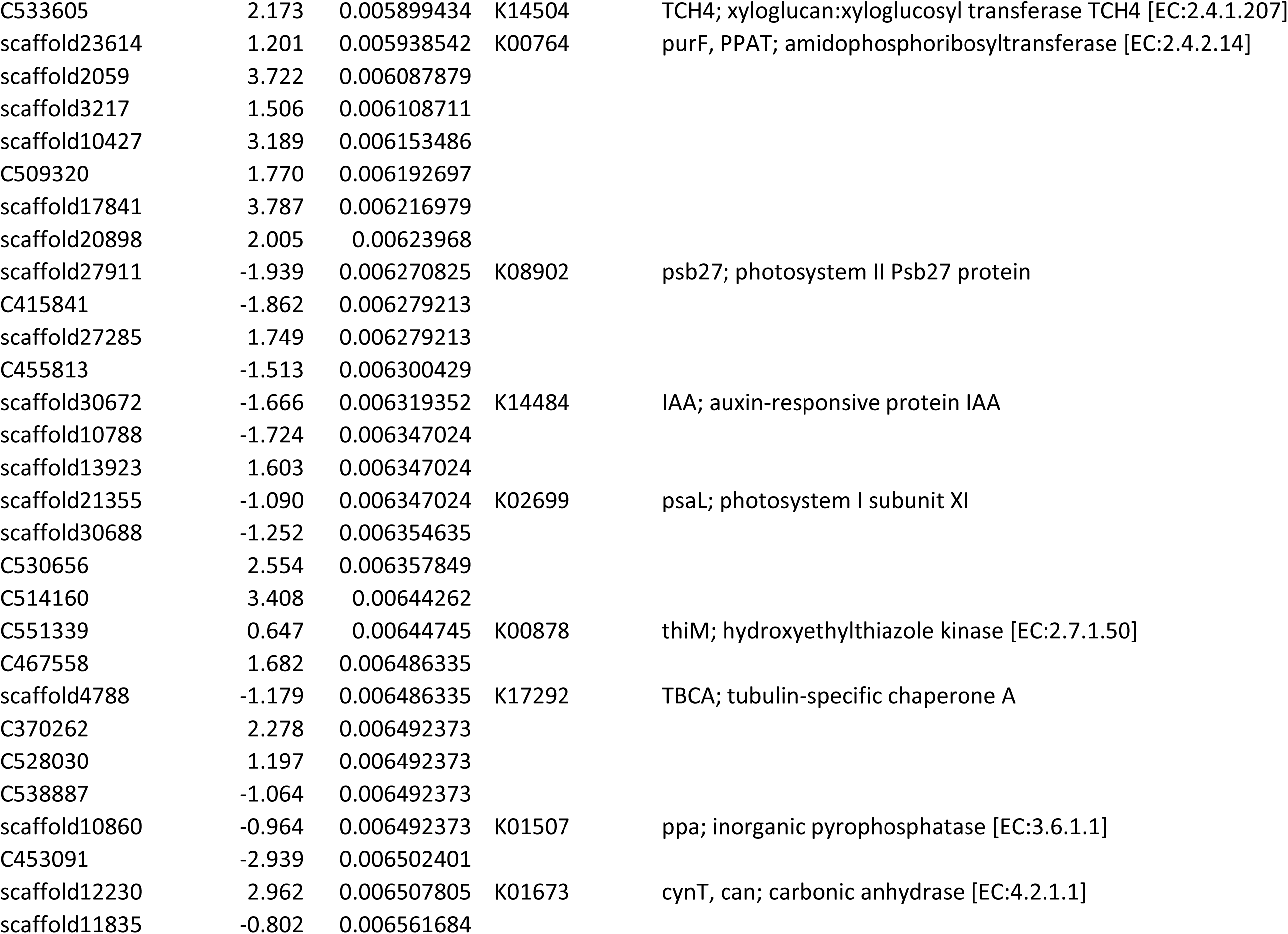
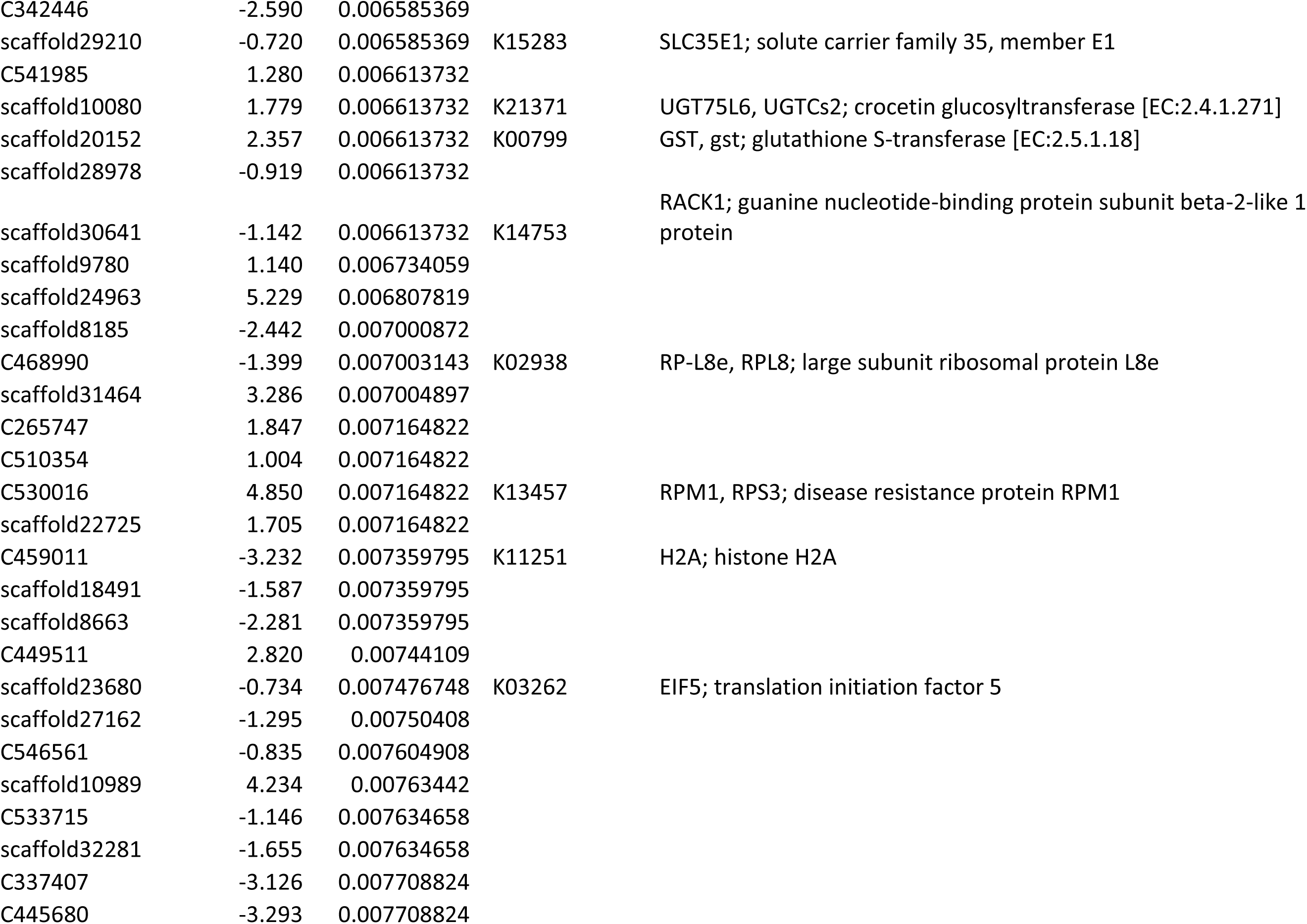
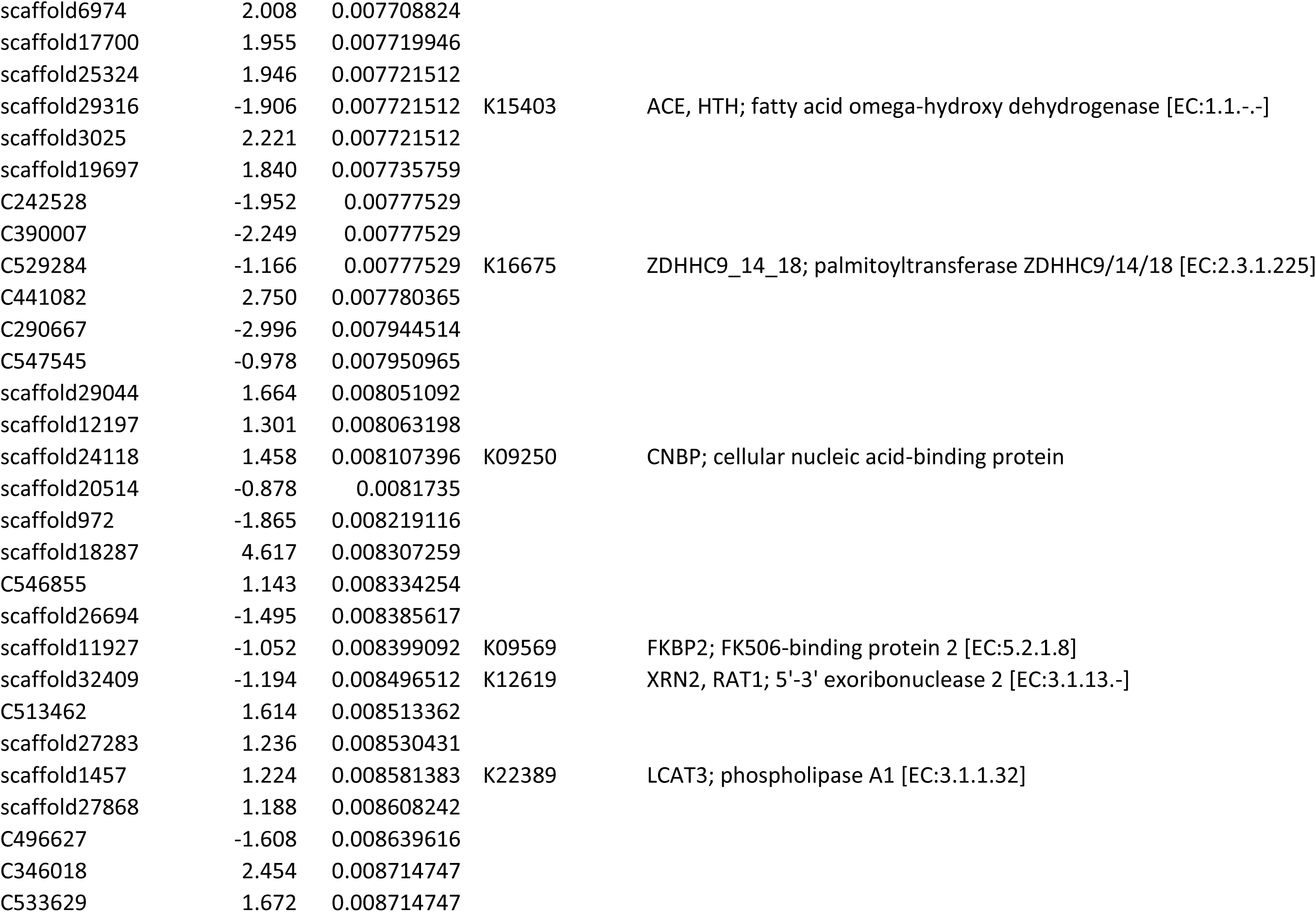
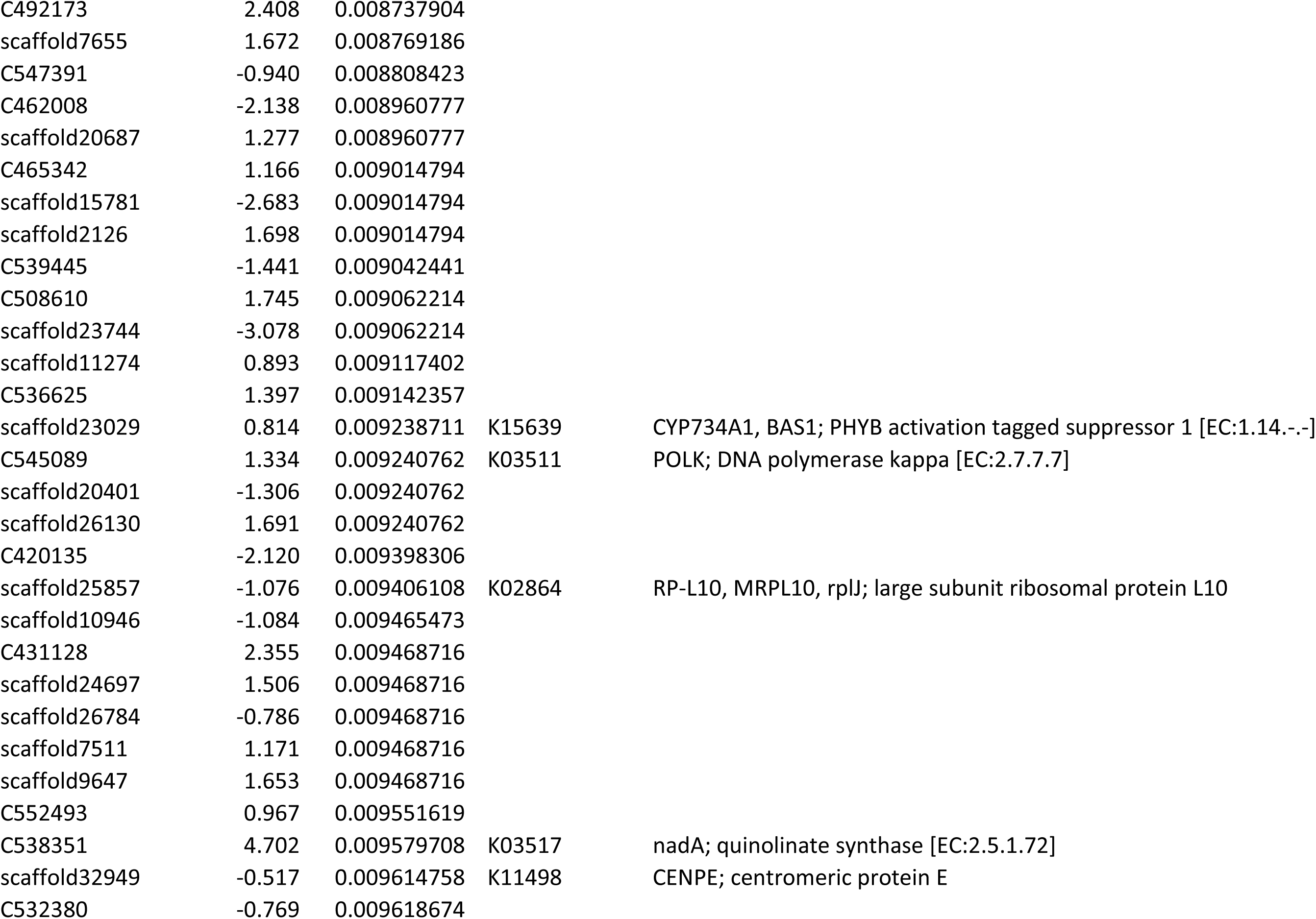
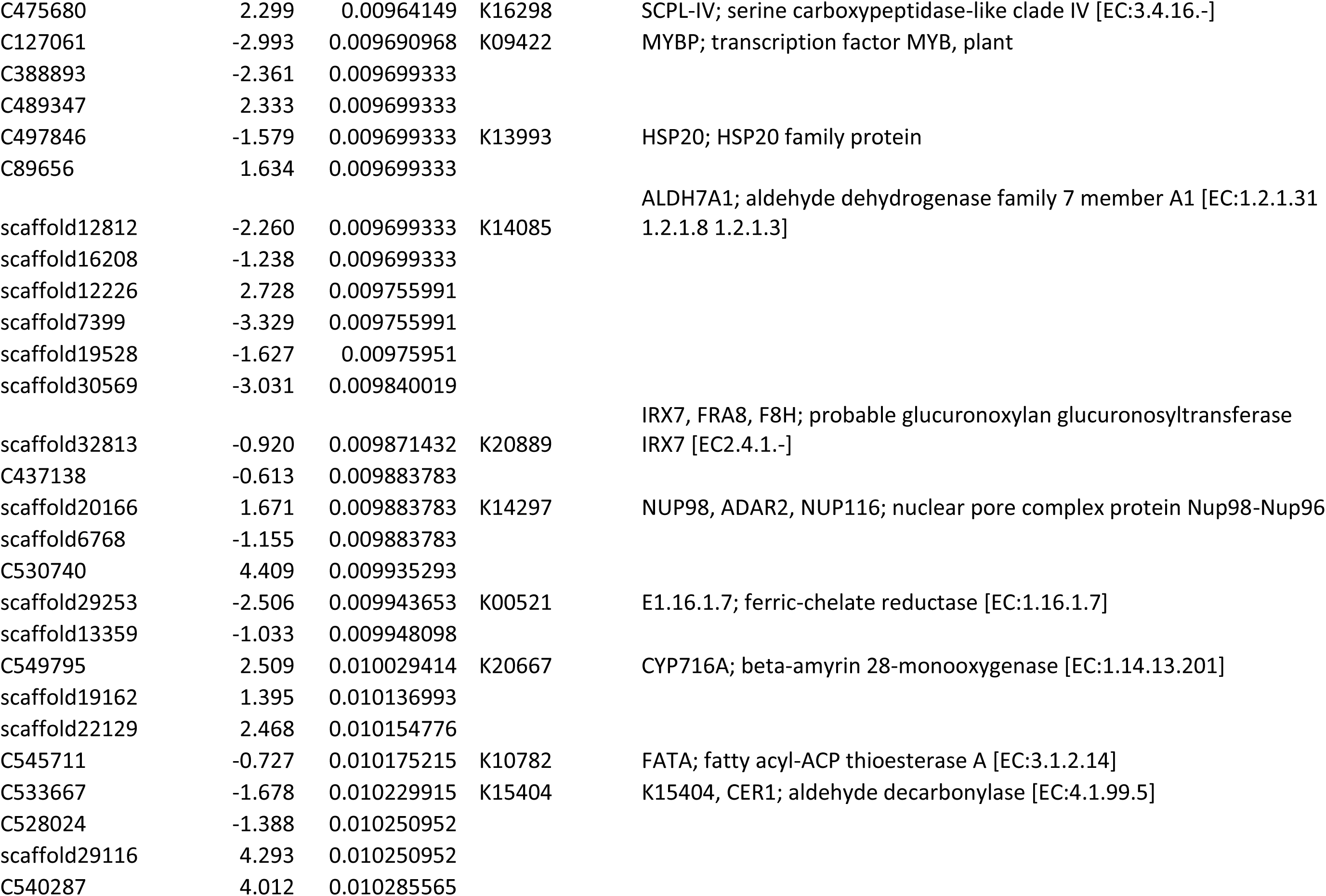
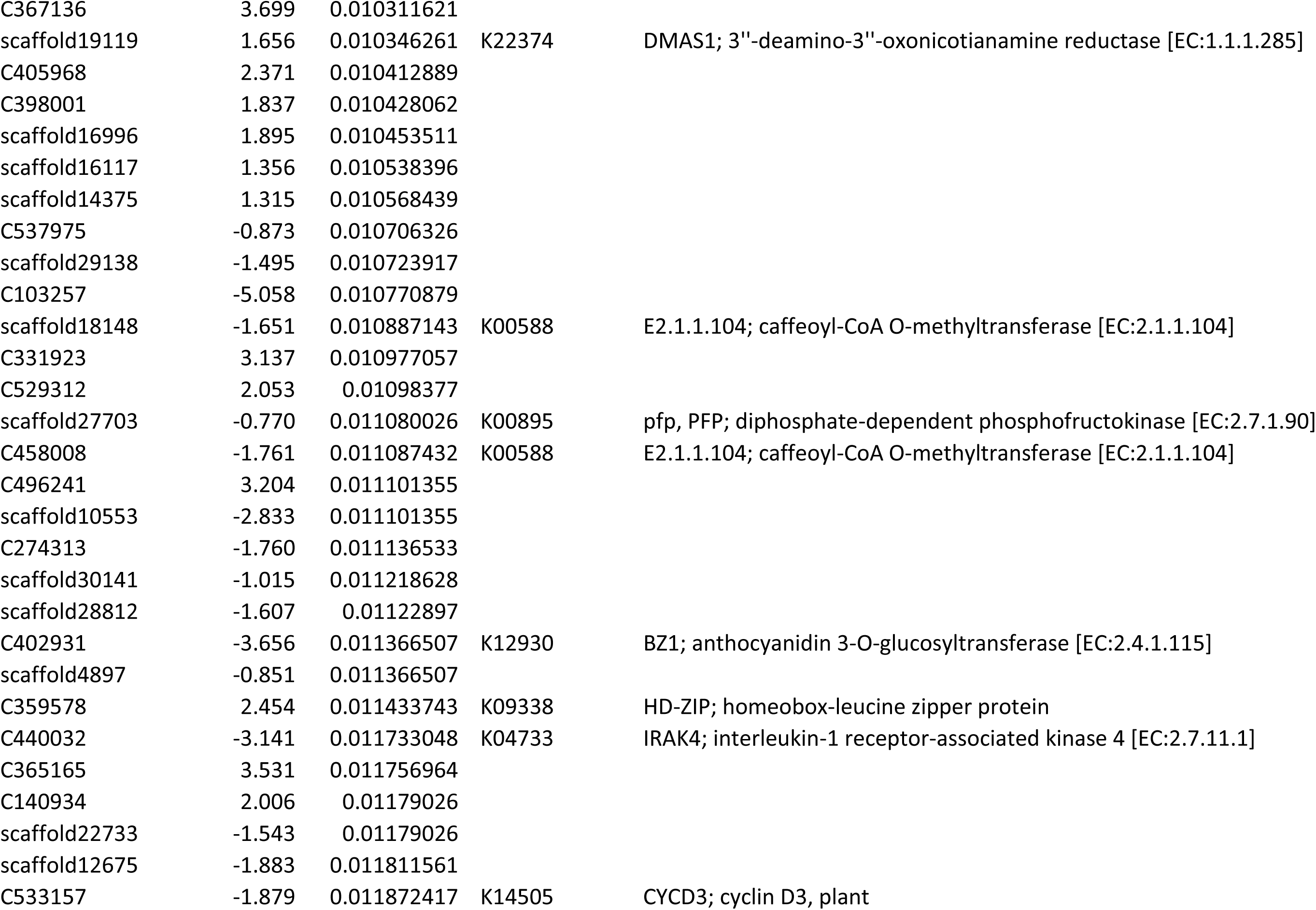
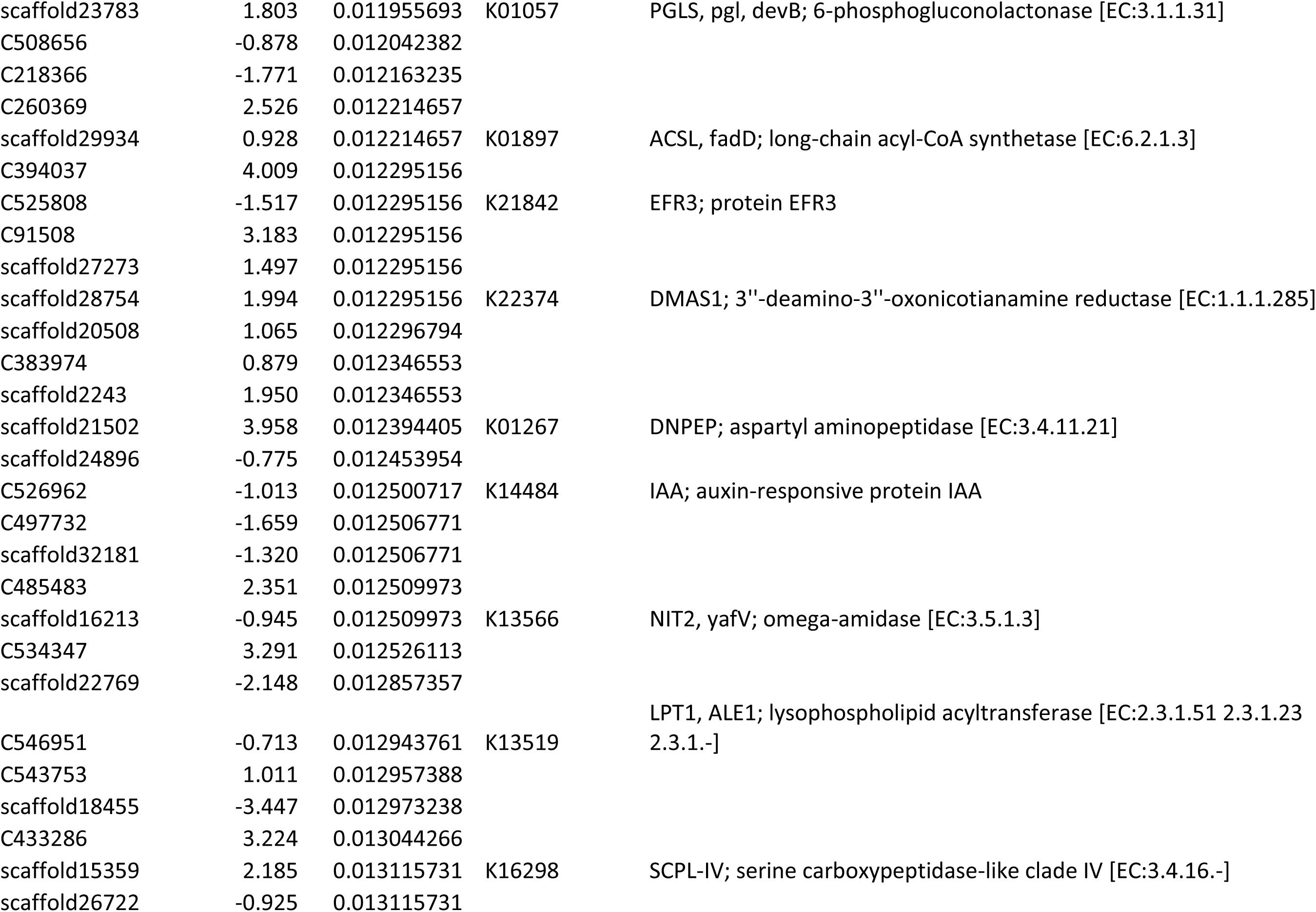
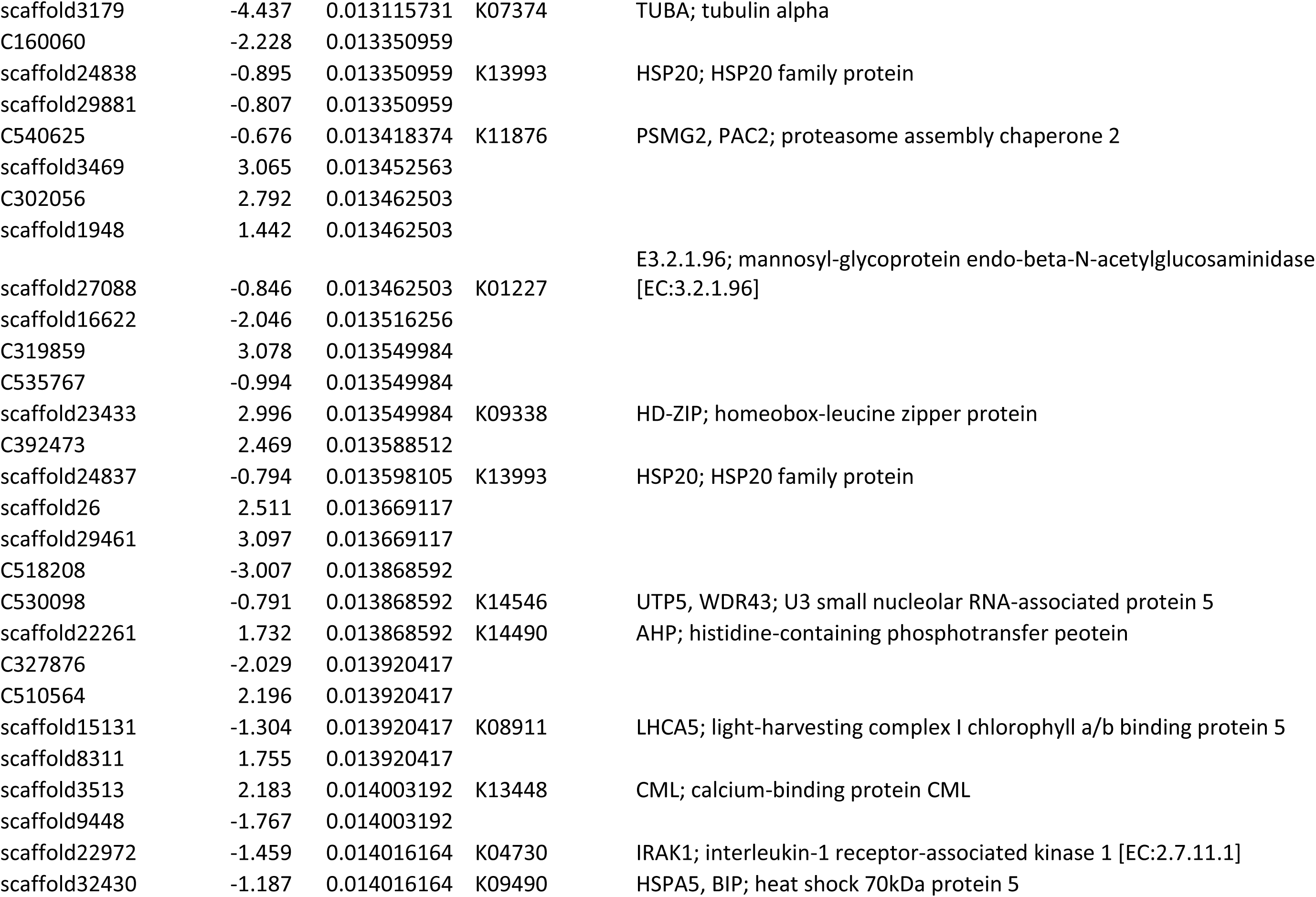
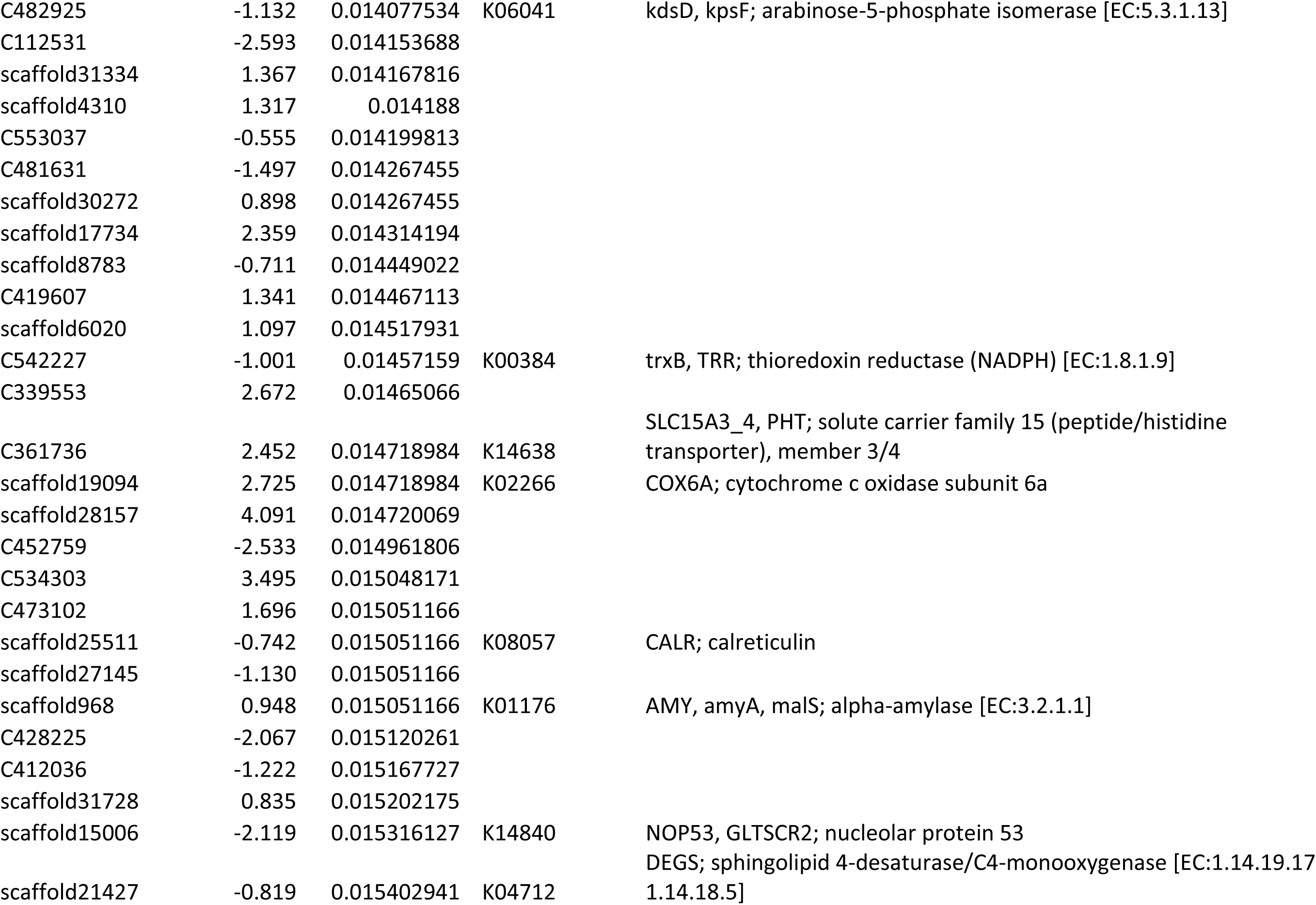
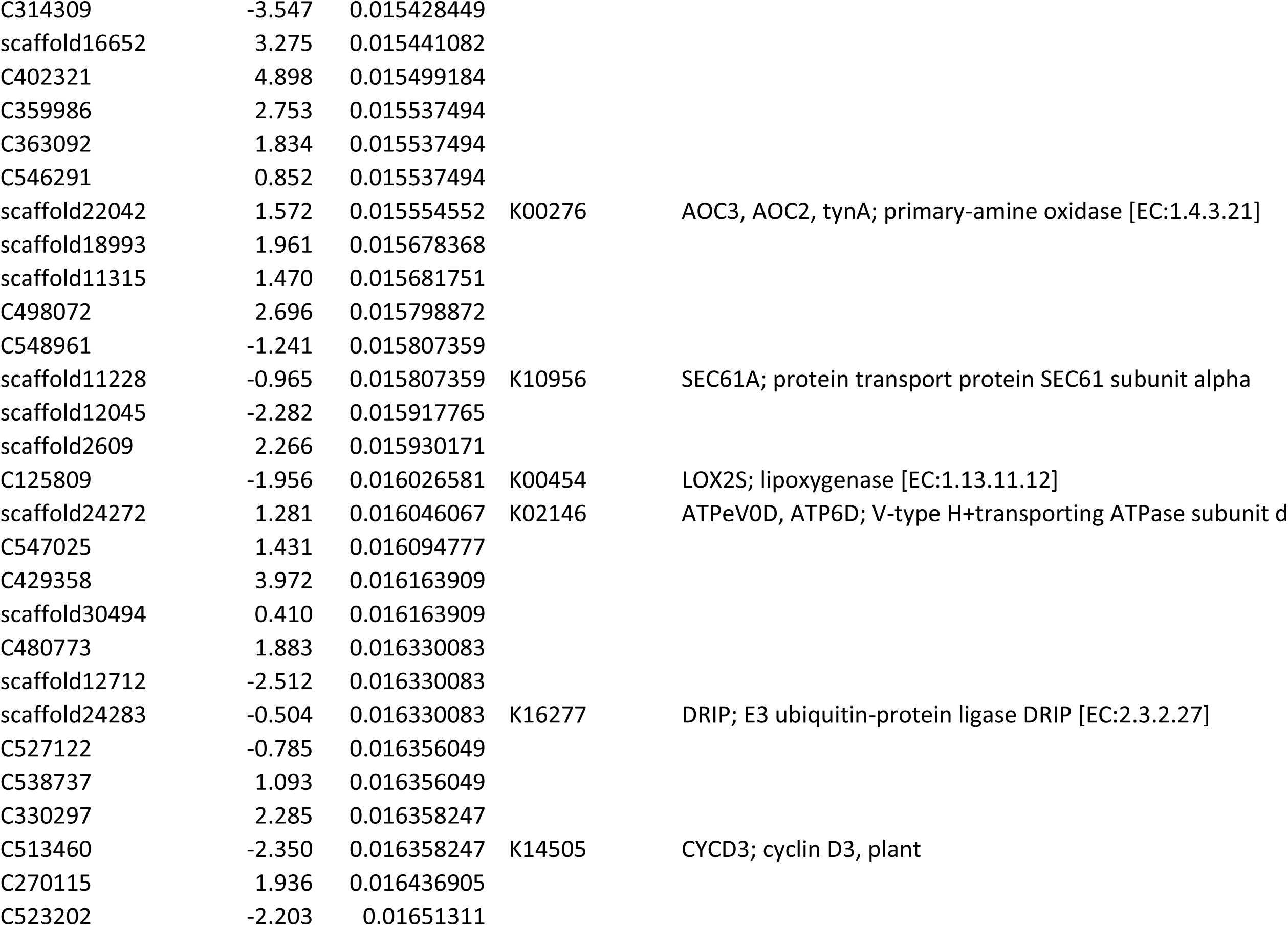
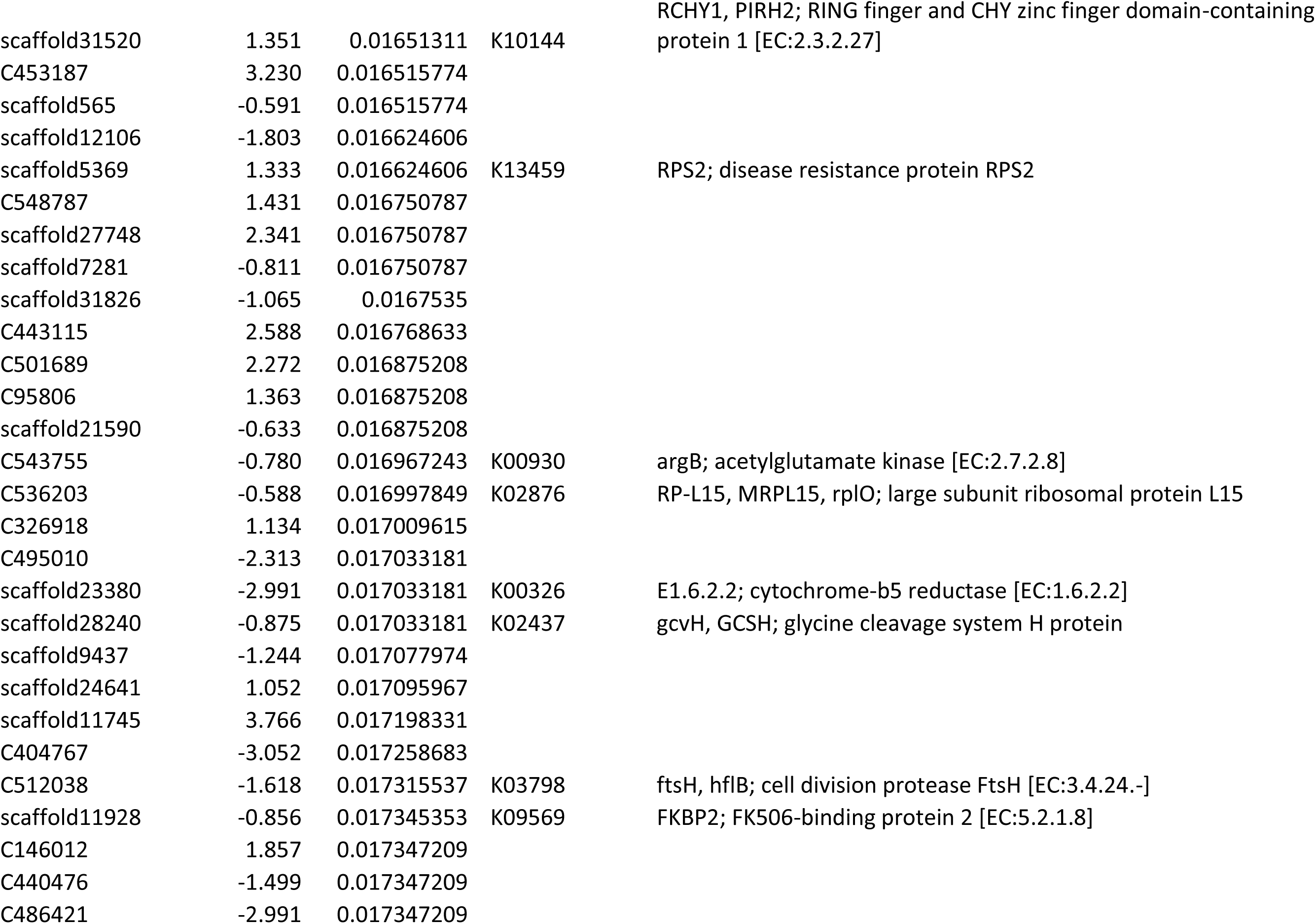
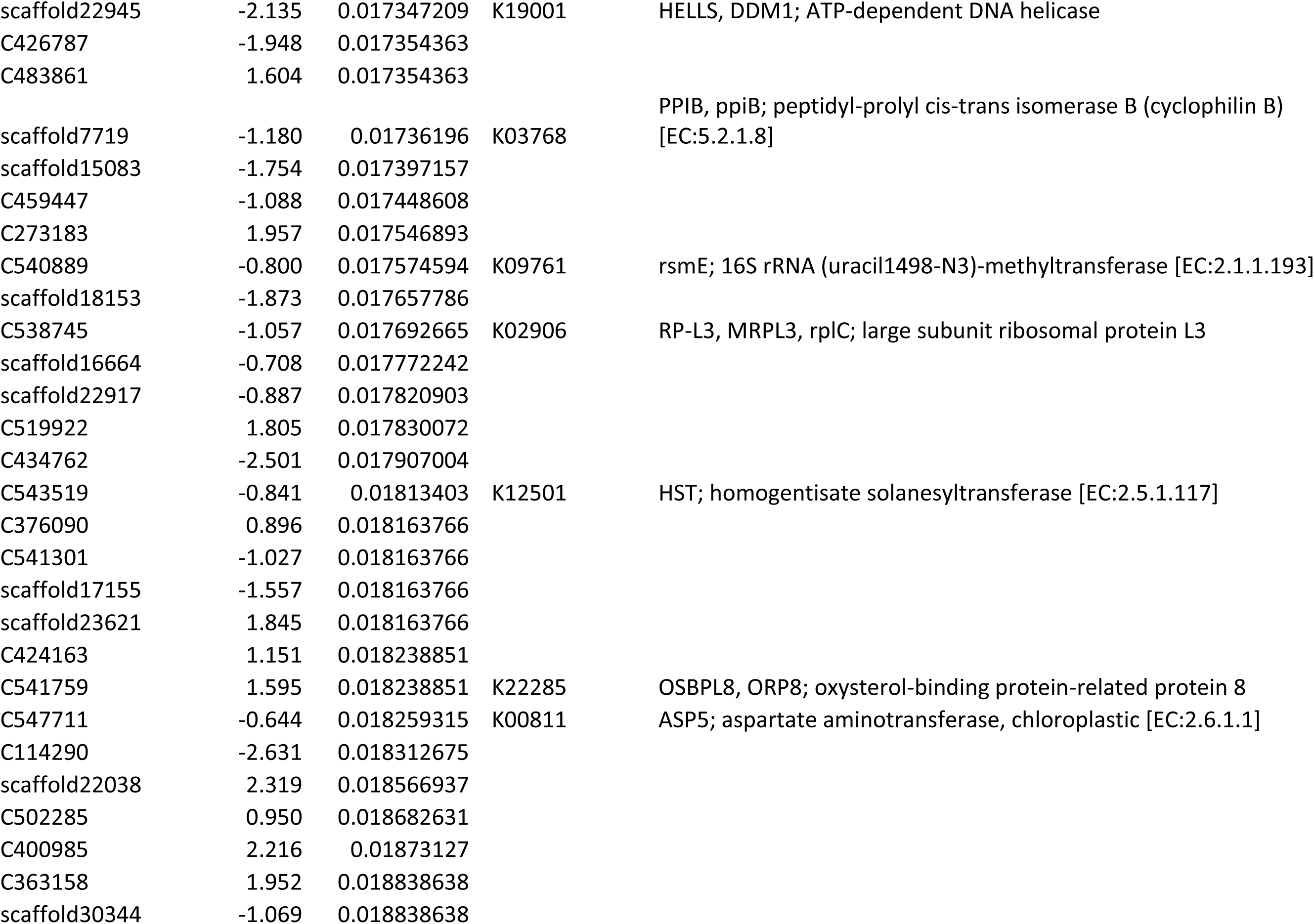
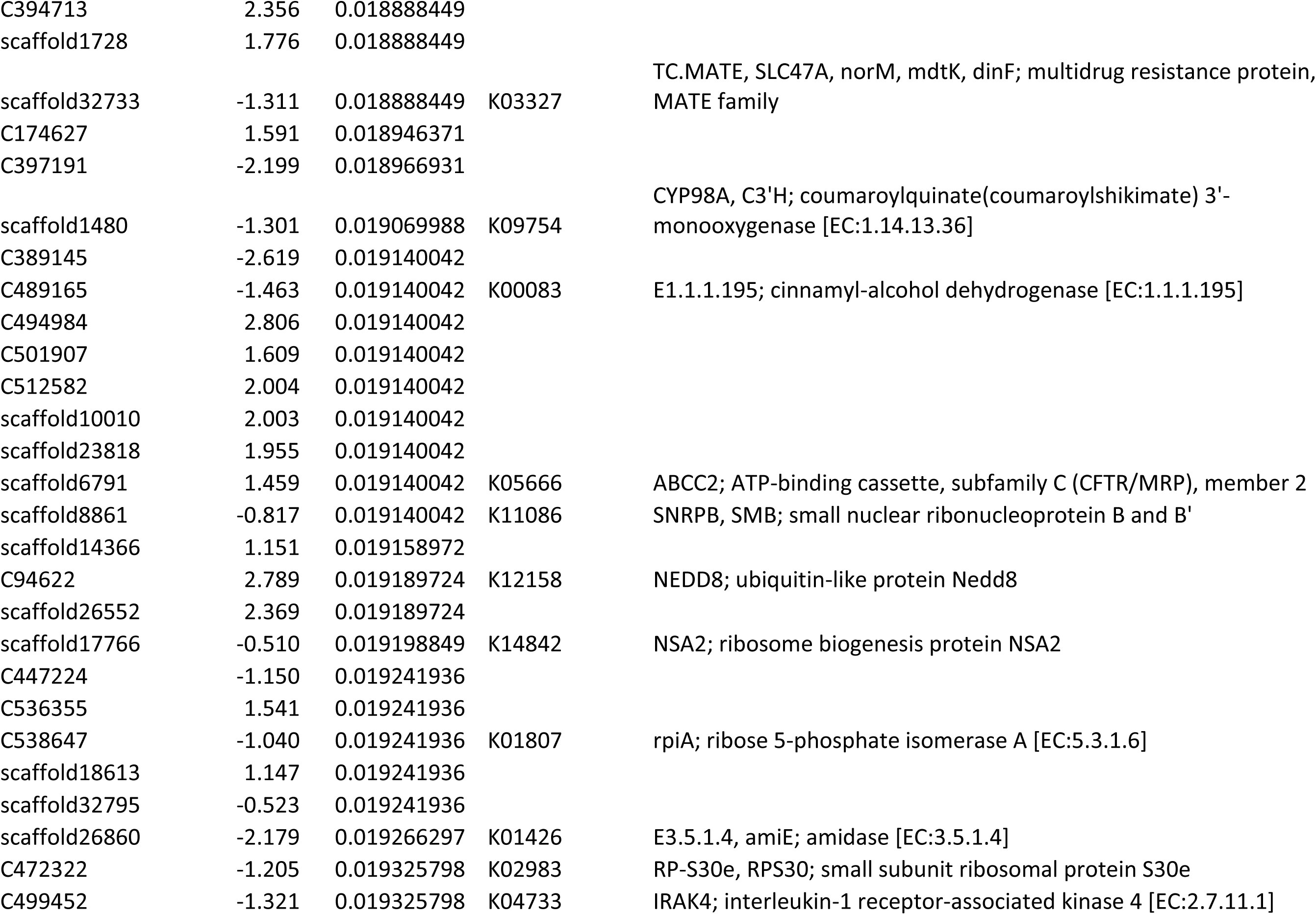
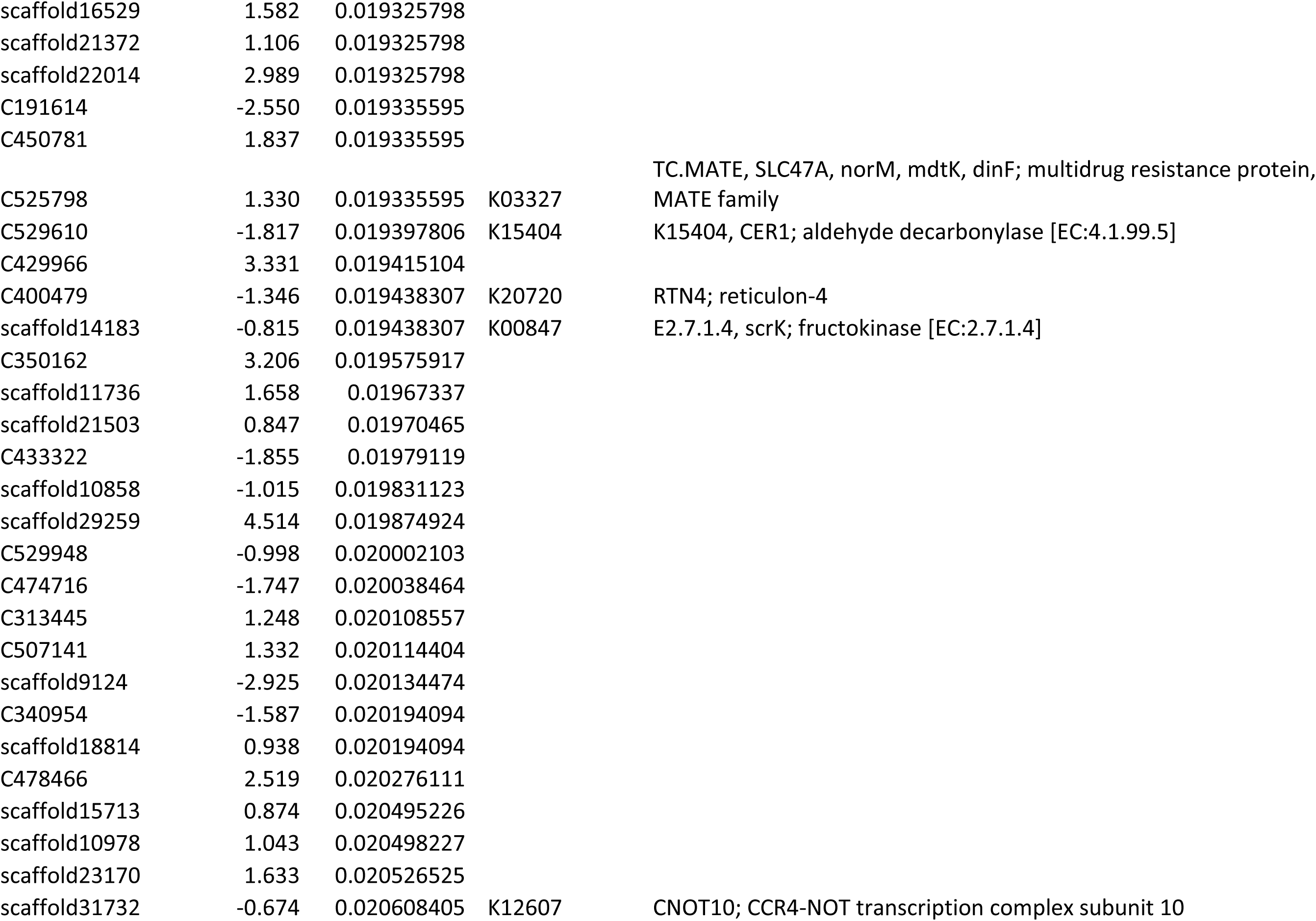
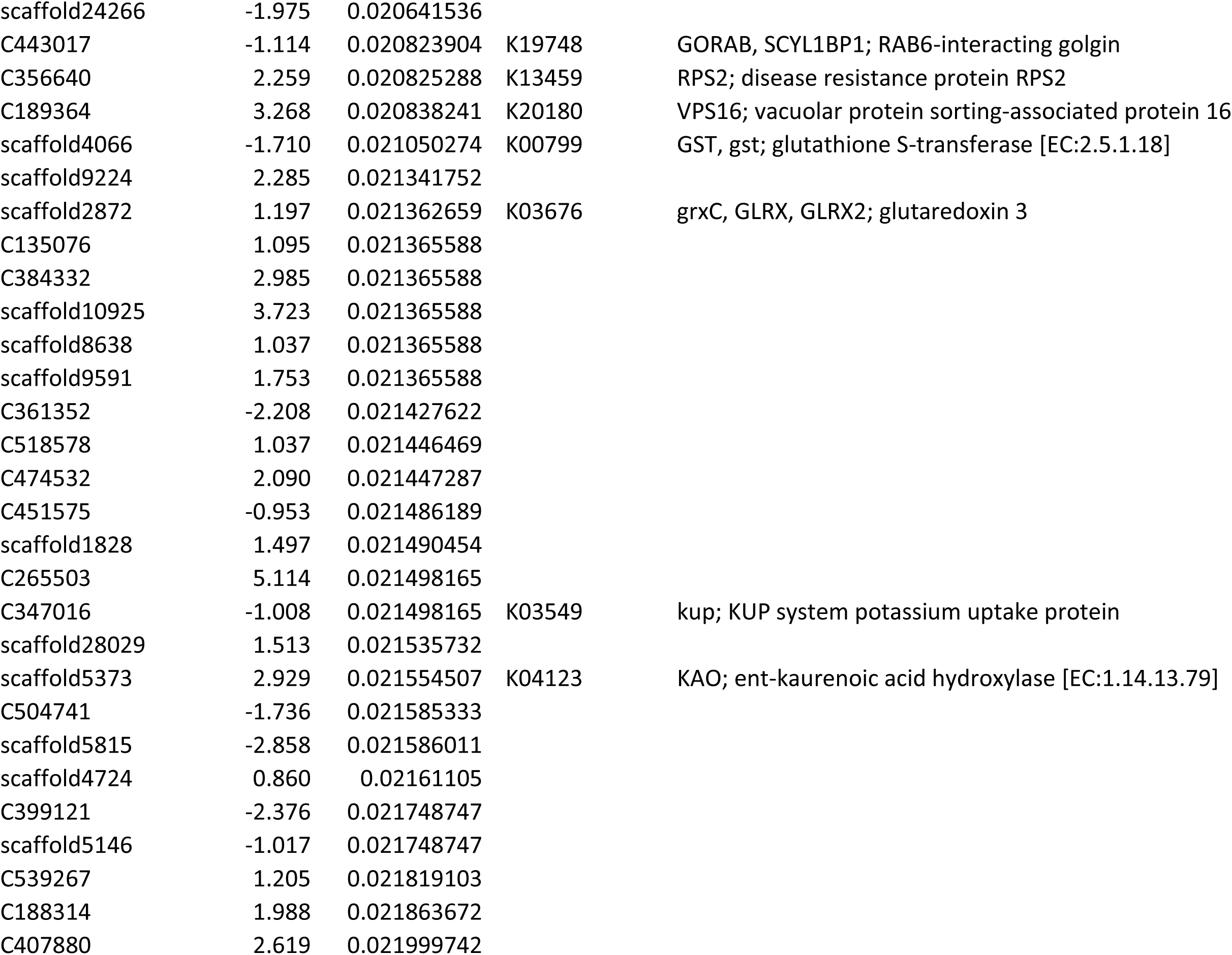
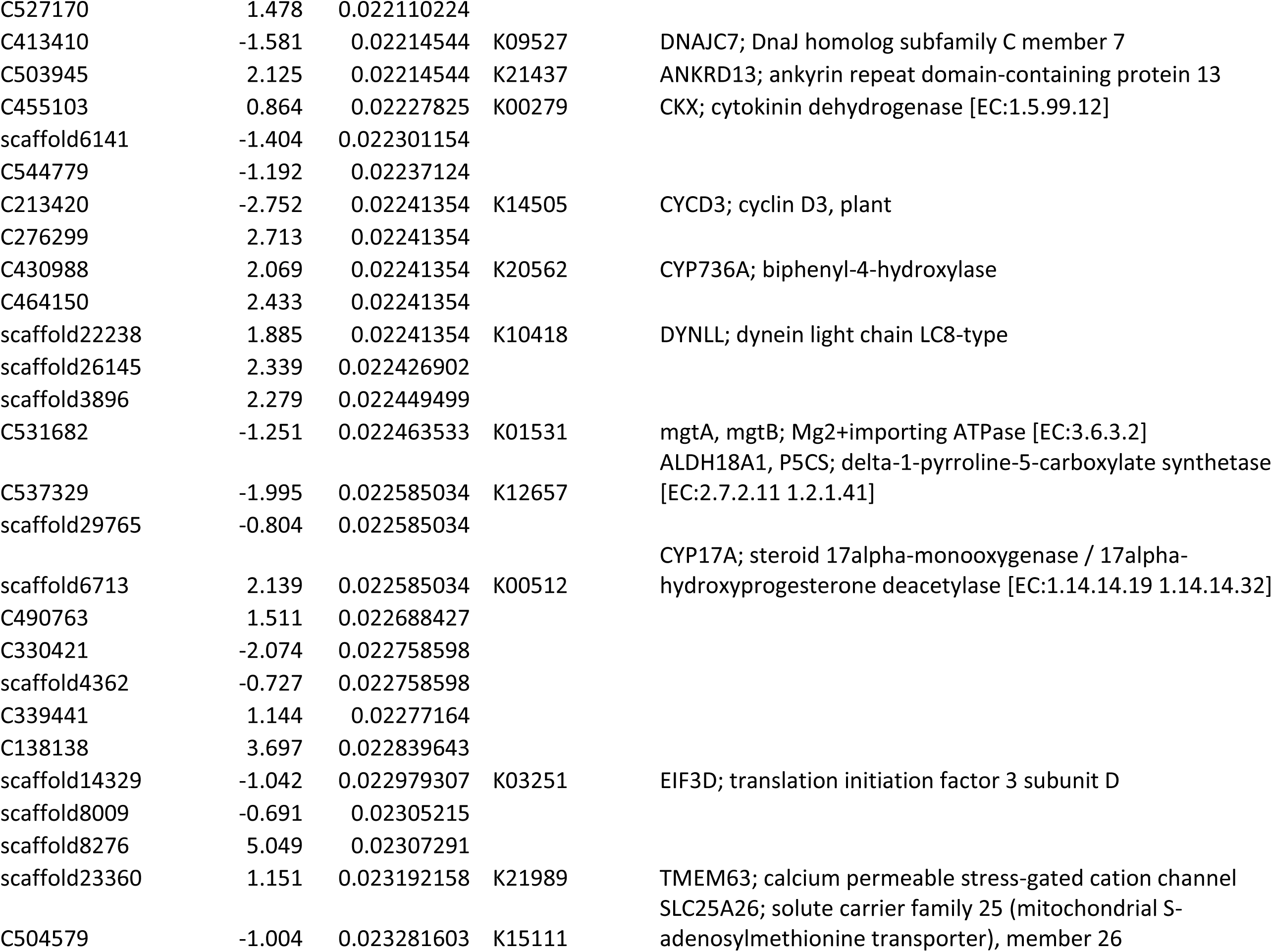
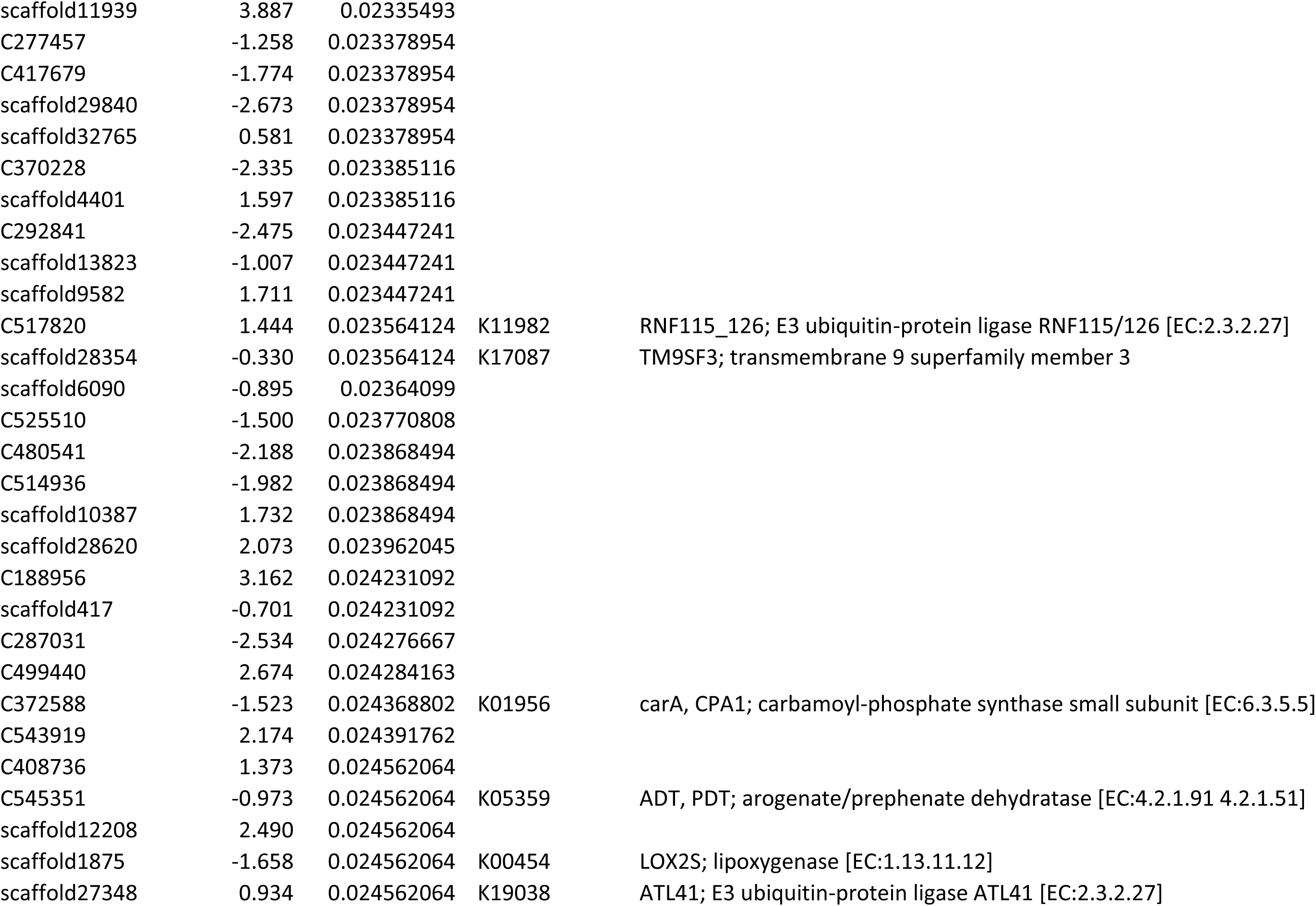
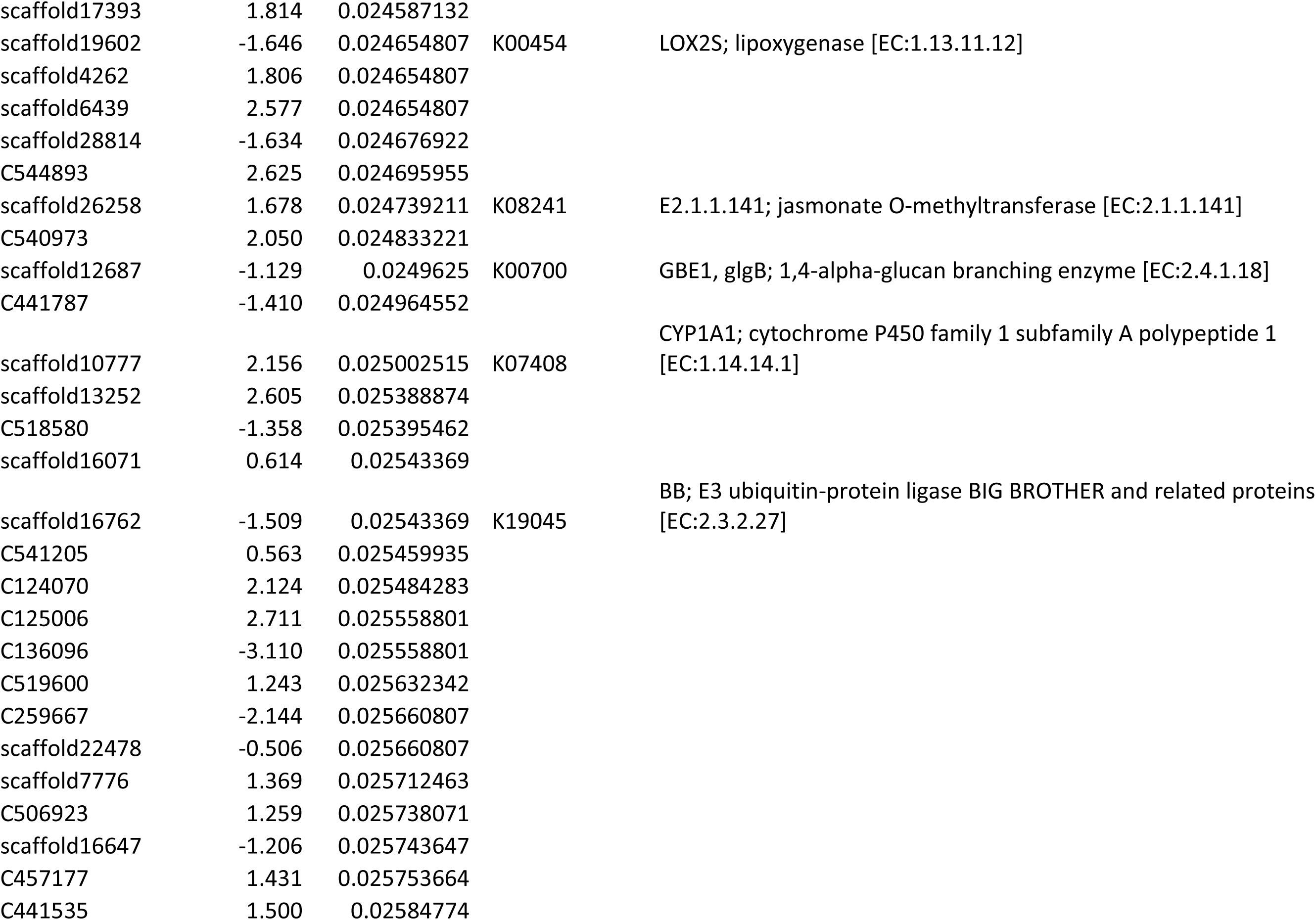
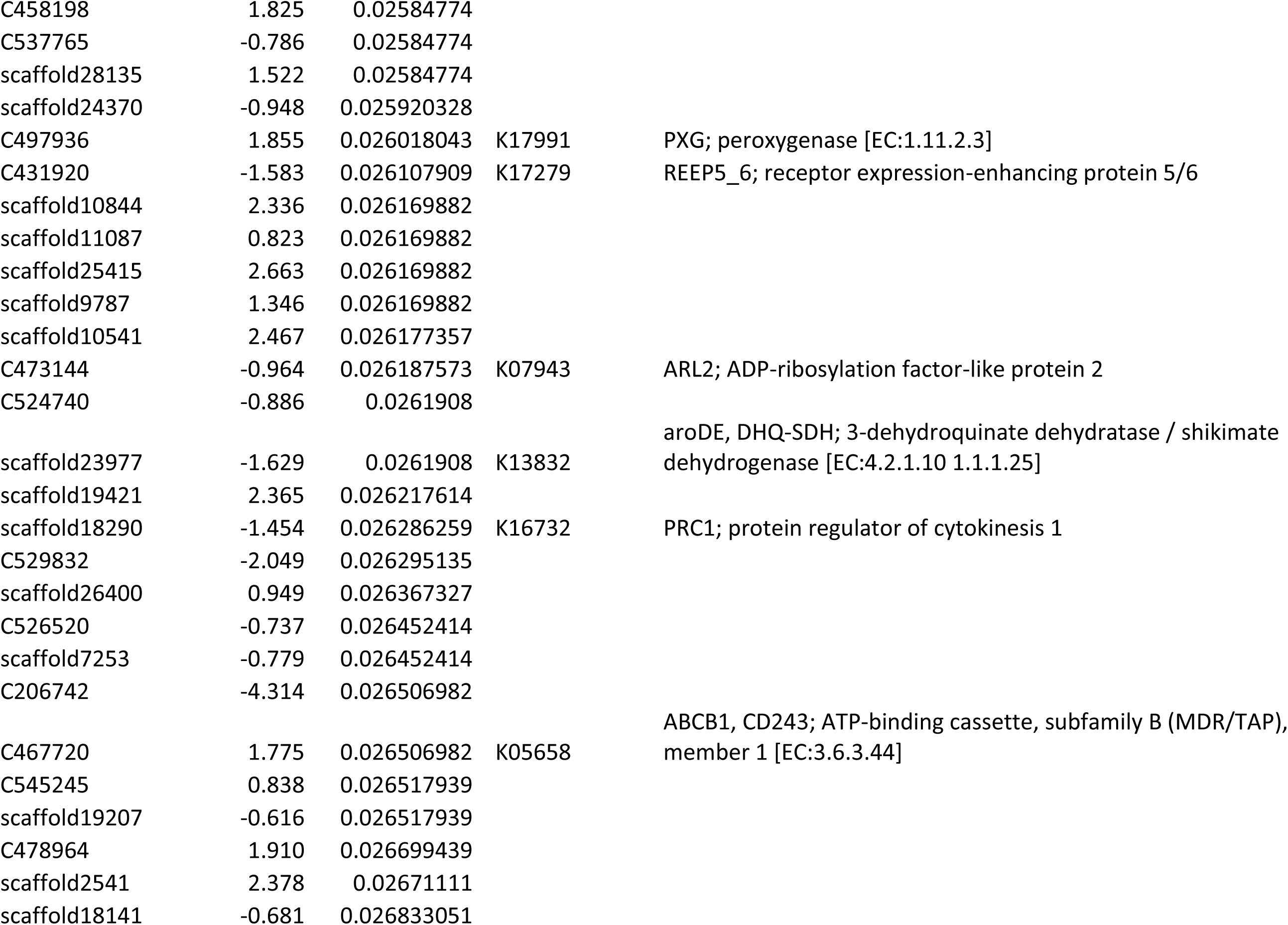
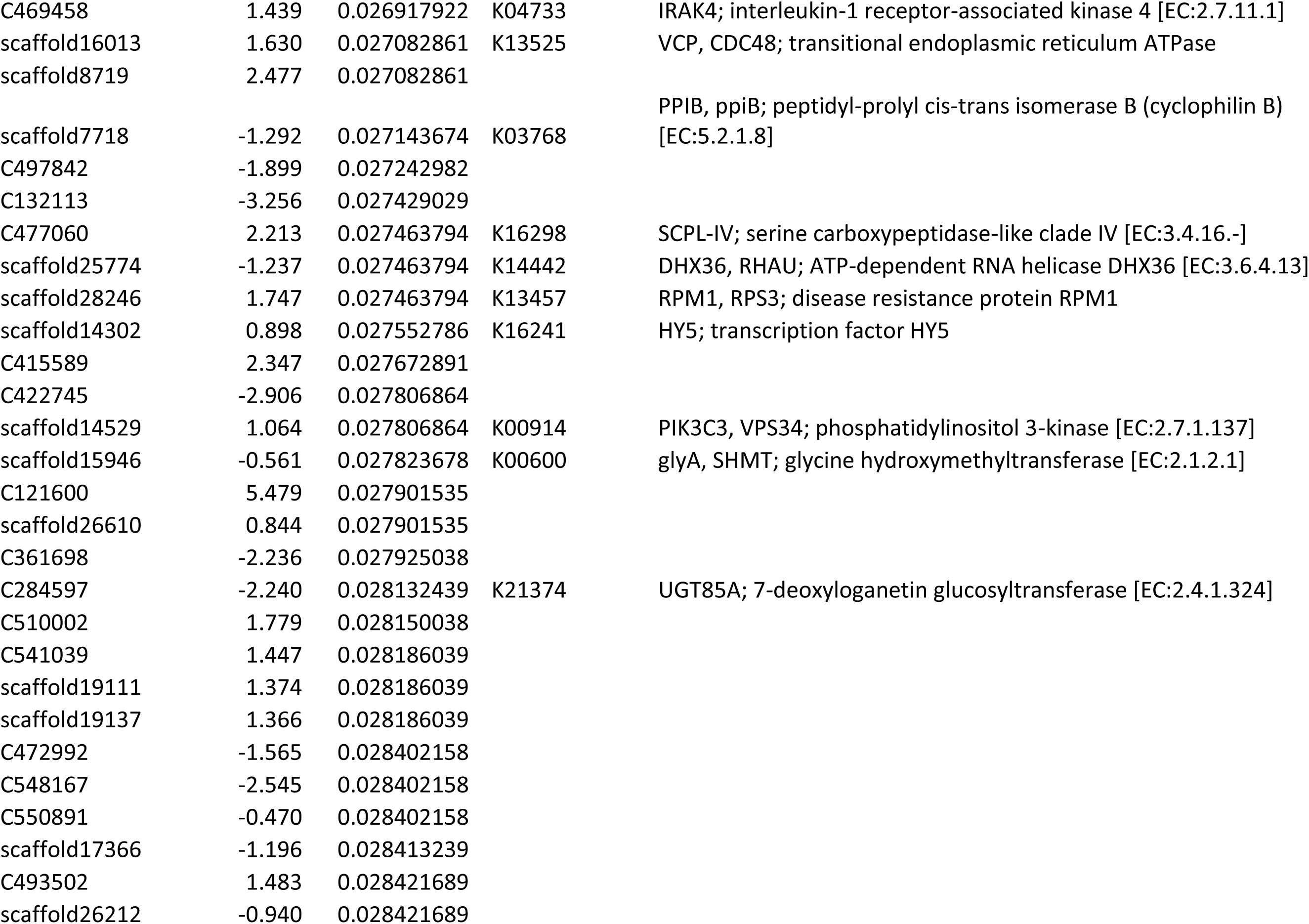
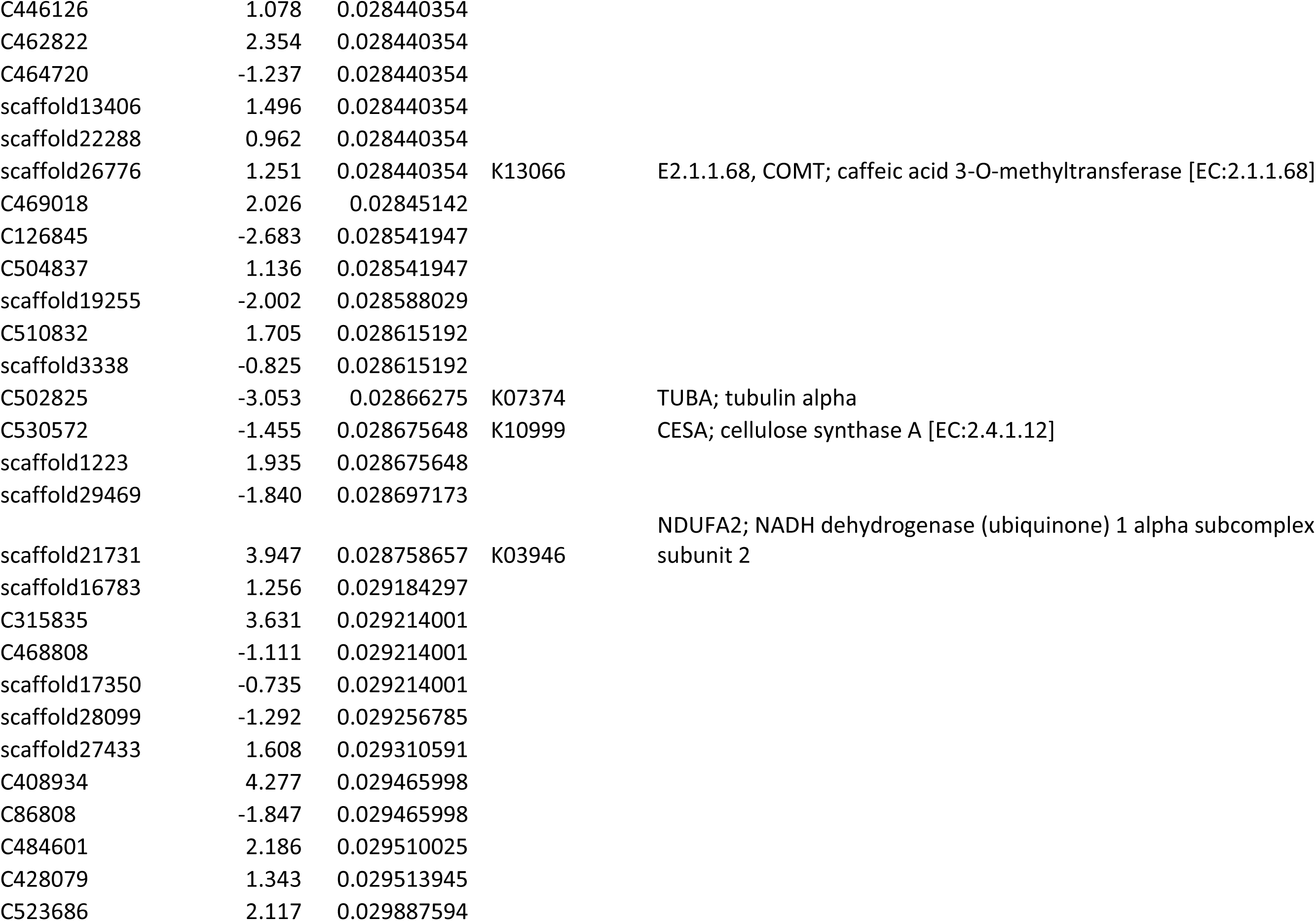
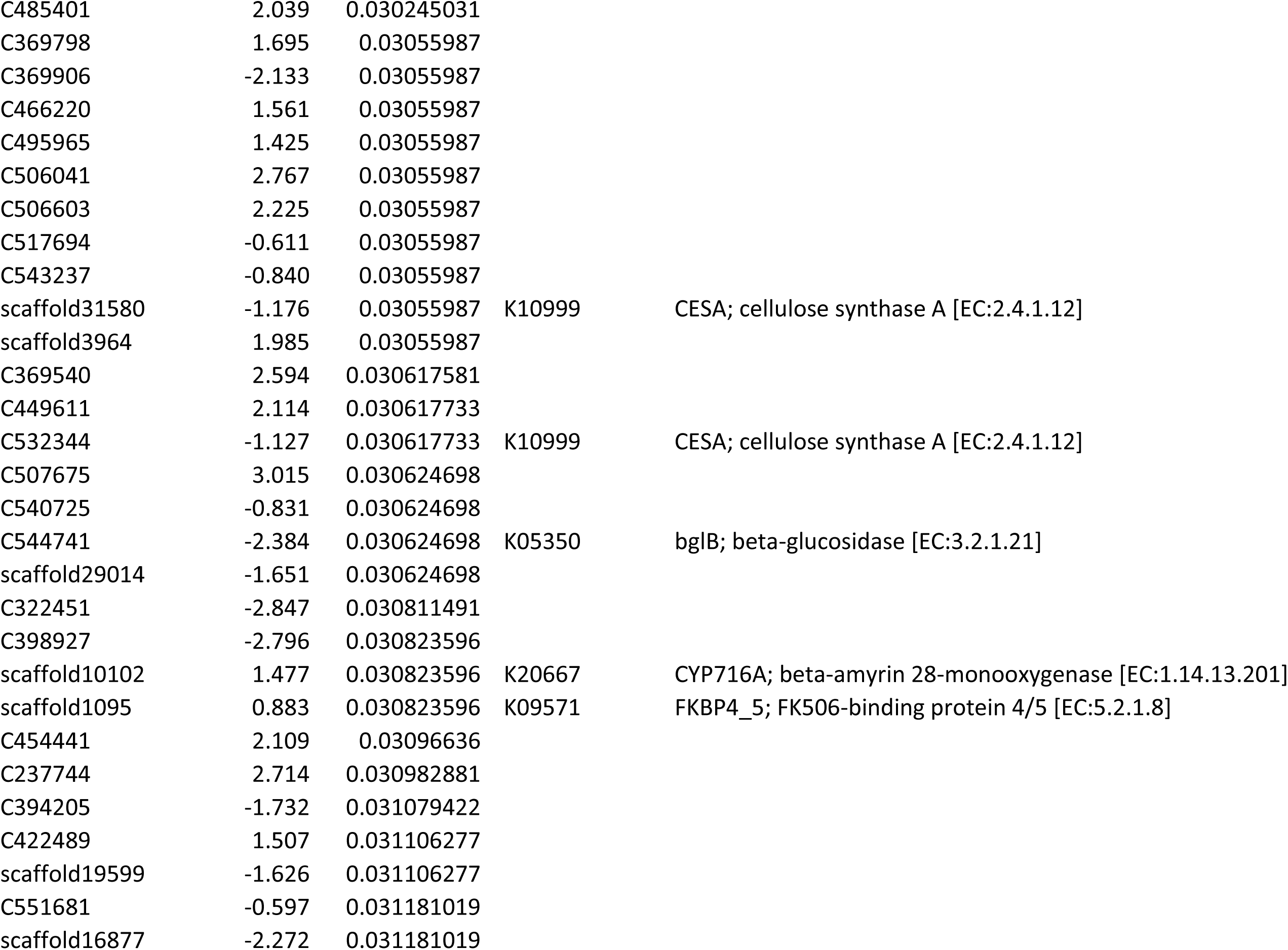
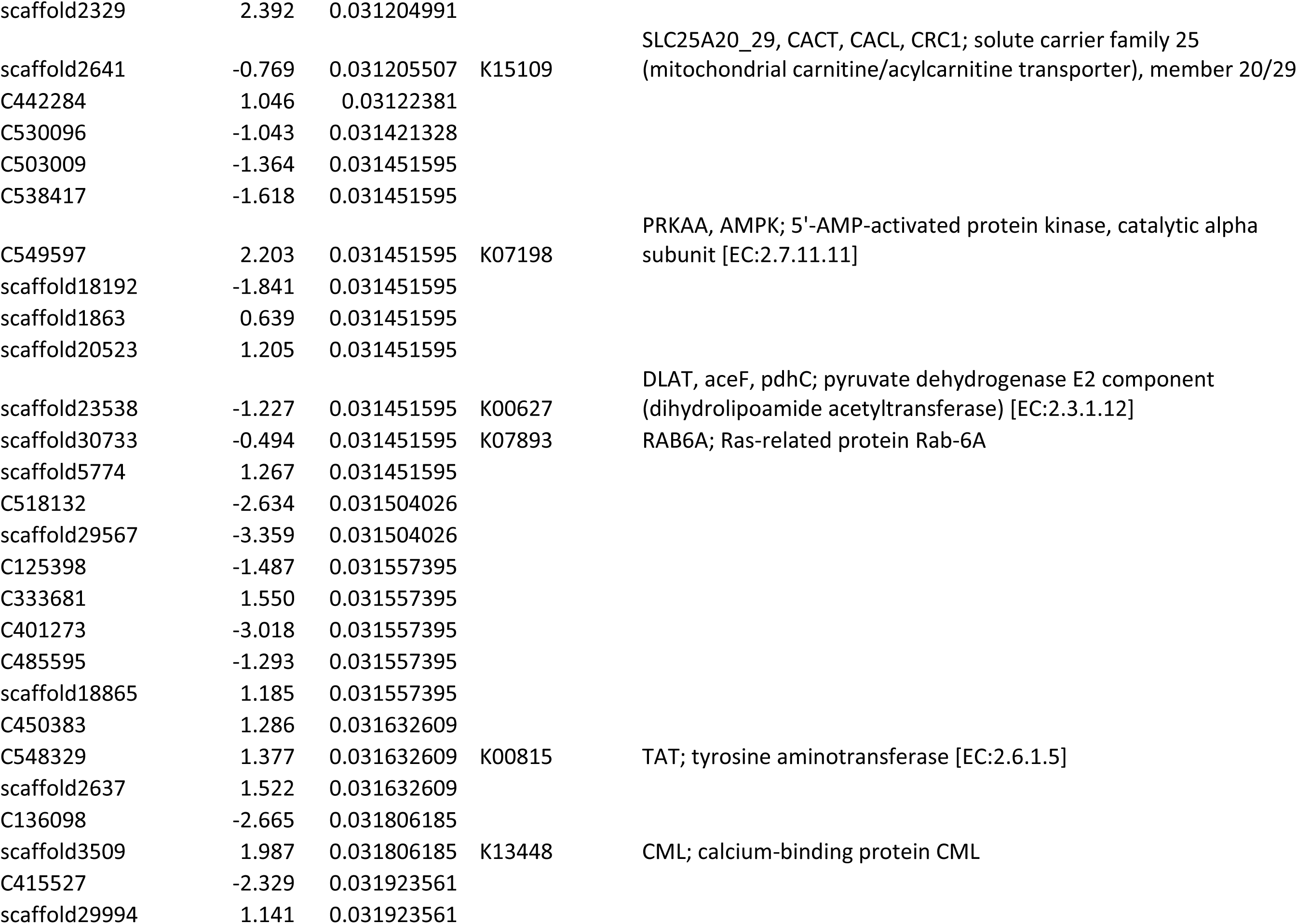
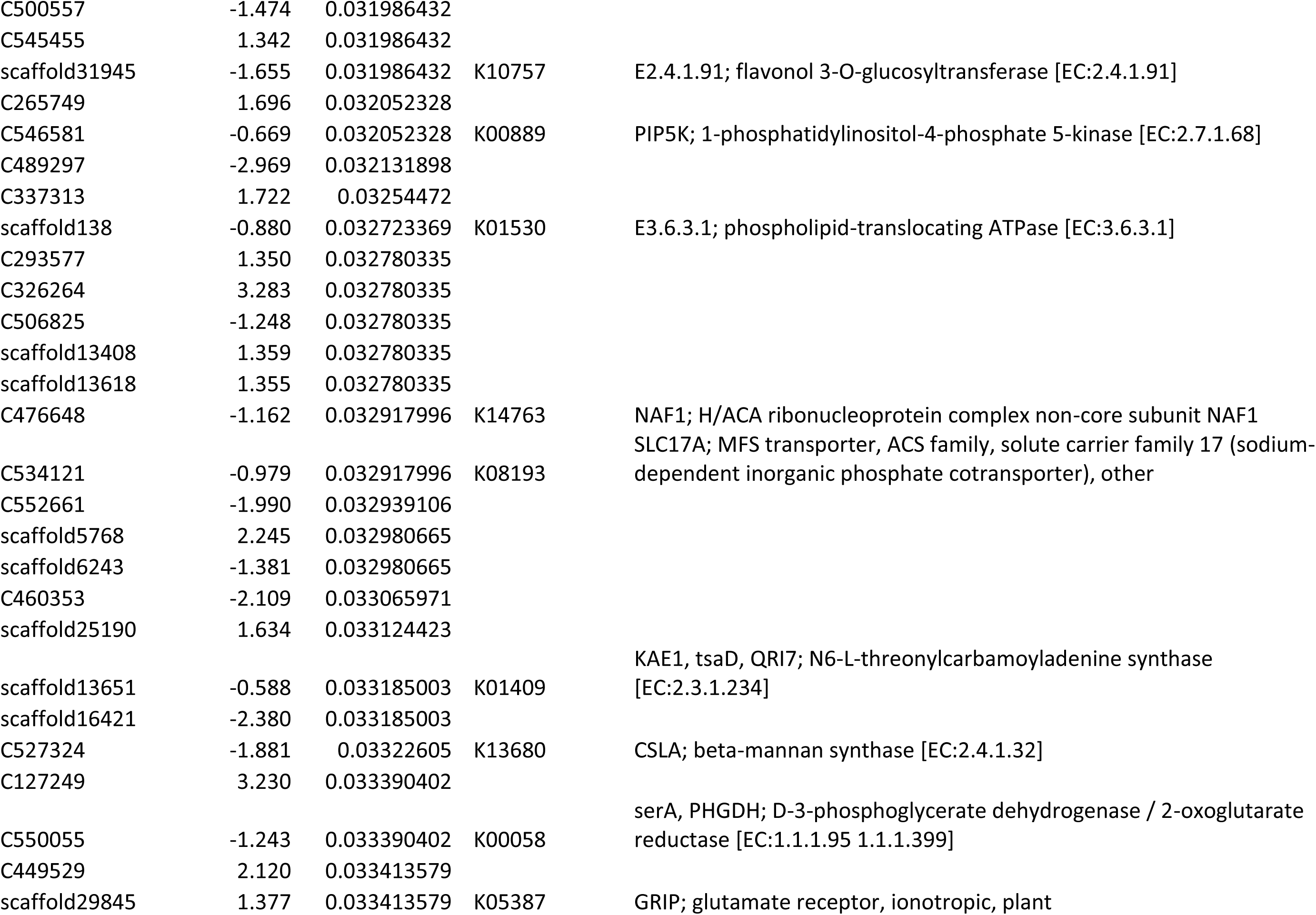
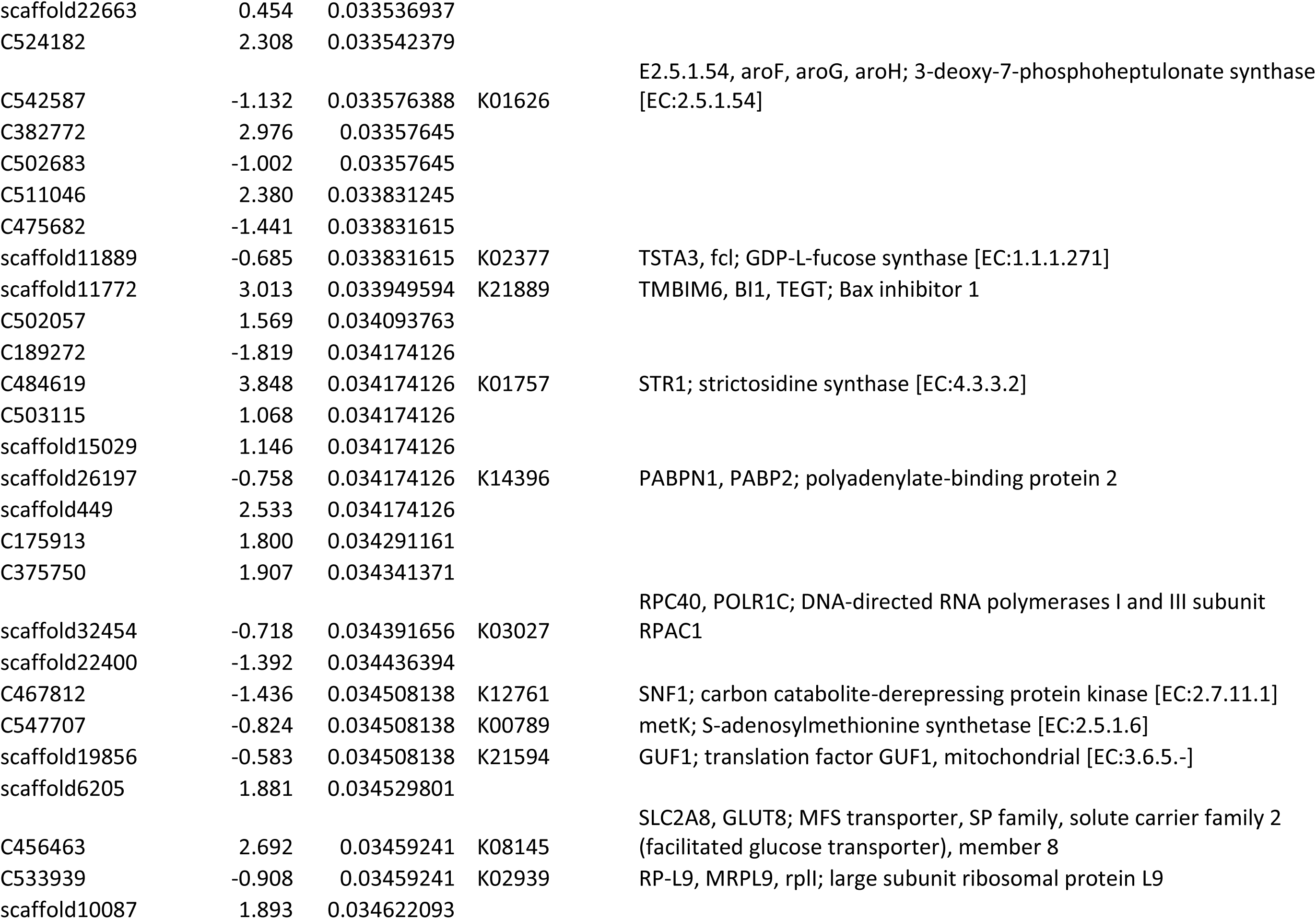
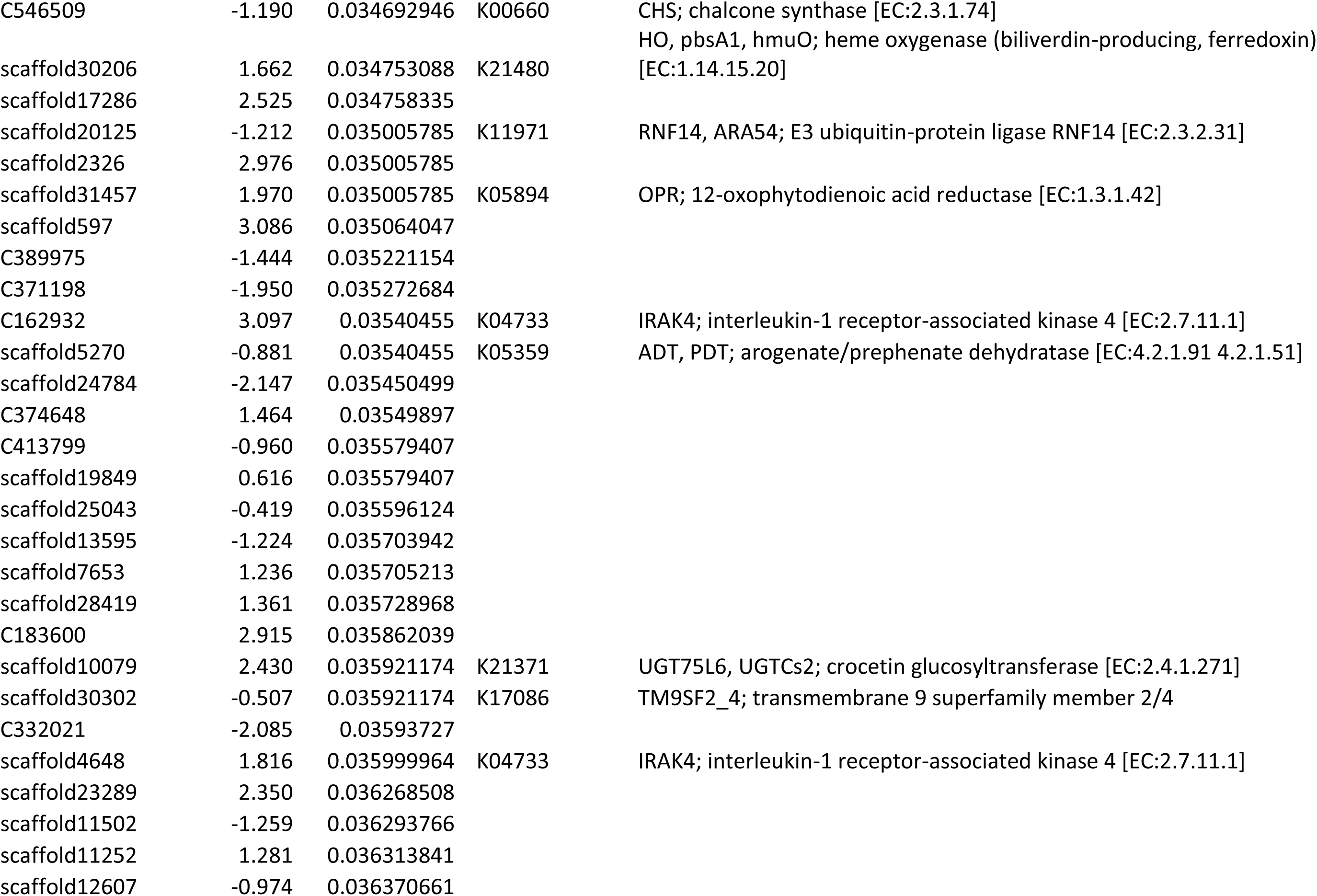
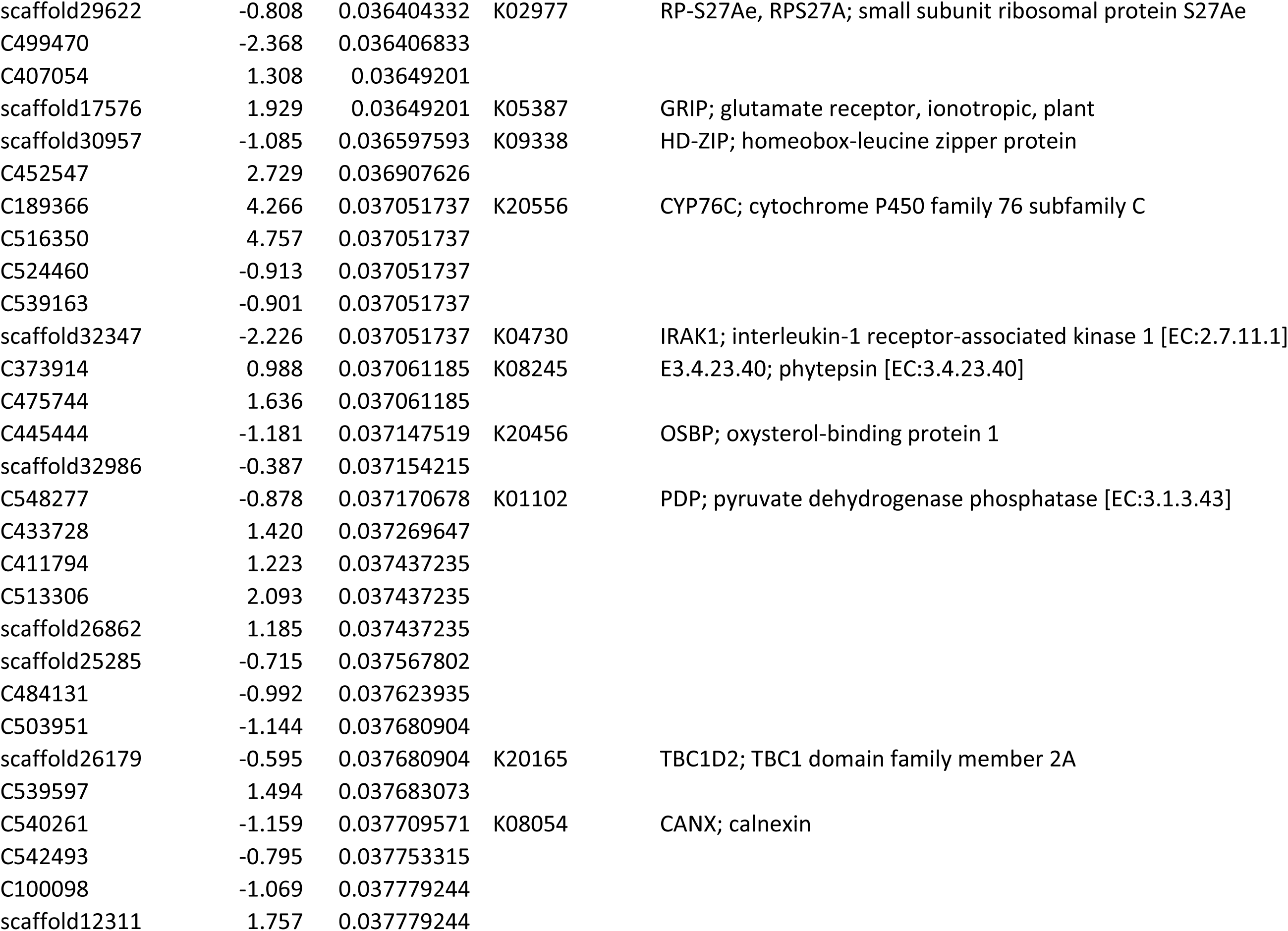
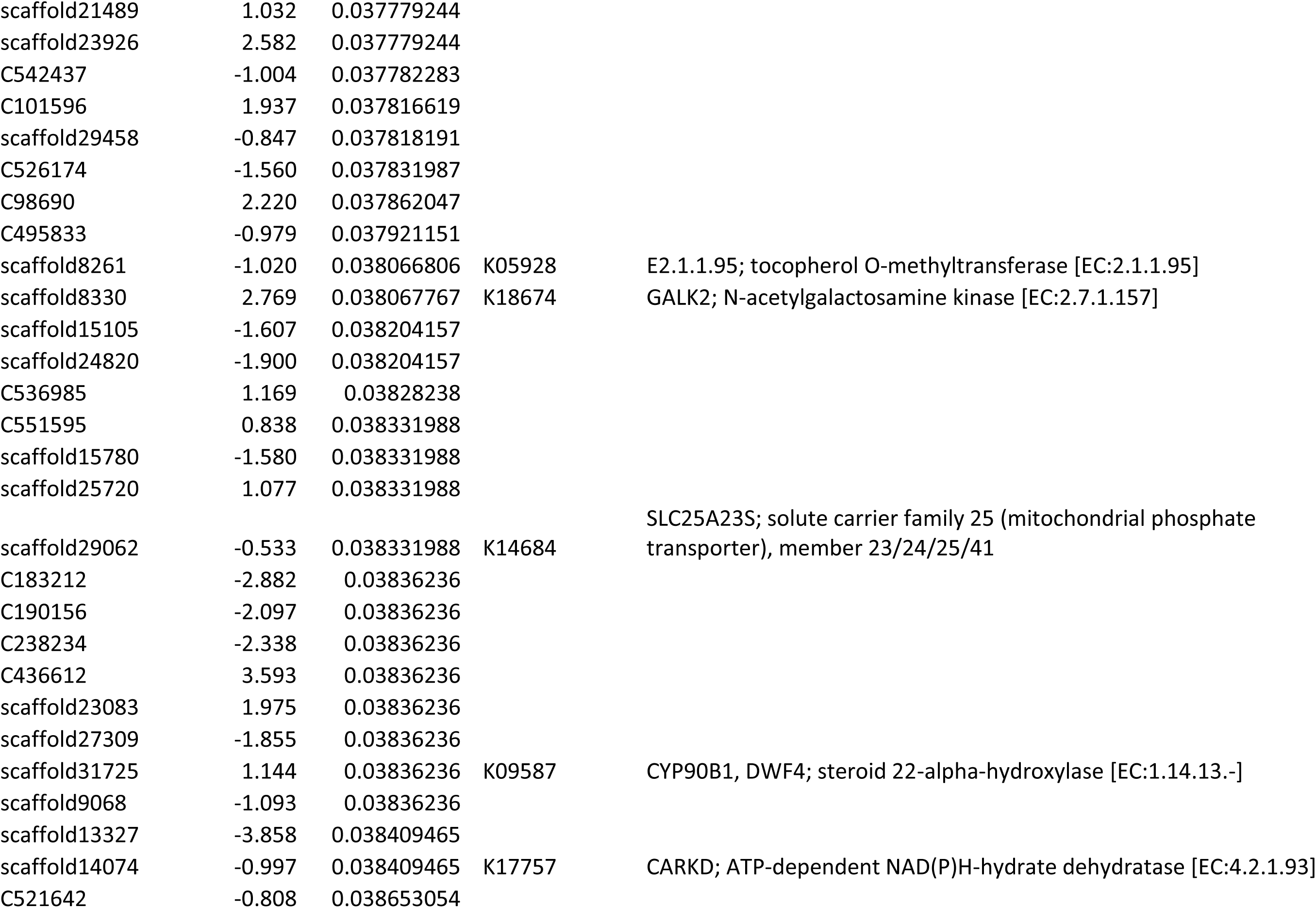
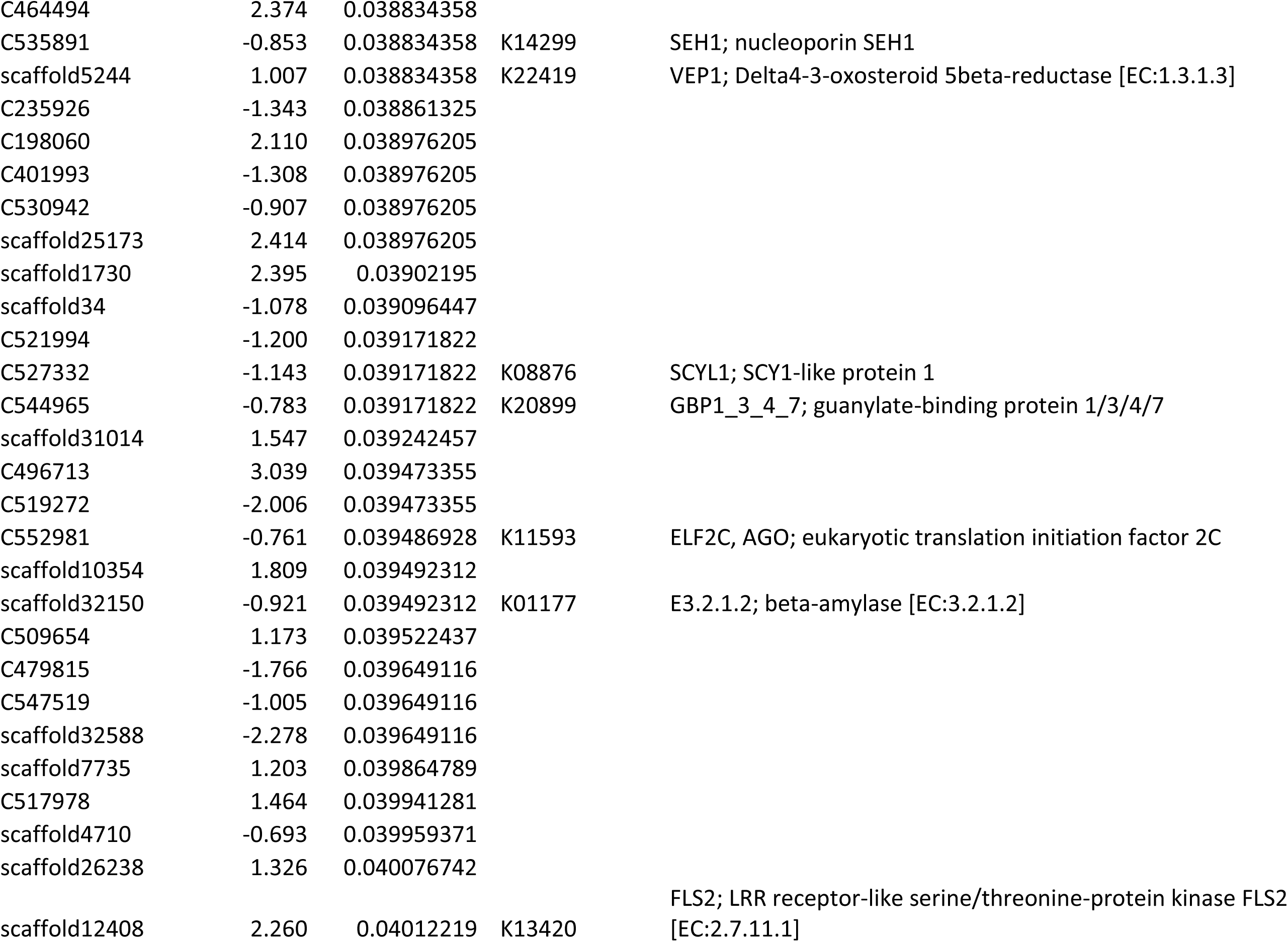
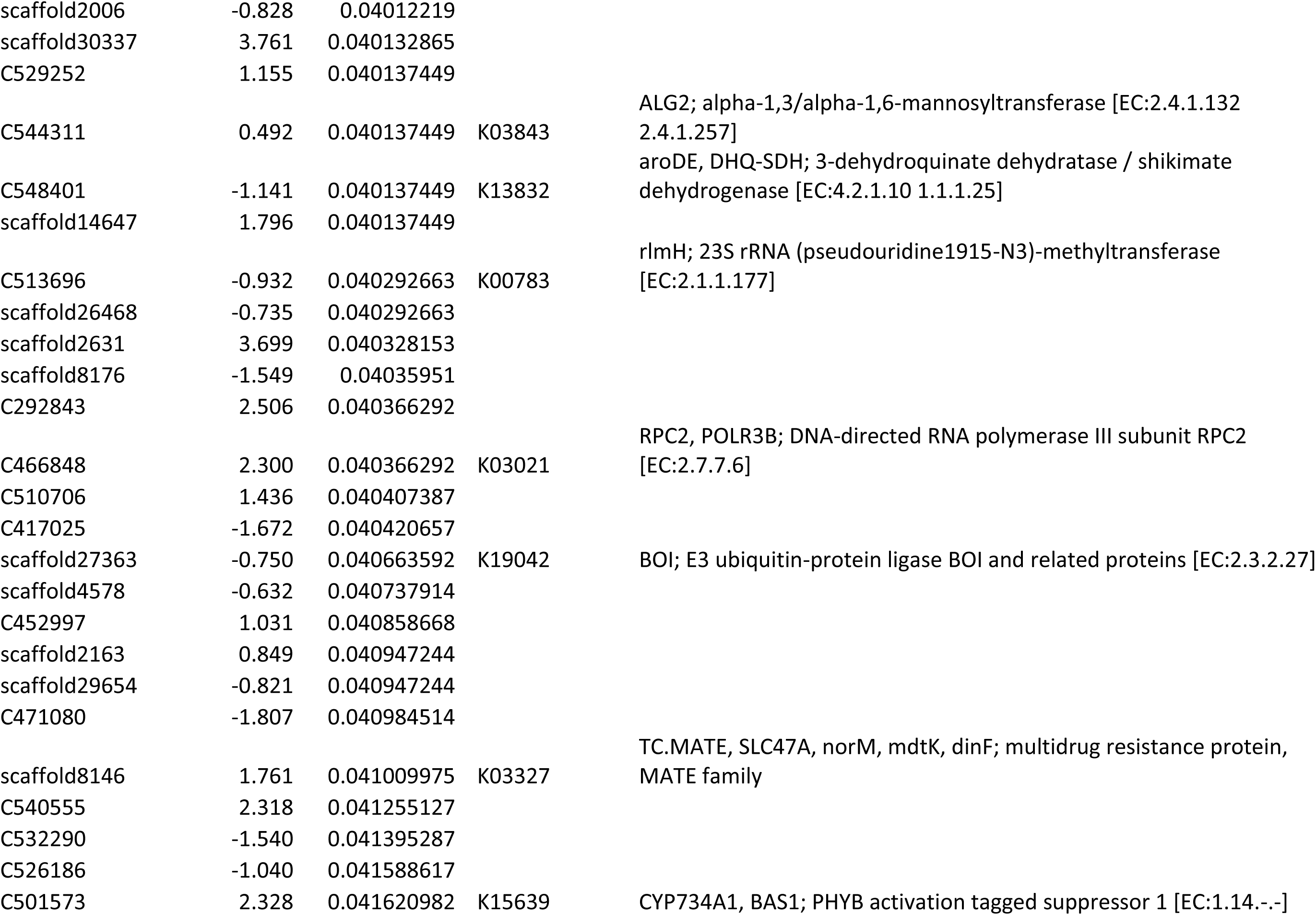
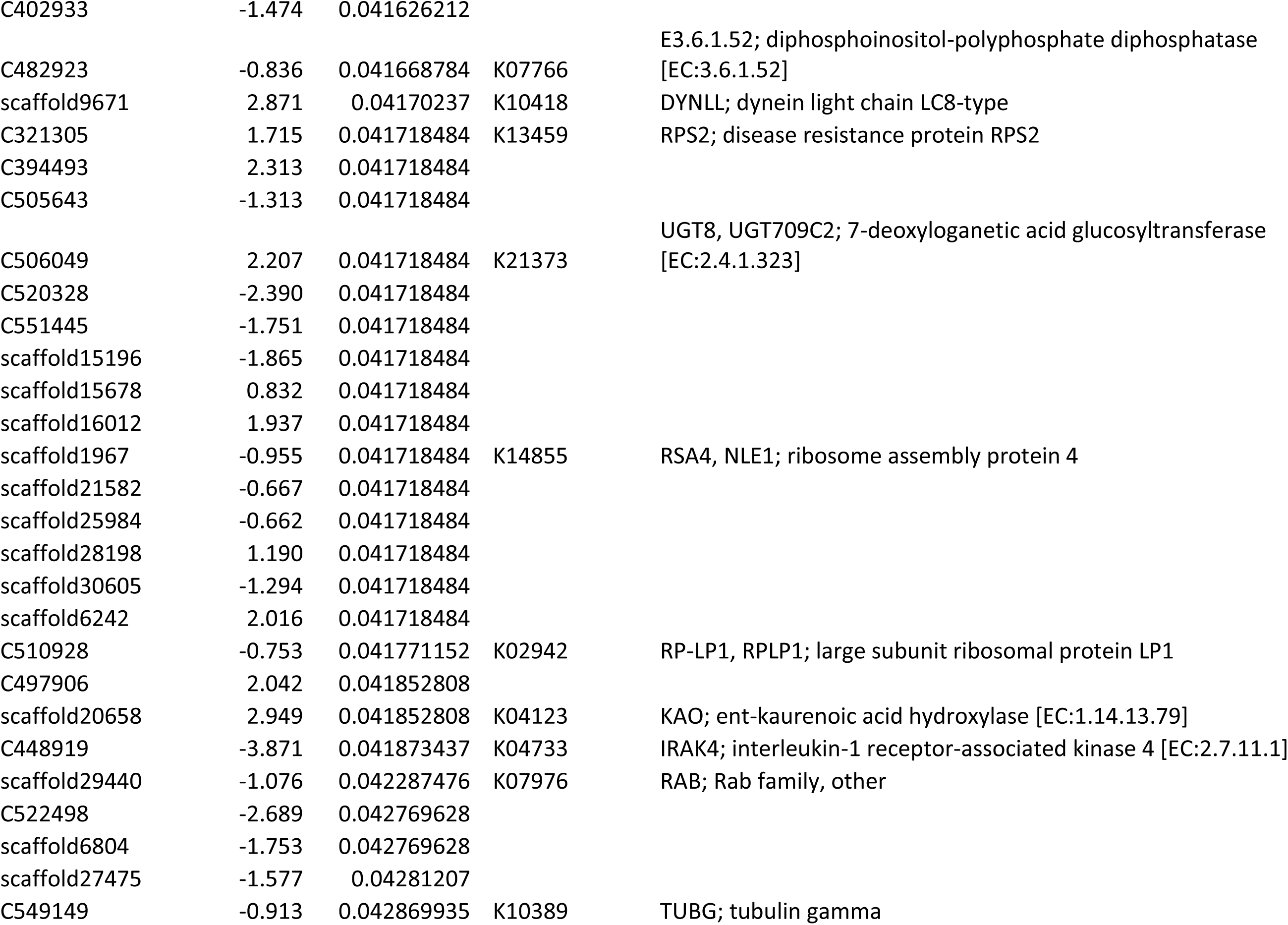
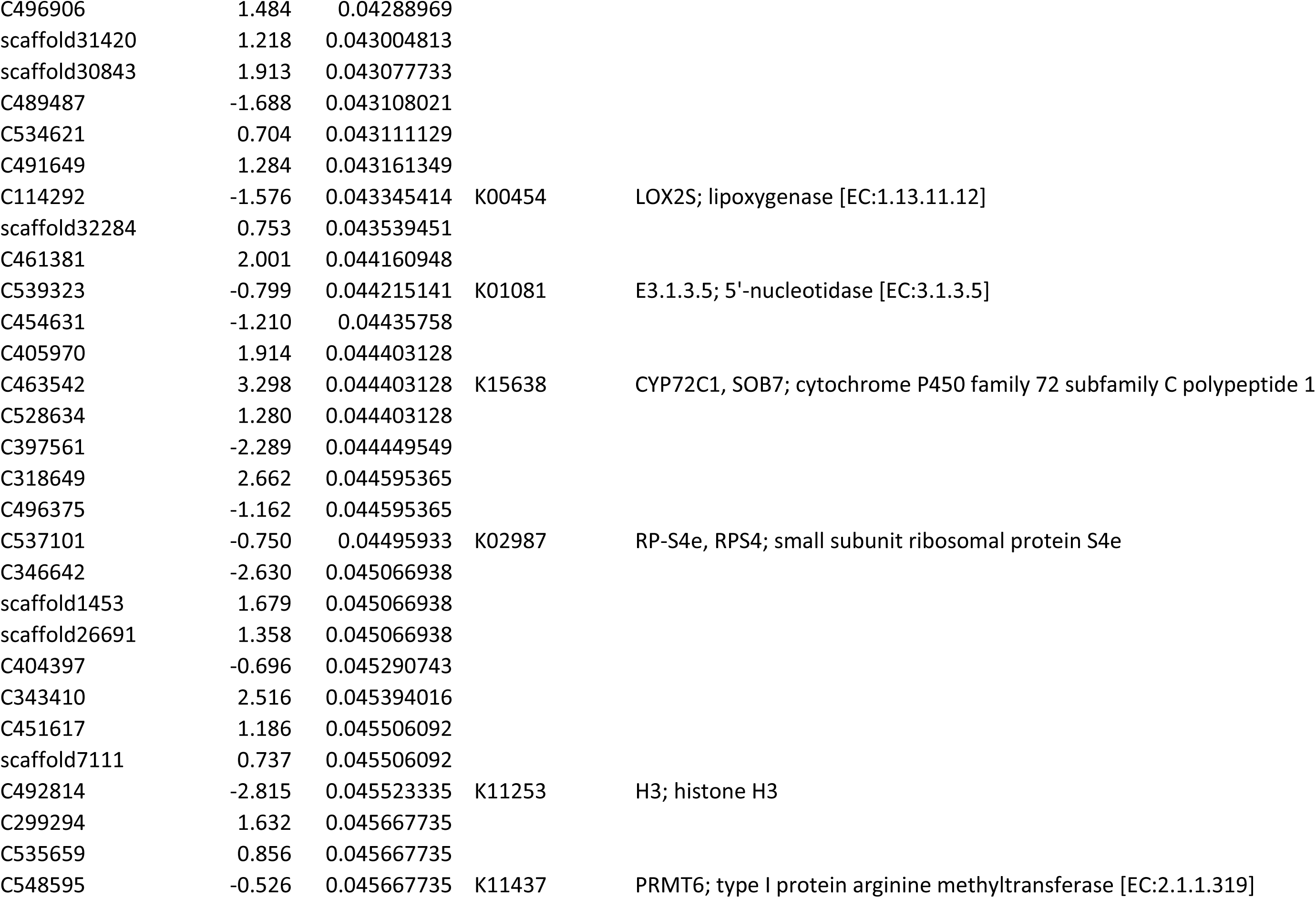
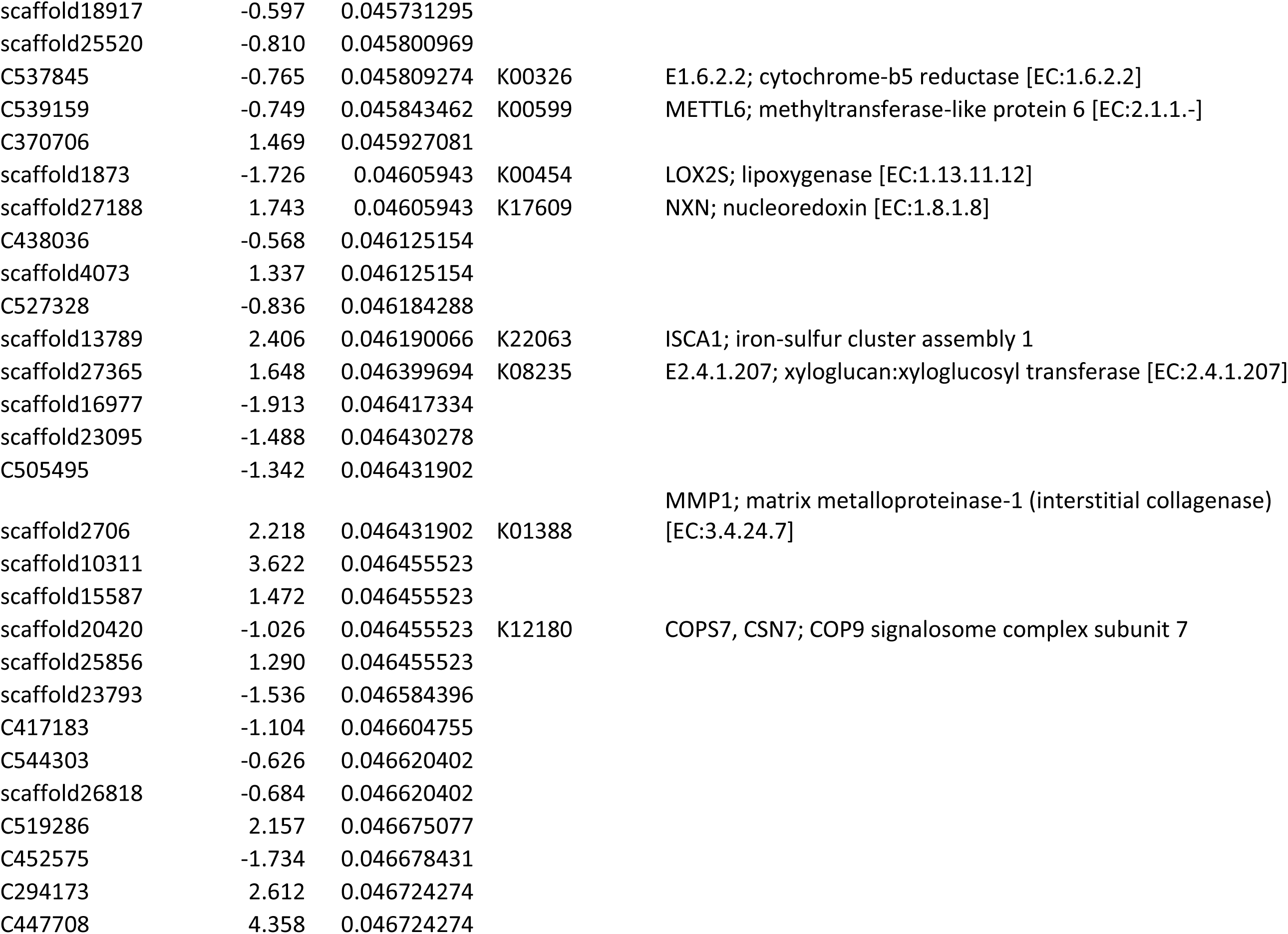
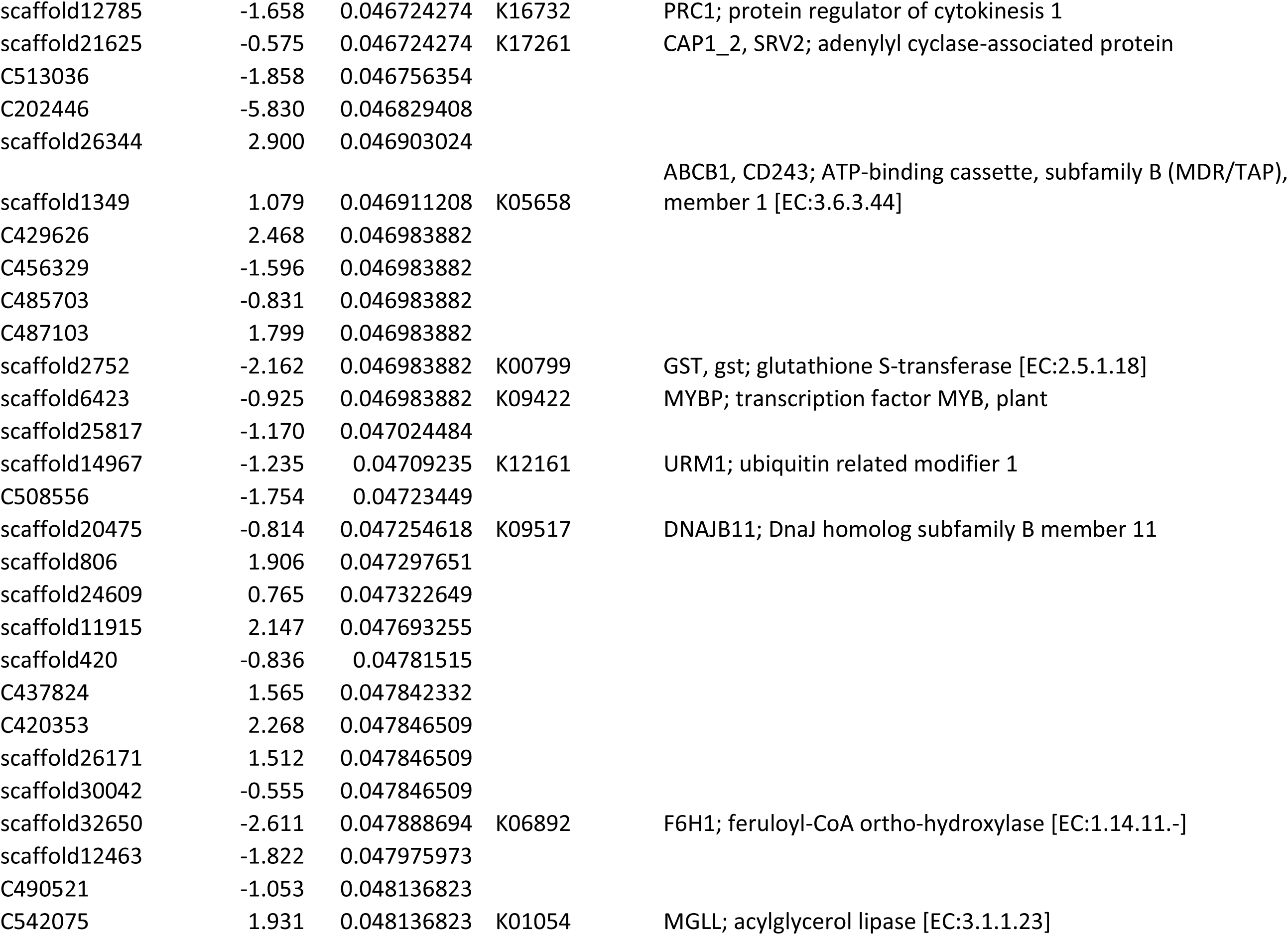
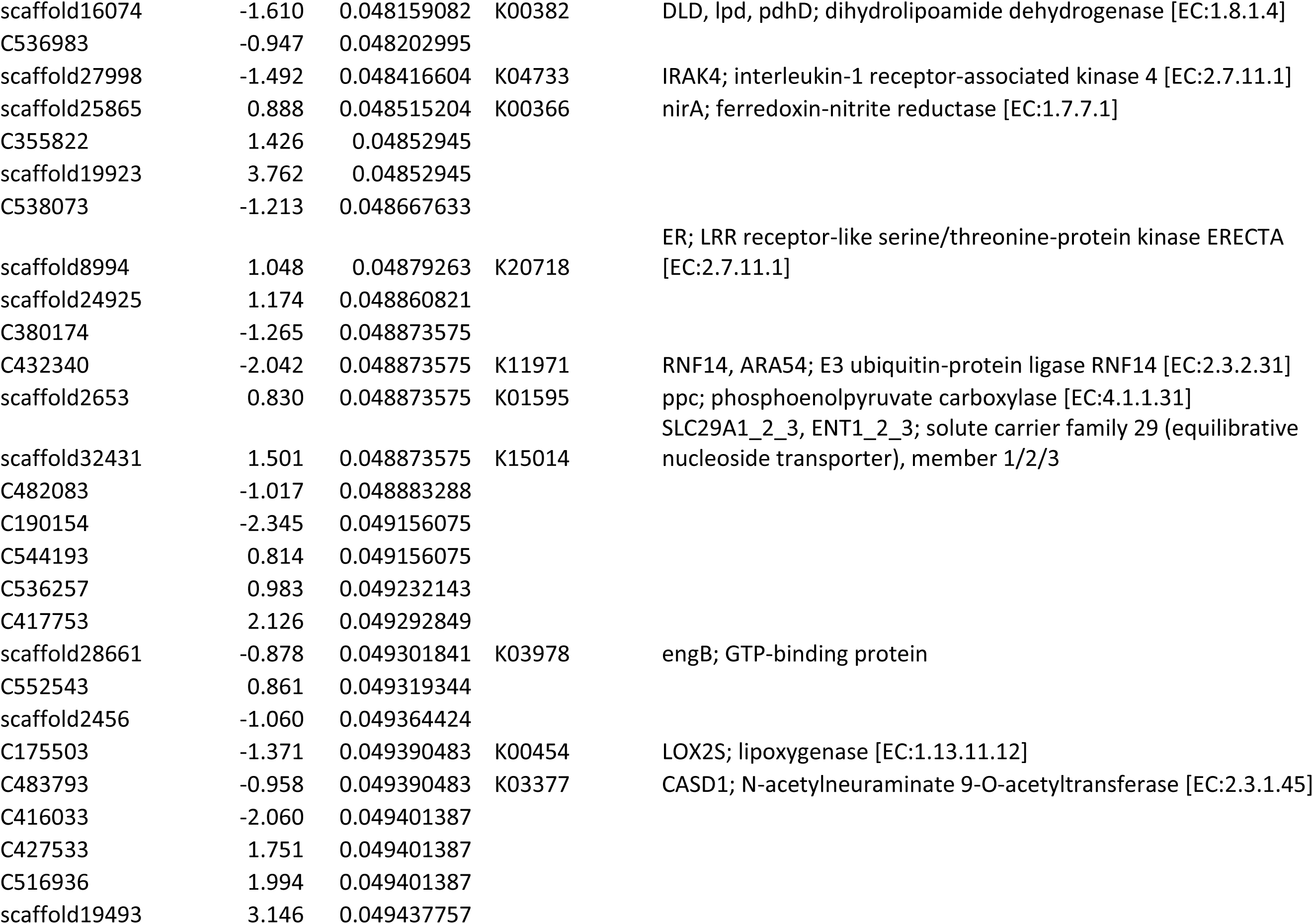
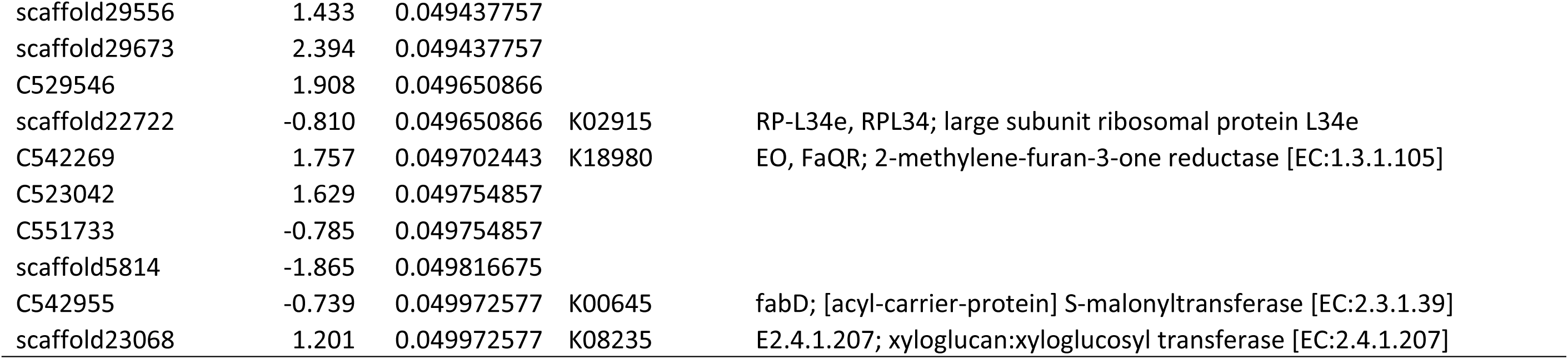
Differentially expressed genes (DEGs) identified between light green- and pink-bracted cultivars of C. kousa. Includes transcript ID, log2 fold change, adjusted p-value, KEGG ortholog, and functional annotation.

**Supplementary Table T7.**
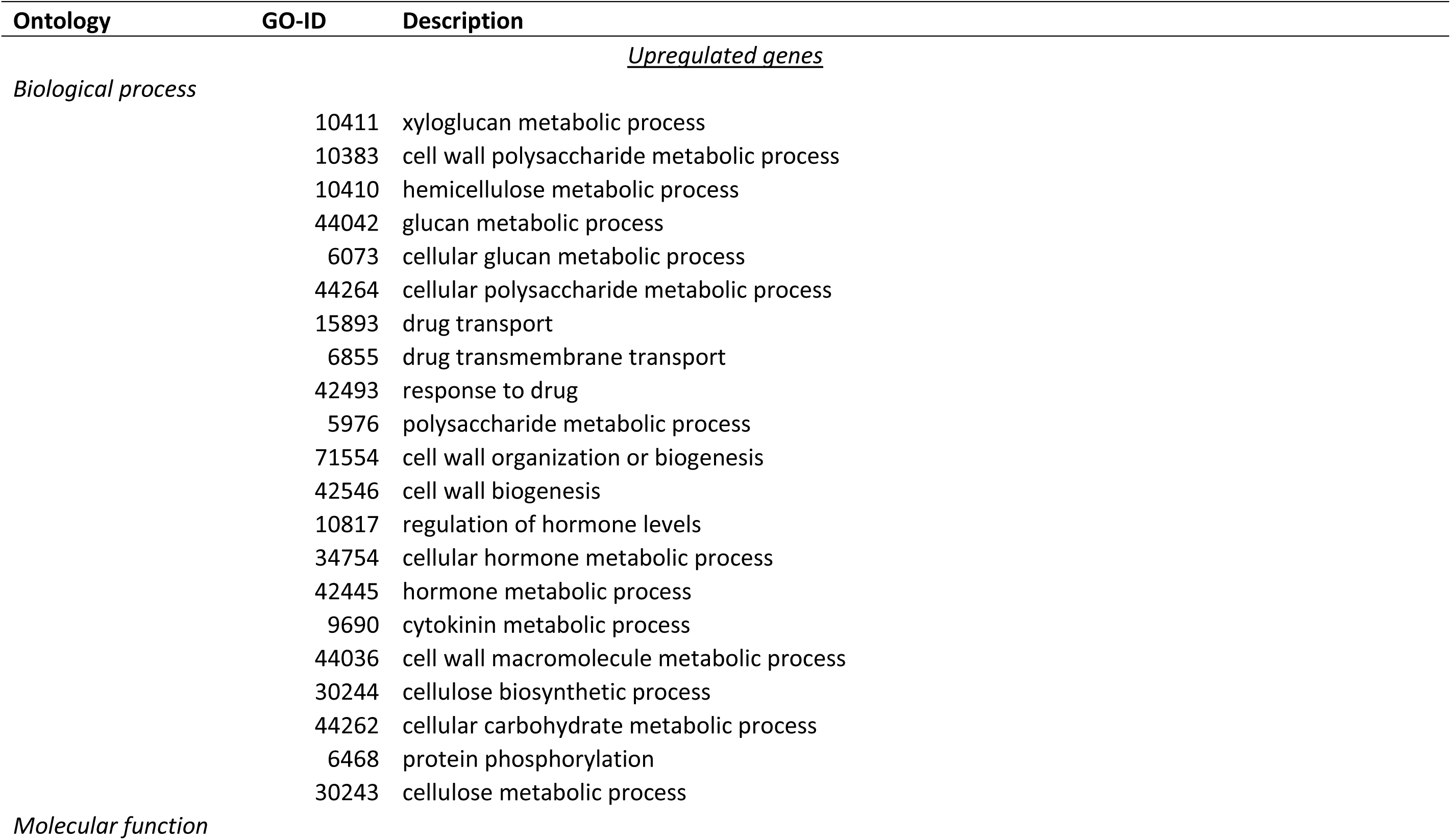
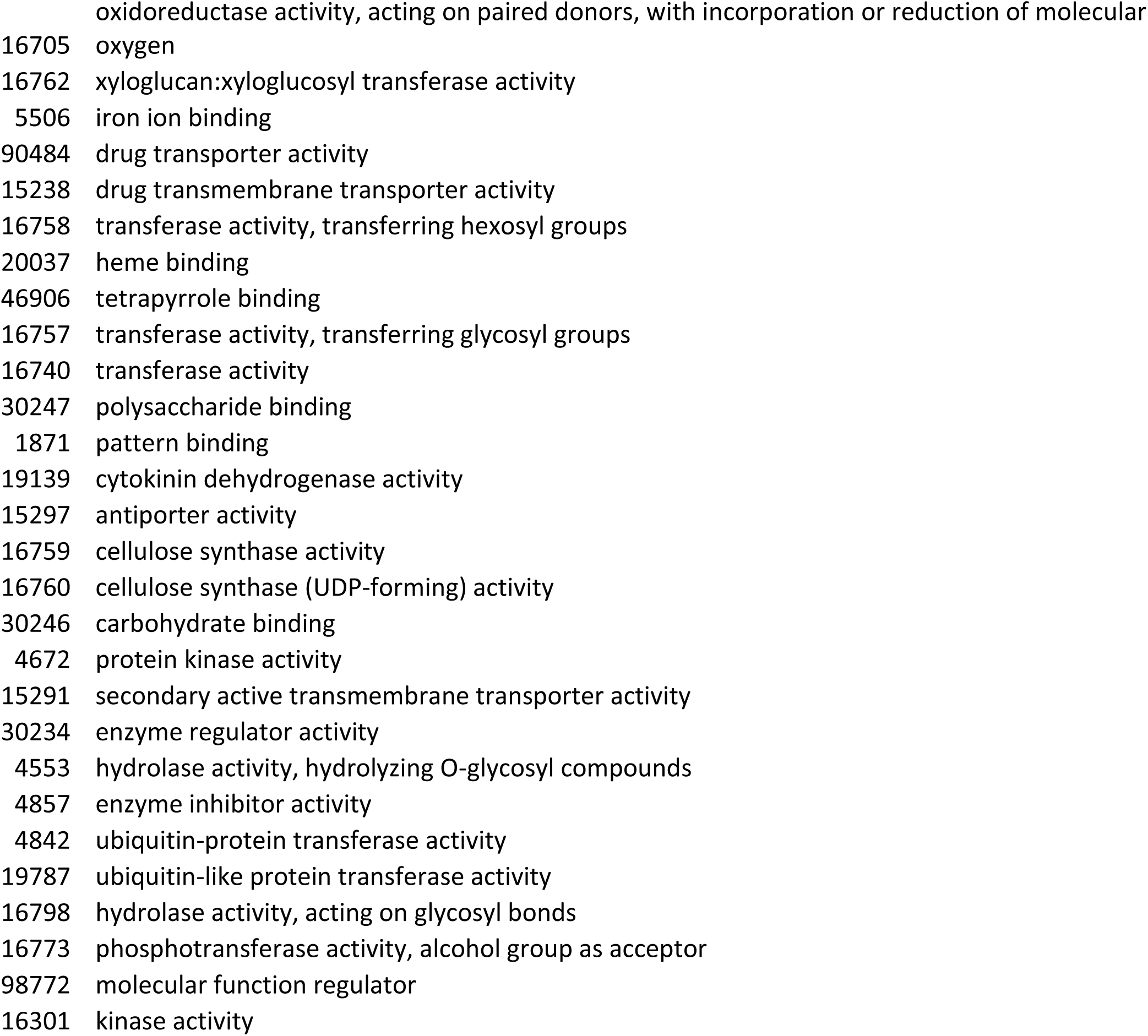
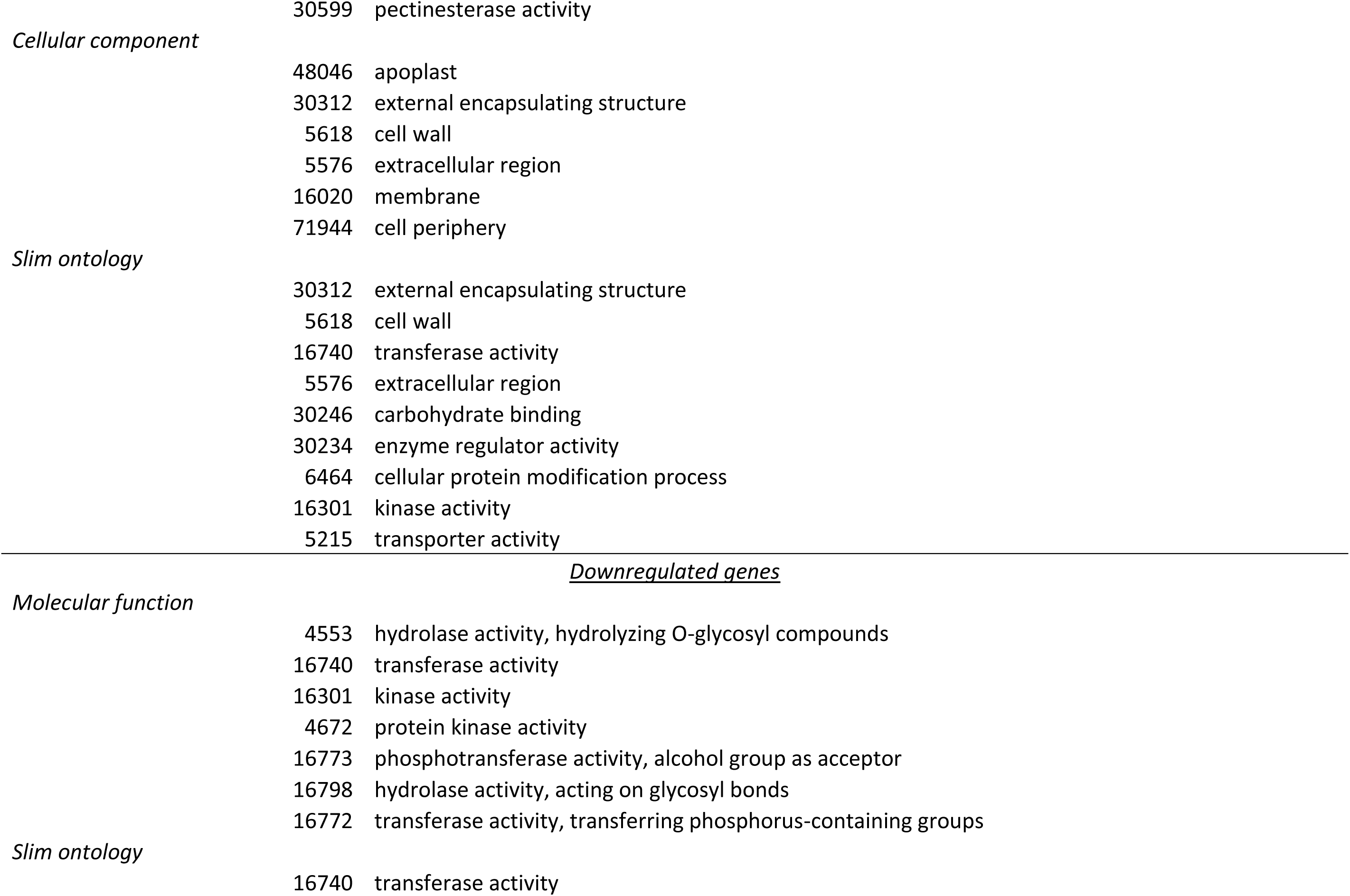
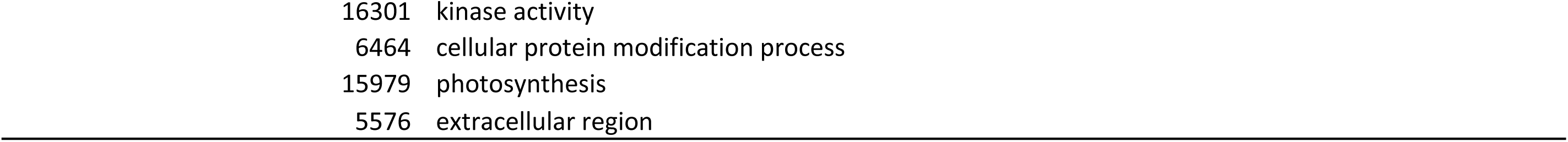
Gene Ontology (GO) terms significantly enriched among DEGs upregulated in C. florida, identified using BiNGO plugin in Cytoscape.

**Supplementary Table T8.**
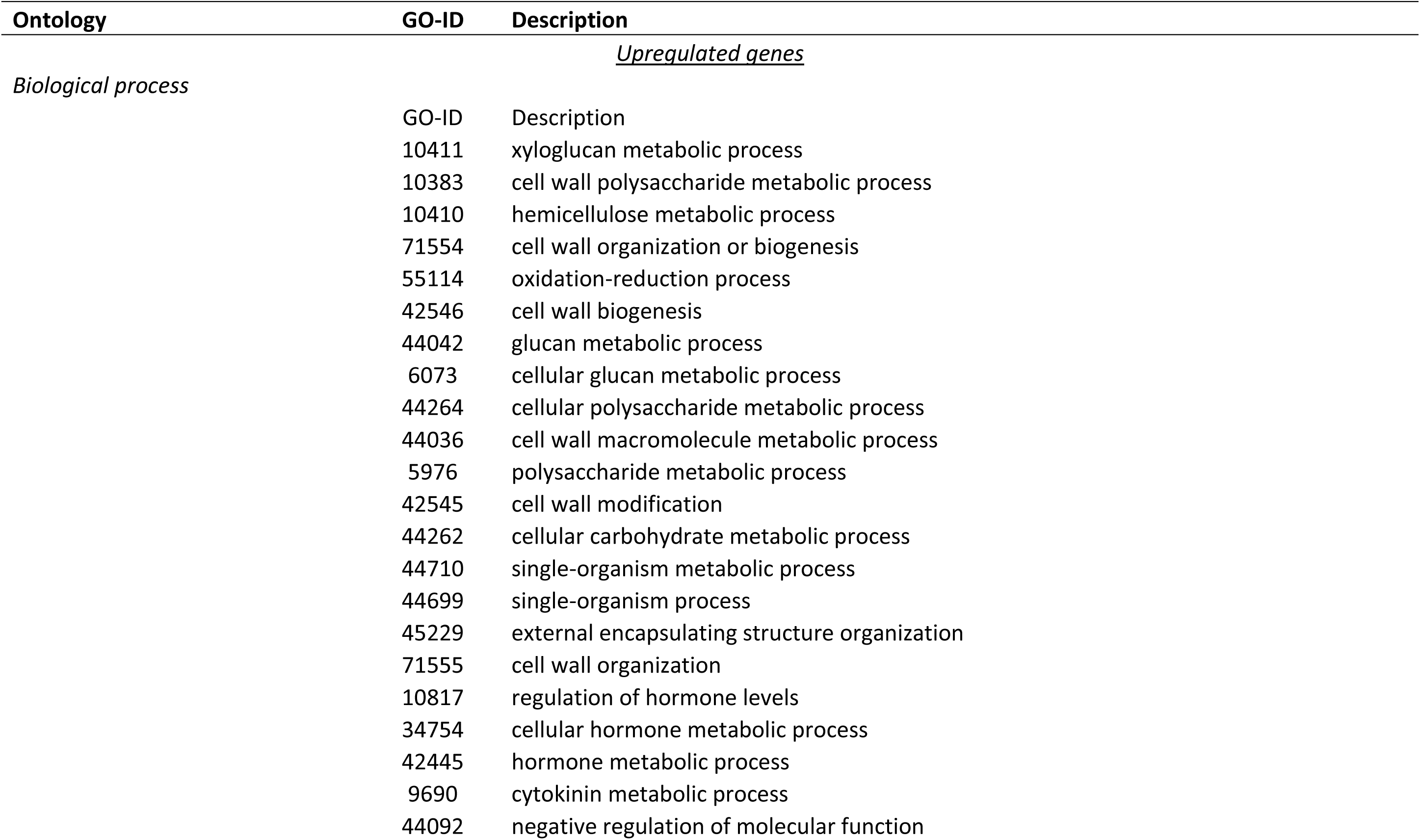
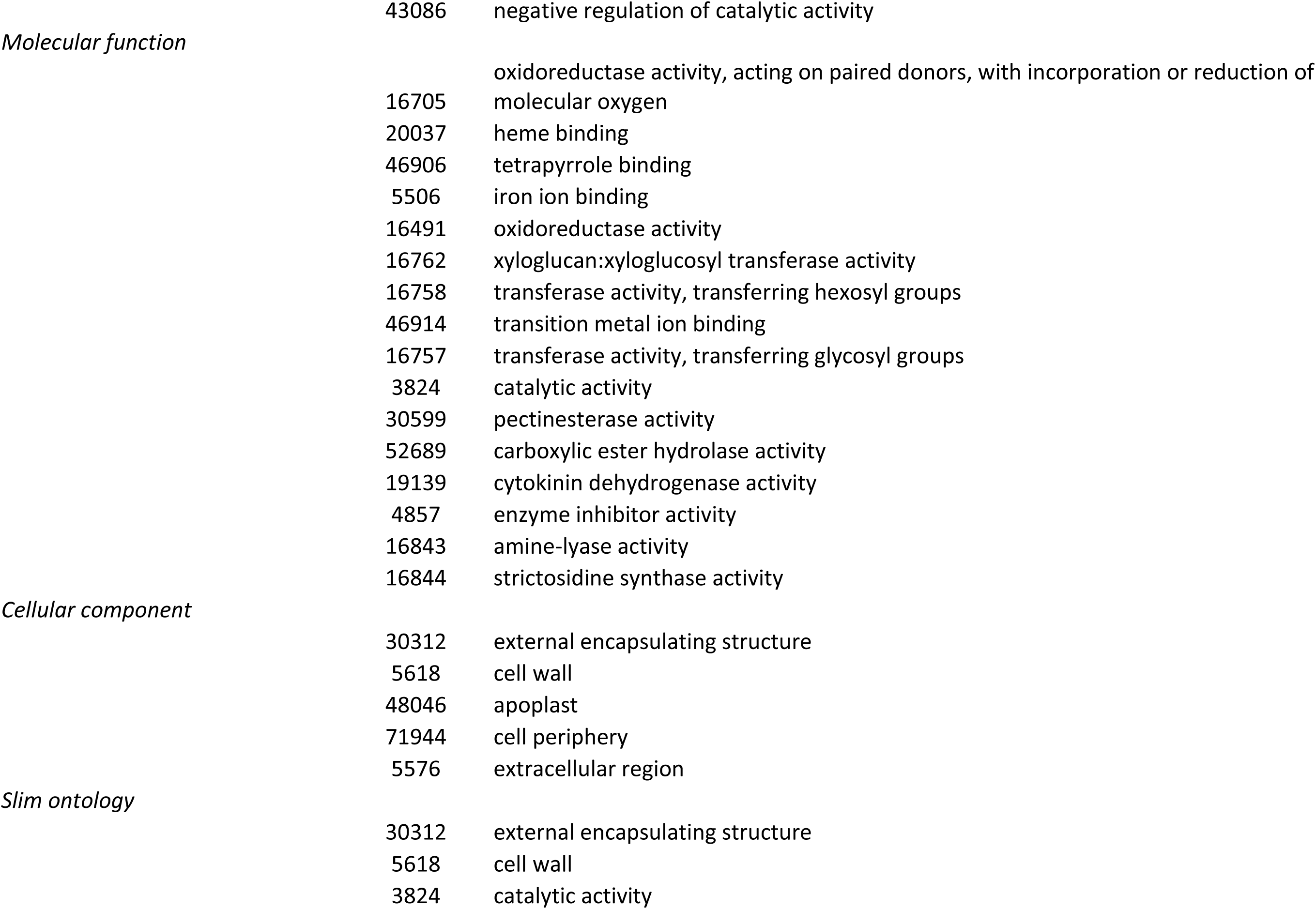
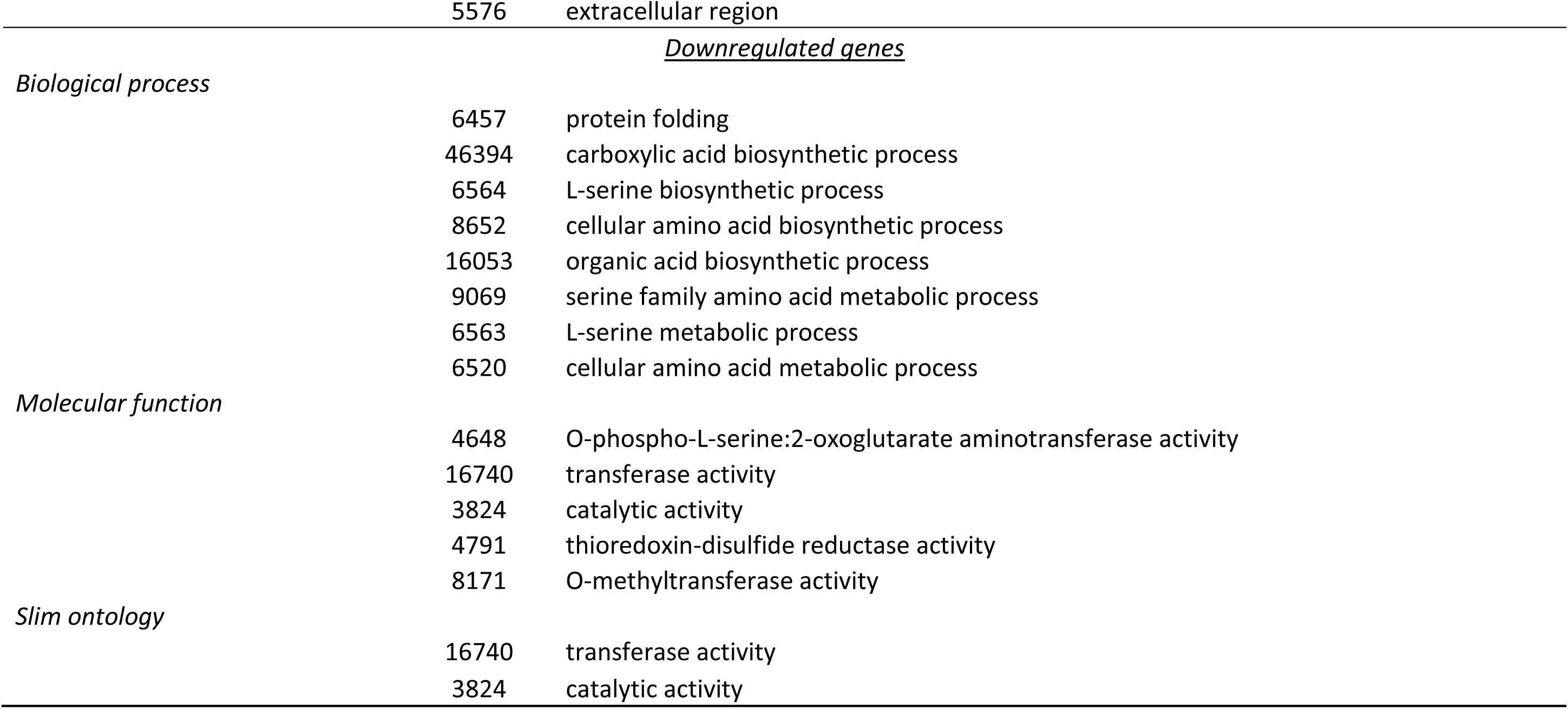
Gene Ontology (GO) terms significantly enriched among DEGs upregulated in C. kousa, identified using BiNGO plugin in Cytoscape.

**Supplementary Table T9.**
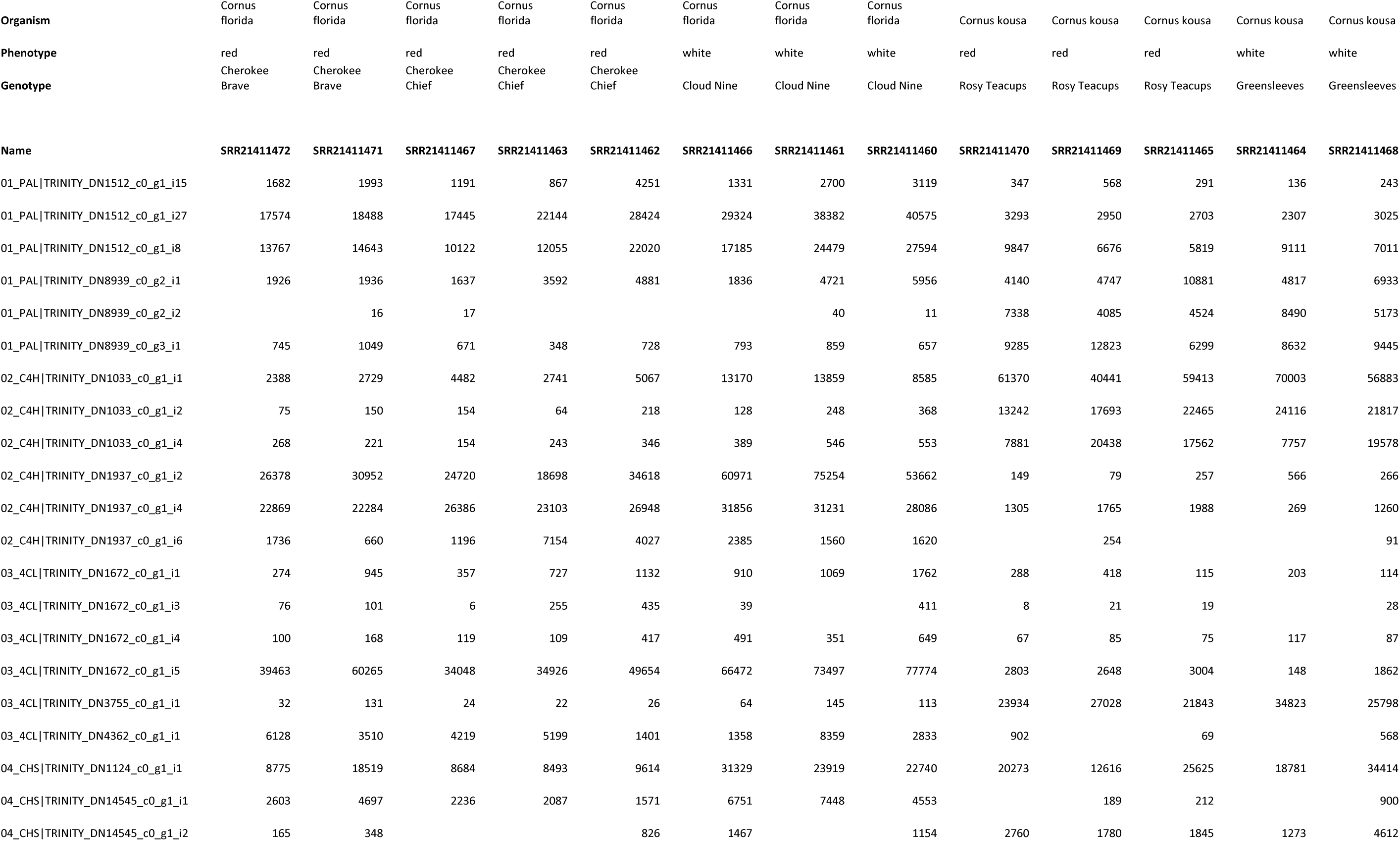
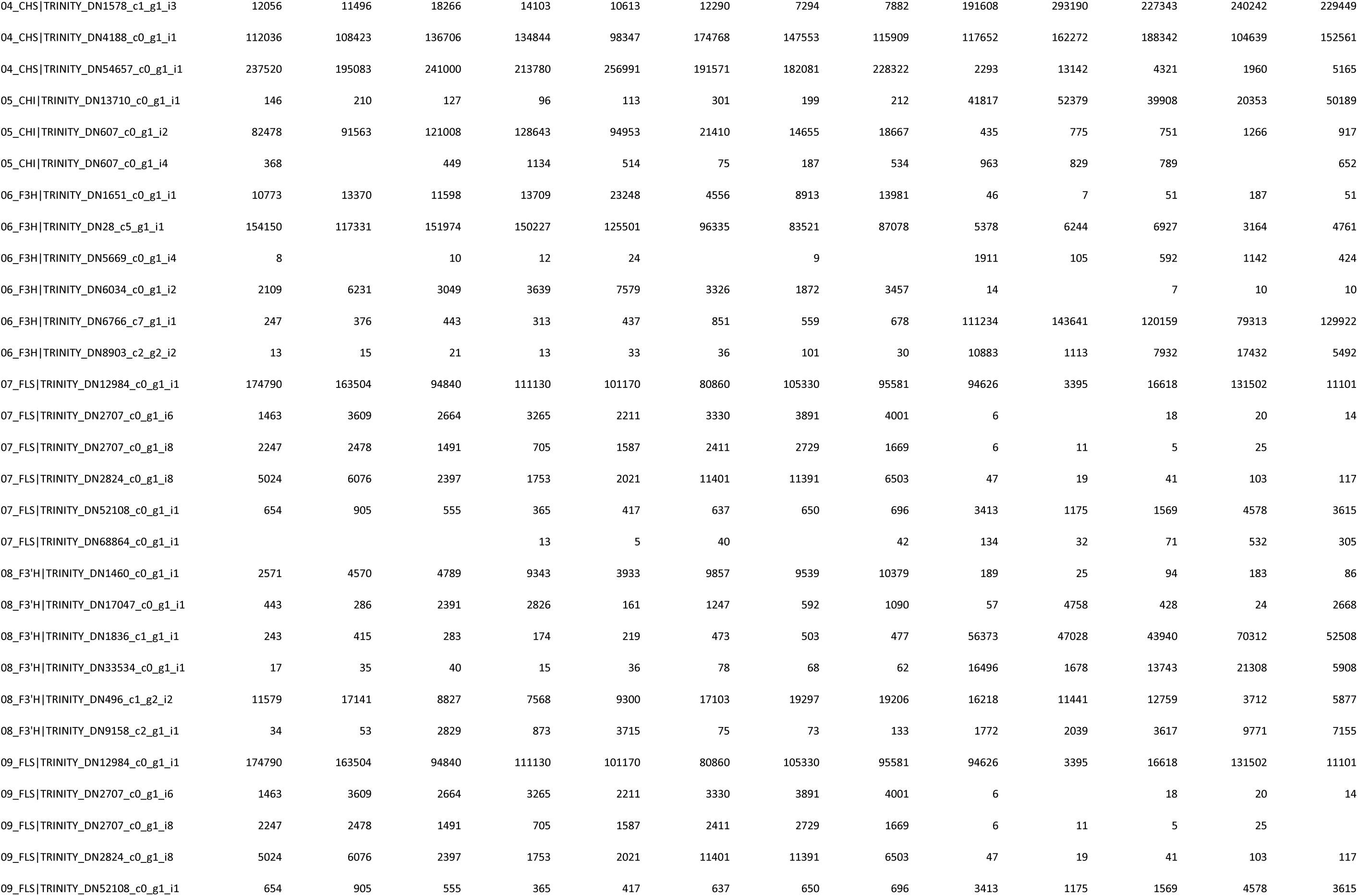
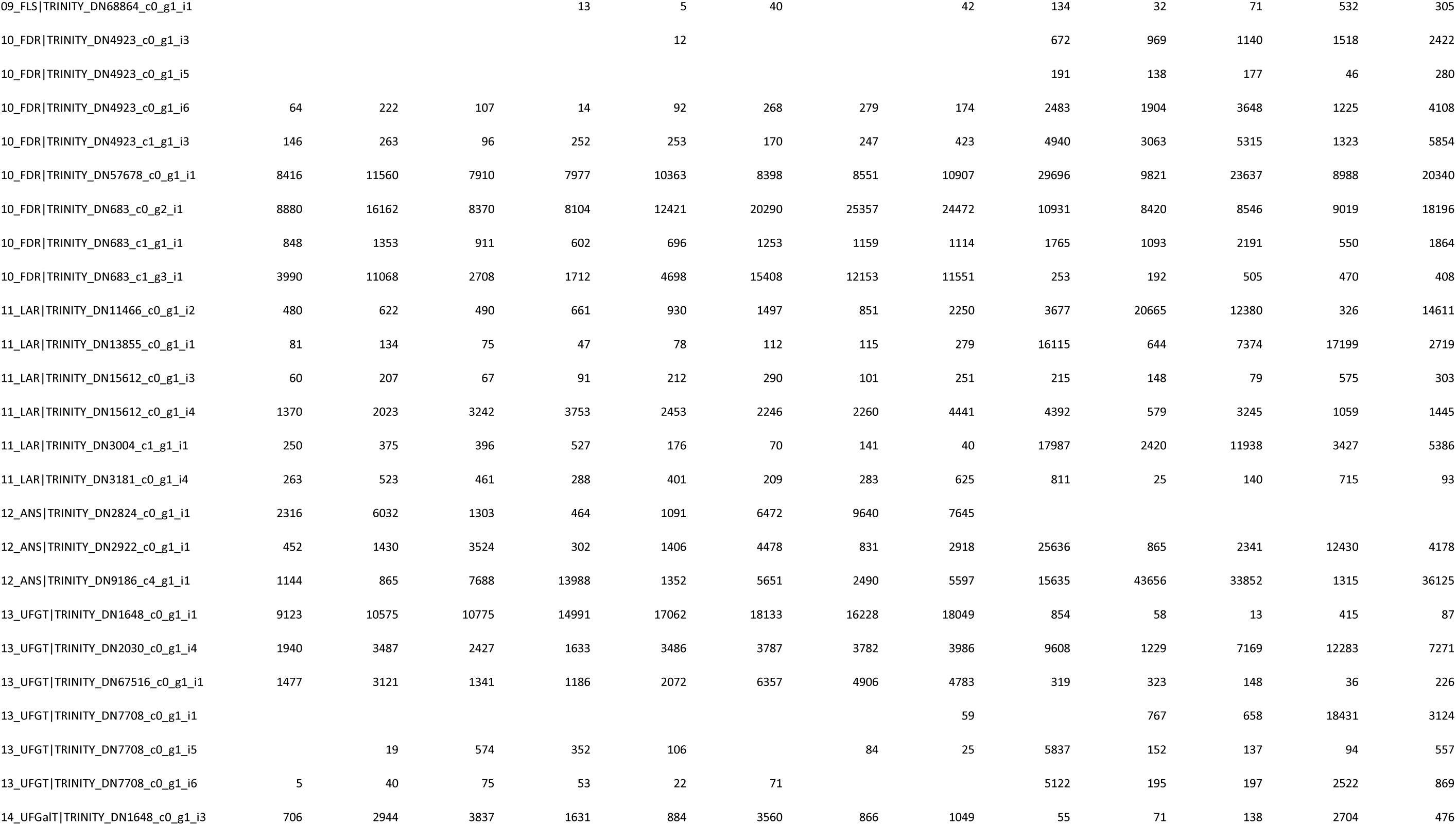
Normalized expression matrix of 14 phenylpropanoid pathway genes across 13 RNA-seq samples from five Cornus cultivars. Includes gene IDs and expression values used for heatmap visualization (Figure 5).

**Supplementary Table T10.**
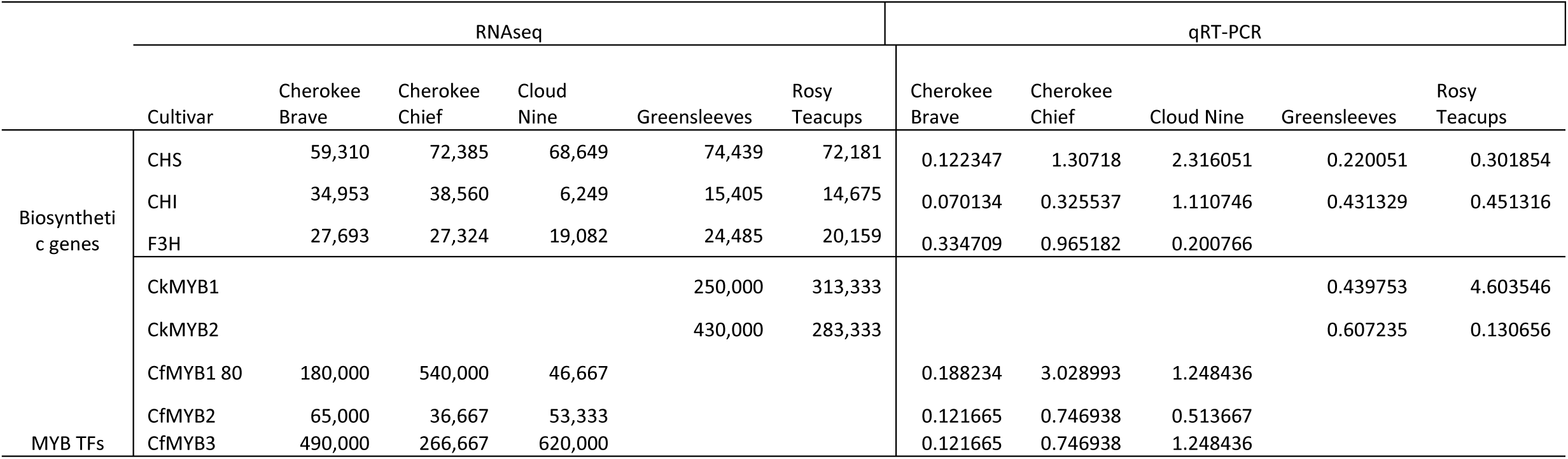
Summary of gene expression patterns and regulatory roles of candidate MYB transcription factors and structural genes used in the regulatory network diagram (Figure 6).

